# Modeling Regulatory Network Topology Improves Genome-Wide Analyses of Complex Human Traits

**DOI:** 10.1101/2020.03.13.990010

**Authors:** Xiang Zhu, Zhana Duren, Wing Hung Wong

## Abstract

Genome-wide association studies (GWAS) have cataloged many significant associations between genetic variants and complex traits. However, most of these findings have unclear biological significance, because they often have small effects and occur in non-coding regions. Integration of GWAS with gene regulatory networks addresses both issues by aggregating weak genetic signals within regulatory programs. Here we develop a Bayesian framework that integrates GWAS summary statistics with regulatory networks to infer genetic enrichments and associations simultaneously. Our method improves upon existing approaches by explicitly modeling network topology to assess enrichments, and by automatically leveraging enrichments to identify associations. Applying this method to 18 human traits and 38 regulatory networks shows that genetic signals of complex traits are often enriched in interconnections specific to trait-relevant cell types or tissues. Prioritizing variants within enriched networks identifies known and new trait-associated genes revealing novel biological and therapeutic insights.

## Introduction

Genome-wide association studies (GWAS) have catalogued many significant and reproducible associations between common genetic variants, notably single-nucleotide polymorphisms (SNPs), and diverse human complex traits^1^. However, it remains challenging^2^ to translate these findings into biological mechanisms and clinical applications, because most trait-associated variants have individually small effects and map to non-coding sequences.

One interpretation is that non-coding variants cumulatively affect complex traits through cell type- or tissue-specific^3^ gene regulation^4^. To test this hypothesis, large-scale epigenomic^5,6^ and transcriptomic^7–10^ data have been made available spanning diverse human cell types and tissues. Exploiting these regulatory genomic data, many studies have shown enrichments of trait-associated SNPs in chromatin regions^11–13^ and genes^14–16^ that are active in trait-relevant cell types or tissues. These studies simply overlap regulatory maps with GWAS data and often ignore functional interactions among loci within regulatory programs.

Gene regulatory networks^17–20^ have proven useful in mining functional interactions of genes from genomic data. Transcriptional regulatory interactions, rather than gene expression alone, drive tissue specificity^19^. Further, context-specific regulatory networks have emerged as promising tools to dissect the genetics of complex traits^21–23^. Network-connectivity analyses in GWAS have shown that trait-associated genes are more highly interconnected than expected^18^ and highly interconnected genes are enriched for trait heritability^24^. A major limitation of these analyses, however, is that they do not leverage observed enrichments to enhance trait-associated gene discovery.

To unleash the potential of regulatory networks in GWAS, we develop a novel framework for simultaneous genome-wide network enrichment and gene prioritization analysis. Through extensive simulations on the new method, we show its flexibility in various genetic architectures, its robustness to a wide range of model mis-specification, and its improved performance over existing methods. Applying the method to 18 human traits and 38 regulatory networks, we identify strong enrichments of genetic associations in network topology specific to trait-relevant cell types or tissues. By prioritizing variants within enriched networks we identify trait-associated genes that were not implicated by the same GWAS. Many of these putatively novel genes have strong support from multiple lines of external evidence; some are further validated by follow-up GWAS of the same traits with increased sample sizes. Together, these results demonstrate the potential for our method to yield novel biological and therapeutic insights from existing data.

## Results

### Method overview

Figure 1 shows the method schematic. In brief, we develop a new model dissecting the total effect of a single SNP on a trait into effects of multiple (nearby and distal) genes through a regulatory network, and then we combine it with a multiple-SNP regression likelihood^25^ based on GWAS summary statistics to perform Bayesian inference.

**Fig 1:**
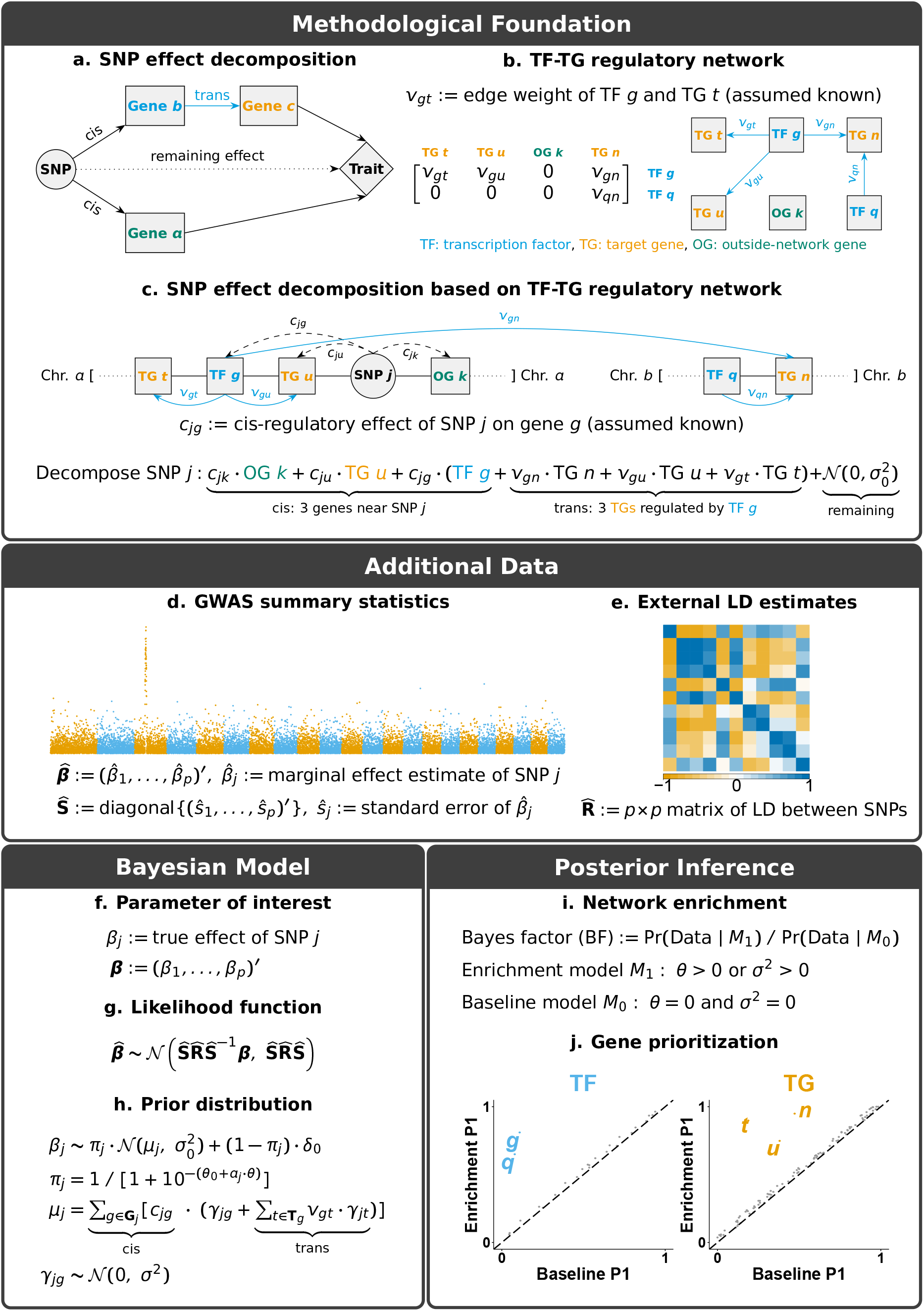
Schematic of RSS-NET. **a** Decomposition of the total effect of a common SNP on a complex trait through multiple nearby and distal genes. **b** Gene regulatory network defined as a weighted and directed bipartite graph linking TFs to TGs. **c** Given a TF-TG network, RSS-NET exploits its topology to decompose the total genetic effect into cis and trans regulatory components. Both the SNP-gene (*c_jg_*) and TF-TG (*v_gt_*) weights in the decomposition are assumed known and they are specified by existing omics data (Methods). **d-e** In addition to TF-TG networks, RSS-NET also requires GWAS summary statistics and ancestry-matching LD estimates as input. **f-h** Bayesian hierarchical models underlying RSS-NET. An in-depth description is provided in Methods. **i** Given a network, RSS-NET produces a BF comparing the baseline (M0) and enrichment (*M*_1_) models to summarize the evidence for network enrichment. **j** RSS-NET prioritizes loci within an enriched network by computing *P*_1_, the posterior probability that at least one SNP *j* in a locus is trait-associated (*β_j_* = 0). Differences between *P*_1_ under *M*_0_ and *M*_1_ reflect the influence of a regulatory network on genetic associations, highlighting putatively novel trait-associated genes.

We start with a conceptual decomposition of the total effect of a common SNP on a complex trait into three components: a cis-regulatory component through nearby genes, a trans-regulatory component through distal genes that are regulated by genes near this SNP, and a remaining component due to other factors (Fig. 1a). Since common genetic variation contributes to complex traits primarily via gene regulation^22^, we find this decomposition a sensible approximation to the genetic basis of complex traits.

Despite various ways to model the regulatory components, here we use cell type- or tissue-specific regulatory networks^18,20^ linking transcription factors (TFs) to target genes (TGs). Specifically, we define a regulatory network as a directed bipartite graph with weighted edges from TFs to TGs (Fig. 1b). Given a TF-TG network, we use its topology to decompose the total effect of each SNP into effects of multiple interconnected genes (Fig. 1c). For example, we represent the expected total effect of SNP *j* shown in Figure 1c as a weighted sum of cis effects of three nearby genes (outside-network gene *k*, TG *u* and TF *g*) and trans effects of three TGs (*n, u, t*) that are directly regulated by TF *g*. For identifiability we assume the SNP-gene (*c_jg_*) and TF-TG (*v_gt_*) weights in the decomposition are known, specified by existing omics data (Methods).

To implement this regulatory decomposition (Fig. 1c) in GWAS, we formulate a network-induced prior for SNP-level effects (*β*), and combine it with a multiple regression likelihood^25^ for *β* based on single-SNP association statistics from a GWAS (Fig. 1d) and linkage disequilibrium (LD) estimates from a reference panel with ancestry matching the GWAS (Fig. 1e). We refer to the resulting Bayesian framework (Fig. 1f-h) as Regression with Summary Statistics exploiting NEtwork Topology (RSS-NET).

RSS-NET accomplishes two tasks simultaneously: (1) testing whether a network is enriched for genetic associations (Fig. 1i); (2) identifying which genes within this network drive the enrichment (Fig. 1j). Specifically, RSS-NET estimates two independent enrichment parameters: *θ* and *σ*^2^, which measure the extent to which, SNPs near network genes and regulatory elements (REs) have increased likelihood to be associated with the trait, and, SNPs near network edges have larger effect sizes, respectively. To assess network enrichment, RSS-NET computes a Bayes factor (BF) comparing the “enrichment model” (*M*_1_: *θ* > 0 or *σ*^2^ > 0) against the “baseline model” (*M*_0_: *θ* = 0 and *σ*^2^ = 0). To prioritize genes within enriched networks, RSS-NET contrasts posterior distributions of *β* estimated under *M*_0_ and *M*_1_.

RSS-NET improves upon its predecessor RSS-E^16^. Specifically, RSS-NET exploits the full network topology, whereas RSS-E ignores the edge information. By explicitly modeling regulatory interconnections, RSS-NET outperforms RSS-E in both simulated and real datasets. Despite different treatments of network information, RSS-NET and RSS-E share computation schemes (Supplementary Notes), allowing us to reuse the efficient algorithm of RSS-E. Software is available at https://github.com/suwonglab/rss-net.

### Method comparison through simulations

The novelty of RSS-NET resides in a unified framework that leverages network topology to infer enrichments from whole-genome association statistics and prioritizes loci in light of inferred enrichments automatically. We are not aware of any published method with the same features. However, one could ignore topology and simply annotate SNPs based on their proximity to network genes and REs (Methods). For these SNP-level annotations there are methods to assess global enrichments or local associations on GWAS summary data. Here we use Pascal^26^, LDSC^13,27^ and RSS-E^16^ to benchmark RSS-NET.

Given a network, we first simulated SNP effects (*β*) from either RSS-NET assumed or mis-specified priors, and then combined them with real genotypes to simulate phenotypes from a genome-wide multiple-SNP model. We computed the corresponding single-SNP association statistics, on which we compared RSS-NET with other methods. Since RSS-NET is a model-based approach, we designed a large array of simulation scenarios for both correctly- and mis-specified *β*. To reduce computation of this large-scale design, we mainly used real genotypes^28^ of 348,965 genome-wide common SNPs and a whole-genome regulatory network inferred for B cell (436 TFs, 3,018 TGs)^20,29^. We also performed simulations on real genotypes^30^ of 1 million common SNPs^31^ or different networks, and obtained similar results.

We started with simulations where RSS-NET modeling assumptions were satisfied. We considered two genetic architectures: a sparse scenario with most SNPs being null and a polygenic scenario with most SNPs being trait-associated. For each architecture, we created negative datasets by simulating SNP effects (*β*) from *M*_0_ and positive datasets by simulating *β* from three *M*_1_ patterns (only *θ* > 0; only *σ*^2^ > 0; both *θ* > 0 and *σ*^2^ > 0) of the target network, and applied the methods to detect *M*_1_ from all datasets (Fig. 2 and Supplementary Fig.s 1-2). Existing methods tend to perform well in select settings. For example, Pascal and LDSC perform poorly when genetic signals are very sparse (Fig. 2b); RSS-E performs poorly when enrichment patterns are inconsistent with its modeling assumptions (Fig. 2c). Except for datasets with weak genetic signals on the network (Fig. 2d), RSS-NET performs consistently well in all scenarios. This is expected because the flexible models underlying RSS-NET can capture various genetic architectures and enrichment patterns. In practice, one rarely knows beforehand the correct genetic or enrichment architecture. This makes the flexibility of RSS-NET appealing.

**Fig 2:**
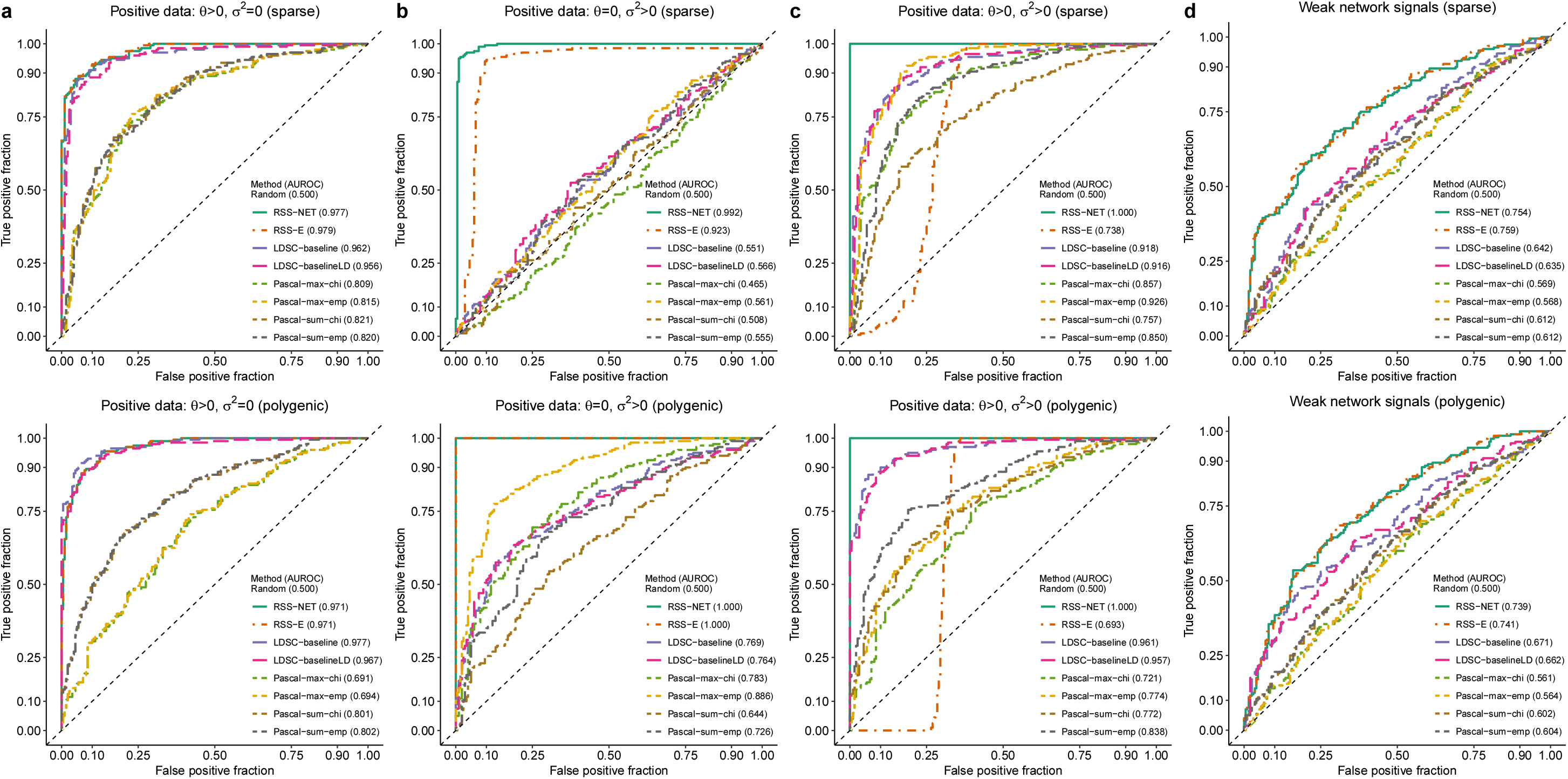
Flexibility of RSS-NET to identify network-based enrichments from GWAS summary statistics. We used a B cell-specific regulatory network and real genotypes of 348,965 genome-wide SNPs to simulate negative and positive individual-level data under two genetic architectures (“sparse” and “polygenic”). We used the baseline model (*M*_0_: 0 = 0 and *σ*^2^ = 0) to simulate SNP effects (*β*) for negative datasets. We simulated *β* for positive datasets from the enrichment model (*M*_1_:*θ*: 0 > 0 or *σ*^2^ > 0) for the target network under three scenarios: **a** 0 > 0, *σ*^2^ = 0; **b** 0 = 0, *σ*^2^ > 0; **c** 0 > 0, *σ*^2^ > 0. We computed the corresponding single-SNP association statistics, on which we compared RSS-NET with RSS-E^16^, LDSC-baseline^13^, LDSC-baselineLD^27^ and Pascal^26^ using their default setups (Methods). Pascal includes two gene scoring options: maximum-of-χ^2^ (`-max`) and sum-of-y^2^ (`-sum`), and two pathway scoring options: *χ*^2^ approximation (`-chi`) and empirical sampling (`-emp`). For each dataset, Pascal and LDSC methods produced *P*-values, whereas RSS-E and RSS-NET produced BFs; these statistics were used to rank the significance of enrichments. A false and true positive occurs if a method identifies enrichment of the target network in a negative and positive dataset respectively. Each panel displays the trade-off between false and true enrichment discoveries, namely receiver operating characteristics (ROC) curves, for all methods in 200 negative and 200 positive datasets of a simulation scenario, and also reports the corresponding areas under ROC curves (AUROCs, a higher value indicating better performance). Dashed diagonal lines denote random ROC curves (AUROC=0.5). d RSS-NET, as well as other methods, does not perform well when the target network harbors weak genetic associations. Simulation details and additional results are provided in Supplementary Figures 1-2.

Genetic associations of complex traits are enriched in regulatory regions^5,6^ Since a regulatory network is a set of genes linked by REs, it is important to confirm that network enrichments identified by RSS-NET are not driven by general regulatory enrichments. To this end we simulated negative datasets with enriched associations in random SNPs that are near genes (Fig. 3a; Supplementary Fig. 3) or REs (Fig. 3b; Supplementary Fig. 4). The results show that RSS-NET is unlikely to yield false discoveries due to arbitrary regulatory enrichments, and it is yet more powerful than other methods.

**Fig 3:**
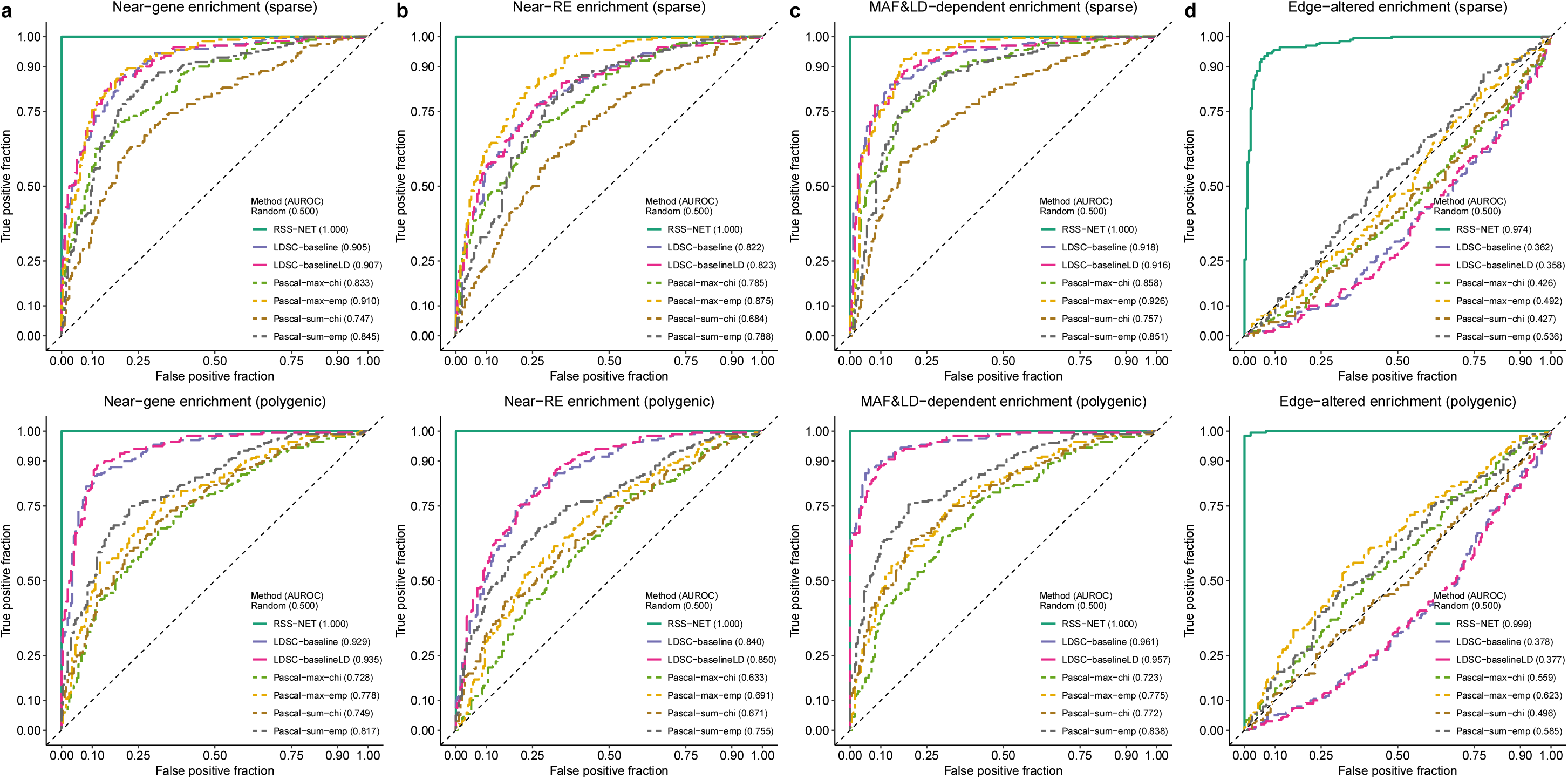
Robustness of RSS-NET to model mis-specification in enrichment analyses. Here positive datasets were generated from a special case of *M*_1_ with *θ* > 0 and *σ*^2^ > 0. Negative datasets were simulated from four scenarios where genetic associations were enriched in: **a** a random set of near-gene SNPs; **b** a random set of near-RE SNPs; **c** SNPs with MAF- and LD-dependent effects; **d** a random edge-altered network. By this design, RSS-NET was mis-specified in all four scenarios. Similar to positive datasets, the simulated false enrichments in all negative datasets manifested in both association proportion (more frequent) and magnitude (larger effect). RSS-E was excluded here because of its poor performance shown in Figure 2c. The rest of this simulation study is the same as Figure 2. Simulation details and additional results are provided in Supplementary Figures 3-6.

Minor allele frequency (MAF)- and LD-dependent genetic architectures have been identified in complex traits^27^. To assess the impact of MAF- and LD-dependence on RSS-NET results, we simulated MAF- and LD-dependent SNP effects (*β*) from an additive model of 10 MAF bins and 6 LD-related annotations^27^, which were then used to create negative datasets (Fig. 3c; Supplementary Fig. 5). Similarly, enrichments identified by RSS-NET are unlikely to be false positives induced by MAF- and LD-dependence.

Interconnections within regulatory programs play key roles in driving context specificity^19^ and propagating disease risk^22^, but existing methods often ignore the edge information. In contrast, RSS-NET leverages the full topology of a given network. The topology-aware feature increases the potential of RSS-NET to identify the most relevant network for a trait among candidates that share many nodes but differ in edges. To illustrate this feature, we designed a scenario where a real target network and random candidates had the same nodes and edge counts, but different edges. We simulated positive and negative datasets where genetic associations were enriched in the target network and random candidates respectively, and then tested enrichment of the target network on all datasets. As expected, only RSS-NET can reliably distinguish true enrichments of the target network from enrichments of its edge-altered counterparts (Fig. 3d; Supplementary Fig. 6).

To benchmark its prioritization component, we compared RSS-NET with gene-based association modules in RSS-E^16^ and Pascal^26^ (Fig. 4; Supplementary Fig.s 7-9). Consistent with previous work^16^, RSS methods outperform Pascal methods even without network enrichment (Fig. 4a). This is because RSS-NET and RSS-E exploit a multiple regression framework^25^ to learn the genetic architecture from data of all genes and assess their effects jointly, whereas Pascal only uses data of a single gene to estimate its effect. Similar to enrichment simulations (Fig. 2), RSS-NET outperforms RSS-E in prioritizing genes across different enrichment patterns (Fig. 4b-d). This again highlights the flexibility of RSS-NET.

**Fig 4:**
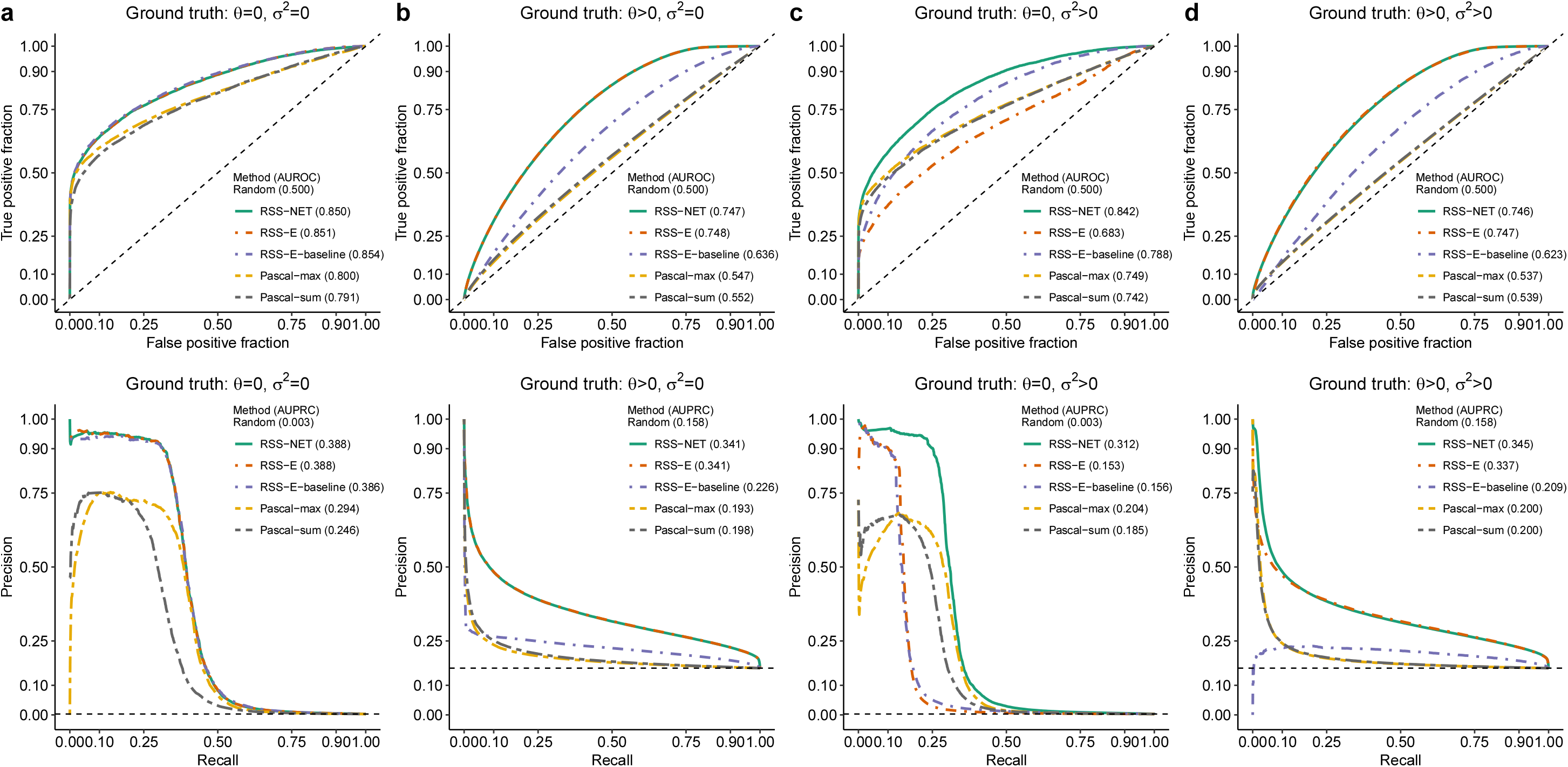
Power of RSS-NET to identify gene-based associations from GWAS summary statistics. We used a B cell-specific regulatory network and real genotypes of 348,965 genome-wide SNPs to simulate individual-level GWAS data under four scenarios: **a** *θ* = 0, *σ*^2^ = 0; **b** *θ* > 0, *σ*^2^ = 0; **c** *θ* = 0, *σ*^2^ > 0; **d** *θ* > 0, *σ*^2^ > 0. We computed the corresponding single-SNP summary statistics on which we compared RSS-NET with gene-based association components of RSS-E^16^ and Pascal^26^. RSS-E is a special case of RSS-NET assuming *σ*^2^ = 0, and RSS-E-baseline is a special case of RSS-E assuming *θ* = 0. Pascal includes two gene scoring options: maximum-of-of-*χ*^2^ (`-max`) and sum-of-*χ*^2^ (`-sum`). Given a network, Pascal and RSS-E-baseline do not leverage any network information, RSS-E ignores the edge information, and RSS-NET exploits the full topology. Each scenario contains 200 datasets and each dataset contains 16,954 autosomal protein-coding genes for testing. We defined a gene as “trait-associated” if at least one SNP *j* within 100 kb of the transcribed region of this gene had non-zero effect (*β_j_* ≠ 0). For each gene in each dataset, RSS methods produced posterior probabilities that the gene was trait-associated (*P*_1_), whereas Pascal methods produced association *P*-values; these statistics were used to rank the significance of gene-level associations. The upper half of each panel displays ROC curves and AUROCs for all methods, with dashed diagonal lines indicating random performance (AUROC=O.5). The lower half of each panel displays precision-recall (PRC) curves and areas under PRC curves (AUPRCs) for all methods, with dashed horizontal lines indicating random performance. For AUROC and AUPRC, higher value indicates better performance. Simulation details and additional results are provided in Supplementary Figures 7-8.

Finally, since RSS-NET uses a regulatory network as is, and, most networks to date are algorithmically inferred, we performed simulations to assess the robustness of RSS-NET under noisy networks. Specifically we simulated datasets from a real target network, created noisy networks by randomly removing edges from this real target, and then used the noisy networks (rather than the real one) in RSS-NET analyses. By exploiting retained true nodes and edges, RSS-NET produces reliable results in identifying both network enrichments and genetic associations, and unsurprisingly, its performance drops as the noise level increases (Supplementary Fig. 10).

In conclusion, RSS-NET is adaptive to various genetic architectures and enrichment patterns, it is robust to a wide range of model mis-specification, and it outperforms existing related methods. To further investigate its real-world utility, we applied RSS-NET to analyze 18 complex traits and 38 regulatory networks.

### Enrichment analyses of 38 networks across 18 traits

We first inferred^20^ whole-genome regulatory networks for 38 human cell types and tissues (Methods; Supplementary Table 1) from public data^29^ of paired expression and chromatin accessibility (PECA). On average each network has 431 TFs, 3,298 TGs and 93,764 TF-TG weighted edges. Clustering showed that networks recapitulated context similarity, with immune cells and brain regions grouping together as two units (Fig. 5a; Supplementary Fig. 11).

**Fig 5:**
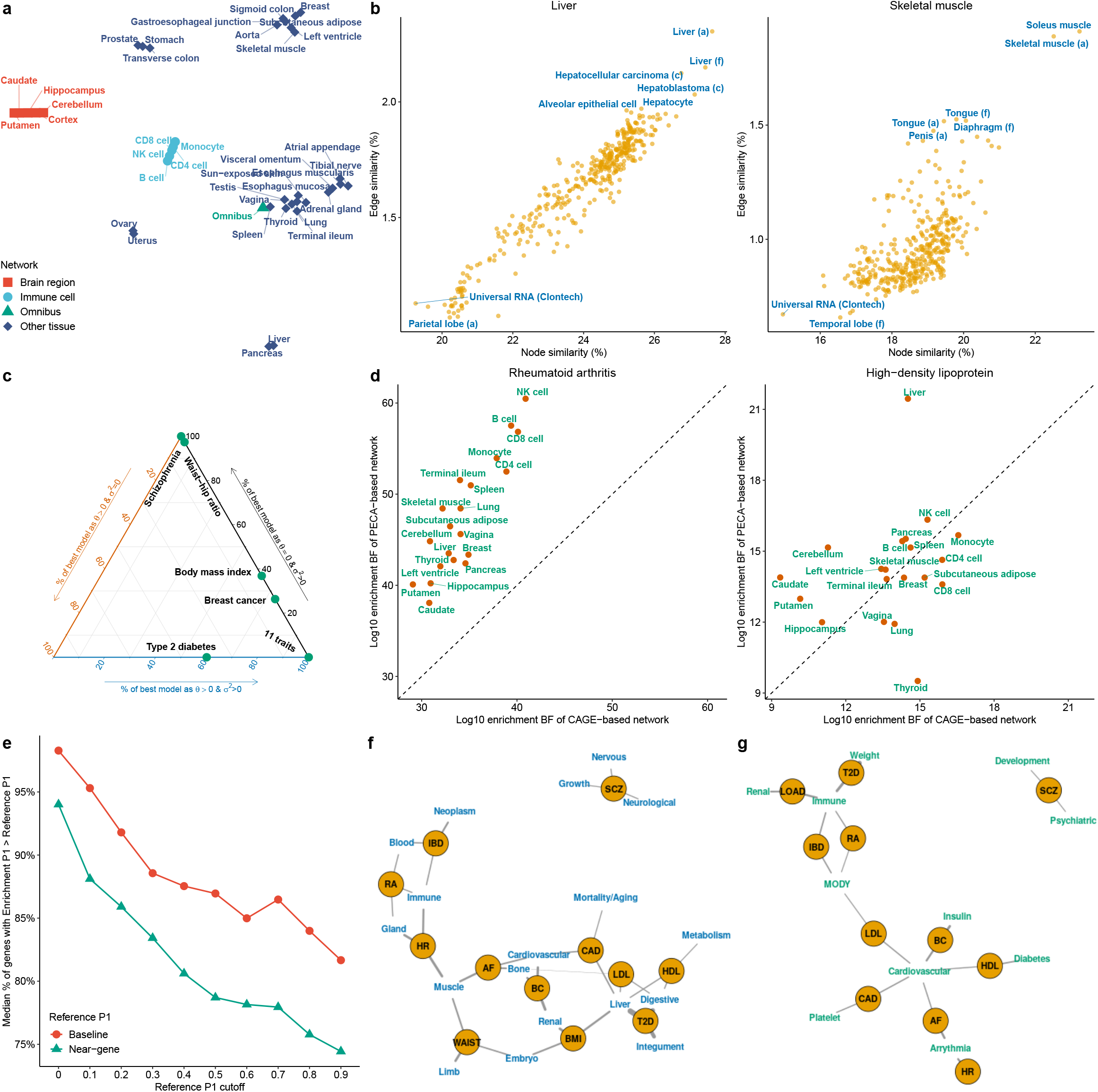
RSS-NET analyses of 18 complex traits and 38 regulatory networks. **a** Clustering of 38 regulatory networks based on í-distributed stochastic neighbour embedding. Details are provided in Supplementary Figure 11. **b** Similarity between a given tissue-specific PECA-based network and 394 CAGE-based networks for various cell types and tissues (a: adult samples; c: cell lines; f: fetal samples). The similarity between a PECA- and CAGE-based network is summarized by Jaccard indices of their node sets (*x*-axis) and edge sets (y-axis). Additional results are provided in Supplementary Figure 12. **c** Ternary diagram showing, for each trait, percentages of the “best” enrichment model (showing the largest BF) as *M*_11_: *θ* > 0, *σ*^2^ = 0, *M*_12_: *θ* = 0, *σ*^2^ > 0 and *M*_13_: *θ* > 0, *σ*^2^ > 0 across networks. See Supplementary Table 6 for numerical values. Shown are 16 traits that had multiple networks more enriched than the near-gene control. **d** Comparison of context-matched PECA-based (y-axis) and CAGE-based (x-axis) network enrichments on the same GWAS data. Dashed lines have slope 1 and intercept 0. Additional results are provided in Supplementary Figure 14. **e** Median proportion of genes with 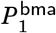 higher than reference estimates (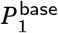 or 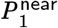), among genes with reference estimates higher than a given cutoff. Medians are evaluated among 16 traits that had multiple networks more enriched than the near-gene control. See Supplementary Table 8 for numerical values. **f-g** Overlap of RSS-NET prioritized genes 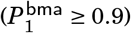 with genes implicated in knockout mouse phenotypes^38^ (**f**) and human Mendelian diseases^39,40^ (**g**). An edge indicates that a category of knockout mouse or Mendelian genes is significantly enriched for genes prioritized for a GWAS trait (FDR ≤ 0.1). Thicker edges correspond to stronger enrichment odds ratios. To simplify visualization, only top-ranked categories are shown for each trait (**f**: 3; **g**: 2). See Supplementary Tables 12-13 for full results. Trait abbreviations are defined in Supplementary Table 2.

As a validation, we assessed the pairwise similarity between the 38 PECA-based networks and 394 human cell type- and tissue-specific regulatory networks^18^ reconstructed from independent cap analysis of gene expression (CAGE) data^7,8^. As expected, PECA- and CAGE-based networks often reached maximum overlap when derived from biosamples of matched cell or tissue types (Fig. 5b; Supplementary Fig. 12), showing that the context specificity of PECA-based networks is replicable.

On the 38 networks, we applied RSS-NET to analyze 1.1 million common SNPs^31^ for 18 traits, using GWAS summary statistics from 20,883 to 253,288 European-ancestry individuals (Supplementary Table 2) and LD estimates from the European panel of 1000 Genomes Project^30^. For each trait-network pair we computed a BF assessing network enrichment. Full results of 684 trait-network pairs are available online (Data Availability).

To check whether observed enrichments could be driven by general regulatory enrichments, we created a “near-gene” control network with 18,334 protein-coding autosomal genes as nodes and no edges, and then analyzed this control with RSS-NET on the same GWAS data. For most traits, the near-gene control has substantially weaker enrichment than the actual networks. In particular, 512 out of 684 trait-network pairs (one-sided binomial *P* = 2.2 × 10^−40^) showed stronger enrichments than their near-gene counterparts (average log10 BF increase: 13.94; one-sided *t P* = 5.1 × 10^−15^), and, 16 out of 18 traits had multiple networks more enriched than the near-gene control (minimum: 5; one-sided Wilcoxon *P* = 1.2 × 10^−4^). In contrast, LDSC and Pascal methods identified fewer trait-network pairs passing the neargene enrichment control (LDSC maximum: 389, one-sided *χ*^2^ *P* = 1.7 × 10^−12^; Pascal maximum: 69, *P* = 2.0 × 10^−129^; Supplementary Table 3). Consistent with simulations (Fig. 3a-b), these results indicate that network enrichments identified by RSS-NET are unlikely driven by generic regulatory enrichments harbored in the vicinity of genes.

Among 512 trait-network pairs passing the near-gene enrichment control, we further examined whether the observed enrichments could be confounded by network properties or genomic annotations. We did not observe any correlation between BFs and three network features (proportion of SNPs in a network: Pearson *R* = −3.0 × 10^−2^, *P* = 0.49; node counts: *R* = −5.4 × 10^−2^, *P* = 0.23; edge counts: *R* = −9.2 × 10^−3^, *P* = 0.84). To check confounding effects of genomic annotations, we computed the correlation between BFs and proportions of SNPs falling into both a network and each of 73 functional categories^27^, and we did not find any significant correlation (−0.13 < *R* < −0.01, *P* > 0.05/73). Similar patterns hold for all 684 trait-network pairs (Supplementary Tables 4-5). Altogether, the results suggest that observed enrichments are unlikely driven by generic network or genome features.

For each trait-network pair, we also computed BFs comparing the baseline (*M*_0_) against three disjoint models where enrichment (*M*_1_) was contributed by (1) network genes and REs only (*M*_11_: *θ* > 0, *σ*^2^ = 0); (2) TF-TG edges only (*M*_12_: *θ* = 0, *σ*^2^ > 0); (3) network genes, REs and TF-TG edges (*M*_13_: *θ* > 0, *σ*^2^ > 0). We found that *M*_13_ was the most supported model by data (with the largest BF) for 411 out of 512 trait-network pairs (one-sided binomial *P* = 1.2 × 10^−45^), highlighting the key role of TF-TG edges in driving enrichments. To further confirm this finding, we repeated RSS-NET analyses by fixing all TF-TG edge weights as zero (*vtg* = 0) and we observed substantially weaker enrichments (average log10 BF decrease: 30.46; one-sided *tP* = 8.6× 10^−35^; Supplementary Fig. 13). Together the results corroborate the “omnigenic” model that genetic signals of complex traits are distributed via regulatory interconnections^22^.

When stratifying results by traits, however, we found that enrichment patterns varied considerably (Fig. 5c; Supplementary Table 6). For type 2 diabetes (T2D), two of five networks passing the near-gene enrichment control showed the strongest support for *M*_11_. Many networks showed the strongest support for *M*_12_ in breast cancer (10), body mass index (BMI, 14), waist-hip ratio (37) and schizophrenia (38). Since one rarely knows the true enrichment patterns a priori, and *M*_1_ includes {*M*_11_,*M*_12_,*M*_13_} as special cases, we used *M*_1_-based BFs throughout this study. Collectively, these results highlight the heterogeneity of network enrichments across complex traits, which can be potentially learned from data by flexible approaches like RSS-NET.

Top-ranked enrichments recapitulated many trait-context links reported in previous GWAS. Genetic associations with BMI were enriched in the networks of pancreas (BF = 2.07 × 10^13^), bowel (BF = 8.02 × 10^12^) and adipose (BF = 4.73 × 10^12^), consistent with the roles of obesity-related genes in insulin biology and energy metabolism. Networks of immune cells showed enrichments for rheumatoid arthritis (RA, BF = 2.95 × 10^60^), inflammatory bowel disease (IBD, BF = 5.07 × 10^35^) and Alzheimer’s disease (BF = 8.31 × 10^26^). Networks of cardiac and other muscle tissues showed enrichments for coronary artery disease (CAD, BF = 9.78 × 10^28^), atrial fibrillation (AF, BF = 8.55 × 10^14^), and heart rate (BF = 2.43 × 10^7^). Other examples include brain network with neuroticism (BF = 2.12 × 10^19^), and, liver network with high- and low-density lipoprotein (HDL, BF = 2.81 × 10^21^; LDL, BF = 7.66 × 10^27^).

Some top-ranked enrichments were not identified in the original GWAS, but they are biologically relevant. For example, natural killer (NK) cell network showed the strongest enrichment among 38 networks for BMI (BF = 3.95 × 10^13^), LDL (BF = 5.18 × 10^30^) and T2D (BF = 1.49 × 10^77^). This result supports a recent mouse study^32^ revealing the role of NK cell in obesity-induced inflammation and insulin resistance, and adds to the considerable evidence unifying metabolism and immunity in many pathological states^33^. Other examples include adipose network with CAD^34^ (BF = 1.67 × 10^29^), liver network with Alzheimer’s disease^16,35^ (BF = 1.09 × 10^20^) and monocyte network with AF^36,37^ (BF = 4.84 × 10^12^).

Some networks show enrichments in multiple traits. To assess network co-enrichments among traits, we tested correlations for all trait pairs using their BFs of 38 networks (Supplementary Table 7). In total 29 of 153 trait pairs were significantly correlated (*P* < 0.05/153). Reassuringly, subtypes of the same disease showed strongly correlated enrichments, as in IBD subtypes (*R* = 0.96, *P* = 1.3 × 10^−20^) and CAD subtypes (*R* = 0.90, *P* = 3.3 × 10^−14^). The results also recapitulated known genetic correlations including RA with IBD (*R* = 0.79, *P* = 5.3 × 10^−9^)and neuroticism with schizophrenia (*R* = 0.73, *P* = 1.6 × 10^−7^). Network enrichments of CAD were correlated with enrichments of its known risk factors such as heart rate (*R* = 0.75, *P* = 5.1 × 10^−8^), BMI (*R* = 0.71, *P* = 5.1 × 10^−7^), AF (*R* = 0.65, *P* = 9.2 × 10^−6^) and height (*R* = 0.64, *P* = 1.6 × 10^−5^). Network enrichments of Alzheimer’s disease were strongly correlated with enrichments of LDL (*R* = 0.90, *P* = 2.6 × 10^−14^) and IBD (*R* = 0.78, *P* = 8.3 × 10^−9^), consistent with roles of lipid metabolism and inflammation in Alzheimer’s disease^35^. Genetic correlations among traits are not predictive of correlations based on network enrichments (*R* = 0.12, *P* = 0.18), suggesting the additional explanatory power from regulatory networks to reveal trait similarities in GWAS.

To show that RSS-NET can be applied more generally, we analyzed the CAGE-based networks^18^ of 20 cell types and tissues that were also present in 38 PECA-based networks (Fig. 5d; Supplementary Fig. 14). PECA-based networks often produced larger BFs than their CAGE-based counterparts on the same GWAS data (average log10 BF increase: 17.36; one-sided *t P* = 1.4 × 10^−11^), suggesting that PECA-based networks are more enriched in genetic signals. Reassuringly, PECA- and CAGE-based networks consistentlyhighlighted known trait-context links (e.g. immune cells and autoimmune diseases, muscle tissues and heart diseases). For some traits PECA-based networks produced more informative results. For example, CAGE-based analysis of HDL showed a broad enrichment pattern across cell types and tissues (consistent with previous connectivity analysis^18^ of the same data), whereas PECA-based analysis identified liver as the top-enriched context by a wide margin. Although not our main focus, these results highlight the potential for RSS-NET to systematically evaluate different network inferences in GWAS.

### Enrichment-informed prioritization of network genes

A key feature of RSS-NET is that inferred network enrichments automatically contribute to prioritize associations of network genes (Method). Specifically, for each locus RSS-NET produces 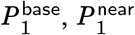 and 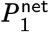, the posterior probability that at least one SNP in the locus is associated with the trait, assuming *M*_0_, *M*_1_ for the near-gene control network, and *M*_1_ for a given network, respectively. When multiple networks are enriched, RSS-NET produces 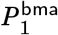 by averaging 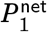 over all networks passing the near-gene control, weighted by their BFs. This allows us to assess genetic associations in light of enrichment without having to select a single enriched network. Differences between enrichment estimates 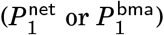 and reference estimates 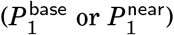 reflect the impact of network on a locus.

RSS-NET enhances genetic association detection by leveraging inferred enrichments. To quantify this improvement, for each trait we calculated the proportion of genes with higher 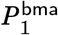 than reference estimates 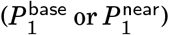, among genes with reference *P*_1_ passing a given cutoff (Fig. 5e). When using 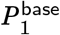 as reference, we observed high proportions of genes with 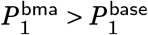 (median: 82 – 98%) across a wide range of 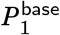-cutoffs (0 – 0.9), and as expected, the improvement decreased as the reference cutoff increased. When using 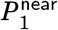 as reference, we observed less genes with improved *P*_1_ than using 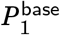 (one-sided Wilcoxon *P* = 9.8 × 10^−4^), suggesting the observed improvement might be partially due to general near-gene enrichments, but proportions of genes with 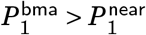 remained high (median: 74 – 94%) nonetheless. Similar patterns occurred when we repeated the analysis with 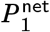 across 512 trait-network pairs (Supplementary Table 8). Together the results demonstrate the strong influence of network enrichments on nominating additional trait-associated genes.

RSS-NET tends to promote more genes in networks with stronger enrichments. For each trait the proportion of genes with 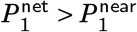 in a network is often positively correlated with its enrichment BF (R: 0.28 – 0.91; Supplementary Table 9). When a gene belongs to multiple networks, its highest 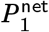 often occurs in the top-enriched networks. We illustrate this coherent pattern with *MT1G,* a liver-active^9^ gene that was prioritized for HDL by RSS-NET and also implicated in a recent multi-ancestry genome-wide interaction analysis of HDL^41^. Although *MT1G* belongs to regulatory networks of 18 contexts, only the top enrichment in liver (BF = 2.81 × 10^21^) informs a strong association between *MT1G* and HDL 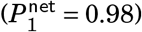, and remaining networks with weaker enrichments yield minimal improvement 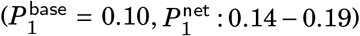. Figure 6 shows additional examples.

RSS-NET recapitulates many genes implicated in the same GWAS. For each analyzed dataset we downloaded the GWAS-implicated genes from the GWAS Catalog^1^ and computed the proportion of these genes with high 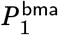. With a stringent cutoff 0.9, we observed a significant overlap (median across traits: 69%; median Fisher exact *P* = 1.2 × 10^−26^; Supplementary Table 10). Reassuringly, many recapitulated genes are well-established for the traits (Supplementary Table 11), such as *CACNA1C* for schizophrenia, *TCF7L2* for T2D, *APOB* for lipids and *STAT4* for autoimmune diseases.

RSS-NET also uncovers putative associations that were not reported in the same GWAS. To demonstrate that many of these new associations are potentially real we exploited 15 analyzed traits that each had a updated GWAS with larger sample size. In each case we obtained newly implicated genes from the GWAS Catalog^1^ and computed the proportion of these genes that were identified by RSS-NET 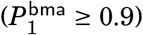. The overlap proportions remained significant (median: 12%; median Fisher exact *P* = 1.9 × 10^−5^; Supplementary Table 10), showing the potential of RSS-NET to identify trait-associated genes that can be validated by later GWAS with additional samples. Among these validated genes, many are strongly supported by multiple lines of external evidence. A particular example is *NR0B2,* a liver-active^9^ gene prioritized for HDL (BF = 2.81 × 10^21^, 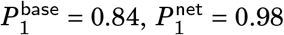), which was not identified by standard GWAS^43^ of the same data (minimum single-SNP *P* = 1.4 × 10^−7^ within 100 kb, *n* = 99,900). *NR0B2* was associated with mouse lipid traits^44–46^ and human obesity^47^, and identified in a later GWAS of HDL^48^ with doubled sample size (P = 9.7 × 10^−16^, *n* = 187,056). Table 1 lists additional examples.

**Table 1.**
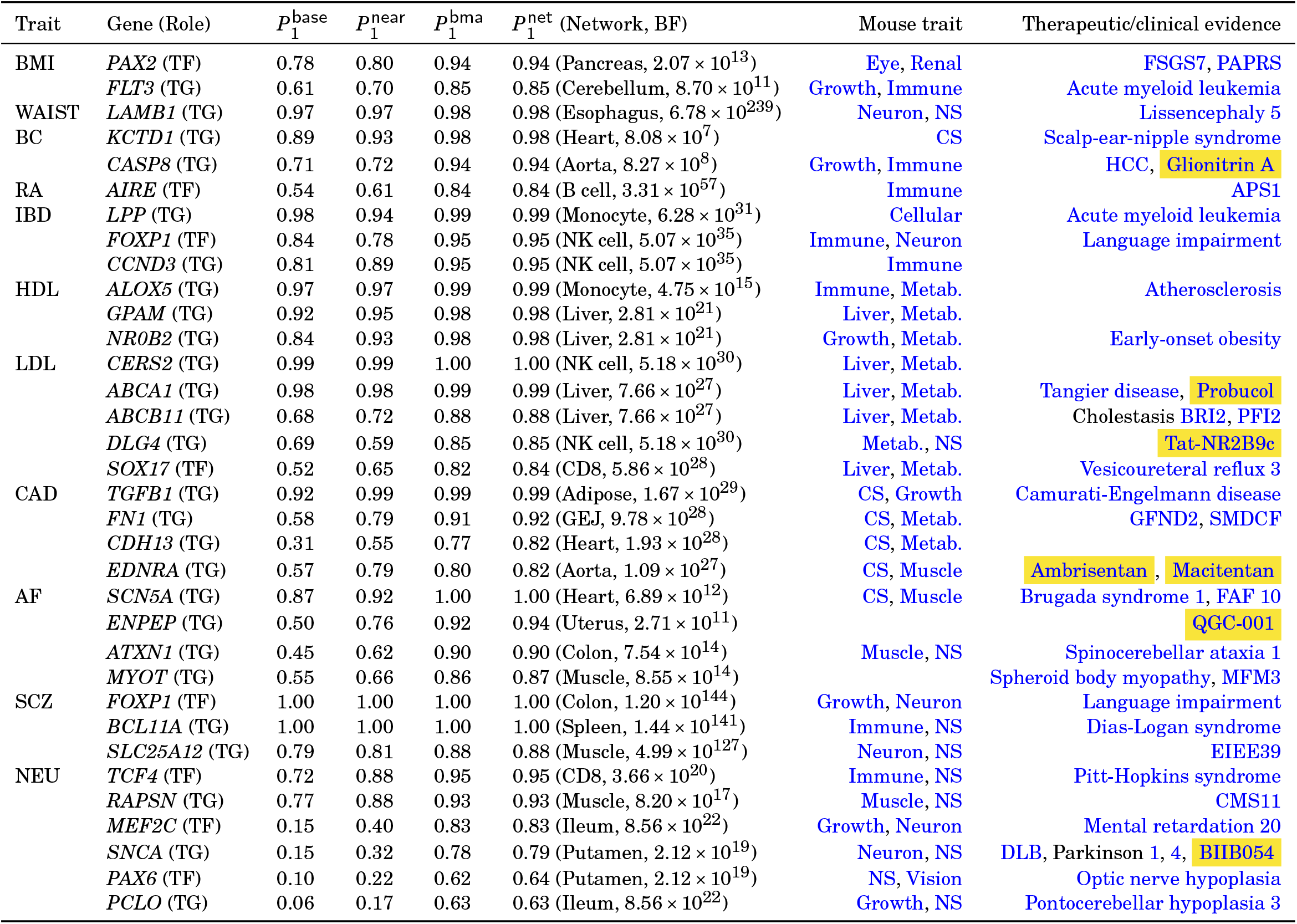
Examples of RSS-NET highlighted genes that were not reported in GWAS of the same data (*P*≥ 5×10^−8^) but were implicated in later GWAS with increased sample sizes (*P* < 5×10^−8^). The “mouse trait” column is based on the Mouse Genome Informatics 38. The “therapeutic/clinical evidence” column is based on the Online Mendelian Inheritance in Man39 and Therapeutic Target Database42. Click blue links to view details online. Drugs are highlighted in yellow. Abbreviations of GWAS traits are defined in Supplementary Table 2. GEJ: gastroesophageal junction; CS: cardiovascular system; DS: digestive/alimentary system; Metab.: metabolism; NS: nervous system.

### Biological and clinical relevance of prioritized genes

Besides looking up overlaps with GWAS publications, we cross-referenced RSS-NET prioritized genes 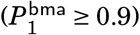 with multiple orthogonal databases to systematically assess their biological and therapeutic themes.

Mouse phenomics provides important resources to study genetics of human traits^49^. Here we evaluated overlap between RSS-NET prioritized genes and genes implicated in 27 categories of knockout mouse phenotypes^38^. Network-informed genes 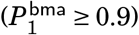 were significantly enriched in 128 mouse-human trait pairs (FDR ≤ 0.1; Supplementary Table 12). Fewer significant pairs were identified without network information (119 for 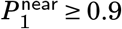; 80 for 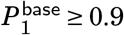). For many human traits, top enrichments of network-prioritized genes occurred in closely related mouse phenotypes (Fig. 5f). Schizophrenia-associated genes were strongly enriched in nervous, neurological and growth phenotypes (OR: 1.77 – 2.04). Genes prioritized for autoimmune diseases were strongly enriched in immune and hematopoietic phenotypes (OR: 2.05 – 2.35). The cardiovascular system showed strong enrichments of genes associated with heart conditions (OR: 2.45 – 2.92). The biliary system showed strong enrichments of genes associated with lipids, BMI, CAD and T2D (OR: 2.16 −10.78). The phenotypically matched cross-species enrichments strengthen the biological relevance of RSS-NET results.

Mendelian disease-causing genes often contribute to complex traits^50^. Here we quantified overlap between RSS-NET prioritized genes and genes causing 19 categories^40^ of Mendelian disorders^39^. Leveraging regulatory networks 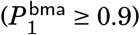, we observed 47 significantly enriched Mendelian-complex trait pairs (FDR ≤ 0.1; 44 for 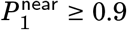; 31 for 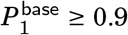; Supplementary Table 13), among which the top-ranked ones were often phenotypically matched (Fig. 5g). Schizophrenia-associated genes were strongly enriched in Mendelian development and psychiatric disorders (OR: 2.22-2.23). Genes prioritized for AF and heart rate were strongly enriched in arrhythmia (OR: 7.16 – 8.28). Genes prioritized for autoimmune diseases were strongly enriched in monogenic immune dysregulation (OR: 3.11 −4.32). Monogenic cardiovascular diseases showed strong enrichments of genes associated with lipids and heart conditions (OR: 2.69 – 3.70). We also identified pairs where Mendelian and complex traits seemed unrelated but were indeed linked. Examples include Alzheimer’s disease with immune dysregulation^35^ (OR = 7.32) and breast cancer with insulin disorders^51^ (OR = 9.71). The results corroborate that Mendelian and complex traits exist on a continuum.

Human genetics has proven valuable in therapeutic development^52^. To evaluate their potential in drug discovery, we examined whether RSS-NET prioritized genes are pharmacologically active targets with known clinical indications^42^. We identified genes with perfectly matched drug indications and GWAS traits. The most illustrative identical match is *EDNRA,* a gene that is prioritized for CAD (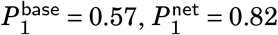 in aorta network), and is also a successful target of approved drugs for cardiovascular diseases (Table 1). We identified genes with closely related drug indications and GWAS traits. For example, *TTR* is prioritized for Alzheimer 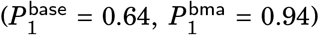, and is also a successful target of approved drugs for amyloidosis (Table 2). For early-stage development, overlaps between drug indications and GWAS traits may provide additional genetic confidence. For example, HCAR3 is prioritized for HDL 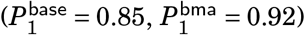, and is also a clinical trial target for lipid metabolism disorders (Table 2). Other examples include CASP8 with cancer, NFKB2 with IBD, and DLG4 with stroke (Tables 1–2). We also found mismatches between drug indications and GWAS traits, which could suggest drug repurposing opportunities^53^. For example, CSF3 is prioritized for AF 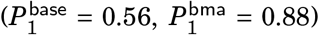, and is also a successful target of an approved drug for aplastic anemia (AA). Since CSF3 is associated with various blood cell traits in mouse^54^ and human^55^, and inflammation plays a role in both AA and AF etiology^36,37,56^, it is tempting to assess effects of the approved AA drug on AF. Mechanistic evaluations are required to understand the prioritized therapeutic genes, but they could form a useful basis for future studies.

**Table 2.** Examples of RSS-NET highlighted genes that have not reached genome-wide significance in the GWAS Catalog1 (*P* ≥ 5×10^8^) at the time of analysis. The rest is the same as Table 1.

| Trait | Gene (Role) | $P_1^{\text{base}}$ | $P_1^{\text{near}}$ | $P_1^{\text{bma}}$ | $P_1^{\text{net}}$ (Network, BF) | Mouse trait | Therapeutic/clinical evidence |
| --- | --- | --- | --- | --- | --- | --- | --- |
| BMI | *NEXN* (TG) | 0.71 | 0.79 | 0.89 | 0.90 (Muscle, $9.31 \times 10^{12}$) | CS, Muscle | Cardiomyopathy D1CC, H20 |
| | *CDX2* (TF) | 0.61 | 0.70 | 0.83 | 0.86 (NK cell, $3.95 \times 10^{13}$) | DS, Growth | |
| WAIST | *BSCL2* (TG) | 0.80 | 0.68 | 0.87 | 0.87 (Esophagus, $6.78 \times 10^{239}$) | Adipose, Growth | Lipodystrophy CG2 |
| | *FOXP2* (TF) | 0.56 | 0.59 | 0.73 | 0.73 (Esophagus, $6.78 \times 10^{239}$) | Growth, NS | Speech-language disorder 1 |
| BC | *ADSL* (TG) | 0.76 | 0.80 | 0.91 | 0.92 (Aorta, $8.27 \times 10^8$) | CS, Eye | Adenylosuccinase deficiency |
| | *SYNE1* (TG) | 0.57 | 0.63 | 0.89 | 0.90 (Esophagus, $6.30 \times 10^7$) | Growth, Muscle | AMCM, EDMD4, SCAR8 |
| RA | *TAL1* (TF) | 0.71 | 0.79 | 0.91 | 0.93 (CD4, $3.02 \times 10^{52}$) | Immune, Tumor | Acute lymphocytic leukemia |
| | *FHIT* (TG) | 0.30 | 0.60 | 0.90 | 0.91 (CD4, $3.02 \times 10^{52}$) | Immune, Tumor | |
| | *FLT3* (TG) | 0.33 | 0.57 | 0.73 | 0.73 (B cell, $3.31 \times 10^{57}$) | Immune, Tumor | Acute myeloid leukemia |
| IBD | *FHIT* (TG) | 0.63 | 0.87 | 0.95 | 0.95 (CD4, $5.32 \times 10^{33}$) | Immune, Tumor | |
| | *GATA3* (TF) | 0.85 | 0.83 | 0.94 | 0.94 (NK cell, $5.07 \times 10^{35}$) | Immune, Renal | Barakat syndrome |
| | *RORA* (TF) | 0.66 | 0.78 | 0.87 | 0.90 (B cell, $1.49 \times 10^{32}$) | Immune, NS | IDDECA |
| | *NFKB2* (TF) | 0.74 | 0.85 | 0.84 | 0.88 (B cell, $1.49 \times 10^{32}$) | Immune | CVID10, DIMS-0150 |
| | *LRBA* (TG) | 0.42 | 0.58 | 0.72 | 0.72 (NK cell, $5.07 \times 10^{35}$) | Immune | Immunodeficiency CV8 |
| | *DOCK2* (TG) | 0.38 | 0.53 | 0.71 | 0.71 (NK cell, $5.07 \times 10^{35}$) | Immune | Immunodeficiency 40 |
| HDL | *MT1G* (TG) | 0.10 | 0.09 | 0.98 | 0.98 (Liver, $2.81 \times 10^{21}$) | CS, Metab. | |
| | *RETSAT* (TG) | 0.79 | 0.80 | 0.95 | 0.95 (Liver, $2.81 \times 10^{21}$) | Adipose, Metab. | |
| | *ESR1* (TF) | 0.77 | 0.82 | 0.95 | 0.95 (Liver, $2.81 \times 10^{21}$) | CS, Metab. | Myocardial infarction |
| | *HCAR3* (TG) | 0.85 | 0.85 | 0.92 | 0.92 (Monocyte, $4.75 \times 10^{15}$) | Metab. | ARI-3037MO |
| | *TNNC1* (TG) | 0.48 | 0.45 | 0.78 | 0.78 (Liver, $2.81 \times 10^{21}$) | CS, Muscle | CMD1Z, CMH13, Levosimendan |
| LDL | *RAF1* (TG) | 0.79 | 0.83 | 0.90 | 0.90 (Aorta, $3.71 \times 10^{27}$) | CS, Immune | CMD1NN, Semapimod |
| | *APOA1* (TG) | 0.70 | 0.76 | 0.90 | 0.90 (Liver, $7.66 \times 10^{27}$) | CS, Metab. | Amyloidosis, HDL deficiency |
| | *ACADVL* (TG) | 0.69 | 0.59 | 0.85 | 0.85 (NK cell, $5.18 \times 10^{30}$) | Liver, Metab. | VLCAD deficiency |
| T2D | *ITGB6* (TG) | 0.75 | 0.99 | 0.99 | 0.99 (Ileum, $4.52 \times 10^{62}$) | Immune, Metab. | AI1H |
| HR | *TKT* (TG) | 0.65 | 0.67 | 0.92 | 0.93 (Aorta, $2.43 \times 10^7$) | CS, Growth | SDDHD |
| CAD | *OSM* (TG) | 0.56 | 0.78 | 0.86 | 0.86 (Aorta, $1.09 \times 10^{27}$) | Immune, Metab. | GSK2330811 |
| | *TRIB1* (TG) | 0.43 | 0.68 | 0.85 | 0.85 (Adipose, $1.67 \times 10^{29}$) | Adipose, Metab. | |
| | *TAB2* (TG) | 0.19 | 0.43 | 0.61 | 0.61 (CD8, $1.13 \times 10^{25}$) | CS | Congenital heart defects |
| AF | *TPMT* (TG) | 0.88 | 0.93 | 0.99 | 0.99 (Ileum, $4.43 \times 10^{13}$) | Metab. | THPM1 |
| | *RUNX1* (TF) | 0.44 | 0.60 | 0.88 | 0.89 (Heart, $2.15 \times 10^{14}$) | CS, Immune | Acute myeloid leukemia, FPDMM |
| | *CSF3* (TG) | 0.56 | 0.72 | 0.88 | 0.88 (Muscle, $8.55 \times 10^{14}$) | Blood, Immune | Interleukin-3 |
| LOAD | *CASP2* (TG) | 0.99 | 1.00 | 1.00 | 1.00 (CD8, $8.31 \times 10^{26}$) | Cellular, NS | Caspase-2 |
| | *TTR* (TG) | 0.64 | 0.92 | 0.94 | 0.94 (Pancreas, $3.53 \times 10^{20}$) | Metab. | FAP, Inotersen, Patisiran |
| SCZ | *RORA* (TF) | 1.00 | 1.00 | 1.00 | 1.00 (Cortex, $5.39 \times 10^{128}$) | Neuron, NS | IDDECA |
| | *ERBB4* (TG) | 1.00 | 1.00 | 1.00 | 1.00 (Putamen, $7.22 \times 10^{116}$) | Neuron, NS | ALS19 |
| | *NFIB* (TF) | 0.97 | 0.97 | 0.98 | 0.98 (Cortex, $5.39 \times 10^{128}$) | NS | MACID |
| | *GRIK2* (TG) | 0.90 | 0.94 | 0.97 | 0.97 (Cerebellum, $3.15 \times 10^{129}$) | Neuron, NS | Mental retardation 6 |
| | *SYT1* (TG) | 0.84 | 0.89 | 0.93 | 0.93 (Cerebellum, $3.15 \times 10^{129}$) | Neuron, NS | Baker-Gordon syndrome |
| | *ESR1* (TF) | 0.80 | 0.84 | 0.93 | 0.93 (Colon, $1.07 \times 10^{141}$) | Neuron, NS | Migraine |
| | *NTRK2* (TG) | 0.78 | 0.84 | 0.91 | 0.91 (Cerebellum, $3.15 \times 10^{129}$) | Neuron, NS | EIEE58 |
| | *LRRK2* (TG) | 0.73 | 0.78 | 0.86 | 0.86 (Monocyte, $5.85 \times 10^{131}$) | Neuron, NS | Parkinson 8, DNL151, DNL201 |
| | *C9orf72* (TG) | 0.74 | 0.78 | 0.83 | 0.83 (Spleen, $1.44 \times 10^{141}$) | Neuron, NS | FTDALS1 |
| | *SNCA* (TG) | 0.60 | 0.66 | 0.74 | 0.74 (Cerebellum, $3.15 \times 10^{129}$) | Neuron, NS | DLB, Parkinson 1, 4 |
| NEU | *LMBRD1* (TG) | 0.42 | 0.66 | 0.94 | 0.94 (Ileum, $8.56 \times 10^{22}$) | Metab. | MAHCF |
| | *PRKCQ* (TG) | 0.36 | 0.56 | 0.90 | 0.91 (Spleen, $2.13 \times 10^{19}$) | Immune, NS | |
| | *ATP1A2* (TG) | 0.33 | 0.39 | 0.76 | 0.78 (Putamen, $2.12 \times 10^{19}$) | Neuron, NS | AHC1, FHM2 |

## Discussion

We present RSS-NET, a new topology-aware method for integrative analysis of regulatory networks and GWAS summary data. We demonstrate the improvement of RSS-NET over existing methods through extensive simulations, and illustrate its potential to yield novel insights via analyses of 38 networks and 18 traits. With multi-omics integration becoming a routine in GWAS, we expect that researchers will find RSS-NET useful.

Compared with existing integrative approaches, RSS-NET has several key strengths. First, unlike many methods that require loci passing a significance threshold^11,12,17^, RSS-NET uses data from genome-wide common variants. This potentially allows RSS-NET to identify subtle enrichments even in studies with few significant hits. Second, RSS-NET models enrichments directly as increased rates *(θ)* and sizes (*σ*^2^) of SNP-level associations, and thus bypasses the issue of converting SNP-level summary data to gene-level statistics^17,18,26^. Third, RSS-NET inherits from RSS-E^16^ an important feature that inferred enrichments automatically highlight which network genes are most likely to be trait-associated. This prioritization component, though useful, is missing in current polygenic analyses^13,15,24,27^. Fourth, by making flexible modeling assumptions, RSS-NET is adaptive to unknown genetic and enrichment architectures.

RSS-NET provides a new view of complex trait genetics through the lens of regulatory topology. Complementing previous connectivity analyses^17–19,24^, RSS-NET highlights a consistent pattern where genetic signals of complex traits often distribute across genome via regulatory topology. RSS-NET further leverages topology enrichments to enhance trait-associated gene discovery. The topology awareness of RSS-NET relies on a novel model that decomposes effect size of a single SNP into effects of multiple (cis or trans) genes through a regulatory network. Other than a theoretical perspective^22^, we are not aware of any publication implementing the topology-aware model in practice.

RSS-NET depends critically on the quality of input regulatory networks. The more accurate networks are, the better performance RSS-NET achieves. Currently our understanding of regulatory networks remains incomplete, and most of available networks are algorithmically inferred^17–20^. Artifacts in inferred networks can bias RSS-NET results; however our simulations confirm the robustness of RSS-NET when input networks are not severely deviated from ground truth. The modular design of RSS-NET enables systematic assessment of various networks in the same GWAS and provides interpretable performance metrics, as illustrated in our comparison of PECA- and CAGE-based networks. As more accurate networks become available in diverse cellular contexts, the performance of RSS-NET will be markedly enhanced.

Like any method, RSS-NET has several limitations in its current form. First, despite its prioritization feature, RSS-NET does not attempt to pinpoint associations to causal SNPs within prioritized loci. For this task we recommend off-the-shelf fine-mapping methods^57^. Second, the computation time of RSS-NET increases as the total number of analyzed SNPs increases, and thus our analyses focused on 1.1 million common SNPs^31^. Relaxing the complexity will allow RSS-NET to analyze more SNPs jointly. Third, RSS-NET uses a simple method to derive SNP-gene relevance (*c_jg_*) from expression quantitative trait loci (eQTL). A more principled approach would be applying the RSS likelihood^25^ to eQTL summary data (as we did in GWAS) and using the estimated SNP effects to specify *c_jg_*. However, our initial assessments indicated that the model-based approach was limited by the small sample sizes of current eQTL studies^9,10^. With eQTL studies reaching large sample sizes^58^ comparable to current GWAS^1^, this approach may improve *c jg* specification in RSS-NET. Fourth, RSS-NET analyzes one network at a time. Since a complex disease typically manifests in various sites, multiple cellular networks are likely to mediate disease risk jointly. To extend RSS-NET to incorporate multiple networks, an intuitive idea would be representing the total effect of a SNP as an average of its effect size in each network, weighted by network relevance for a disease. Fifth, RSS-NET does not include known SNP-level^13,24,27^ or gene-level^14–16^ annotations. Although our mis-specification simulations and near-gene control analyses confirm that RSS-NET is robust to generic enrichments of known features, accounting for known annotations can help interpret observed network enrichments^24^. Our preliminary experiments, however, showed that incorporating additional networks or annotations in RSS-NET increased computation costs. Hence, we view developing computationally efficient multi-network, multi-annotation methods as an important area for future work.

In summary, improved understanding of human complex trait genetics requires biologically-informed models beyond the standard one employed in GWAS. By modeling tissue-specific regulatory topology, RSS-NET is a step forward in this direction.

## Methods

### Gene and SNP information

This study used genes and SNPs from the human genome assembly GRCh37. This study used 18,334 protein-coding autosomal genes(http://ftp.ensembl.org/pub/grch37/release-94/gtf/homo_sapiens, accessed January 3, 2019). Simulations used 348,965 genome-wide SNPs^28^ (https://www.wtccc.org.uk), and data analyses used 1,289,786 genomewide HapMap3^31^ SNPs (https://data.broadinstitute.org/alkesgroup/LDSCORE/w_hm3.snplist.bz2, accessed November 27, 2018). As discussed later, these SNP sets were chosen to reduce computation. This study also excluded SNPs on sex chromosomes, SNPs with minor allele frequency less than 1%, and SNPs in the human leukocyte antigen region.

### GWAS summary statistics and LD estimates

The European-ancestry GWAS summary statistics (Supplementary Table 2) and LD estimates used in the present study were processed as in previous work^16^. Data sources and references are provided in Supplementary Notes.

### Gene regulatory networks

In this study a regulatory network is a directed bipartite graph {**V**_TF_, **V**_TG_,**E**_TF→TG_}, where **V**_TF_ denotes the node set of TFs, **V**_TG_ denotes the node set of TGs, and **E**_TF→TG_ denotes the set of directed TF-TG edges, summarizing how TFs regulate TGs through REs (Fig. 1b, Supplementary Notes). Each edge has a weight between 0 and 1, measuring the relative regulation strength of a TF on a TG.

Here we inferred 38 regulatory networks from context-matched high-throughput sequencing data of gene expression (e.g., RNA-seq) and chromatin accessibility (e.g., DNAse-seq or ATAC-seq). We obtained these PECA data from portals of ENCODE^29^ (https://www.encodeproject.org, accessed December 14, 2018) and GTEx^9^ (https://gtexportal.org, accessed July 13, 2019); see Supplementary Table 1. The network-construction software and TF-motif information are available at https://github.com/suwonglab/PECA. The 38 networks are available at https://github.com/suwonglab/rss-net, with descriptive statistics provided in Supplementary Tables 14-16.

We first constructed an “omnibus” network from PECA data of 201 biosamples across 80 cell types and tissues, using a regression-based method^20^. In brief, by modeling the distribution of TG expression levels conditional on RE accessibility levels and TF expression levels, we estimated a regression coefficient for each TF-TG pair. We selected a TF-TG pair as the network edge if this estimated coefficient was significantly non-zero, and divided its estimate by the maximum of estimates for all TF-TG pairs to set a (0, 1)-scale edge weight. We also estimated a regression coefficient for each RE-TG pair, which reflected the regulating strengths of REs on TGs and was later used to construct context-specific networks (i.e., {*I_it_*} in (1)). Here we defined REs as open chromatin peaks called from accessibility sequencing data by MACS2^59^.

With the omnibus network in place, we then constructed context-specific networks for 5 immune cell types, 5 brain regions and 27 non-brain tissues. For each context (tissue or cell type), we computed a trans-regulation score (TRS) between TF *g* and TG *t*:

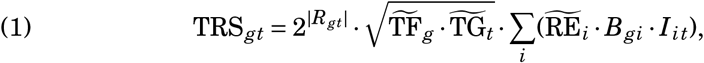

where *R_gt_* is the correlation of TF *g* and TG *t* expression levels across all contexts; 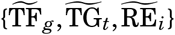 are normalized context-specific expression (TF *g*, TG *t*) and accessbility (RE *i*) levels (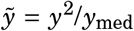, *y*: acutal RE accessibility or gene expression level in the given context, *y*_med_: median level across all contexts); *B_gi_* reflects the motif binding strength of TF *g* on RE *i*, defined as the sum of motif position weight matrix-based log-odds probabilities of all binding sites on RE *i* (calculated by HOMER^60^); and *I_it_* reflects the overall regulating strength of RE *i* on TG *t*, provided by the omnibus network above. TRS offers a natural way to rank and select context-specific TF-TG edges because a larger value of TRS_*gt*_ indicates a stronger regulating strength of TF *g* on TG *t* in the given context. We further set (0,1)-scale TF-TG edge weights by computing log_2_(1 + TRS_*gt*_) / max_(*i;j*)_{log_2_(1 + TRSs_*ij*_)}.

To benchmark PECA-based networks and illustrate RSS-NET as a generally applicable tool, we analyzed 394 published human cell type- and tissuespecific TF-TG circuits^18^ inferred from independent CAGE data^7,8^ (http://regulatorycircuits.org/, accessed May 8, 2019). When evaluating the similarity between PECA- and CAGE-based networks (Fig. 5b, Supplementary Fig. 12), we used their full node and edge sets to compute Jaccard indices. When running RSS-NET on context-matched PECA- and CAGE-based networks (Fig. 5d, Supplementary Fig. 14), we selected top-ranked CAGE-based edges to match PECA-based edge counts (Supplementary Table 15) and normalized CAGE-based edge weights 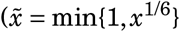, *x*: original weight) to match the scale of PECA-based edge weights (Supplementary Table 16).

### External databases for cross-reference

To validate and interpret RSS-NET results, we used the following external databases (accessed November 28,2019): GWAS Catalog^1^ (https://www.ebi.ac.uk/gwas/), Mouse Genome Informatics^38^ (http://www.informatics.jax.org/), phenotype-specific Mendelian gene sets^40^ (https://github.com/bogdanlab/gene_sets/), Online Mendelian Inheritance in Man^39^ (https://www.omim.org/), Therapeutic Target Database^42^ (http://db.idrblab.net/ttd/).

When quantifying overlaps between RSS-NET prioritized genes and mouse or Mendelian genes, we used all genes for each GWAS trait. We repeated the overlap analysis under the same significance cutoff (FDR≤ 0.1) after excluding genes that were implicated in the same or later GWAS. Since GWAS-implicated genes overlap significantly with phenotypically-matched mouse and Mendelian genes (median Fisher exact *P* = 7.1 × 10^−7^), we identified fewer discoveries as expected (mouse-human pairs: 26, Mendelian-complex pairs: 4; Supplementary Tables 12-13), but we obtained consistent odds ratio estimates nonetheless (mouse *R* = 0.78, *P* = 8.6 × 10^−73^; Mendelian *R* = 0.89, *P* = 9.0 × 10^−74^; Supplementary Fig. 15).

### Network-induced effect size distribution

We model the total effect of SNP *j* on a given trait, *β_j_*, as

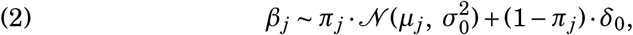

where *π_j_* denotes the probability that SNP *j* is associated with the trait (*β_j_* = 0), 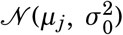 denotes a normal distribution with mean *μ_j_* and variance 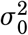 specifying the effect size of a trait-associated SNP *j*, and *δ*_0_ denotes point mass at zero (*β_j_* = 0).

We model the trait-association probability *π_j_* as

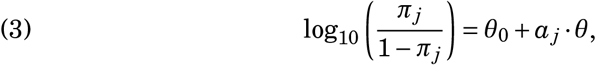

where *θ*_0_ < 0 captures the genome-wide background proportion of trait-associated SNPs, *θ* > 0 reflects the increase in probability, on the log10-odds scale, that a SNP near network genes and REs is trait-associated, and *α_j_* reflects the proximity of SNP *j* to a network. Following previous analyses^15,16,24^, we let *α_j_* = 1 if SNP *j* is within 100 kb of any member gene (TF, TG) or RE for a given network. The idea of (3) is that if a cell type or tissue plays an important role in a trait then genetic associations may occur more often in SNPs involved in the corresponding network and REs than expected by chance.

We model the mean effect size *μ_j_* as

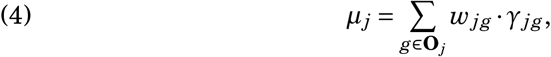

where **O**_*j*_ is the set of all nearby or distal genes contributing to the total effect of SNP *j, w_jg_* measures the relevance between SNP *j* and gene *g*, and *γ_jg_* denotes the effect of SNP *j* on a trait due to gene *g*. We note that (4) provides a generic model to decompose the total effect of a SNP into effects of genes through {**O**_*j*_, *w_jg_*}.

Here we use a TF-TG regulatory network to specify {**O**_*j*_, *w_jg_*} in (4):

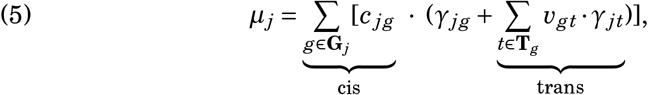

where **G**_*j*_ is the set of all genes within 1 Mb window of SNP *j* (a standard cis-eQTL window size^9,10,58^), *c_jg_* measures the relative impact of a SNP *j* on gene *g*, **T**_*g*_ is the set of all genes directly regulated by TF *g* in a given network (**T**_*g*_ is empty if gene *g* is not a TF), and *v_gt_* measures the relative impact of a TF g on its TG *t*. Since a genome-wide analysis typically involves many SNPs and genes, we fix {**T**_*g*_, *v_gt_, c_jg_*} to ensure the identifiability of (5). We use inferred edges and weights of a context-specific TF-TG network^20,29^ to specify **T**_*g*_ and *v_gt_* respectively. We use context-matched cis-eQTL^9,10,58^ to specify *c_jg_* (Supplementary Notes and Tables 17-18). The idea of (5) is that the true effect of a SNP may fan out through some regulatory network of multiple (nearby or distal) genes to affect the trait^22^.

We model *γ_jg_*, the random effect of SNP *j* due to gene *g* as

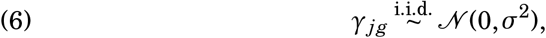

where the SNP-level subscript *j* in *γ_jg_* ensures the exchangeability of *β_j_* in (2); see Supplementary Notes. We use a constant variance *σ*^2^ in (6) for computational convenience. (One could potentially improve (6) by letting *σ*^2^ depend on functional annotations^13,27^ of SNP *j* and/or context-specific expression^14–16^ of gene *g*, though possibly at higher computational cost.)

Combining (2), (4) and (6) yields a variance decomposition for SNP effect:

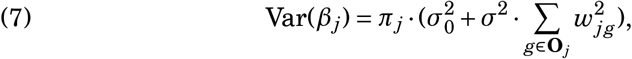

We hypothesize that (7) may provides an alternative approach to heritability analyses^13,24,27^ and we plan to investigate this idea elsewhere.

### Bayesian hierarchical modeling

Consider a GWAS with n unrelated individuals measured on *p* SNPs. In practice we do not know the true SNP-level effects *β*:= (*β*_1_,..., *β_p_*)’ in (2), but we can infer them from GWAS summary statistics and LD estimates. Specifically, we perform Bayesian inference for *β* by combining the network-based prior (2)-(6) with the RSS likelihood^25^:

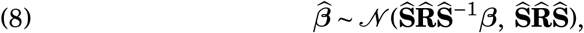

where 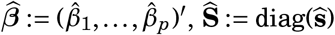 is a *p × p* diagonal matrix with diagonal elements being 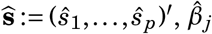 and 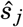 are estimated single-SNP effect size of each SNP *j* and its standard error from the GWAS, and 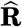 is the *p × p* LD matrix estimated from a reference panel with ancestry matching the GWAS.

RSS-NET, defined by the hierarchical model (2)-(6) and (8), consists of four unknown hyper-parameters: 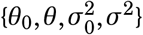. To specify hyper-priors, we first introduce two free parameters {*η, ρ*} ∈ [0,1] to re-parameterize 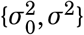:

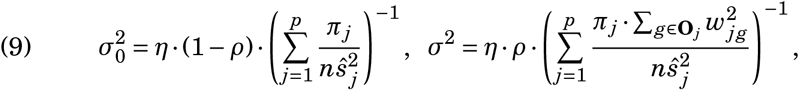

where, roughly, *η* represents the proportion of the total phenotypic variation explained by *p* SNPs, and *ρ* represents the proportion of total genetic variation explained by network annotations {**O**j, *w_jg_*}. Because 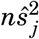 is roughly the ratio of phenotype variance to genotype variance, (9) ensures that SNP effects (*β*) do not rely on sample size *n* and have the same measurement unit as the trait. See Supplementary Notes for a rigorous derivation of (9).

We then place independent uniform grid priors on {*θ*_0_, *θ, η, ρ*} (Supplementary Table 19). These simple hyper-priors produce accurate posterior estimates for hyper-parameters in simulations (Supplementary Fig. 16). RSS-NET results are robust to grid choice in both simulated and real data (Supplementary Fig.s 17-18). (If one had specific information about {*θ*_0_,*θ,η,ρ*} in a given setting then this could be incorporated in the hyper-priors.)

### Network enrichment

To assess whether a regulatory network is enriched for genetic associations with a trait, we evaluate a Bayes factor (BF):

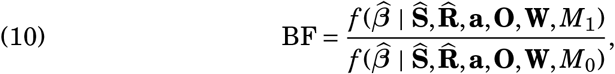

where *f* (·) denotes probability densities, **a** is defined in (3), {**O,W**} are defined in (4), *M*_1_ denotes the enrichment model where *θ* > 0 or *σ*^2^ > 0, and *M*_0_ denotes the baseline model where *θ* = 0 and *σ*^2^ = 0. The observed data are BF times more likely under *M*_1_ than under *M*_0_, and so the larger the BF, the stronger evidence for network enrichment. See Supplementary Notes for details of computing BF. To compute BFs used in Figure 5c, we replace *M*_1_ in (10) with three restricted enrichment models (*M*_11_, *M*_12_, *M*_13_). Unless otherwise specified, all BFs reported in this work are based on *M*_1_.

Given a BF cutoff false positive rates vary considerably across genetic architectures and enrichment patterns in simulations (Supplementary Table 20). As the genetic basis of most complex traits remains unknown, we find it impractical to fix some significance threshold. Instead we recommend an adaptive approach. Specifically, for a given GWAS we run RSS-NET on a near-gene control network containing all genes as nodes and no edges (i.e., *α_j_* = 1 for all SNPs within 100 kb of any gene and *v_gt_* = 0 for all TF-TG pairs), and we use the resulting BF as the enrichment threshold in this GWAS. As shown in our analyses, this approach has three main advantages. First, it is adaptive to study heterogeneity such as differences in traits and sample sizes (Supplementary Table 2). Second, it accounts for generic regulatory enrichments of genetic signals residing near genes. Third, it facilitates comparisons with non-Bayesian methods based on P-values (Supplementary Table 3).

### Locus association

To identify association between a locus and a trait, we compute *P*_1_, the posterior probability that at least one SNP in the locus is associated with the trait:

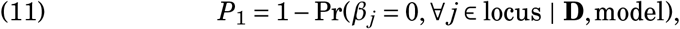

where **D** is a shorthand for the input data of RSS-NET including GWAS summary statistics 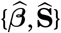, LD estimates 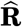 and network annotations {**a, O, W**}. See Supplementary Notes for details of computing *P*_1_. For a locus, 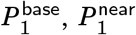 and 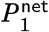 correspond to *P*_1_ evaluated under the baseline model *M*_0_, the enrichment model *M*_1_ for the near-gene control network, and *M*_1_ for a given TF-TG network. In this study we defined a locus as the transcribed region of a gene plus 100 kb up and downstream, and thus we used “locus” and “gene” interchangeably.

For *K* networks with enrichments stronger than the near-gene control, we use Bayesian model averaging (BMA) to compute 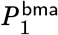 for each locus:

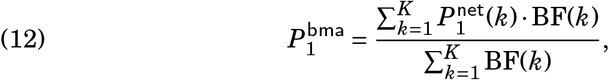

where 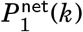 and BF(*k*) are enrichment *P*_1_ and BF for network *k*. The ability to average across networks in (12) is an advantage of our Bayesian framework, because it allows us to assess associations in light of network enrichment without having to select a single enriched network.

In this study we used *P*_1_ ≥ 0.9 as the significance cutoff, yielding a median false-positive rate 1.24 × 10^−4^ in simulations (Supplementary Table 21). We also highlighted genes with 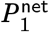 much larger than 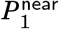 (Fig. 6 and Tables 1–2), because they showcase the influence of tissue-specific regulatory topology on prioritizing genetic associations.

**Fig 6:**
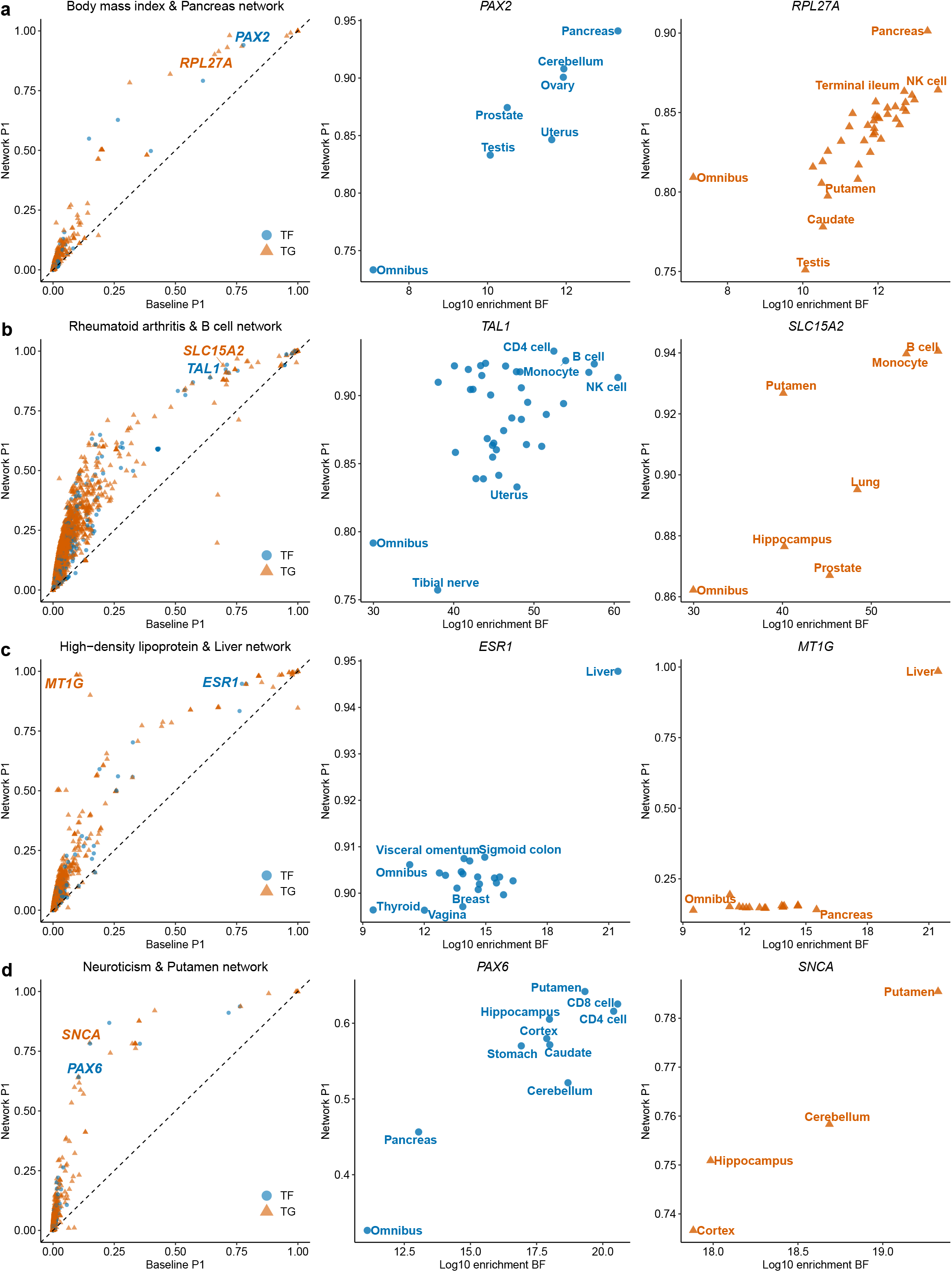
RSS-NET gene prioritization results of select trait-network pairs. In the left column, each dot represents a member gene of a given network. Dashed lines have slope 1 and intercept 0. In the center and right columns, each dot represents a network to which a select gene belongs. Numerical values of *P*_1_ and BF are available online (Data Availability).

### Computation time

The total computational time of RSS-NET to analyze a pair of trait and network is determined by the number of genomewide SNPs analyzed, the size of hyper-parameter grid, and the number of variational iterations till convergence, all of which can vary considerably among studies. It is thus hard to make general statements about computational time. However, to give a specific example, we finished the analysis of 1,032,214 HapMap3 SNPs and liver network for HDL within 12 hours in a standard computer cluster (60 nodes, 8 CPUs and 32 Gb memory per node).

The number of genome-wide SNPs analyzed (*p*) affects the computation time of RSS-NET in two distinct ways. First, the per-iteration complexity of RSS-NET is linear with *p* (Supplementary Notes). Second, a large *p* defines a large optimization problem, often requiring many iterations to converge. To quantify the impact of *p* on computation time, we simulated datasets from different sets of genome-wide common SNPs, analyzed them with RSS-NET on identical computers, and compared the computation time (Supplementary Fig. 9). When *p* increased from 348,965 to 1,030,397, on average the total computation time was four times longer (one-sided Wilcoxon *P* = 8.0 × 10^−132^).

### Simulation overview

To assess the new model for SNP effects (*β*) in RSS-NET, we simulated a large array of correctly- and mis-specified *β* for a given target network. Specifically, we generated “positive” datasets where the underlying *β* was simulated from *M*_1_ for the target network, and “negative” datasets where β was simulated from either *M*_0_ or the following scenarios: (1) random enrichments of near-gene SNPs; (2) random enrichments of near-RE SNPs; (3) MAF-and LD-dependent effect sizes; (4) M_1_ for edge-altered copies of the target network. For a fair comparison in each scenario, we matched positive and negative datasets by i) the number of trait-associated SNPs and ii) proportion of phenotypic variation explained by all SNPs. Simulation details are provided in Supplementary Figures 1-9.

We combined the simulated *β* with genotypes of 348,965 genome-wide SNPs from 1,458 individuals^28^ to simulate phenotypes using an additive multiple-SNP model with Gaussian noise. For simulated individual-level data, we performed the standard single-SNP analysis to generate GWAS summary statistics, on which we compared RSS-NET with external methods.

### External software for benchmarking

This study used the following software to benchmark RSS-NET: RSS-E (https://github.com/stephenslab/rss, accessed October 19,2018), Pascal (https://www2.unil.ch/cbg/index.php?title=Pascal, accessed October 5, 2017) and LDSC with two sets of baseline annotations as covariates (version 1.0.0, https://github.com/bulik/ldsc; baseline model v1.1, https://data.broadinstitute.org/alkesgroup/LDSC0RE/1000G_Phase3_baseline_v1.1_ldscores.tgz; baselineLD model v2.1, https://data.broadinstitute.org/alkesgroup/LDSC0RE/1000G_Phase3_baselineLD_v2.1_ldscores.tgz; accessed November 27, 2018). Versions of all packages and files were up-to-date at the time of analysis.

Given a context-specific TF-TG network, RSS-E and LDSC methods use the same binary SNP-level annotations *{aj*} as defined in (3) of RSS-NET. The interface design of Pascal does not allow direct usage of {*α_j_*}. Here we supplied Pascal program with a GMT file containing all member genes of the network and set SNP-to-gene window sizes as 100 kb (`‘-up=100000 -down=100000’`). In this study all external methods were used with their default setups, and did not include the edge information of a network.

RSS-E outputs the same statistics as RSS-NET (namely, BF and P_1_). Pascal implements two gene scoring methods, maximum-of-*χ*^2^ and sum-of-*χ*^2^, each producing gene-based association *P*-values. Given gene scores, Pascal provides two gene set scoring options, *χ*^2^ approximation and empirical sampling, to produce enrichment *P*-values. LDSC methods output enrichment *P*-values and coefficient *Z*-scores, which produced consistent results in simulations (LDSC-baseline: *R* = 0.98, *P* = 1.2 × 10^−67^; LDSC-baselineLD: *R* = 0.98, *P* = 9.1 × 10^−63^; Supplementary Fig. 19). Due to its higher power shown in simulations (LDSC-baseline: average AUROC increase= 0.012, one-sided *t P* = 4.0× 10^−3^; LDSC-baselineLD: average AUROC increase= 0.023, one-sided *t P* = 1.5 × 10^−5^), we used LDSC enrichment *P*-values throughout this study.

## Data availability

Network files used in this study are available at https://github.com/suwonglab/rss-net. Analysis results of this study are available at https://suwonglab.github.io/rss-net/results. Other data are specified in Methods and Supplementary Notes.

## Code availability

The RSS-NET software is available at https://github.com/suwonglab/rss-net. Tutorials of installing and using RSS-NET are available at https://suwonglab.github.io/rss-net. Results of this study were generated from MATLAB version 9.3.0.713579 (R2017b), on a Linux system with Intel E5-2650V2 2.6 GHz and E5-2640V4 2.4 GHz processors. Other codes are specified in Methods and Supplementary Notes.

## Author contributions

X.Z. and W.H.W. conceived the study. X.Z. developed the methods and implemented the software. X.Z. conducted the simulation experiments. Z.D. provided the 38 regulatory networks. X.Z. performed the data analyses. X.Z. prepared the supplementary materials and online resources. X.Z. wrote the manuscript. X.Z. and W.H.W. revised the manuscript.

## Acknowledgments

This study is supported by Stein Fellowship to X.Z. and NIH grants P50HG007735 and R01HG010359 to W.H.W. This study uses computational resources provided by the Stanford Research Computing Center. This study uses data generated by the WTCCC, 1000 Genomes, ENCODE, GTEx, DICE, eQTLGen and multiple GWAS consortia. We thank them for making their data publicly available (Supplementary Notes). We thank X. He for helpful comments on a draft manuscript.

## Notes

https://suwonglab.github.io/rss-net/

### Competing Interest Statement

The authors have declared no competing interest.

### Summary of Updates

We summarize our revisions by: (1) new simulation studies to illustrate the robustness of RSS-NET; (2) additional replication and comparative analyses of RSS-NET on real data; (3) expanded discussions of methodology intuition and implementation details. The new results are consistent with conclusions in our original manuscript, and the revised texts improve the readability of this work.

