## Supplementary Information for "Modeling Regulatory Network Topology Improves Genome-Wide Analyses of Complex Human Traits"

Xiang Zhu

*The Pennsylvania State University, University Park, PA 16802, USA*

**Summary**. This document contains Supplementary Notes, Figures 1-19 and Tables 1-21 associated with the manuscript entitled *"Modeling regulatory network topology improves genome-wide analyses of complex human traits"* (last edited on 2020-12-08).

### Supplementary Notes

#### Computations underlying RSS-NET

RSS-NET is given by the following Bayesian model:

$$\begin{aligned}
\widehat{\boldsymbol{\beta}} &\sim \mathcal{N}(\widehat{\mathbf{S}}\widehat{\mathbf{R}}\widehat{\mathbf{S}}^{-1}\boldsymbol{\beta},\ \widehat{\mathbf{S}}\widehat{\mathbf{R}}\widehat{\mathbf{S}}), && (1)\\
\beta_j &\sim \pi_j \cdot \mathcal{N}(\mu_j,\ \sigma_0^2) + (1-\pi_j)\cdot\delta_0, && (2)\\
\pi_j &= 1\,/\,[1+10^{-(\theta_0+a_j\cdot\theta)}], && (3)\\
\mu_j &= \textstyle\sum_{g\in\mathbf{O}_j} w_{jg}\cdot\gamma_{jg}, && (4)\\
\gamma_{jg} &\sim \mathcal{N}(0,\ \sigma^2), && (5)
\end{aligned}$$

where $\mathcal{N}(\boldsymbol{\mu},\ \boldsymbol{\Sigma})$ denotes a normal distribution with mean vector $\boldsymbol{\mu}$ and covariance matrix $\boldsymbol{\Sigma}$, $\widehat{\boldsymbol{\beta}} := (\hat{\beta}_1,\ldots,\hat{\beta}_p)'$ is a $p\times 1$ vector, $\widehat{\mathbf{S}} := \text{diag}\{(\hat{s}_1,\ldots,\hat{s}_p)'\}$ is a $p\times p$ diagonal matrix, $\{\hat{\beta}_j,\hat{s}_j\}$ are single-SNP effect estimate and its standard error for each SNP $j$ in a GWAS, $\widehat{\mathbf{R}}$ is a $p\times p$ LD matrix estimated from a reference panel with ancestry matching the GWAS, $\boldsymbol{\beta} := (\beta_1,\ldots,\beta_p)'$ is a $p\times 1$ vector, $\beta_j$ is the true effect of each SNP $j$, $\pi_j$ denotes the prior probability that $\beta_j \neq 0$, $\mu_j$ denotes the prior expected value of $\beta_j$ conditioning on $\beta_j \neq 0$, $\delta_0$ denotes point mass at zero, $\{a_j,\mathbf{O}_j,w_{jg}\}$ are known annotations derived from a given regulatory network (see main text Equations 3-5), $\gamma_{jg}$ denotes the random effect of SNP $j$ due to gene $g$, and $\{\theta_0,\theta,\sigma_0,\sigma\}$ are hyper-parameters.

To fit the RSS-NET model, we first integrate out $\gamma_{jg}$:

$$\begin{aligned}
\widehat{\boldsymbol{\beta}} &\sim \mathcal{N}(\widehat{\mathbf{S}}\widehat{\mathbf{R}}\widehat{\mathbf{S}}^{-1}\boldsymbol{\beta},\ \widehat{\mathbf{S}}\widehat{\mathbf{R}}\widehat{\mathbf{S}}), && (6)\\
\beta_j &\sim \pi_j \cdot \mathcal{N}(0,\ \sigma_j^2) + (1-\pi_j)\cdot\delta_0, && (7)\\
\pi_j &= 1\,/\,[1+10^{-(\theta_0+a_j\theta)}], && (8)\\
\sigma_j^2 &= \sigma_0^2 + \sigma^2\cdot\textstyle\sum_{g\in\mathbf{O}_j} w_{jg}^2. && (9)
\end{aligned}$$

We then write the posterior distribution of $\boldsymbol{\beta}$ as

$$p(\boldsymbol{\beta}\mid\mathbf{D}) = \int p(\boldsymbol{\beta}\mid\mathbf{D},\theta_0,\theta,\sigma_0,\sigma)\,p(\theta_0,\theta,\sigma_0,\sigma\mid\mathbf{D})\,\mathrm{d}\theta_0\mathrm{d}\theta\mathrm{d}\sigma_0\mathrm{d}\sigma, \qquad (10)$$

where $\mathbf{D}$ is a shorthand for the input data of RSS-NET including GWAS summary statistics $\{\widehat{\boldsymbol{\beta}},\widehat{\mathbf{S}}\}$, LD estimates $\widehat{\mathbf{R}}$ and network annotations $\{\mathbf{a},\mathbf{O},\mathbf{W}\}$ (see main text Equations 3-5). For a given set of $\{\theta_0,\theta,\sigma_0,\sigma\}$, we use the mean-field variational approximation algorithm from RSS-E (Zhu and Stephens 2018) to estimate $p(\boldsymbol{\beta} \mid \mathbf{D},\theta_0,\theta,\sigma_0,\sigma)$:

$$p(\boldsymbol{\beta} \mid \mathbf{D},\theta_0,\theta,\sigma_0,\sigma) \approx \prod_{j=1}^{p} q_j^\star(\beta_j \mid \mathbf{D},\theta_0,\theta,\sigma_0,\sigma), \tag{11}$$

$$q_j^\star(\beta_j \mid \mathbf{D},\theta_0,\theta,\sigma_0,\sigma) = \alpha_j^\star \cdot \mathcal{N}(\beta_j;\, \nu_j^\star,(\tau_j^\star)^2)+(1-\alpha_j^\star)\cdot\delta_0(\beta_j), \tag{12}$$

where the optimal variational parameters $\{\alpha_j^\star,\nu_j^\star,\tau_j^\star\}$ are given by:

$$(\tau_j^\star)^2 = \frac{1}{\hat{s}_j^{-2}+\sigma_j^{-2}}, \tag{13}$$

$$\nu_j^\star = (\tau_j^\star)^2\cdot\left(\frac{\hat{\beta}_j}{\hat{s}_j^2}-\sum_{i\neq j}\frac{\widehat{\mathbf{R}}_{ij}\alpha_i^\star\nu_i^\star}{\hat{s}_i\hat{s}_j}\right), \tag{14}$$

$$\frac{\alpha_j^\star}{1-\alpha_j^\star} = \frac{\pi_j}{1-\pi_j}\cdot\frac{\tau_j^\star}{\sigma_j}\cdot\exp\left\{\frac{(\nu_j^\star)^2}{2(\tau_j^\star)^2}\right\}. \tag{15}$$

The per-iteration complexity of the coordinate descent algorithm (13)-(15) is linear with $p$, the number of genome-wide SNPs analyzed.

We use $F^\star(\mathbf{D},\theta_0,\theta,\sigma_0,\sigma)$, the variational lower bound corresponding to the optimal variational parameters $\{\alpha_j^\star,\nu_j^\star,\tau_j^\star\}$ in (13)-(15) to approximate the log marginal likelihood $\log p(\widehat{\boldsymbol{\beta}} \mid \widehat{\mathbf{S}},\widehat{\mathbf{R}},\mathbf{a},\mathbf{O},\mathbf{W},\theta_0,\theta,\sigma_0,\sigma)$:

$$\begin{aligned}F^\star(\mathbf{D},\theta_0,\theta,\sigma_0,\sigma) =& F_0(\mathbf{D})+\widehat{\boldsymbol{\beta}}'\widehat{\mathbf{S}}^{-2}\mathrm{E}_{q^\star}(\boldsymbol{\beta})-\frac{1}{2}\mathrm{E}'_{q^\star}(\boldsymbol{\beta})\widehat{\mathbf{S}}^{-1}\widehat{\mathbf{R}}\widehat{\mathbf{S}}^{-1}\mathrm{E}_{q^\star}(\boldsymbol{\beta})-\frac{1}{2}\sum_{j=1}^{p}\frac{\mathrm{Var}_{q^\star}(\beta_j)}{\hat{s}_j^2}\\ &-\sum_{j=1}^{p}\alpha_j^\star\log\left(\frac{\alpha_j^\star}{\pi_j}\right)-\sum_{j=1}^{p}(1-\alpha_j^\star)\log\left(\frac{1-\alpha_j^\star}{1-\pi_j}\right)\\ &+\sum_{j=1}^{p}\frac{\alpha_j^\star}{2}\left\{1+\log\left[\frac{(\tau_j^\star)^2}{\sigma_j^2}\right]-\frac{(\tau_j^\star)^2+(\nu_j^\star)^2}{\sigma_j^2}\right\},\end{aligned}$$

$F_0(\mathbf{D})=-[\log|2\pi\cdot\widehat{\mathbf{S}}\widehat{\mathbf{R}}\widehat{\mathbf{S}}|+\widehat{\boldsymbol{\beta}}'(\widehat{\mathbf{S}}\widehat{\mathbf{R}}\widehat{\mathbf{S}})^{-1}\widehat{\boldsymbol{\beta}}]\,/\,2$, $\mathrm{E}_{q^\star}(\boldsymbol{\beta})=(\mathrm{E}_{q^\star}(\beta_1),\ldots,\mathrm{E}_{q^\star}(\beta_p))'$, $\mathrm{E}_{q^\star}(\beta_j)=\alpha_j^\star\nu_j^\star$ and $\mathrm{Var}_{q^\star}(\beta_j)=\alpha_j^\star[(\tau_j^\star)^2+(\nu_j^\star)^2]-(\alpha_j^\star\nu_j^\star)^2$. Of note, RSS-NET and RSS-E have the same posterior computation scheme if we set $\sigma_j^2=\sigma_\beta^2$ for all $j=1,\ldots,p$ in (13)-(15) and $F^\star$.

Finally, we approximate the joint posterior of hyper-parameters $p(\theta_0,\theta,\sigma_0,\sigma \mid \mathbf{D})$ as:

$$p(\theta_0,\theta,\sigma_0,\sigma \mid \mathbf{D}) \approx \omega^\star(\theta_0,\theta,\sigma_0,\sigma) \propto \exp\{F^\star(\mathbf{D},\theta_0,\theta,\sigma_0,\sigma)\}, \tag{16}$$

since we use independent uniform grid hyper-priors (**Supplementary Table 19**).

#### Bayes factor for network enrichments

To assess whether a regulatory network is enriched for genetic associations with a trait, we evaluate the following Bayes factor (BF):

$$\mathrm{BF}=\frac{p(\widehat{\boldsymbol{\beta}} \mid \widehat{\mathbf{S}},\widehat{\mathbf{R}},\mathbf{a},\mathbf{O},\mathbf{W},M_1)}{p(\widehat{\boldsymbol{\beta}} \mid \widehat{\mathbf{S}},\widehat{\mathbf{R}},\mathbf{a},\mathbf{O},\mathbf{W},M_0)}, \tag{17}$$

where $M_1$ denotes the enrichment model where $\theta > 0$ or $\sigma^2 > 0$, and $M_0$ denotes the baseline model where $\theta = 0$ and $\sigma^2 = 0$. To compute BF (17), we approximate intractable marginal likelihoods by corresponding variational lower bounds $F^\star$:

$$\begin{aligned}\text{BF} &= \frac{\int p(\widehat{\boldsymbol{\beta}} \mid \widehat{\mathbf{S}}, \widehat{\mathbf{R}}, \mathbf{a}, \mathbf{O}, \mathbf{W}, \theta_0, \theta, \sigma_0, \sigma) p(\theta_0) p(\theta) p(\sigma_0) p(\sigma) \mathrm{d}\theta \mathrm{d}\theta_0 \mathrm{d}\sigma_0 \mathrm{d}\sigma}{\int p(\widehat{\boldsymbol{\beta}} \mid \widehat{\mathbf{S}}, \widehat{\mathbf{R}}, \mathbf{a}, \mathbf{O}, \mathbf{W}, \theta_0, \theta = 0, \sigma_0, \sigma = 0) p(\theta_0) p(\sigma_0) \mathrm{d}\theta_0 \mathrm{d}\sigma_0} \\ &\approx \frac{n_1^{-1} \sum_{s=1}^{n_1} \exp\{F^\star(\mathbf{D}, \theta_0^{(s)}, \theta^{(s)}, \sigma_0^{(s)}, \sigma^{(s)})\}}{n_0^{-1} \sum_{t=1}^{n_0} \exp\{F^\star(\mathbf{D}, \theta_0^{(t)}, \theta = 0, \sigma_0^{(t)}, \sigma = 0)\}},\end{aligned} \tag{18}$$

where $\{\theta_0^{(s)}, \theta^{(s)}, \sigma_0^{(s)}, \sigma^{(s)}\}$ and $\{\theta_0^{(t)}, \sigma_0^{(t)}\}$ are pre-defined grids (**Supplementary Table 19**).

#### Posterior probability for genetic associations

To identify association between a locus and a trait, we compute $P_1$, the posterior probability that at least one SNP in the locus is associated with the trait:

$$P_1 = 1 - \Pr(\beta_j = 0, \forall j \in \text{locus} \mid \mathbf{D}, \text{model}), \tag{19}$$

where the "model" here can be the baseline model $M_0$ (yielding $P_1^{\text{base}}$), the enrichment model $M_1$ for the near-gene control network (yielding $P_1^{\text{near}}$) and a given network (yielding $P_1^{\text{net}}$). Given a grid $\{\theta_0^{(s)}, \theta^{(s)}, \sigma_0^{(s)}, \sigma^{(s)}\}$, $P_1$ is estimated as

$$\begin{aligned} P_1 &= 1 - \int \Pr(\beta_j = 0, \ \forall j \in \text{locus} \mid \mathbf{D}, \theta_0, \theta, \sigma_0, \sigma) p(\theta_0, \theta, \sigma_0, \sigma \mid \mathbf{D}) \mathrm{d}\theta \mathrm{d}\theta_0 \mathrm{d}\sigma_0 \mathrm{d}\sigma \\ &\approx 1 - \sum_{s=1}^{n_1} \prod_{j \in \text{locus}} \left[1 - \alpha_j^\star(\theta_0^{(s)}, \theta^{(s)}, \sigma_0^{(s)}, \sigma^{(s)})\right] \cdot \omega^\star(\theta_0^{(s)}, \theta^{(s)}, \sigma_0^{(s)}, \sigma^{(s)}). \end{aligned} \tag{20}$$

#### Regulatory network as a bipartite graph

In this study a regulatory network is a directed bipartite graph $\{\mathbf{V}_{\text{TF}}, \mathbf{V}_{\text{TG}}, \mathbf{E}_{\text{TF}\to\text{TG}}\}$, where $\mathbf{V}_{\text{TF}}$ denotes the node set of transcription factors (TFs), $\mathbf{V}_{\text{TG}}$ denotes the node set of target genes (TGs), and $\mathbf{E}_{\text{TF}\to\text{TG}}$ denotes the set of directed TF-to-TG edges, summarizing how TFs regulate TGs through regulatory elements (REs). Each edge has a weight between 0 and 1, measuring the relative regulation strength of a TF on a TG.

Below is a small subset of B cell regulatory network (`https://github.com/suwonglab/rss-net/tree/master/data`). In this example, $\mathbf{V}_{\text{TF}}$ corresponds to the column `TF`, $\mathbf{V}_{\text{TG}}$ corresponds to the column `TG`, $\mathbf{E}_{\text{TF}\to\text{TG}}$ corresponds to all rows of `TF` and `TG`, and the edge weights are specified by the column `Score`.

```
        TF     TG    Score
1:    RARG EEF1A1 0.723037
2:    SOX9   CD74 0.690921
3:    EBF1  FXYD5 0.659009
4:    PAX5   CD44 0.704366
5:  TRIM28   GSAP 0.629798
6:  POU6F1  RPS3A 0.632690
```

In this study, for a given TF, we only consider the downstream effects of all genes that are **directly** regulated by this TF (see main text Figure 1c and Equation 5). For example, it is biologically possible that a TF 1 regulates a target gene which is also a TF, say TF 2; when specifying the RSS-NET SNP-level effect size distribution, we only incorporate the direct downstream effect of TF 1 on TF 2, and ignore all other indirect downstream effect of TF 1 on genes that are only directly regulated by TF 2.

#### Quantify *cis* impact of a SNP on a gene

To specify $c_{jg}$, the relative impact of a *cis* SNP $j$ on a gene $g$ (main text Equation 5), we use a simple approach based on published *cis* expression quantitative trait loci (eQTL). Specifically, if a pair of SNP $j$ and gene $g$ is available in an eQTL database, we define

$$c_{jg} = 1 + \sqrt{z_{jg}^2 / (z_{jg}^2 + n_{jg})}, \tag{21}$$

where $n_{jg}$ is the number of individuals with both genotype of SNP $j$ and expression of gene $g$ available and $z_{jg}$ is the single-SNP $z$-score measuring the marginal association between genotype of SNP $j$ and expression of gene $g$ across $n_{jg}$ individuals; if the pair does not have eQTL association data available, we set $c_{jg} = 1$. When deriving $c_{jg}$ from *cis*-eQTL data, we consider all available SNP-gene pairs, not just the significant ones.

In this study we specify $c_{jg}$ in a context-matched manner. For example, if we analyze the regulatory network derived from liver samples, we also use *cis*-eQTL data from liver samples to define $c_{jg}$. For networks of 5 brain regions and 27 non-brain tissues, we use the tissue-matched *cis*-eQTL data from GTEx (https://storage.googleapis.com/gtex_analysis_v7/single_tissue_eqtl_data/GTEx_Analysis_v7_eQTL_all_associations.tar.gz, accessed April 7, 2019). For networks of 5 immune cells, we use the cell-type-matched *cis*-eQTL data from DICE (http://downloads.dice-database.org/downloads/DICE_DB_1/eqtl/unfiltered/, accessed July 3, 2019). For the "omnibus" network, we use the *cis*-eQTL data of blood samples from the eQTLGen Consortium (http://www.eqtlgen.org/cis-eqtls.html, accessed May 17, 2019). Summary statistics of $c_{jg}$ are provided in **Supplementary Tables 17-18**.

The small sample sizes of eQTL studies make it hard to produce reliable model-based estimates of $c_{jg}$. Hence, we keep our specification of $c_{jg}$ deliberately simple. We acknowledge that (21) could be potentially improved, although we have not fully investigated it here. Due to the modular design of RSS-NET, once improved estimates of $c_{jg}$ become available, they can be incorporated into RSS-NET in a straightforward manner.

#### Explain SNP-level subscript $j$ in $\gamma_{jg}$

In this work we model $\gamma_{jg}$, the random effect of SNP $j$ due to gene $g$ as

$$\gamma_{jg} \overset{\text{i.i.d.}}{\sim} \mathcal{N}(0, \sigma^2).$$

We emphasize that the SNP-level subscript $j$ in $\gamma_{jg}$ ensures the exchangeability of $\beta_j$ required by (2), as shown in the following example. Suppose there are two nearby SNPs 1 and 2, and their effects are sums of random effects of genes 1 and 2. If we ignore the SNP-level subscript $j$ and use the $\gamma_g$ notation, then

$$\begin{aligned}
\beta_1 &= 0.18 \cdot \gamma_1 + 0.36 \cdot \gamma_2, \\
\beta_2 &= 0.36 \cdot \gamma_1 + 0.72 \cdot \gamma_2, \\
\gamma_g &\overset{\text{i.i.d.}}{\sim} \mathcal{N}(0, \sigma^2), \; g = 1, 2.
\end{aligned}$$

This is inconsistent with the independent prior (2) placed on $\beta_j$, because under this notation, $\beta_2 = 2\beta_1$ and the correlation of $\beta_1$ and $\beta_2$ is exactly 1. In contrast, this inconsistency disappears if we use the $\gamma_{jg}$ notation:

$$\beta_1 = 0.18 \cdot \gamma_{11} + 0.36 \cdot \gamma_{12},$$
$$\beta_2 = 0.36 \cdot \gamma_{21} + 0.72 \cdot \gamma_{22},$$
$$\gamma_{jg} \overset{\text{i.i.d.}}{\sim} \mathcal{N}(0,\ \sigma^2),\ j = 1,2,\ g = 1,2.$$

Now $\beta_1$ and $\beta_2$ are independent, because $\gamma_{jg}$'s are all i.i.d for $j = 1,2$ and $g = 1,2$.

#### Induced prior distribution for $\sigma_0^2$ and $\sigma^2$

Here we assume a multiple regression model for the phenotype-genotype relationship:

$$\mathbf{y} = \mathbf{X}\boldsymbol{\beta} + \boldsymbol{\epsilon}, \tag{22}$$

where $\mathbf{y}$ is an $n \times 1$ centered vector of phenotype, $\mathbf{X}$ is an $n \times p$ column-centered matrix of genotype, $\boldsymbol{\beta}$ is a $p \times 1$ vector of multiple regression coefficients (i.e. SNP-level effect sizes), and $\boldsymbol{\epsilon}$ is an independent error term. Let $\sigma_{x,j}^2$ denote the population variance of genotype for SNP $j$, $j = 1, \ldots, p$, and let $\sigma_y^2$ denote the population variance of phenotype.

PROPOSITION 1. If SNP-level effect sizes $\boldsymbol{\beta}$ follow the prior distribution specified in RSS-NET (2), (4) and (5), then, for all $i, j = 1, \ldots, p$,

$$\mathrm{E}(\beta_j) = 0,\ \ \mathrm{Var}(\beta_j) = \pi_j \left(\sigma_0^2 + \sigma^2 \cdot \textstyle\sum_{g \in \mathbf{O}_j} w_{jg}^2\right),\ \ \mathrm{Cov}(\beta_i,\ \beta_j) = 0. \tag{23}$$

PROOF. This is a direct application of the law of total expectation. ■

PROPOSITION 2. $\mathrm{E}[V(\mathbf{X}\boldsymbol{\beta})] = \mathrm{E}^{\text{base}}[V(\mathbf{X}\boldsymbol{\beta})] + \mathrm{E}^{\text{net}}[V(\mathbf{X}\boldsymbol{\beta})]$, where

$$\mathrm{E}^{\text{base}}[V(\mathbf{X}\boldsymbol{\beta})] = \sigma_0^2 \cdot \sum_{j=1}^{p} \pi_j \cdot \mathrm{E}[V(\mathbf{X}_j)], \tag{24}$$

$$\mathrm{E}^{\text{net}}[V(\mathbf{X}\boldsymbol{\beta})] = \sigma^2 \cdot \sum_{j=1}^{p} \pi_j \cdot \left(\textstyle\sum_{g \in \mathbf{O}_j} w_{jg}^2\right) \cdot \mathrm{E}[V(\mathbf{X}_j)], \tag{25}$$

$\mathbf{X}_j$ is the $j$th column of $\mathbf{X}$, $j = 1, \ldots, p$, and $V(\mathbf{d})$ denotes the sample variance of a vector $\mathbf{d}$.

PROOF. Define a $p \times p$ diagonal matrix $\boldsymbol{\Sigma}_\beta$ where the $j$th diagonal entry is $\mathrm{Var}(\beta_j)$ in PROPOSITION 1. It suffices to notice that $V(\mathbf{X}\boldsymbol{\beta}) = n^{-1}\boldsymbol{\beta}'\mathbf{X}'\mathbf{X}\boldsymbol{\beta}$, and,

$$\begin{aligned} \mathrm{E}[V(\mathbf{X}\boldsymbol{\beta})] &= \mathrm{E}\{\mathrm{E}[V(\mathbf{X}\boldsymbol{\beta}) \mid \mathbf{X}]\} = \mathrm{E}[\mathrm{trace}(n^{-1}\mathbf{X}'\mathbf{X} \cdot \boldsymbol{\Sigma}_\beta)], \\ &= \mathrm{E}\left[\sum_{j=1}^{p} n^{-1}\mathbf{X}_j'\mathbf{X}_j \cdot \mathrm{Var}(\beta_j)\right] = \sum_{j=1}^{p} \mathrm{Var}(\beta_j) \cdot \mathrm{E}[V(\mathbf{X}_j)]. \end{aligned} \tag{26}$$

Use PROPOSITION 1 to complete the proof. ■

REMARK 1. PROPOSITION 2 shows that the total genetic variation $\mathrm{E}[V(\mathbf{X}\boldsymbol{\beta})]$ can be decomposed into a component (24) that is shared by all $p$ SNPs and a component (25) that is due to network annotations $\{\mathbf{O}_j, w_{jg}\}$.

PROPOSITION 3. Let $ns_j^2 = \sigma_y^2/\sigma_{x,j}^2$, for $j = 1,\dots,p$. If $\{\sigma_0^2, \sigma^2\}$ are re-parameterized as

$$\sigma_0^2 = \eta \cdot (1-\rho) \cdot \left( \sum_{j=1}^{p} \frac{\pi_j}{ns_j^2} \right)^{-1}, \quad \sigma^2 = \eta \cdot \rho \cdot \left( \sum_{j=1}^{p} \frac{\pi_j \cdot \sum_{g \in \mathbf{O}_j} w_{jg}^2}{ns_j^2} \right)^{-1}, \tag{27}$$

then

$$\eta = \frac{\mathrm{E}[V(\mathbf{X}\boldsymbol{\beta})]}{\mathrm{E}[V(\mathbf{y})]}, \quad \rho = \frac{\mathrm{E}^{\mathrm{net}}[V(\mathbf{X}\boldsymbol{\beta})]}{\mathrm{E}[V(\mathbf{X}\boldsymbol{\beta})]}. \tag{28}$$

PROOF. It suffices to show that, for all $j = 1,\dots,p$,

$$ns_j^2 = \frac{\sigma_y^2}{\sigma_{x,j}^2} = \frac{\mathrm{E}[V(\mathbf{y})]}{\mathrm{E}[V(\mathbf{X}_j)]}. \tag{29}$$

Use PROPOSITION 2 to complete the proof. ■

REMARK 2. PROPOSITION 3 provides the mathematical basis to interpret $\{\eta, \rho\}$. Specifically, $\eta$ represents the proportion of the total phenotypic variation explained by $p$ SNPs, and $\rho$ represents the proportion of total genetic variation explained by network annotations $\{\mathbf{O}_j, w_{jg}\}$.

REMARK 3. Since $ns_j^2 = \sigma_y^2/\sigma_{x,j}^2$, where $\sigma_y^2$ is the population variance of phenotype and $\sigma_{x,j}^2$ is the population variance of genotype at SNP $j$, this re-parameterization based on $\{\eta, \rho\}$ ensures that the induced-prior (27) of $\{\sigma_0^2, \sigma^2\}$ does not depend on GWAS sample size $n$, and the resulting genetic effect sizes ($\boldsymbol{\beta}$) have the same measurement unit as the phenotype.

REMARK 4. The induced prior (27) of $\{\sigma_0^2, \sigma^2\}$ contains a quantity $ns_j^2 = \sigma_y^2/\sigma_{x,j}^2$ that depends on unknown population parameters $\{\sigma_y^2, \sigma_{x,j}^2\}$. As shown in Zhu and Stephens (2017), $ns_j^2$ can be reliably estimated from GWAS summary data: $ns_j^2 \approx n\hat{s}_j^2$. In practice, we replace the unknown $ns_j^2$ in (27) with known $n\hat{s}_j^2$ (see main text Equation 9).

#### Acknowledgments and data sources

This study uses data generated by the Wellcome Trust Case Control Consortium, 1000 Genomes Project, ENCODE Consortium, GTEx Project, DICE Project, eQTLGen Consortium, and multiple GWAS consortia. We thank them for making their data publicly available. Detailed acknowledgments and data sources are listed below.

- **Wellcome Trust Case Control Consortium (WTCCC).** This study use individual-level genotype data generated by WTCCC to design simulations. A full list of the investigators who contributed to the generation of the data is available from `https://www.wtccc.org.uk/`. The WTCCC data are available at the European Genome-phenome Archive (`https://www.ebi.ac.uk/ega/`).

- **1000 Genomes Project.** This study uses haplotypes of individuals with European ancestry from the 1000 Genomes Project Phase 3 to estimate LD. The 1000 Genomes Phase 3 haplotype data have been contributed by 1000 Genomes investigators and have been downloaded from `ftp://ftp.1000genomes.ebi.ac.uk/vol1/ftp/release/20130502/`.

- **Encyclopedia of DNA Elements (ENCODE) Consortium.** This study uses gene expression and chromatin accessibility data generated by the ENCODE Consortium and the ENCODE production laboratories. The ENCODE data are available at the ENCODE portal (https://www.encodeproject.org/).

- **Genotype-Tissue Expression (GTEx) Project.** This study uses *cis*-eQTL data generated by the GTEx Project. The GTEx Project was supported by the Common Fund of the Office of the Director of the National Institutes of Health, and by NCI, NHGRI, NHLBI, NIDA, NIMH, and NINDS. The GTEx data are available at the GTEx Portal (https://gtexportal.org/home/).

- **Database of Immune Cell Expression, Expression quantitative trait loci and Epigenomics (DICE) Project.** This study uses *cis*-eQTL data generated by the DICE Project. The DICE Project was supported by NIH R24AI108564. The DICE data are available at https://dice-database.org/.

- **eQTLGen Consortium.** This study uses *cis*-eQTL data generated by the eQTLGen Consortium. The eQTLGen data are available at http://www.eqtlgen.org/.

- **Genetic Investigation of ANthropometric Traits (GIANT) Consortium.** GWAS summary statistics on adult human height (Wood et al. 2014), body mass index (Locke et al. 2015) and body fat distribution (Shungin et al. 2015) have been contributed by GIANT investigators and have been downloaded from http://portals.broadinstitute.org/collaboration/giant.

- **Psychiatric Genomics Consortium (PGC).** GWAS summary statistics on schizophrenia (Ripke et al. 2014) have been contributed by PGC investigators and have been downloaded from http://www.med.unc.edu/pgc.

- **International Inflammatory Bowel Disease Genetics Consortium (IIBDGC).** GWAS summary statistics on inflammatory bowel disease, Crohn's disease and ulcerative colitis (Liu et al. 2015) have been contributed by IIBDGC investigators and have been downloaded from https://www.ibdgenetics.org.

- **Coronary ARtery DIsease Genome wide Replication and Meta-analysis (CARDIoGRAM) plus The Coronary Artery Disease (C4D) Genetics (CARDIoGRAMplusC4D) Consortium.** GWAS summary statistics on coronary artery disease and myocardial infarction (Nikpay et al. 2015) have been contributed by CARDIoGRAMplusC4D investigators and downloaded from http://www.cardiogramplusc4d.org/.

- **GWAS summary statistics of heart rate.** GWAS summary statistics on heart rate have been contributed by authors of Den Hoed et al. (2013) and have been downloaded from https://walker05.u.hpc.mssm.edu/.

- **International Genomics of Alzheimer's Project (IGAP).** GWAS summary statistics on Alzheimer's disease (Lambert et al. 2013) have been contributed by IGAP investigators and have been downloaded from http://web.pasteur-lille.fr/en/recherche/u744/igap/igap_download.php. We thank the International Genomics of Alzheimer's Project (IGAP) for providing summary results data for these analyses. The investigators within IGAP contributed to the design and implementation of IGAP and/or provided data but did not participate in analysis or writing of this report. IGAP was made possible by the generous participation of the control subjects, the patients, and their families. The i-Select chips was funded by the French National Foundation on Alzheimer's disease and related disorders. EADI was supported by the LABEX (laboratory of excellence program investment for the future)

DISTALZ grant, Inserm, Institut Pasteur de Lille, Universite de Lille 2 and the Lille University Hospital. GERAD was supported by the Medical Research Council (Grant no. 503480), Alzheimer's Research UK (Grant no. 503176), the Wellcome Trust (Grant no. 082604/2/07/Z) and German Federal Ministry of Education and Research (BMBF): Competence Network Dementia (CND) grant no. 01GI0102, 01GI0711, 01GI0420. CHARGE was partly supported by the NIH/NIA grant R01 AG033193 and the NIA AG081220 and AGES contract N01-AG-12100, the NHLBI grant R01 HL105756, the Icelandic Heart Association, and the Erasmus Medical Center and Erasmus University. ADGC was supported by the NIH/NIA grants: U01 AG032984, U24 AG021886, U01 AG016976, and the Alzheimer's Association grant ADGC-10-196728.

- **Social Science Genetic Association Consortium (SSGAC).** GWAS summary statistics on neuroticism (Okbay et al. 2016) have been contributed by SSGAC investigators and have been downloaded from `https://www.thessgac.org`. For financial support, the SSGAC thanks the U.S. National Science Foundation, the U.S. National Institutes of Health (National Institute on Aging, and the Office for Behavioral and Social Science Research), the Ragnar Soderberg Foundation, the Swedish Research Council, The Jan Wallander and Tom Hedelius Foundation, the European Research Council, and the Pershing Square Fund of the Foundations of Human Behavior.

- **GWAS summary statistics of rheumatoid arthritis.** GWAS summary statistics on rheumatoid arthritis have been contributed by authors of Okada et al. (2014) and have been downloaded from `http://plaza.umin.ac.jp/yokada/datasource/software.htm`.

- **DIAbetes Genetics Replication And Meta-analysis (DIAGRAM) Consortium.** GWAS summary statistics on type 2 diabetes (Morris et al. 2012) have been contributed by DIAGRAM investigators and have been downloaded from `http://www.diagram-consortium.org`.

- **Global Lipids Genetics Consortium (GLGC).** GWAS summary statistics on low-density and high-density lipoprotein cholesterol (Teslovich et al. 2010) have been contributed by GLGC investigators and have been downloaded from `http://csg.sph.umich.edu//abecasis/public/lipids2010`.

- **GWAS summary statistics of atrial fibrillation.** GWAS summary statistics on atrial fibrillation have been contributed by authors of Christophersen et al. (2017) and are available from the corresponding author on reasonable request; see "Data availability" section of Christophersen et al. (2017).

- **Genetic Associations and Mechanisms in Oncology (GAME-ON), Discovery, Biology, and Risk of Inherited Variants in Breast Cancer (DRIVE) Project.** GWAS summary statistics on breast cancer have been contributed by the GAME-ON/DRIVE Project and have been downloaded from `http://gameon.dfci.harvard.edu`. The DRIVE project acknowledges the following GWASs and investigators that shared genome-wide summary data as part of the breast-cancer GWAS meta-analysis: the Australian Breast Cancer Family Study (ABCFS) (John L. Hopper, Melissa C. Southey, Enes Makalic, Daniel F. Schmidt), the British Breast Cancer Study (BBCS) (Olivia Fletcher, Julian Peto, Lorna Gibson, Isabel dos Santos Silva), the Breast and Prostate Cancer Cohort Consortium (BPC3) (David J. Hunter, Sara Lindstrom, Peter Kraft), the Breast Cancer Family Registries (BCFR) (Habib Ahsan, Alice Whittemore), the Dutch Familial Bilateral Breast Cancer Study (DFBBCS) (Quinten Waisfisz, Hanne Meijers-Heijboer, Muriel Adank, Rob B. van der Luijt, Andre G. Uitterlinden, Albert Hofman), German Consortium for Hereditary Breast and Ovarian Cancer (GC-HBOC) (Alfons Meindl, Rita K. Schmutzler, Bertram Muller-Myhsok, Peter Lichtner), the Helsinki Breast Cancer Study (HEBCS) (Heli Nevanlinna, Taru A. Muranen, Kristiina Aittomaki, Carl Blomqvist), the Mammary Carcinoma Risk Factor Investigation (MARIE) (Jenny Chang-Claude, Rebecca Hein, Norbert Dahmen, Lars Beckman), SardiNIA (Laura Crisponi), the Singapore and Sweden Breast Cancer Study (SASBAC) (Per Hall, Kamila Czene, Astrid Irwanto, Jianjun Liu), and the UK2 (Douglas F. Easton, Clare Turnbull, Nazneen Rahman).

### Supplementary Figures

#### Supplementary Figure 1

**Simulation details and additional results of Figure 2.** We use real genotypes of 348,965 genome-wide SNPs on chromosomes 1-22 from 1,458 individuals in the UK Blood Service Control Group (Wellcome Trust Case Control Consortium 2007) to simulate phenotype data, and then compute single-SNP association summary statistics. On these summary data, we compare RSS-NET with existing enrichment methods.

We infer a B cell regulatory network from the paired gene expression and chromatin accessibility data of primary B cells from peripheral blood (Duren et al. 2017; Luo et al. 2020). We use the B cell network to create SNP-level annotation for 348,965 SNPs. Specifically, we let $a_j = 1$ if SNP $j$ is within 100 kb of either a regulatory element or the transcribed region of a member gene in the B cell network, and let $a_j = 0$ otherwise. There are 121,308 SNPs mapped to the B cell network with annotation $a_j = 1$.

To ensure that simulated datasets have roughly the same proportion of true genetic signals, we simulate causal indicators for negative and positive datasets in a paired way. We first simulate the causal indicator $\zeta_j$ of each SNP $j$ in a positive dataset as:

$$\zeta_j \sim \text{Bernoulli}(\pi_j), \;\; \pi_j = 1 / [1 + 10^{-(\theta_0 + a_j\theta)}],$$

where $\theta_0$ and $\theta$ are the baseline and enrichment proportion parameters respectively. We then count the number of causal SNPs in this positive dataset as $n_c := \sum_j \zeta_j$, and randomly choose $n_c$ SNPs as causal SNPs for the corresponding negative dataset.

For positive datasets, we simulate the true effect $\beta_j$ of each SNP $j$ as

$$\begin{aligned} \beta_j \mid \zeta_j = 0 &\sim \delta_0, \\ \beta_j \mid \zeta_j = 1 &\sim \mathcal{N}(0,\, \sigma_0^2 + \sigma^2 \cdot \textstyle\sum_{g \in \mathbf{O}_j} w_{jg}^2), \end{aligned}$$

where $\delta_0$ denotes point mass at zero, $\sigma_0^2$ and $\sigma^2$ are the baseline and enrichment magnitude parameter respectively, $\{\mathbf{O}_j, w_{jg}\}$ are network annotations (see main text Equations 4-5). For negative datasets, we simulate $\beta_j$ from the same model above with $\sigma^2 = 0$.

To ensure that negative and positive datasets have roughly the same magnitude of total genetic signals, we simulate phenotypes by matching signal-to-noise ratios. Specifically, we simulate phenotype $y_i$ of individual $i$ as follows:

$$y_i = \textstyle\sum_{j=1}^{p} x_{ij} \cdot \beta_j + \epsilon_i, \;\; \epsilon_i \sim \mathcal{N}(0, \tau^{-1}),$$

where $p$ is 348,965, $x_{ij}$ is the genotype of SNP $j$ for individual $i$. The true value of residual variance $\tau^{-1}$ is determined by the total proportion of variance in phenotype $\mathbf{y}$ explained by effects of all available SNPs in $\mathbf{X}$: PVE $= V(\mathbf{X}\boldsymbol{\beta}) / [\tau^{-1} + V(\mathbf{X}\boldsymbol{\beta})]$, where $V(\mathbf{X}\boldsymbol{\beta})$ is the sample variance of $\mathbf{X}\boldsymbol{\beta}$.

We set the true values of hyper-parameter as follows: baseline proportion parameter $\theta_0 \in \{-2,-3,-4\}$, enrichment proportion parameter $\theta \in \{0,2\}$, baseline magnitude parameter $\sigma_0 = 1$, enrichment magnitude parameter $\sigma \in \{0,0.5,2\}$, and PVE $\in \{0.2,0.6\}$. For each combination of $\{\theta_0,\theta,\sigma_0,\sigma,\text{PVE}\}$ (excluding negative cases where $\theta = \sigma = 0$), we simulate 200 negative and 200 positive independent datasets. The caption of each panel in this supplementary figure shows the true hyper-parameter values of positive datasets. The table below lists true values of positive datasets for each panel in **Figure 2** ($\sigma_0 = 1$).

| | a | b | c | d |
|---|---|---|---|---|
| sparse: $\theta_0 = -4$ | $\theta = 2, \sigma = 0, \text{PVE} = 0.6$ | $\theta = 0, \sigma = 0.5, \text{PVE} = 0.6$ | $\theta = 2, \sigma = 0.5, \text{PVE} = 0.6$ | $\theta = 2, \sigma = 0, \text{PVE} = 0.2$ |
| polygenic: $\theta_0 = -2$ | $\theta = 2, \sigma = 0, \text{PVE} = 0.6$ | $\theta = 0, \sigma = 0.5, \text{PVE} = 0.6$ | $\theta = 2, \sigma = 0.5, \text{PVE} = 0.6$ | $\theta = 2, \sigma = 0, \text{PVE} = 0.2$ |

We apply RSS-NET to simulated datasets with the following hyper-parameter grids. The baseline parameters $\theta_0$ and $\sigma_0$ are set as their true values. The grids on the enrichment parameters $\theta$ and $\sigma$ are `(0:0.25:1)` if their true values are zero; otherwise they are `[0 ((truth-0.5):0.25:(truth+0.5))]`. For the same simulation scenario, we use the same hyper-parameter grid in RSS-NET analyses of all negative and positive datasets.

For all datasets the target of enrichment testing is the B cell network. A false positive occurs if a method identifies enrichment in a negative dataset. A true positive occurs if a method identifies enrichment in a positive dataset. We evaluate the performance of enrichment methods by plotting the receiver operating characteristic (ROC) curve and computing the area under the ROC curve (AUROC) for each method. Both metrics are implemented in the R package `plotROC` (Sachs 2017).

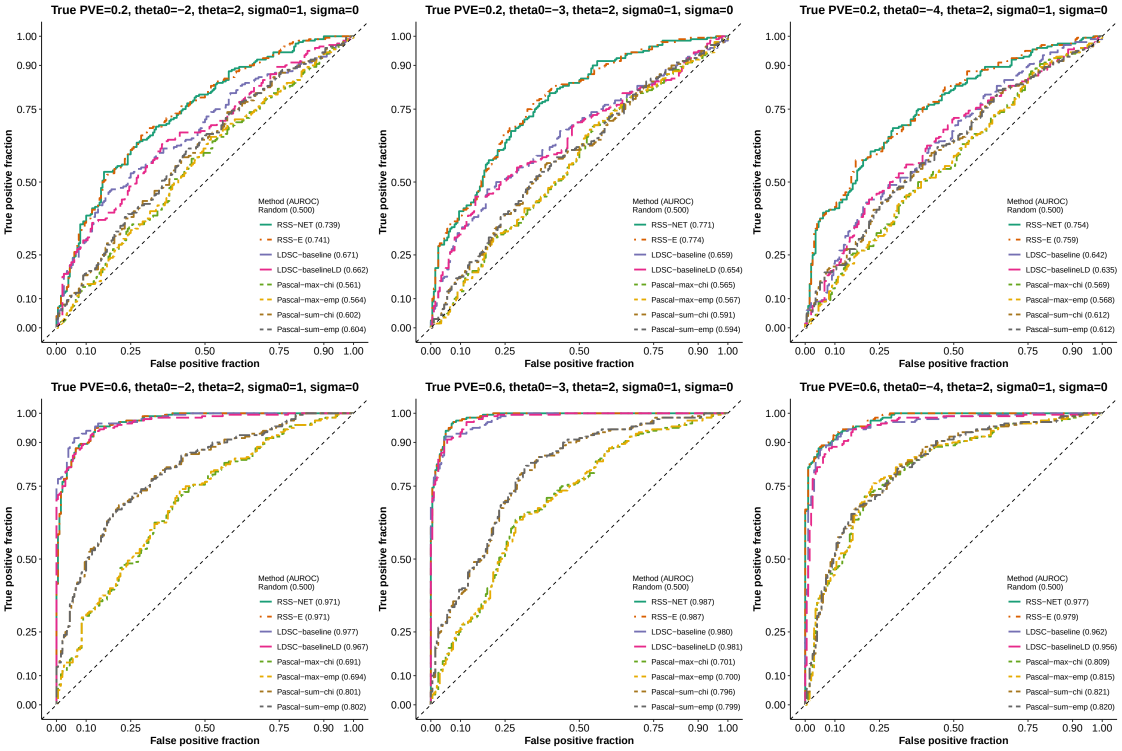

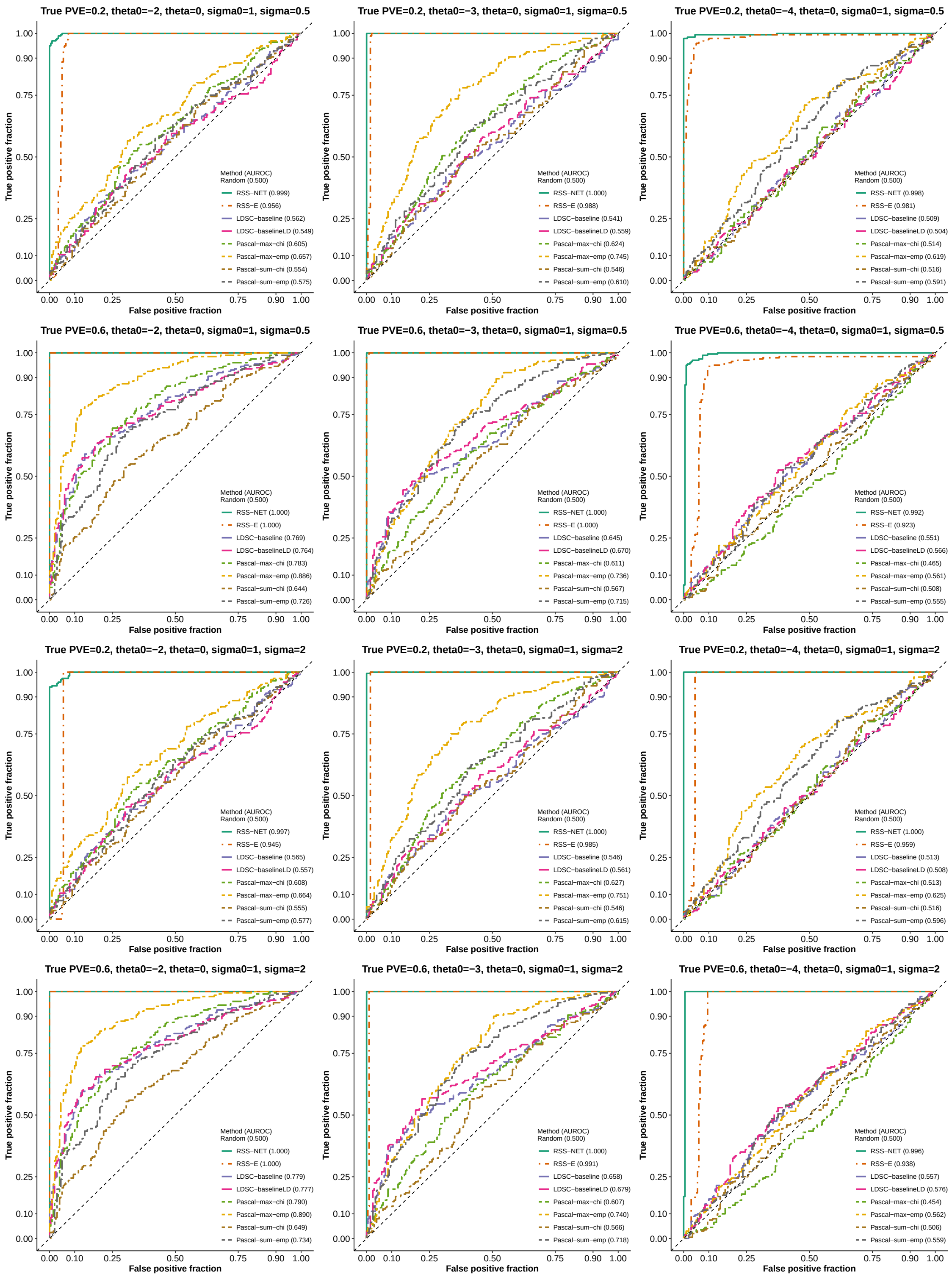
True PVE=0.2, theta0=−2, theta=0, sigma0=1, sigma=0.5
True positive fraction
False positive fraction
Method (AUROC)
Random (0.500)
RSS−NET (0.999)
RSS−E (0.956)
LDSC−baseline (0.562)
LDSC−baselineLD (0.549)
Pascal−max−chi (0.605)
Pascal−max−emp (0.657)
Pascal−sum−chi (0.554)
Pascal−sum−emp (0.575)
True PVE=0.2, theta0=−3, theta=0, sigma0=1, sigma=0.5
Method (AUROC)
Random (0.500)
RSS−NET (1.000)
RSS−E (0.988)
LDSC−baseline (0.541)
LDSC−baselineLD (0.559)
Pascal−max−chi (0.624)
Pascal−max−emp (0.745)
Pascal−sum−chi (0.546)
Pascal−sum−emp (0.610)
True PVE=0.2, theta0=−4, theta=0, sigma0=1, sigma=0.5
Method (AUROC)
Random (0.500)
RSS−NET (0.998)
RSS−E (0.981)
LDSC−baseline (0.509)
LDSC−baselineLD (0.504)
Pascal−max−chi (0.514)
Pascal−max−emp (0.619)
Pascal−sum−chi (0.516)
Pascal−sum−emp (0.591)
True PVE=0.6, theta0=−2, theta=0, sigma0=1, sigma=0.5
Method (AUROC)
Random (0.500)
RSS−NET (1.000)
RSS−E (1.000)
LDSC−baseline (0.769)
LDSC−baselineLD (0.764)
Pascal−max−chi (0.783)
Pascal−max−emp (0.886)
Pascal−sum−chi (0.644)
Pascal−sum−emp (0.726)
True PVE=0.6, theta0=−3, theta=0, sigma0=1, sigma=0.5
Method (AUROC)
Random (0.500)
RSS−NET (1.000)
RSS−E (1.000)
LDSC−baseline (0.645)
LDSC−baselineLD (0.670)
Pascal−max−chi (0.611)
Pascal−max−emp (0.736)
Pascal−sum−chi (0.567)
Pascal−sum−emp (0.715)
True PVE=0.6, theta0=−4, theta=0, sigma0=1, sigma=0.5
Method (AUROC)
Random (0.500)
RSS−NET (0.992)
RSS−E (0.923)
LDSC−baseline (0.551)
LDSC−baselineLD (0.566)
Pascal−max−chi (0.465)
Pascal−max−emp (0.561)
Pascal−sum−chi (0.508)
Pascal−sum−emp (0.555)
True PVE=0.2, theta0=−2, theta=0, sigma0=1, sigma=2
Method (AUROC)
Random (0.500)
RSS−NET (0.997)
RSS−E (0.945)
LDSC−baseline (0.565)
LDSC−baselineLD (0.557)
Pascal−max−chi (0.608)
Pascal−max−emp (0.664)
Pascal−sum−chi (0.555)
Pascal−sum−emp (0.577)
True PVE=0.2, theta0=−3, theta=0, sigma0=1, sigma=2
Method (AUROC)
Random (0.500)
RSS−NET (1.000)
RSS−E (0.985)
LDSC−baseline (0.546)
LDSC−baselineLD (0.561)
Pascal−max−chi (0.627)
Pascal−max−emp (0.751)
Pascal−sum−chi (0.546)
Pascal−sum−emp (0.615)
True PVE=0.2, theta0=−4, theta=0, sigma0=1, sigma=2
Method (AUROC)
Random (0.500)
RSS−NET (1.000)
RSS−E (0.959)
LDSC−baseline (0.513)
LDSC−baselineLD (0.508)
Pascal−max−chi (0.513)
Pascal−max−emp (0.625)
Pascal−sum−chi (0.516)
Pascal−sum−emp (0.596)
True PVE=0.6, theta0=−2, theta=0, sigma0=1, sigma=2
Method (AUROC)
Random (0.500)
RSS−NET (1.000)
RSS−E (1.000)
LDSC−baseline (0.779)
LDSC−baselineLD (0.777)
Pascal−max−chi (0.790)
Pascal−max−emp (0.890)
Pascal−sum−chi (0.649)
Pascal−sum−emp (0.734)
True PVE=0.6, theta0=−3, theta=0, sigma0=1, sigma=2
Method (AUROC)
Random (0.500)
RSS−NET (1.000)
RSS−E (0.991)
LDSC−baseline (0.658)
LDSC−baselineLD (0.679)
Pascal−max−chi (0.607)
Pascal−max−emp (0.740)
Pascal−sum−chi (0.566)
Pascal−sum−emp (0.718)
True PVE=0.6, theta0=−4, theta=0, sigma0=1, sigma=2
Method (AUROC)
Random (0.500)
RSS−NET (0.996)
RSS−E (0.938)
LDSC−baseline (0.557)
LDSC−baselineLD (0.576)
Pascal−max−chi (0.454)
Pascal−max−emp (0.562)
Pascal−sum−chi (0.506)
Pascal−sum−emp (0.559)

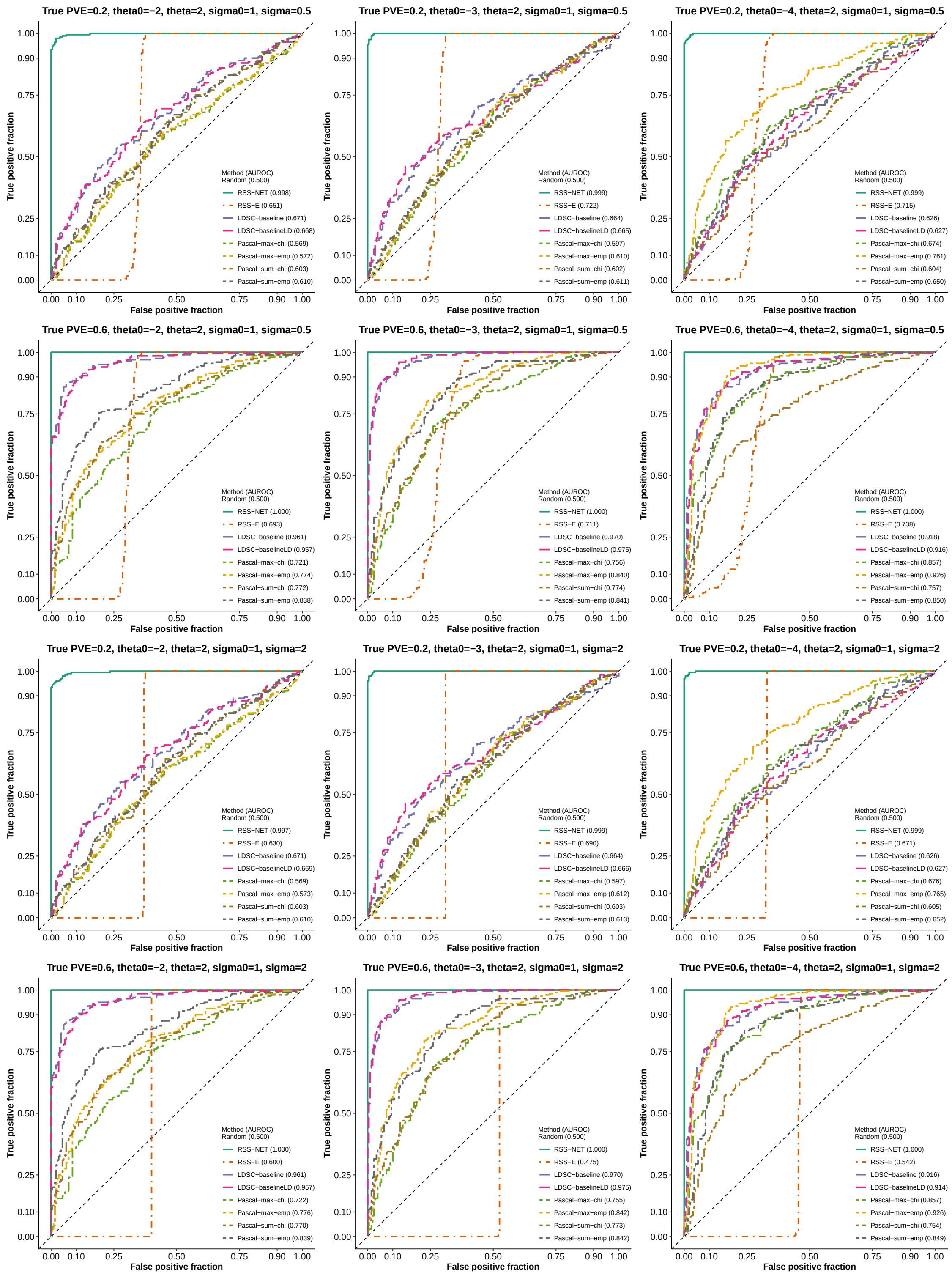
True PVE=0.2, theta0=−2, theta=2, sigma0=1, sigma=0.5
Method (AUROC)
Random (0.500)
RSS−NET (0.998)
RSS−E (0.651)
LDSC−baseline (0.671)
LDSC−baselineLD (0.668)
Pascal−max−chi (0.569)
Pascal−max−emp (0.572)
Pascal−sum−chi (0.603)
Pascal−sum−emp (0.610)
True PVE=0.2, theta0=−3, theta=2, sigma0=1, sigma=0.5
Method (AUROC)
Random (0.500)
RSS−NET (0.999)
RSS−E (0.722)
LDSC−baseline (0.664)
LDSC−baselineLD (0.665)
Pascal−max−chi (0.597)
Pascal−max−emp (0.610)
Pascal−sum−chi (0.602)
Pascal−sum−emp (0.611)
True PVE=0.2, theta0=−4, theta=2, sigma0=1, sigma=0.5
Method (AUROC)
Random (0.500)
RSS−NET (0.999)
RSS−E (0.715)
LDSC−baseline (0.626)
LDSC−baselineLD (0.627)
Pascal−max−chi (0.674)
Pascal−max−emp (0.761)
Pascal−sum−chi (0.604)
Pascal−sum−emp (0.650)
True PVE=0.6, theta0=−2, theta=2, sigma0=1, sigma=0.5
Method (AUROC)
Random (0.500)
RSS−NET (1.000)
RSS−E (0.693)
LDSC−baseline (0.961)
LDSC−baselineLD (0.957)
Pascal−max−chi (0.721)
Pascal−max−emp (0.774)
Pascal−sum−chi (0.772)
Pascal−sum−emp (0.838)
True PVE=0.6, theta0=−3, theta=2, sigma0=1, sigma=0.5
Method (AUROC)
Random (0.500)
RSS−NET (1.000)
RSS−E (0.711)
LDSC−baseline (0.970)
LDSC−baselineLD (0.975)
Pascal−max−chi (0.756)
Pascal−max−emp (0.840)
Pascal−sum−chi (0.774)
Pascal−sum−emp (0.841)
True PVE=0.6, theta0=−4, theta=2, sigma0=1, sigma=0.5
Method (AUROC)
Random (0.500)
RSS−NET (1.000)
RSS−E (0.738)
LDSC−baseline (0.918)
LDSC−baselineLD (0.916)
Pascal−max−chi (0.857)
Pascal−max−emp (0.926)
Pascal−sum−chi (0.757)
Pascal−sum−emp (0.850)
True PVE=0.2, theta0=−2, theta=2, sigma0=1, sigma=2
Method (AUROC)
Random (0.500)
RSS−NET (0.997)
RSS−E (0.630)
LDSC−baseline (0.671)
LDSC−baselineLD (0.669)
Pascal−max−chi (0.569)
Pascal−max−emp (0.573)
Pascal−sum−chi (0.603)
Pascal−sum−emp (0.610)
True PVE=0.2, theta0=−3, theta=2, sigma0=1, sigma=2
Method (AUROC)
Random (0.500)
RSS−NET (0.999)
RSS−E (0.690)
LDSC−baseline (0.664)
LDSC−baselineLD (0.666)
Pascal−max−chi (0.597)
Pascal−max−emp (0.612)
Pascal−sum−chi (0.603)
Pascal−sum−emp (0.613)
True PVE=0.2, theta0=−4, theta=2, sigma0=1, sigma=2
Method (AUROC)
Random (0.500)
RSS−NET (0.999)
RSS−E (0.671)
LDSC−baseline (0.626)
LDSC−baselineLD (0.627)
Pascal−max−chi (0.676)
Pascal−max−emp (0.765)
Pascal−sum−chi (0.605)
Pascal−sum−emp (0.652)
True PVE=0.6, theta0=−2, theta=2, sigma0=1, sigma=2
Method (AUROC)
Random (0.500)
RSS−NET (1.000)
RSS−E (0.600)
LDSC−baseline (0.961)
LDSC−baselineLD (0.957)
Pascal−max−chi (0.722)
Pascal−max−emp (0.776)
Pascal−sum−chi (0.770)
Pascal−sum−emp (0.839)
True PVE=0.6, theta0=−3, theta=2, sigma0=1, sigma=2
Method (AUROC)
Random (0.500)
RSS−NET (1.000)
RSS−E (0.475)
LDSC−baseline (0.970)
LDSC−baselineLD (0.975)
Pascal−max−chi (0.755)
Pascal−max−emp (0.842)
Pascal−sum−chi (0.773)
Pascal−sum−emp (0.842)
True PVE=0.6, theta0=−4, theta=2, sigma0=1, sigma=2
Method (AUROC)
Random (0.500)
RSS−NET (1.000)
RSS−E (0.542)
LDSC−baseline (0.916)
LDSC−baselineLD (0.914)
Pascal−max−chi (0.857)
Pascal−max−emp (0.926)
Pascal−sum−chi (0.754)
Pascal−sum−emp (0.849)
True positive fraction
False positive fraction
0.00
0.10
0.25
0.50
0.75
0.90
1.00

#### Supplementary Figure 2

We repeat the simulations in **Supplementary Figure 1** on a different network based on data from vagina (https://github.com/suwonglab/rss-net/tree/master/data/).

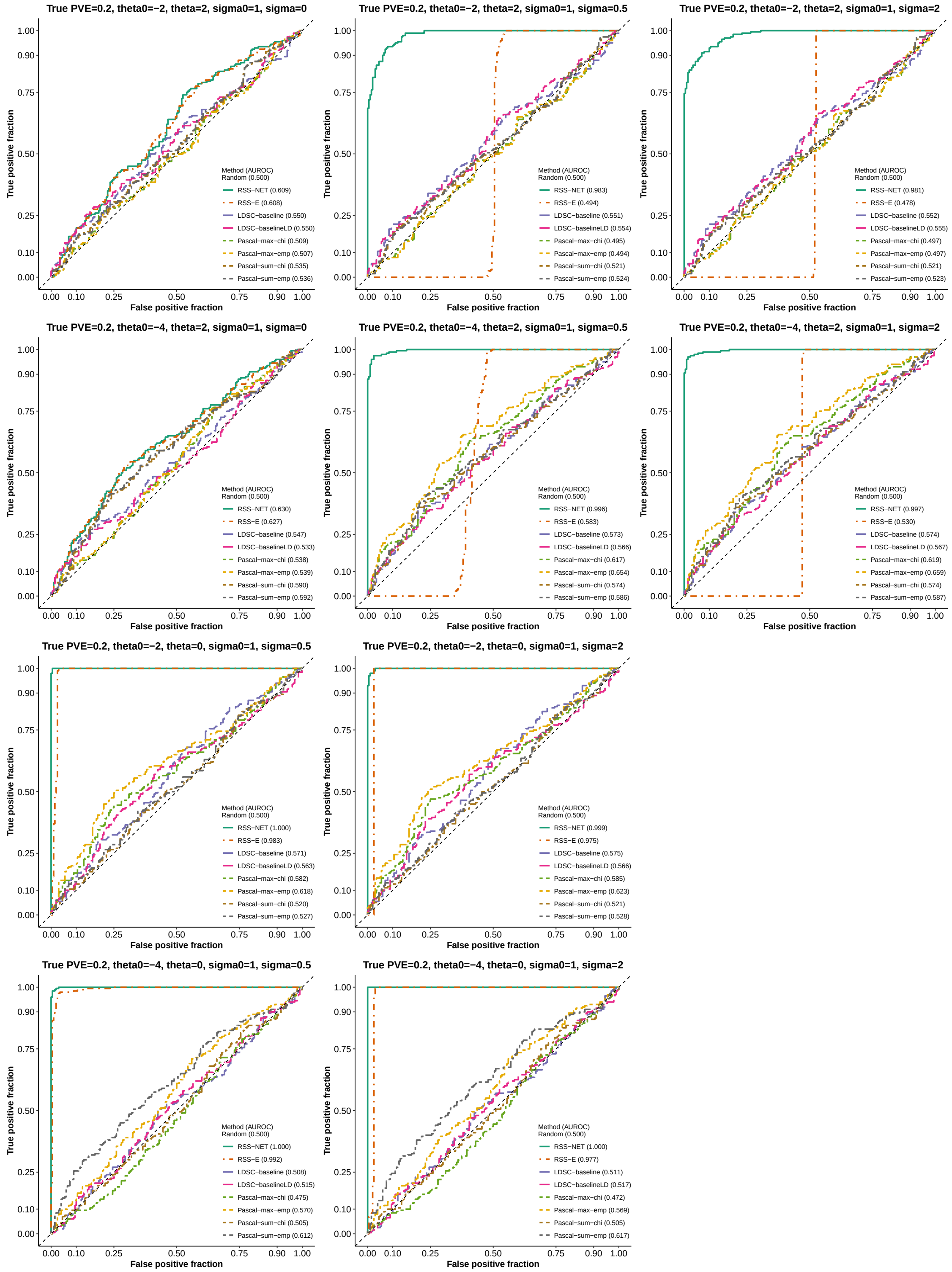

#### Supplementary Figure 3

**Simulation details and additional results of Figure 3(a).** This part aims to assess the robustness of RSS-NET to model mis-specification where a random set of "near-gene" SNPs is enriched for association in negative datasets. Details of this part are almost identical to those in **Supplementary Figure 1**. Here we only highlight the differences.

We define a SNP as "near-gene" if this SNP is within 100 kb of the transcribed region of any autosomal protein-coding gene. In total, there are 254,296 near-gene SNPs among 348,965 genome-wide SNPs from Wellcome Trust Case Control Consortium (2007).

Here we simulate negative and positive datasets in a paired way. We first simulate a positive dataset as in **Supplementary Figure 1**. For this positive dataset, we count the total number of causal SNPs as $n_c = \sum_j \zeta_j$, and count the number of causal SNPs in the target network as $n_p = \sum_j \zeta_j a_j$ (recall that $a_j = 1$ if SNP $j$ is in the network). We then randomly choose $n_p$ SNPs from the 254,296 near-gene SNPs and $(n_c - n_p)$ SNPs from the remaining SNPs, and use them as causal SNPs for the corresponding negative dataset. With causal indicators $\{\zeta_j\}$ in place, we simulate the true genetic effects $\boldsymbol{\beta}$ in a negative dataset as follows

$$\begin{aligned} \beta_j \mid \zeta_j = 0 &\sim \delta_0, \\ \beta_j \mid \zeta_j = 1,\ \alpha_j = 0 &\sim \mathcal{N}(0,\ \sigma_0^2), \\ \beta_j \mid \zeta_j = 1,\ \alpha_j = 1 &\sim \mathcal{N}(0,\ (2\cdot\sigma_0)^2), \end{aligned}$$

where $\zeta_j = 1$ if SNP $j$ is causal and 0 otherwise, $\alpha_j = 1$ if SNP $j$ is near-gene and 0 otherwise. The rest of simulations is the same as **Supplementary Figure 1**.

The caption of each panel in this supplementary figure shows the true hyper-parameter values of positive datasets. The sparse scenario in **Figure 3(a)** corresponds to simulations with true hyper-parameter values of positive datasets being $\theta_0 = -4$, $\theta = 2$, $\sigma_0 = 1$, $\sigma = 0.5$ and PVE = 0.6. The polygenic scenario in **Figure 3(a)** corresponds to simulations with true hyper-parameter values of positive datasets being $\theta_0 = -2$, $\theta = 2$, $\sigma_0 = 1$, $\sigma = 0.5$ and PVE = 0.6.

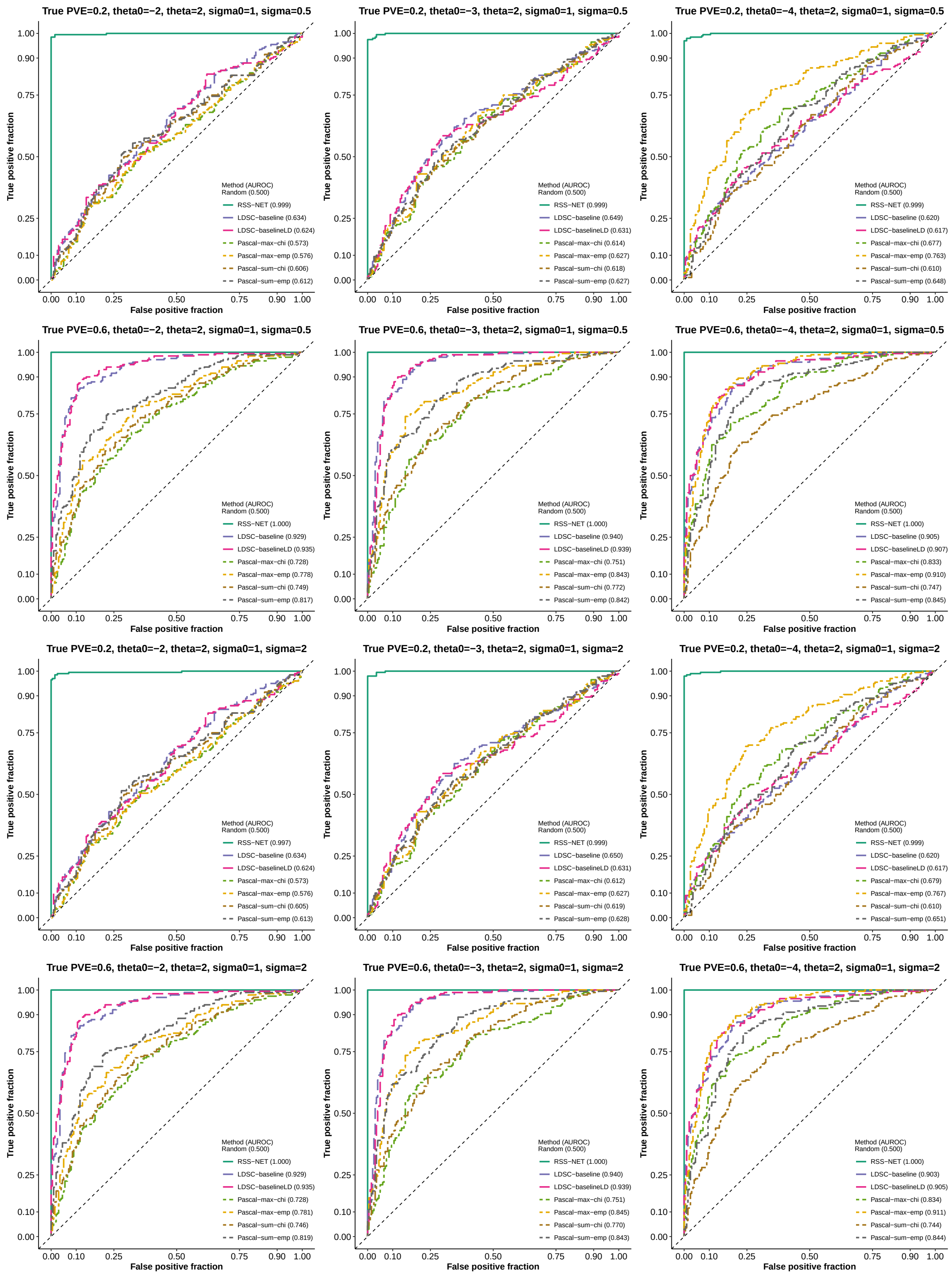
True PVE=0.2, theta0=−2, theta=2, sigma0=1, sigma=0.5
Method (AUROC)
Random (0.500)
RSS−NET (0.999)
LDSC−baseline (0.634)
LDSC−baselineLD (0.624)
Pascal−max−chi (0.573)
Pascal−max−emp (0.576)
Pascal−sum−chi (0.606)
Pascal−sum−emp (0.612)
True PVE=0.2, theta0=−3, theta=2, sigma0=1, sigma=0.5
Method (AUROC)
Random (0.500)
RSS−NET (0.999)
LDSC−baseline (0.649)
LDSC−baselineLD (0.631)
Pascal−max−chi (0.614)
Pascal−max−emp (0.627)
Pascal−sum−chi (0.618)
Pascal−sum−emp (0.627)
True PVE=0.2, theta0=−4, theta=2, sigma0=1, sigma=0.5
Method (AUROC)
Random (0.500)
RSS−NET (0.999)
LDSC−baseline (0.620)
LDSC−baselineLD (0.617)
Pascal−max−chi (0.677)
Pascal−max−emp (0.763)
Pascal−sum−chi (0.610)
Pascal−sum−emp (0.648)
True PVE=0.6, theta0=−2, theta=2, sigma0=1, sigma=0.5
Method (AUROC)
Random (0.500)
RSS−NET (1.000)
LDSC−baseline (0.929)
LDSC−baselineLD (0.935)
Pascal−max−chi (0.728)
Pascal−max−emp (0.778)
Pascal−sum−chi (0.749)
Pascal−sum−emp (0.817)
True PVE=0.6, theta0=−3, theta=2, sigma0=1, sigma=0.5
Method (AUROC)
Random (0.500)
RSS−NET (1.000)
LDSC−baseline (0.940)
LDSC−baselineLD (0.939)
Pascal−max−chi (0.751)
Pascal−max−emp (0.843)
Pascal−sum−chi (0.772)
Pascal−sum−emp (0.842)
True PVE=0.6, theta0=−4, theta=2, sigma0=1, sigma=0.5
Method (AUROC)
Random (0.500)
RSS−NET (1.000)
LDSC−baseline (0.905)
LDSC−baselineLD (0.907)
Pascal−max−chi (0.833)
Pascal−max−emp (0.910)
Pascal−sum−chi (0.747)
Pascal−sum−emp (0.845)
True PVE=0.2, theta0=−2, theta=2, sigma0=1, sigma=2
Method (AUROC)
Random (0.500)
RSS−NET (0.997)
LDSC−baseline (0.634)
LDSC−baselineLD (0.624)
Pascal−max−chi (0.573)
Pascal−max−emp (0.576)
Pascal−sum−chi (0.605)
Pascal−sum−emp (0.613)
True PVE=0.2, theta0=−3, theta=2, sigma0=1, sigma=2
Method (AUROC)
Random (0.500)
RSS−NET (0.999)
LDSC−baseline (0.650)
LDSC−baselineLD (0.631)
Pascal−max−chi (0.612)
Pascal−max−emp (0.627)
Pascal−sum−chi (0.619)
Pascal−sum−emp (0.628)
True PVE=0.2, theta0=−4, theta=2, sigma0=1, sigma=2
Method (AUROC)
Random (0.500)
RSS−NET (0.999)
LDSC−baseline (0.620)
LDSC−baselineLD (0.617)
Pascal−max−chi (0.679)
Pascal−max−emp (0.767)
Pascal−sum−chi (0.610)
Pascal−sum−emp (0.651)
True PVE=0.6, theta0=−2, theta=2, sigma0=1, sigma=2
Method (AUROC)
Random (0.500)
RSS−NET (1.000)
LDSC−baseline (0.929)
LDSC−baselineLD (0.935)
Pascal−max−chi (0.728)
Pascal−max−emp (0.781)
Pascal−sum−chi (0.746)
Pascal−sum−emp (0.819)
True PVE=0.6, theta0=−3, theta=2, sigma0=1, sigma=2
Method (AUROC)
Random (0.500)
RSS−NET (1.000)
LDSC−baseline (0.940)
LDSC−baselineLD (0.939)
Pascal−max−chi (0.751)
Pascal−max−emp (0.845)
Pascal−sum−chi (0.770)
Pascal−sum−emp (0.843)
True PVE=0.6, theta0=−4, theta=2, sigma0=1, sigma=2
Method (AUROC)
Random (0.500)
RSS−NET (1.000)
LDSC−baseline (0.903)
LDSC−baselineLD (0.905)
Pascal−max−chi (0.834)
Pascal−max−emp (0.911)
Pascal−sum−chi (0.744)
Pascal−sum−emp (0.844)
True positive fraction
False positive fraction

#### Supplementary Figure 4

**Simulation details and additional results of Figure 3(b).** This part aims to assess the robustness of RSS-NET to model mis-specification where a random set of "near-RE" SNPs is enriched for association in negative datasets. Details of this part are almost identical to those in **Supplementary Figure 1**. Here we only highlight the differences.

We define a SNP as "near-RE" if this SNP is within 10 kb of any RE. There are 3,895,021 unique autosomal REs associated with 38 regulatory networks analyzed in this study. In total, there are 169,100 near-RE SNPs among 348,965 genome-wide SNPs from Wellcome Trust Case Control Consortium (2007).

Here we simulate negative and positive datasets in a paired way. We first simulate a positive dataset as in **Supplementary Figure 1**. For this positive dataset, we count the total number of causal SNPs as $n_c = \sum_j \zeta_j$, and count the number of causal SNPs in the target network as $n_p = \sum_j \zeta_j a_j$ (recall that $a_j = 1$ if SNP $j$ is in the network). We then randomly choose $n_p$ SNPs from the 169,100 near-RE SNPs and $(n_c - n_p)$ SNPs from the remaining SNPs, and use them as causal SNPs for the corresponding negative dataset. With causal indicators $\{\zeta_j\}$ in place, we simulate the true genetic effects $\boldsymbol{\beta}$ in a negative dataset as follows

$$\begin{aligned}
\beta_j \mid \zeta_j = 0 &\sim \delta_0, \\
\beta_j \mid \zeta_j = 1,\ \alpha_j = 0 &\sim \mathcal{N}(0,\ \sigma_0^2), \\
\beta_j \mid \zeta_j = 1,\ \alpha_j = 1 &\sim \mathcal{N}(0,\ (2\cdot\sigma_0)^2),
\end{aligned}$$

where $\zeta_j = 1$ if SNP $j$ is causal and 0 otherwise, $\alpha_j = 1$ if SNP $j$ is near-RE and 0 otherwise. The rest of simulations is the same as **Supplementary Figure 1**

The caption of each panel in this supplementary figure shows the true hyper-parameter values of positive datasets. The sparse scenario in **Figure 3(b)** corresponds to simulations with true hyper-parameter values of positive datasets being $\theta_0 = -4$, $\theta = 2$, $\sigma_0 = 1$, $\sigma = 0.5$ and PVE = 0.6. The polygenic scenario in **Figure 3(b)** corresponds to simulations with true hyper-parameter values of positive datasets being $\theta_0 = -2$, $\theta = 2$, $\sigma_0 = 1$, $\sigma = 0.5$ and PVE = 0.6.

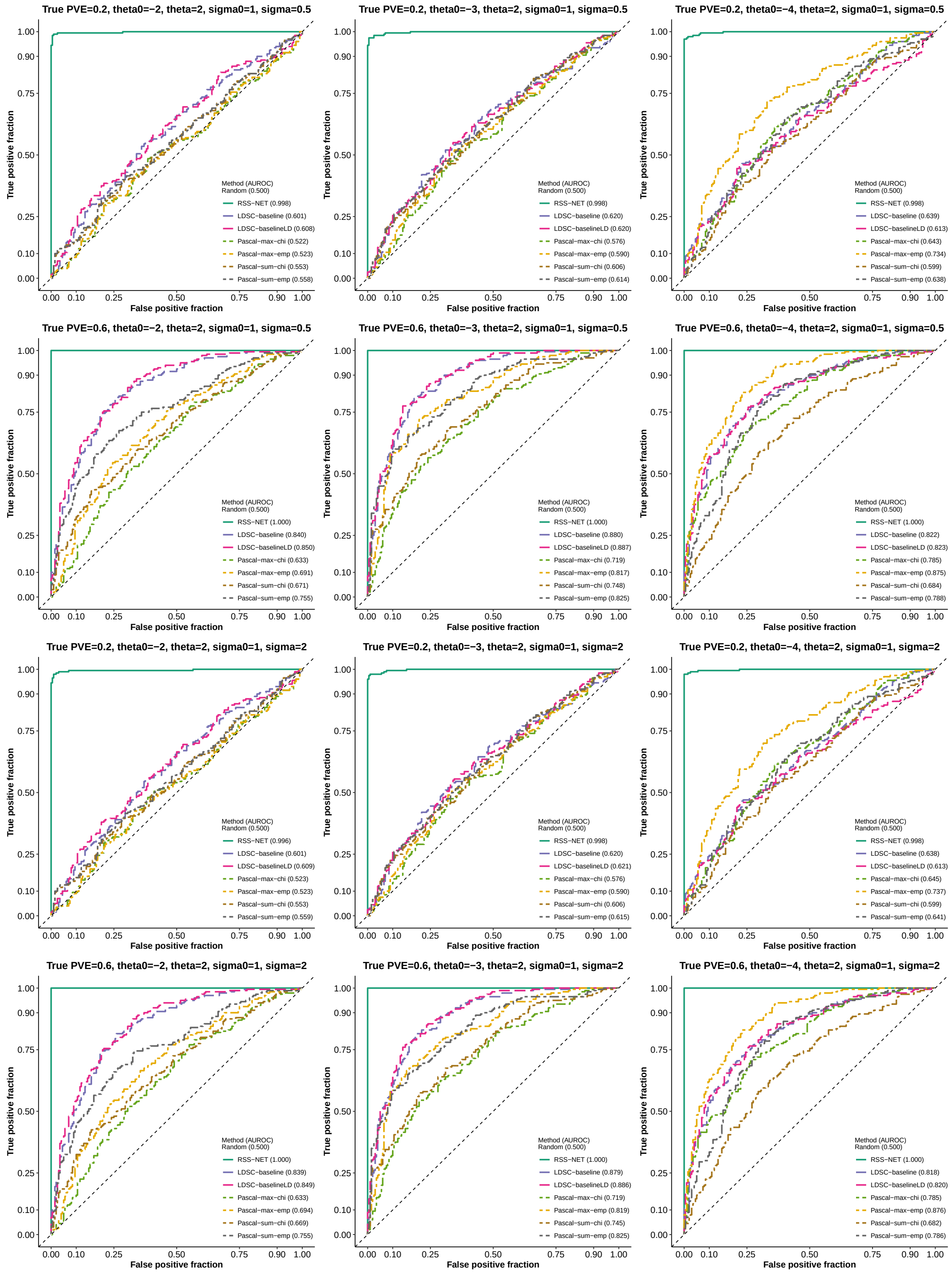
True PVE=0.2, theta0=−2, theta=2, sigma0=1, sigma=0.5
True positive fraction
False positive fraction
0.00 0.10 0.25 0.50 0.75 0.90 1.00
Method (AUROC)
Random (0.500)
RSS−NET (0.998)
LDSC−baseline (0.601)
LDSC−baselineLD (0.608)
Pascal−max−chi (0.522)
Pascal−max−emp (0.523)
Pascal−sum−chi (0.553)
Pascal−sum−emp (0.558)
True PVE=0.2, theta0=−3, theta=2, sigma0=1, sigma=0.5
Method (AUROC)
Random (0.500)
RSS−NET (0.998)
LDSC−baseline (0.620)
LDSC−baselineLD (0.620)
Pascal−max−chi (0.576)
Pascal−max−emp (0.590)
Pascal−sum−chi (0.606)
Pascal−sum−emp (0.614)
True PVE=0.2, theta0=−4, theta=2, sigma0=1, sigma=0.5
Method (AUROC)
Random (0.500)
RSS−NET (0.998)
LDSC−baseline (0.639)
LDSC−baselineLD (0.613)
Pascal−max−chi (0.643)
Pascal−max−emp (0.734)
Pascal−sum−chi (0.599)
Pascal−sum−emp (0.638)
True PVE=0.6, theta0=−2, theta=2, sigma0=1, sigma=0.5
Method (AUROC)
Random (0.500)
RSS−NET (1.000)
LDSC−baseline (0.840)
LDSC−baselineLD (0.850)
Pascal−max−chi (0.633)
Pascal−max−emp (0.691)
Pascal−sum−chi (0.671)
Pascal−sum−emp (0.755)
True PVE=0.6, theta0=−3, theta=2, sigma0=1, sigma=0.5
Method (AUROC)
Random (0.500)
RSS−NET (1.000)
LDSC−baseline (0.880)
LDSC−baselineLD (0.887)
Pascal−max−chi (0.719)
Pascal−max−emp (0.817)
Pascal−sum−chi (0.748)
Pascal−sum−emp (0.825)
True PVE=0.6, theta0=−4, theta=2, sigma0=1, sigma=0.5
Method (AUROC)
Random (0.500)
RSS−NET (1.000)
LDSC−baseline (0.822)
LDSC−baselineLD (0.823)
Pascal−max−chi (0.785)
Pascal−max−emp (0.875)
Pascal−sum−chi (0.684)
Pascal−sum−emp (0.788)
True PVE=0.2, theta0=−2, theta=2, sigma0=1, sigma=2
Method (AUROC)
Random (0.500)
RSS−NET (0.996)
LDSC−baseline (0.601)
LDSC−baselineLD (0.609)
Pascal−max−chi (0.523)
Pascal−max−emp (0.523)
Pascal−sum−chi (0.553)
Pascal−sum−emp (0.559)
True PVE=0.2, theta0=−3, theta=2, sigma0=1, sigma=2
Method (AUROC)
Random (0.500)
RSS−NET (0.998)
LDSC−baseline (0.620)
LDSC−baselineLD (0.621)
Pascal−max−chi (0.576)
Pascal−max−emp (0.590)
Pascal−sum−chi (0.606)
Pascal−sum−emp (0.615)
True PVE=0.2, theta0=−4, theta=2, sigma0=1, sigma=2
Method (AUROC)
Random (0.500)
RSS−NET (0.998)
LDSC−baseline (0.638)
LDSC−baselineLD (0.613)
Pascal−max−chi (0.645)
Pascal−max−emp (0.737)
Pascal−sum−chi (0.599)
Pascal−sum−emp (0.641)
True PVE=0.6, theta0=−2, theta=2, sigma0=1, sigma=2
Method (AUROC)
Random (0.500)
RSS−NET (1.000)
LDSC−baseline (0.839)
LDSC−baselineLD (0.849)
Pascal−max−chi (0.633)
Pascal−max−emp (0.694)
Pascal−sum−chi (0.669)
Pascal−sum−emp (0.755)
True PVE=0.6, theta0=−3, theta=2, sigma0=1, sigma=2
Method (AUROC)
Random (0.500)
RSS−NET (1.000)
LDSC−baseline (0.879)
LDSC−baselineLD (0.886)
Pascal−max−chi (0.719)
Pascal−max−emp (0.819)
Pascal−sum−chi (0.745)
Pascal−sum−emp (0.825)
True PVE=0.6, theta0=−4, theta=2, sigma0=1, sigma=2
Method (AUROC)
Random (0.500)
RSS−NET (1.000)
LDSC−baseline (0.818)
LDSC−baselineLD (0.820)
Pascal−max−chi (0.785)
Pascal−max−emp (0.876)
Pascal−sum−chi (0.682)
Pascal−sum−emp (0.786)

#### Supplementary Figure 5

**Simulation details and additional results of Figure 3(c).** This part aims to assess the robustness of RSS-NET to model mis-specification where true genetic effects are MAF- and LD-dependent in negative datasets. Details of this part are almost identical to those in **Supplementary Figure 1**. Here we only highlight the differences.

Here we simulate negative and positive datasets in a paired way. We first simulate a positive dataset as in **Supplementary Figure 1**. For this positive dataset, we count the total number of causal SNPs as $n_c = \sum_j \zeta_j$. We then randomly choose $n_c$ genome-wide SNPs as causal SNPs for the corresponding negative dataset. With causal indicators $\{\zeta_j\}$ in place, we simulate the true genetic effects $\boldsymbol{\beta}$ in a negative dataset as follows

$$\begin{aligned} \beta_j \mid \zeta_j = 0 &\sim \delta_0, \\ \beta_j \mid \zeta_j = 1 &\sim \mathcal{N}(0,\, \sigma_0^2 + \textstyle\sum_{k=1}^{16} a_{jk} \cdot \tau_k), \\ \tau_k &\sim \mathcal{N}(\hat{\mu}_k,\, \hat{\sigma}_k^2), \end{aligned}$$

where $\zeta_j = 1$ if SNP $j$ is causal and 0 otherwise, $a_{jk}$ is the value of annotation $k$ at SNP $j$, the 16 annotations include 10 MAF bins and 6 LD-related annotations defined in Gazal et al. (2017), $\{\hat{\mu}_k, \hat{\sigma}_k\}$ are meta-analyzed mean and standard error estimates for $\tau_k$ across 31 independent human traits, which are provided in the Supplementary Table 9 of Gazal et al. (2017). The rest of simulations is the same as **Supplementary Figure 1**

The caption of each panel in this supplementary figure shows the true hyper-parameter values of positive datasets. The sparse scenario in **Figure 3(c)** corresponds to simulations with true hyper-parameter values of positive datasets being $\theta_0 = -4$, $\theta = 2$, $\sigma_0 = 1$, $\sigma = 0.5$ and PVE $= 0.6$. The polygenic scenario in **Figure 3(c)** corresponds to simulations with true hyper-parameter values of positive datasets being $\theta_0 = -2$, $\theta = 2$, $\sigma_0 = 1$, $\sigma = 0.5$ and PVE $= 0.6$.

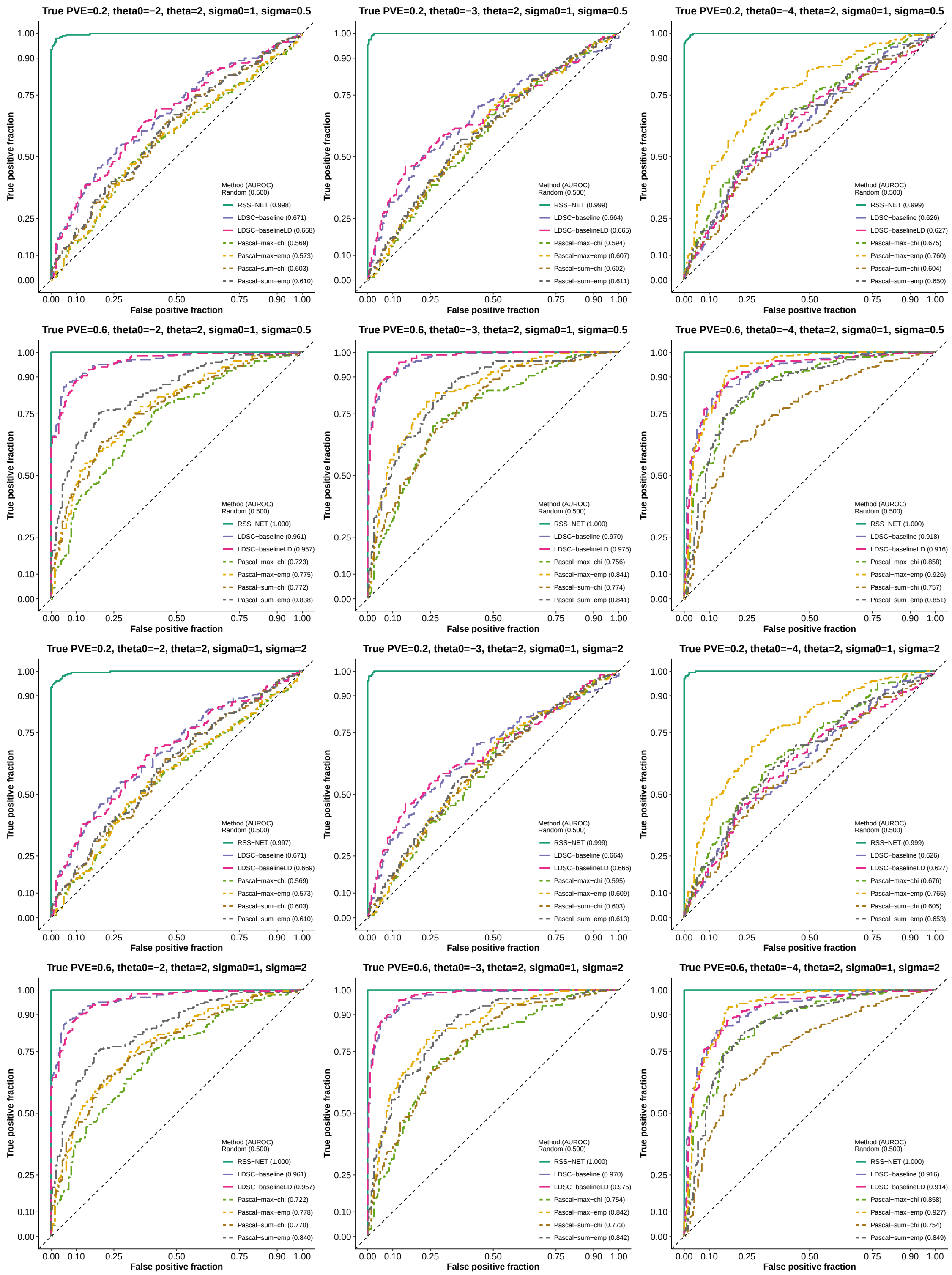
True PVE=0.2, theta0=−2, theta=2, sigma0=1, sigma=0.5
True positive fraction
False positive fraction
0.00
0.10
0.25
0.50
0.75
0.90
1.00
Method (AUROC)
Random (0.500)
RSS−NET (0.998)
LDSC−baseline (0.671)
LDSC−baselineLD (0.668)
Pascal−max−chi (0.569)
Pascal−max−emp (0.573)
Pascal−sum−chi (0.603)
Pascal−sum−emp (0.610)
True PVE=0.2, theta0=−3, theta=2, sigma0=1, sigma=0.5
Method (AUROC)
Random (0.500)
RSS−NET (0.999)
LDSC−baseline (0.664)
LDSC−baselineLD (0.665)
Pascal−max−chi (0.594)
Pascal−max−emp (0.607)
Pascal−sum−chi (0.602)
Pascal−sum−emp (0.611)
True PVE=0.2, theta0=−4, theta=2, sigma0=1, sigma=0.5
Method (AUROC)
Random (0.500)
RSS−NET (0.999)
LDSC−baseline (0.626)
LDSC−baselineLD (0.627)
Pascal−max−chi (0.675)
Pascal−max−emp (0.760)
Pascal−sum−chi (0.604)
Pascal−sum−emp (0.650)
True PVE=0.6, theta0=−2, theta=2, sigma0=1, sigma=0.5
Method (AUROC)
Random (0.500)
RSS−NET (1.000)
LDSC−baseline (0.961)
LDSC−baselineLD (0.957)
Pascal−max−chi (0.723)
Pascal−max−emp (0.775)
Pascal−sum−chi (0.772)
Pascal−sum−emp (0.838)
True PVE=0.6, theta0=−3, theta=2, sigma0=1, sigma=0.5
Method (AUROC)
Random (0.500)
RSS−NET (1.000)
LDSC−baseline (0.970)
LDSC−baselineLD (0.975)
Pascal−max−chi (0.756)
Pascal−max−emp (0.841)
Pascal−sum−chi (0.774)
Pascal−sum−emp (0.841)
True PVE=0.6, theta0=−4, theta=2, sigma0=1, sigma=0.5
Method (AUROC)
Random (0.500)
RSS−NET (1.000)
LDSC−baseline (0.918)
LDSC−baselineLD (0.916)
Pascal−max−chi (0.858)
Pascal−max−emp (0.926)
Pascal−sum−chi (0.757)
Pascal−sum−emp (0.851)
True PVE=0.2, theta0=−2, theta=2, sigma0=1, sigma=2
Method (AUROC)
Random (0.500)
RSS−NET (0.997)
LDSC−baseline (0.671)
LDSC−baselineLD (0.669)
Pascal−max−chi (0.569)
Pascal−max−emp (0.573)
Pascal−sum−chi (0.603)
Pascal−sum−emp (0.610)
True PVE=0.2, theta0=−3, theta=2, sigma0=1, sigma=2
Method (AUROC)
Random (0.500)
RSS−NET (0.999)
LDSC−baseline (0.664)
LDSC−baselineLD (0.666)
Pascal−max−chi (0.595)
Pascal−max−emp (0.609)
Pascal−sum−chi (0.603)
Pascal−sum−emp (0.613)
True PVE=0.2, theta0=−4, theta=2, sigma0=1, sigma=2
Method (AUROC)
Random (0.500)
RSS−NET (0.999)
LDSC−baseline (0.626)
LDSC−baselineLD (0.627)
Pascal−max−chi (0.676)
Pascal−max−emp (0.765)
Pascal−sum−chi (0.605)
Pascal−sum−emp (0.653)
True PVE=0.6, theta0=−2, theta=2, sigma0=1, sigma=2
Method (AUROC)
Random (0.500)
RSS−NET (1.000)
LDSC−baseline (0.961)
LDSC−baselineLD (0.957)
Pascal−max−chi (0.722)
Pascal−max−emp (0.778)
Pascal−sum−chi (0.770)
Pascal−sum−emp (0.840)
True PVE=0.6, theta0=−3, theta=2, sigma0=1, sigma=2
Method (AUROC)
Random (0.500)
RSS−NET (1.000)
LDSC−baseline (0.970)
LDSC−baselineLD (0.975)
Pascal−max−chi (0.754)
Pascal−max−emp (0.842)
Pascal−sum−chi (0.773)
Pascal−sum−emp (0.842)
True PVE=0.6, theta0=−4, theta=2, sigma0=1, sigma=2
Method (AUROC)
Random (0.500)
RSS−NET (1.000)
LDSC−baseline (0.916)
LDSC−baselineLD (0.914)
Pascal−max−chi (0.858)
Pascal−max−emp (0.927)
Pascal−sum−chi (0.754)
Pascal−sum−emp (0.849)

#### Supplementary Figure 6

**Simulation details and additional results of Figure 3(d).** This part aims to assess the robustness of RSS-NET to model mis-specification where an edge-altered version of the true network is enriched for genetic association in negative datasets. Details of this part are almost identical to those in **Supplementary Figure 1**. Here we only highlight the differences.

We simulate random edge-altered networks on the basis of the B cell regulatory network defined in **Supplementary Figure 1**. Specifically, we keep all associated REs and member genes of the B cell network, remove the actual connections between TFs and TGs, and then create fake edges by randomly connecting TFs to TGs with random edge weights. We ensure that the edge-altered network has the same number of edges as the actual B cell network, and their edge weights have the same distribution. For each negative dataset, we simulate a new edge-altered network.

We first simulate positive datasets as in **Supplementary Figure 1**. Since the true and edge-altered networks have the same associated REs and member genes, we let a negative dataset and its corresponding positive dataset have the same causal indicators $\{\zeta_j\}$. We then simulate true genetic effects $\boldsymbol{\beta}$ in a negative dataset as:

$$
\begin{aligned}
\beta_j \mid \zeta_j = 0 &\sim \delta_0, \\
\beta_j \mid \zeta_j = 1 &\sim \mathcal{N}(0,\ \sigma_0^2 + \sigma^2 \cdot \textstyle\sum_{g \in \mathbf{O}_j^\dagger} (w_{jg}^\dagger)^2),
\end{aligned}
$$

where $\{\mathbf{O}_j^\dagger, w_{jg}^\dagger\}$ are annotations of the random edge-altered network. For all datasets the target of enrichment testing is the actual B cell network.

The caption of each panel in this supplementary figure shows the true hyper-parameter values of positive datasets. The sparse scenario in **Figure 3(d)** corresponds to simulations with true hyper-parameter values of positive datasets being $\theta_0 = -4$, $\theta = 2$, $\sigma_0 = 1$, $\sigma = 0.5$ and PVE = 0.6. The polygenic scenario in **Figure 3(d)** corresponds to simulations with true hyper-parameter values of positive datasets being $\theta_0 = -2$, $\theta = 2$, $\sigma_0 = 1$, $\sigma = 0.5$ and PVE = 0.6.

The true and random edge-altered networks have the same associated REs and member genes but totally different topology (TF-TG edges and edge weights). Existing methods like LDSC and Pascal only exploits proximity between SNPs and network genes and/or REs, and thus they cannot distinguish the true and random edge-altered networks. In contrast, because of the edge-enrichment parameter $\sigma^2$, RSS-NET is able to capture the topological differences among networks, and thus it can reliably distinguish true enrichments of the B cell network from enrichments of its edge-altered counterparts.

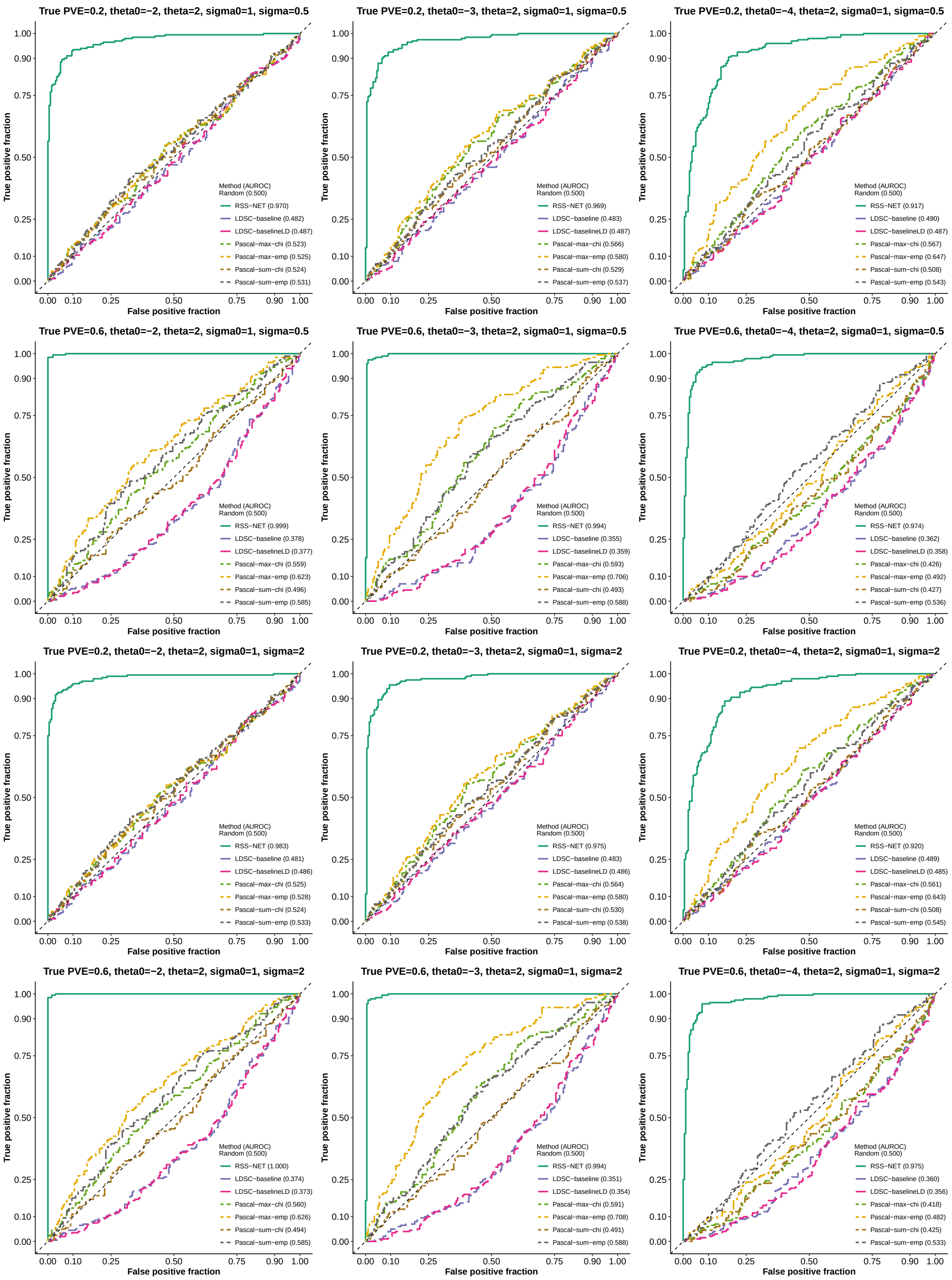
True PVE=0.2, theta0=−2, theta=2, sigma0=1, sigma=0.5
Method (AUROC)
Random (0.500)
RSS−NET (0.970)
LDSC−baseline (0.482)
LDSC−baselineLD (0.487)
Pascal−max−chi (0.523)
Pascal−max−emp (0.525)
Pascal−sum−chi (0.524)
Pascal−sum−emp (0.531)
True PVE=0.2, theta0=−3, theta=2, sigma0=1, sigma=0.5
Method (AUROC)
Random (0.500)
RSS−NET (0.969)
LDSC−baseline (0.483)
LDSC−baselineLD (0.487)
Pascal−max−chi (0.566)
Pascal−max−emp (0.580)
Pascal−sum−chi (0.529)
Pascal−sum−emp (0.537)
True PVE=0.2, theta0=−4, theta=2, sigma0=1, sigma=0.5
Method (AUROC)
Random (0.500)
RSS−NET (0.917)
LDSC−baseline (0.490)
LDSC−baselineLD (0.487)
Pascal−max−chi (0.567)
Pascal−max−emp (0.647)
Pascal−sum−chi (0.508)
Pascal−sum−emp (0.543)
True PVE=0.6, theta0=−2, theta=2, sigma0=1, sigma=0.5
Method (AUROC)
Random (0.500)
RSS−NET (0.999)
LDSC−baseline (0.378)
LDSC−baselineLD (0.377)
Pascal−max−chi (0.559)
Pascal−max−emp (0.623)
Pascal−sum−chi (0.496)
Pascal−sum−emp (0.585)
True PVE=0.6, theta0=−3, theta=2, sigma0=1, sigma=0.5
Method (AUROC)
Random (0.500)
RSS−NET (0.994)
LDSC−baseline (0.355)
LDSC−baselineLD (0.359)
Pascal−max−chi (0.593)
Pascal−max−emp (0.706)
Pascal−sum−chi (0.493)
Pascal−sum−emp (0.588)
True PVE=0.6, theta0=−4, theta=2, sigma0=1, sigma=0.5
Method (AUROC)
Random (0.500)
RSS−NET (0.974)
LDSC−baseline (0.362)
LDSC−baselineLD (0.358)
Pascal−max−chi (0.426)
Pascal−max−emp (0.492)
Pascal−sum−chi (0.427)
Pascal−sum−emp (0.536)
True PVE=0.2, theta0=−2, theta=2, sigma0=1, sigma=2
Method (AUROC)
Random (0.500)
RSS−NET (0.983)
LDSC−baseline (0.481)
LDSC−baselineLD (0.486)
Pascal−max−chi (0.525)
Pascal−max−emp (0.528)
Pascal−sum−chi (0.524)
Pascal−sum−emp (0.533)
True PVE=0.2, theta0=−3, theta=2, sigma0=1, sigma=2
Method (AUROC)
Random (0.500)
RSS−NET (0.975)
LDSC−baseline (0.483)
LDSC−baselineLD (0.486)
Pascal−max−chi (0.564)
Pascal−max−emp (0.580)
Pascal−sum−chi (0.530)
Pascal−sum−emp (0.538)
True PVE=0.2, theta0=−4, theta=2, sigma0=1, sigma=2
Method (AUROC)
Random (0.500)
RSS−NET (0.920)
LDSC−baseline (0.489)
LDSC−baselineLD (0.485)
Pascal−max−chi (0.561)
Pascal−max−emp (0.643)
Pascal−sum−chi (0.508)
Pascal−sum−emp (0.545)
True PVE=0.6, theta0=−2, theta=2, sigma0=1, sigma=2
Method (AUROC)
Random (0.500)
RSS−NET (1.000)
LDSC−baseline (0.374)
LDSC−baselineLD (0.373)
Pascal−max−chi (0.560)
Pascal−max−emp (0.626)
Pascal−sum−chi (0.494)
Pascal−sum−emp (0.585)
True PVE=0.6, theta0=−3, theta=2, sigma0=1, sigma=2
Method (AUROC)
Random (0.500)
RSS−NET (0.994)
LDSC−baseline (0.351)
LDSC−baselineLD (0.354)
Pascal−max−chi (0.591)
Pascal−max−emp (0.708)
Pascal−sum−chi (0.491)
Pascal−sum−emp (0.588)
True PVE=0.6, theta0=−4, theta=2, sigma0=1, sigma=2
Method (AUROC)
Random (0.500)
RSS−NET (0.975)
LDSC−baseline (0.360)
LDSC−baselineLD (0.356)
Pascal−max−chi (0.418)
Pascal−max−emp (0.482)
Pascal−sum−chi (0.425)
Pascal−sum−emp (0.533)
True positive fraction
False positive fraction
0.00
0.10
0.25
0.50
0.75
0.90
1.00

#### Supplementary Figure 7

**Simulation details and additional results of Figure 4.** Here we compare RSS-NET with existing gene-level association testing methods on the same simulated summary data used in **Supplementary Figure 1**. Details of this part are almost identical to those in **Supplementary Figure 1**. Here we only highlight the differences.

For each simulated dataset, we define a gene as "trait-associated" if at lease one SNP $j$ within 100 kb of the transcribed region of this gene has non-zero effect ($\beta_j \neq 0$). For each gene in each simulated dataset, RSS-NET and RSS-E methods (Zhu and Stephens 2018) produce $P_1$, the posterior probability that the gene is trait-associated, whereas Pascal methods (Lamparter et al. 2016) produce $P$-value for the null hypothesis that the gene is not trait-associated; these statistics are used to rank the significance of gene-level associations. If a method identifies association between a non-trait-associated gene and the trait, then it is a "false positive". If a method identifies association between a trait-associated gene and the trait, then it is a "true positive".

Here the true values of baseline proportion parameter $\theta_0$ are $\{-2,-3,-4\}$, the true values of enrichment proportion parameter $\theta$ are $\{0,2\}$, the true value of baseline magnitude parameter $\sigma_0$ is 1, the true values of enrichment magnitude parameter $\sigma$ are $\{0,0.5,2\}$, and the true values of PVE are $\{0.2,0.6\}$. In total there are 36 ($= 3\times 2\times 3\times 2$) simulation scenarios. Each scenario contains 200 independent datasets. For all datasets we test associations of 16,954 autosomal protein-coding genes that are available in Pascal-required database.

The caption of each panel in this supplementary figure shows the true hyper-parameter values of simulated datasets. All panels of **Figure 4** correspond to simulations with true $\theta_0 = -4$, $\sigma_0 = 1$ and PVE = 0.6. **Figure 4(a)** corresponds to simulations with true $\theta = 0$ and $\sigma = 0$. **Figure 4(b)** corresponds to simulations with true $\theta = 2$ and $\sigma = 0$. **Figure 4(c)** corresponds to simulations with true $\theta = 0$ and $\sigma = 2$. **Figure 4(d)** corresponds to simulations with true $\theta = 2$ and $\sigma = 2$.

We evaluate the performance of these gene-level association methods by plotting the receiver operating characteristic (ROC) curve and computing the area under the ROC curve (AUROC) for each method. Both metrics are implemented in the R package `plotROC` (Sachs 2017). Unlike the balanced enrichment simulations (200 positive and 200 negative datasets in each scenario), the numbers of trait-associated genes (true labels) and non-trait-associated genes (false labels) are often very different. Due to this imbalance nature, we also plot the precision-recall (PRC) curve and compute the area under the PRC curve (AUPRC) for each method. Both metrics are implemented in the R package `precrec` (Saito and Rehmsmeier 2017).

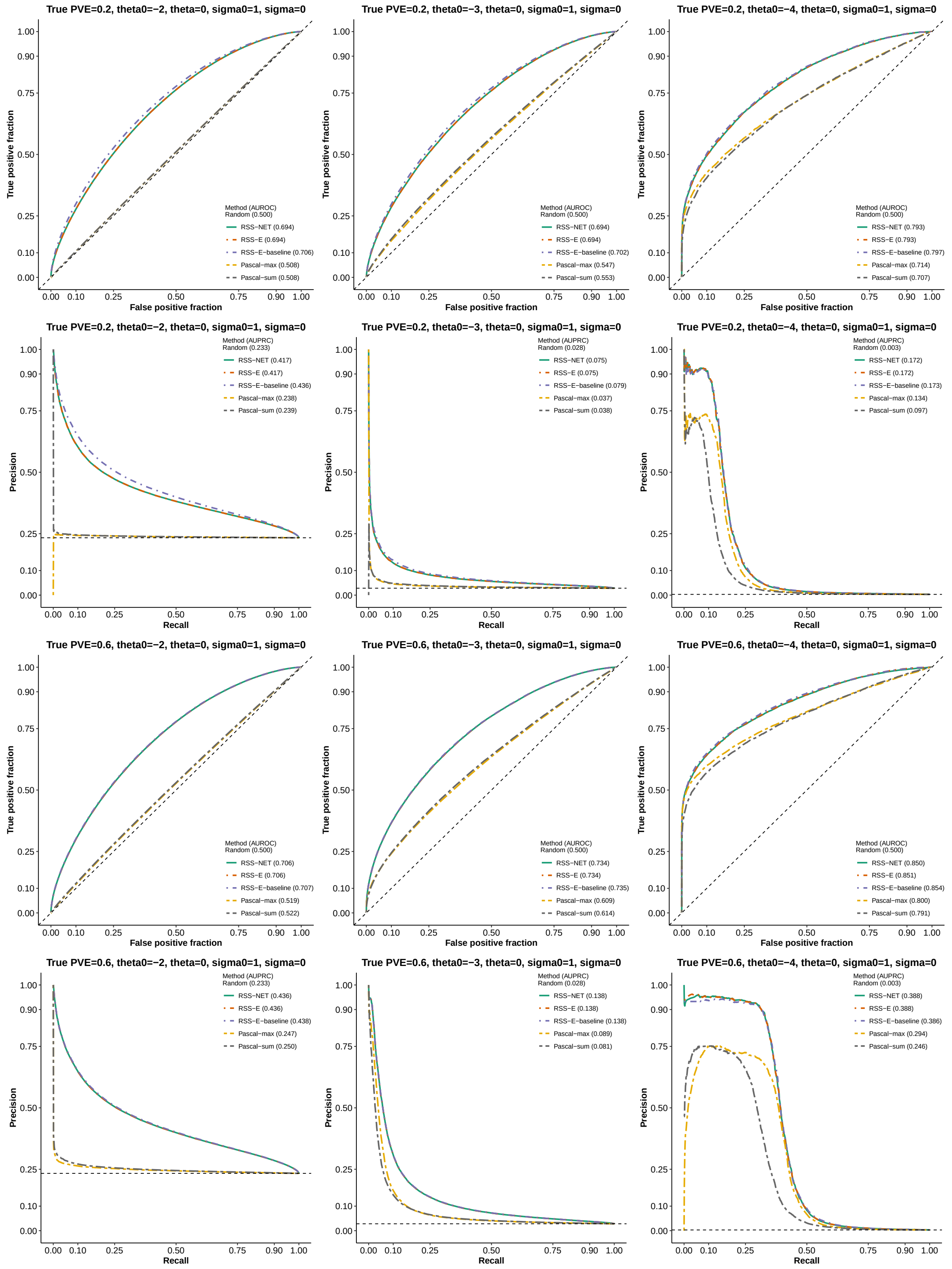
True PVE=0.2, theta0=−2, theta=0, sigma0=1, sigma=0
True positive fraction
False positive fraction
Method (AUROC)
Random (0.500)
RSS−NET (0.694)
RSS−E (0.694)
RSS−E−baseline (0.706)
Pascal−max (0.508)
Pascal−sum (0.508)
True PVE=0.2, theta0=−3, theta=0, sigma0=1, sigma=0
Method (AUROC)
Random (0.500)
RSS−NET (0.694)
RSS−E (0.694)
RSS−E−baseline (0.702)
Pascal−max (0.547)
Pascal−sum (0.553)
True PVE=0.2, theta0=−4, theta=0, sigma0=1, sigma=0
Method (AUROC)
Random (0.500)
RSS−NET (0.793)
RSS−E (0.793)
RSS−E−baseline (0.797)
Pascal−max (0.714)
Pascal−sum (0.707)
True PVE=0.2, theta0=−2, theta=0, sigma0=1, sigma=0
Precision
Recall
Method (AUPRC)
Random (0.233)
RSS−NET (0.417)
RSS−E (0.417)
RSS−E−baseline (0.436)
Pascal−max (0.238)
Pascal−sum (0.239)
True PVE=0.2, theta0=−3, theta=0, sigma0=1, sigma=0
Method (AUPRC)
Random (0.028)
RSS−NET (0.075)
RSS−E (0.075)
RSS−E−baseline (0.079)
Pascal−max (0.037)
Pascal−sum (0.038)
True PVE=0.2, theta0=−4, theta=0, sigma0=1, sigma=0
Method (AUPRC)
Random (0.003)
RSS−NET (0.172)
RSS−E (0.172)
RSS−E−baseline (0.173)
Pascal−max (0.134)
Pascal−sum (0.097)
True PVE=0.6, theta0=−2, theta=0, sigma0=1, sigma=0
Method (AUROC)
Random (0.500)
RSS−NET (0.706)
RSS−E (0.706)
RSS−E−baseline (0.707)
Pascal−max (0.519)
Pascal−sum (0.522)
True PVE=0.6, theta0=−3, theta=0, sigma0=1, sigma=0
Method (AUROC)
Random (0.500)
RSS−NET (0.734)
RSS−E (0.734)
RSS−E−baseline (0.735)
Pascal−max (0.609)
Pascal−sum (0.614)
True PVE=0.6, theta0=−4, theta=0, sigma0=1, sigma=0
Method (AUROC)
Random (0.500)
RSS−NET (0.850)
RSS−E (0.851)
RSS−E−baseline (0.854)
Pascal−max (0.800)
Pascal−sum (0.791)
True PVE=0.6, theta0=−2, theta=0, sigma0=1, sigma=0
Method (AUPRC)
Random (0.233)
RSS−NET (0.436)
RSS−E (0.436)
RSS−E−baseline (0.438)
Pascal−max (0.247)
Pascal−sum (0.250)
True PVE=0.6, theta0=−3, theta=0, sigma0=1, sigma=0
Method (AUPRC)
Random (0.028)
RSS−NET (0.138)
RSS−E (0.138)
RSS−E−baseline (0.138)
Pascal−max (0.089)
Pascal−sum (0.081)
True PVE=0.6, theta0=−4, theta=0, sigma0=1, sigma=0
Method (AUPRC)
Random (0.003)
RSS−NET (0.388)
RSS−E (0.388)
RSS−E−baseline (0.386)
Pascal−max (0.294)
Pascal−sum (0.246)

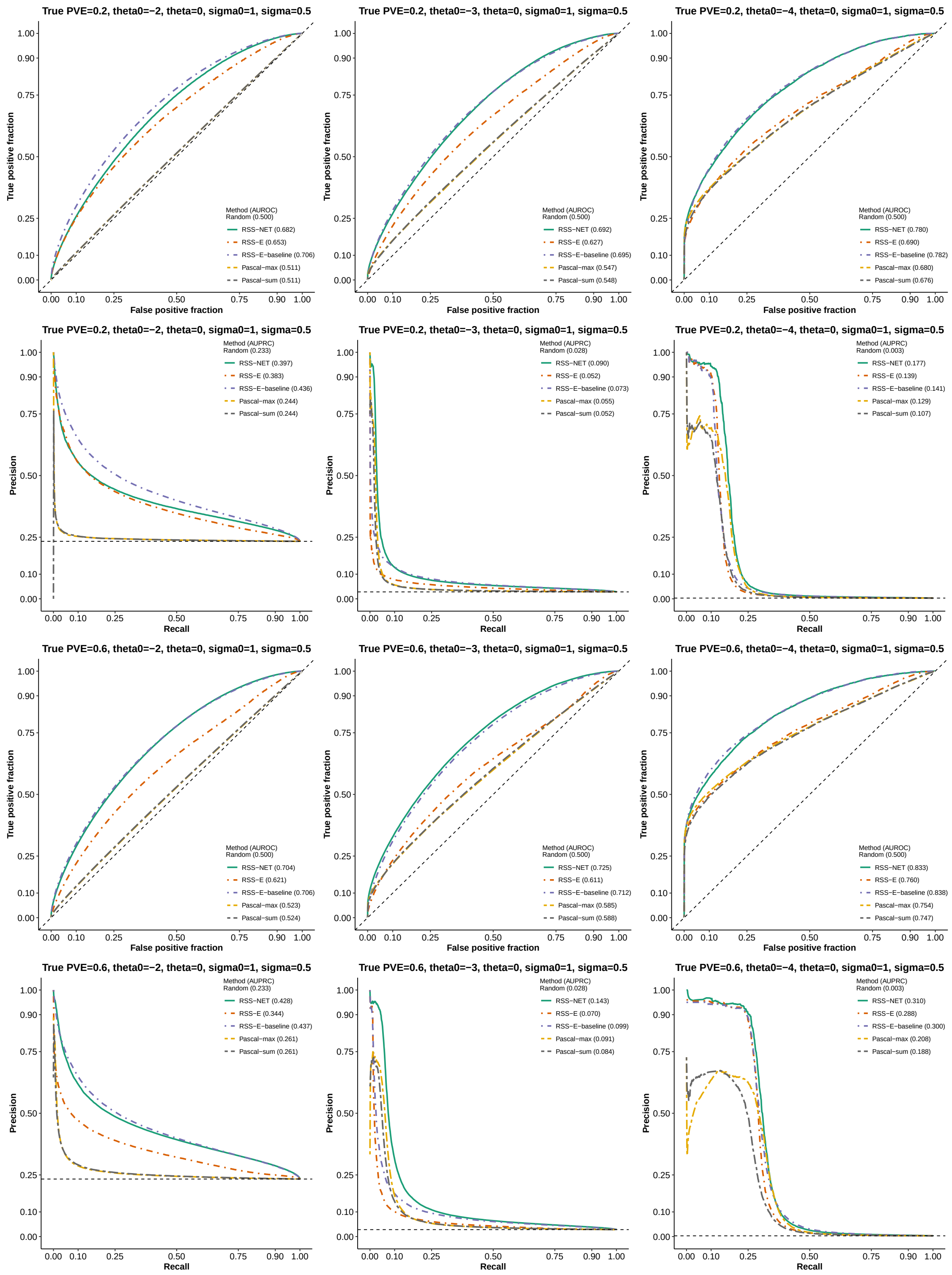
True PVE=0.2, theta0=−2, theta=0, sigma0=1, sigma=0.5
True positive fraction
False positive fraction
Method (AUROC)
Random (0.500)
RSS−NET (0.682)
RSS−E (0.653)
RSS−E−baseline (0.706)
Pascal−max (0.511)
Pascal−sum (0.511)
True PVE=0.2, theta0=−3, theta=0, sigma0=1, sigma=0.5
Method (AUROC)
Random (0.500)
RSS−NET (0.692)
RSS−E (0.627)
RSS−E−baseline (0.695)
Pascal−max (0.547)
Pascal−sum (0.548)
True PVE=0.2, theta0=−4, theta=0, sigma0=1, sigma=0.5
Method (AUROC)
Random (0.500)
RSS−NET (0.780)
RSS−E (0.690)
RSS−E−baseline (0.782)
Pascal−max (0.680)
Pascal−sum (0.676)
True PVE=0.2, theta0=−2, theta=0, sigma0=1, sigma=0.5
Precision
Recall
Method (AUPRC)
Random (0.233)
RSS−NET (0.397)
RSS−E (0.383)
RSS−E−baseline (0.436)
Pascal−max (0.244)
Pascal−sum (0.244)
True PVE=0.2, theta0=−3, theta=0, sigma0=1, sigma=0.5
Method (AUPRC)
Random (0.028)
RSS−NET (0.090)
RSS−E (0.052)
RSS−E−baseline (0.073)
Pascal−max (0.055)
Pascal−sum (0.052)
True PVE=0.2, theta0=−4, theta=0, sigma0=1, sigma=0.5
Method (AUPRC)
Random (0.003)
RSS−NET (0.177)
RSS−E (0.139)
RSS−E−baseline (0.141)
Pascal−max (0.129)
Pascal−sum (0.107)
True PVE=0.6, theta0=−2, theta=0, sigma0=1, sigma=0.5
Method (AUROC)
Random (0.500)
RSS−NET (0.704)
RSS−E (0.621)
RSS−E−baseline (0.706)
Pascal−max (0.523)
Pascal−sum (0.524)
True PVE=0.6, theta0=−3, theta=0, sigma0=1, sigma=0.5
Method (AUROC)
Random (0.500)
RSS−NET (0.725)
RSS−E (0.611)
RSS−E−baseline (0.712)
Pascal−max (0.585)
Pascal−sum (0.588)
True PVE=0.6, theta0=−4, theta=0, sigma0=1, sigma=0.5
Method (AUROC)
Random (0.500)
RSS−NET (0.833)
RSS−E (0.760)
RSS−E−baseline (0.838)
Pascal−max (0.754)
Pascal−sum (0.747)
True PVE=0.6, theta0=−2, theta=0, sigma0=1, sigma=0.5
Method (AUPRC)
Random (0.233)
RSS−NET (0.428)
RSS−E (0.344)
RSS−E−baseline (0.437)
Pascal−max (0.261)
Pascal−sum (0.261)
True PVE=0.6, theta0=−3, theta=0, sigma0=1, sigma=0.5
Method (AUPRC)
Random (0.028)
RSS−NET (0.143)
RSS−E (0.070)
RSS−E−baseline (0.099)
Pascal−max (0.091)
Pascal−sum (0.084)
True PVE=0.6, theta0=−4, theta=0, sigma0=1, sigma=0.5
Method (AUPRC)
Random (0.003)
RSS−NET (0.310)
RSS−E (0.288)
RSS−E−baseline (0.300)
Pascal−max (0.208)
Pascal−sum (0.188)

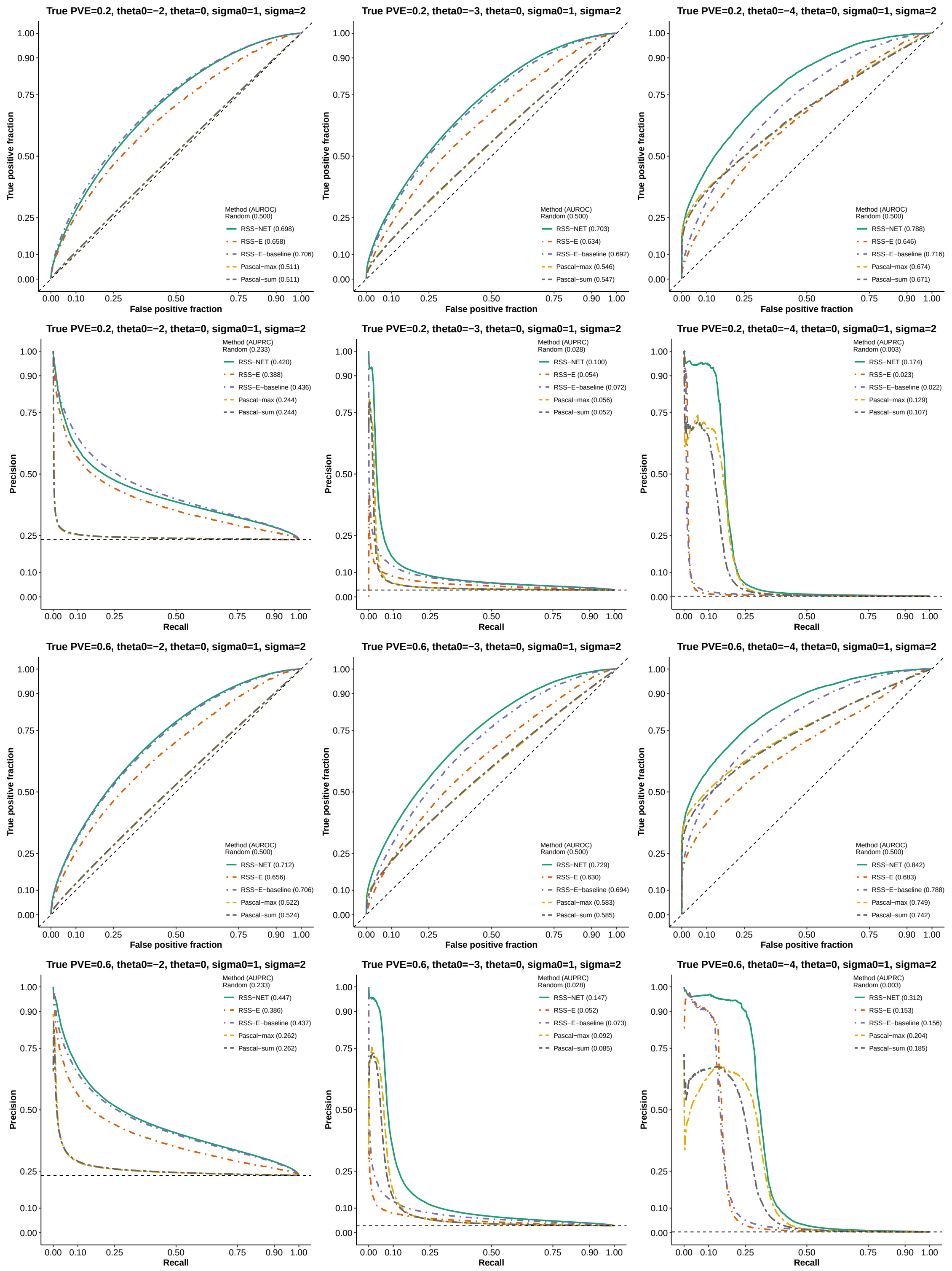
True PVE=0.2, theta0=−2, theta=0, sigma0=1, sigma=2
True positive fraction
False positive fraction
Method (AUROC)
Random (0.500)
RSS−NET (0.698)
RSS−E (0.658)
RSS−E−baseline (0.706)
Pascal−max (0.511)
Pascal−sum (0.511)
True PVE=0.2, theta0=−3, theta=0, sigma0=1, sigma=2
Method (AUROC)
Random (0.500)
RSS−NET (0.703)
RSS−E (0.634)
RSS−E−baseline (0.692)
Pascal−max (0.546)
Pascal−sum (0.547)
True PVE=0.2, theta0=−4, theta=0, sigma0=1, sigma=2
Method (AUROC)
Random (0.500)
RSS−NET (0.788)
RSS−E (0.646)
RSS−E−baseline (0.716)
Pascal−max (0.674)
Pascal−sum (0.671)
True PVE=0.2, theta0=−2, theta=0, sigma0=1, sigma=2
Precision
Recall
Method (AUPRC)
Random (0.233)
RSS−NET (0.420)
RSS−E (0.388)
RSS−E−baseline (0.436)
Pascal−max (0.244)
Pascal−sum (0.244)
True PVE=0.2, theta0=−3, theta=0, sigma0=1, sigma=2
Method (AUPRC)
Random (0.028)
RSS−NET (0.100)
RSS−E (0.054)
RSS−E−baseline (0.072)
Pascal−max (0.056)
Pascal−sum (0.052)
True PVE=0.2, theta0=−4, theta=0, sigma0=1, sigma=2
Method (AUPRC)
Random (0.003)
RSS−NET (0.174)
RSS−E (0.023)
RSS−E−baseline (0.022)
Pascal−max (0.129)
Pascal−sum (0.107)
True PVE=0.6, theta0=−2, theta=0, sigma0=1, sigma=2
Method (AUROC)
Random (0.500)
RSS−NET (0.712)
RSS−E (0.656)
RSS−E−baseline (0.706)
Pascal−max (0.522)
Pascal−sum (0.524)
True PVE=0.6, theta0=−3, theta=0, sigma0=1, sigma=2
Method (AUROC)
Random (0.500)
RSS−NET (0.729)
RSS−E (0.630)
RSS−E−baseline (0.694)
Pascal−max (0.583)
Pascal−sum (0.585)
True PVE=0.6, theta0=−4, theta=0, sigma0=1, sigma=2
Method (AUROC)
Random (0.500)
RSS−NET (0.842)
RSS−E (0.683)
RSS−E−baseline (0.788)
Pascal−max (0.749)
Pascal−sum (0.742)
True PVE=0.6, theta0=−2, theta=0, sigma0=1, sigma=2
Method (AUPRC)
Random (0.233)
RSS−NET (0.447)
RSS−E (0.386)
RSS−E−baseline (0.437)
Pascal−max (0.262)
Pascal−sum (0.262)
True PVE=0.6, theta0=−3, theta=0, sigma0=1, sigma=2
Method (AUPRC)
Random (0.028)
RSS−NET (0.147)
RSS−E (0.052)
RSS−E−baseline (0.073)
Pascal−max (0.092)
Pascal−sum (0.085)
True PVE=0.6, theta0=−4, theta=0, sigma0=1, sigma=2
Method (AUPRC)
Random (0.003)
RSS−NET (0.312)
RSS−E (0.153)
RSS−E−baseline (0.156)
Pascal−max (0.204)
Pascal−sum (0.185)

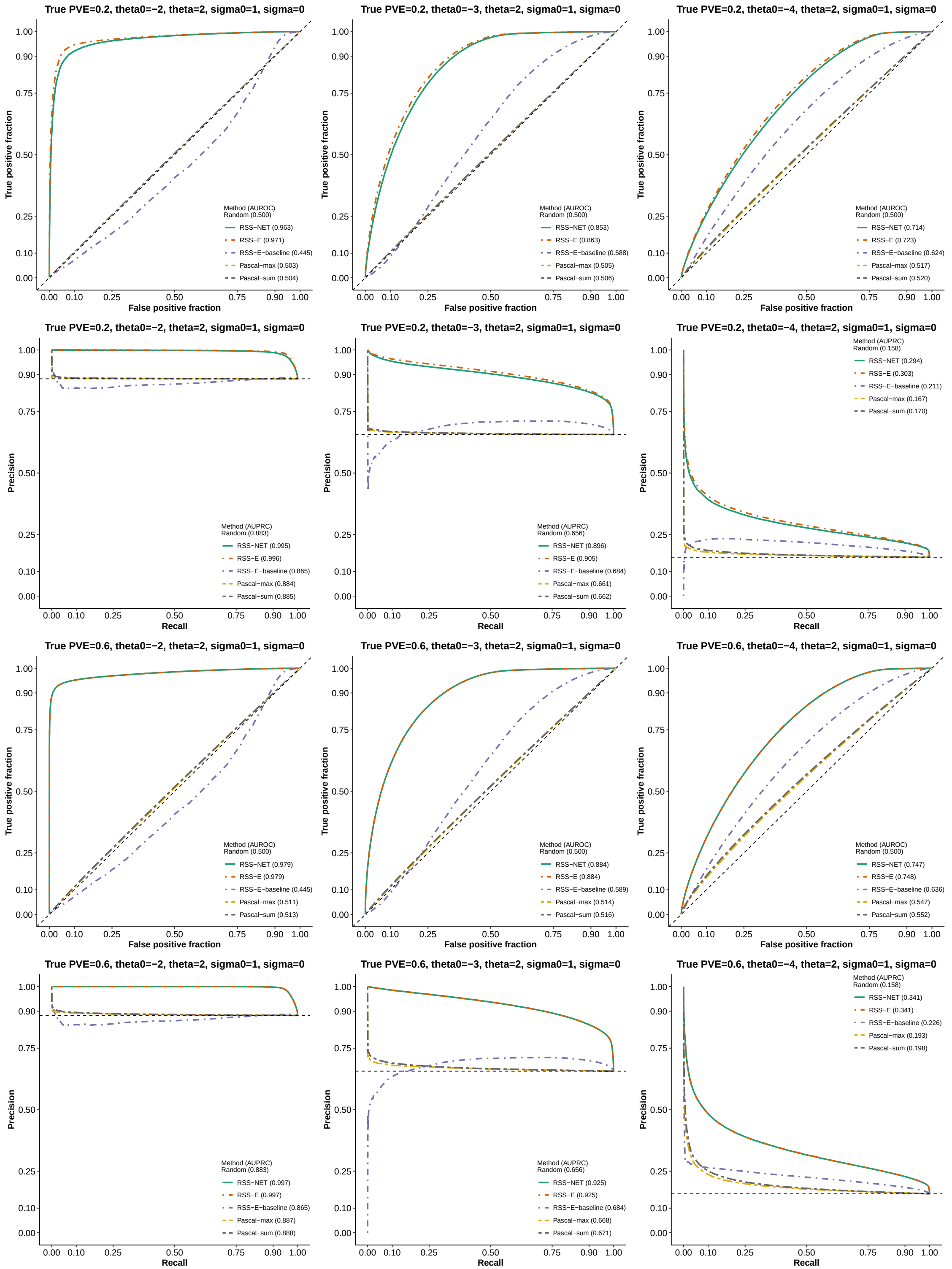
True PVE=0.2, theta0=−2, theta=2, sigma0=1, sigma=0
True positive fraction
False positive fraction
Method (AUROC)
Random (0.500)
RSS−NET (0.963)
RSS−E (0.971)
RSS−E−baseline (0.445)
Pascal−max (0.503)
Pascal−sum (0.504)
True PVE=0.2, theta0=−3, theta=2, sigma0=1, sigma=0
Method (AUROC)
Random (0.500)
RSS−NET (0.853)
RSS−E (0.863)
RSS−E−baseline (0.588)
Pascal−max (0.505)
Pascal−sum (0.506)
True PVE=0.2, theta0=−4, theta=2, sigma0=1, sigma=0
Method (AUROC)
Random (0.500)
RSS−NET (0.714)
RSS−E (0.723)
RSS−E−baseline (0.624)
Pascal−max (0.517)
Pascal−sum (0.520)
True PVE=0.2, theta0=−2, theta=2, sigma0=1, sigma=0
Precision
Recall
Method (AUPRC)
Random (0.883)
RSS−NET (0.995)
RSS−E (0.996)
RSS−E−baseline (0.865)
Pascal−max (0.884)
Pascal−sum (0.885)
True PVE=0.2, theta0=−3, theta=2, sigma0=1, sigma=0
Method (AUPRC)
Random (0.656)
RSS−NET (0.896)
RSS−E (0.905)
RSS−E−baseline (0.684)
Pascal−max (0.661)
Pascal−sum (0.662)
True PVE=0.2, theta0=−4, theta=2, sigma0=1, sigma=0
Method (AUPRC)
Random (0.158)
RSS−NET (0.294)
RSS−E (0.303)
RSS−E−baseline (0.211)
Pascal−max (0.167)
Pascal−sum (0.170)
True PVE=0.6, theta0=−2, theta=2, sigma0=1, sigma=0
Method (AUROC)
Random (0.500)
RSS−NET (0.979)
RSS−E (0.979)
RSS−E−baseline (0.445)
Pascal−max (0.511)
Pascal−sum (0.513)
True PVE=0.6, theta0=−3, theta=2, sigma0=1, sigma=0
Method (AUROC)
Random (0.500)
RSS−NET (0.884)
RSS−E (0.884)
RSS−E−baseline (0.589)
Pascal−max (0.514)
Pascal−sum (0.516)
True PVE=0.6, theta0=−4, theta=2, sigma0=1, sigma=0
Method (AUROC)
Random (0.500)
RSS−NET (0.747)
RSS−E (0.748)
RSS−E−baseline (0.636)
Pascal−max (0.547)
Pascal−sum (0.552)
True PVE=0.6, theta0=−2, theta=2, sigma0=1, sigma=0
Method (AUPRC)
Random (0.883)
RSS−NET (0.997)
RSS−E (0.997)
RSS−E−baseline (0.865)
Pascal−max (0.887)
Pascal−sum (0.888)
True PVE=0.6, theta0=−3, theta=2, sigma0=1, sigma=0
Method (AUPRC)
Random (0.656)
RSS−NET (0.925)
RSS−E (0.925)
RSS−E−baseline (0.684)
Pascal−max (0.668)
Pascal−sum (0.671)
True PVE=0.6, theta0=−4, theta=2, sigma0=1, sigma=0
Method (AUPRC)
Random (0.158)
RSS−NET (0.341)
RSS−E (0.341)
RSS−E−baseline (0.226)
Pascal−max (0.193)
Pascal−sum (0.198)

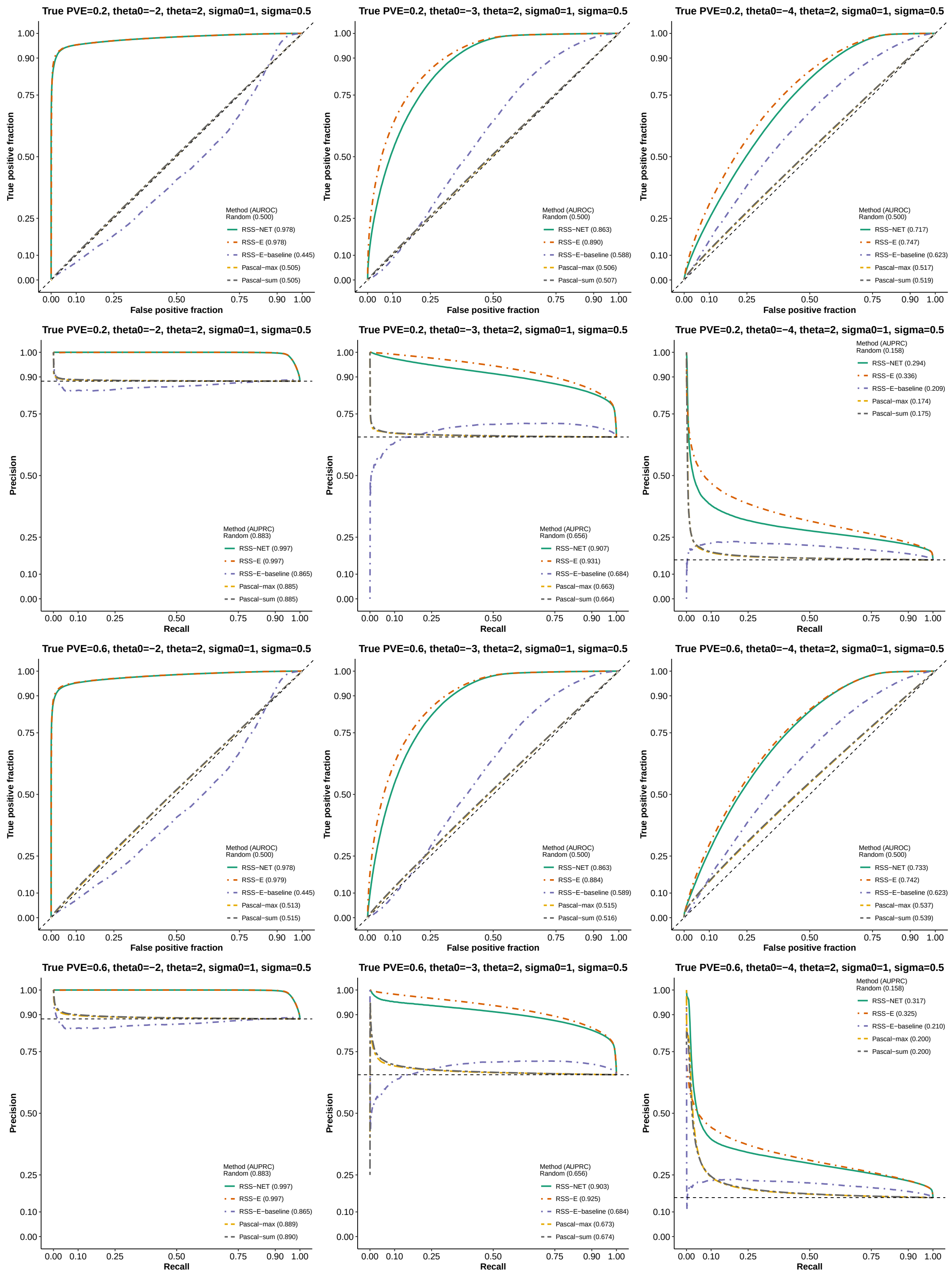
True PVE=0.2, theta0=−2, theta=2, sigma0=1, sigma=0.5
True positive fraction
False positive fraction
Method (AUROC)
Random (0.500)
RSS−NET (0.978)
RSS−E (0.978)
RSS−E−baseline (0.445)
Pascal−max (0.505)
Pascal−sum (0.505)
True PVE=0.2, theta0=−3, theta=2, sigma0=1, sigma=0.5
True positive fraction
False positive fraction
Method (AUROC)
Random (0.500)
RSS−NET (0.863)
RSS−E (0.890)
RSS−E−baseline (0.588)
Pascal−max (0.506)
Pascal−sum (0.507)
True PVE=0.2, theta0=−4, theta=2, sigma0=1, sigma=0.5
True positive fraction
False positive fraction
Method (AUROC)
Random (0.500)
RSS−NET (0.717)
RSS−E (0.747)
RSS−E−baseline (0.623)
Pascal−max (0.517)
Pascal−sum (0.519)
True PVE=0.2, theta0=−2, theta=2, sigma0=1, sigma=0.5
Precision
Recall
Method (AUPRC)
Random (0.883)
RSS−NET (0.997)
RSS−E (0.997)
RSS−E−baseline (0.865)
Pascal−max (0.885)
Pascal−sum (0.885)
True PVE=0.2, theta0=−3, theta=2, sigma0=1, sigma=0.5
Precision
Recall
Method (AUPRC)
Random (0.656)
RSS−NET (0.907)
RSS−E (0.931)
RSS−E−baseline (0.684)
Pascal−max (0.663)
Pascal−sum (0.664)
True PVE=0.2, theta0=−4, theta=2, sigma0=1, sigma=0.5
Precision
Recall
Method (AUPRC)
Random (0.158)
RSS−NET (0.294)
RSS−E (0.336)
RSS−E−baseline (0.209)
Pascal−max (0.174)
Pascal−sum (0.175)
True PVE=0.6, theta0=−2, theta=2, sigma0=1, sigma=0.5
True positive fraction
False positive fraction
Method (AUROC)
Random (0.500)
RSS−NET (0.978)
RSS−E (0.979)
RSS−E−baseline (0.445)
Pascal−max (0.513)
Pascal−sum (0.515)
True PVE=0.6, theta0=−3, theta=2, sigma0=1, sigma=0.5
True positive fraction
False positive fraction
Method (AUROC)
Random (0.500)
RSS−NET (0.863)
RSS−E (0.884)
RSS−E−baseline (0.589)
Pascal−max (0.515)
Pascal−sum (0.516)
True PVE=0.6, theta0=−4, theta=2, sigma0=1, sigma=0.5
True positive fraction
False positive fraction
Method (AUROC)
Random (0.500)
RSS−NET (0.733)
RSS−E (0.742)
RSS−E−baseline (0.623)
Pascal−max (0.537)
Pascal−sum (0.539)
True PVE=0.6, theta0=−2, theta=2, sigma0=1, sigma=0.5
Precision
Recall
Method (AUPRC)
Random (0.883)
RSS−NET (0.997)
RSS−E (0.997)
RSS−E−baseline (0.865)
Pascal−max (0.889)
Pascal−sum (0.890)
True PVE=0.6, theta0=−3, theta=2, sigma0=1, sigma=0.5
Precision
Recall
Method (AUPRC)
Random (0.656)
RSS−NET (0.903)
RSS−E (0.925)
RSS−E−baseline (0.684)
Pascal−max (0.673)
Pascal−sum (0.674)
True PVE=0.6, theta0=−4, theta=2, sigma0=1, sigma=0.5
Precision
Recall
Method (AUPRC)
Random (0.158)
RSS−NET (0.317)
RSS−E (0.325)
RSS−E−baseline (0.210)
Pascal−max (0.200)
Pascal−sum (0.200)

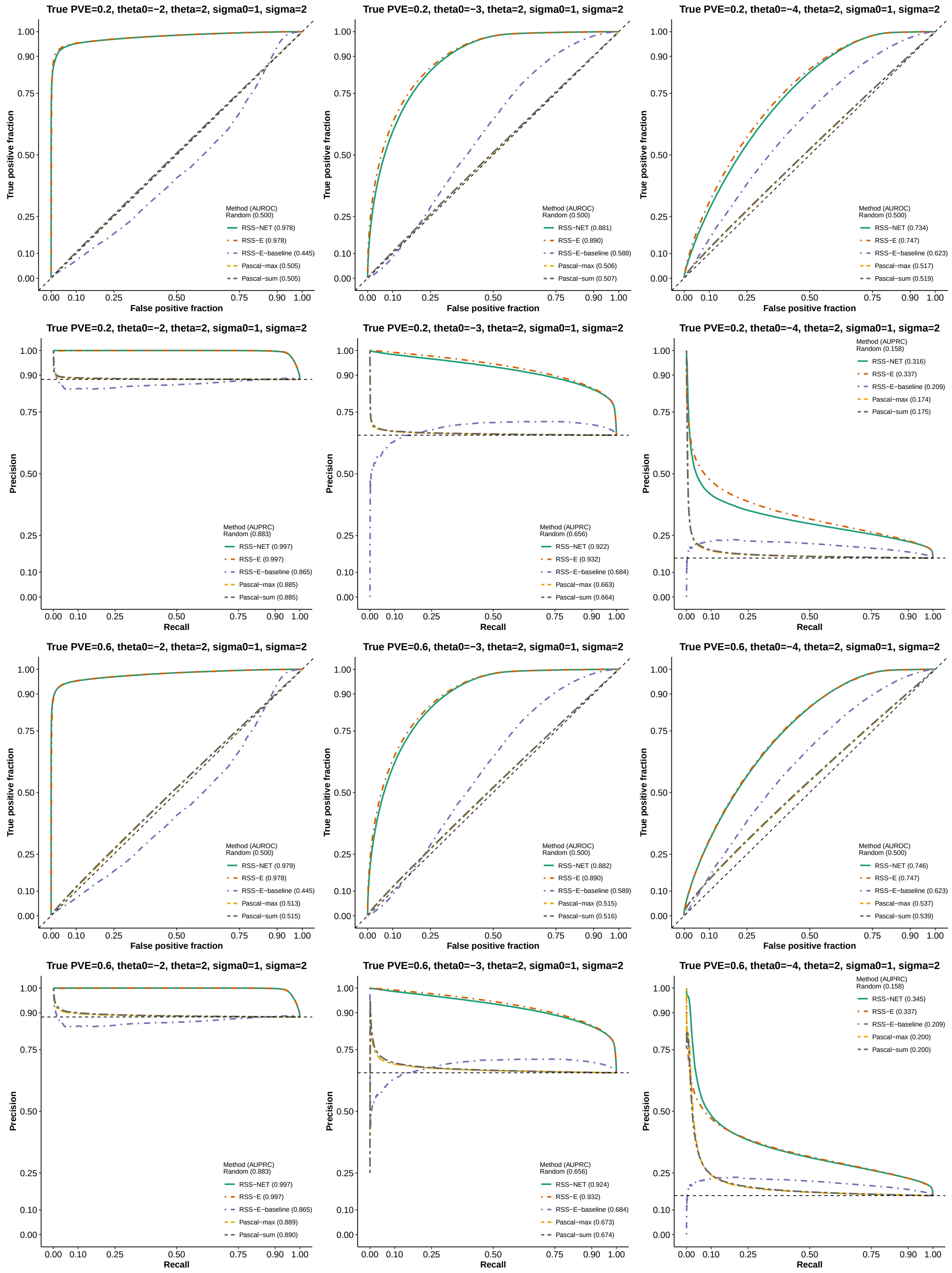
True PVE=0.2, theta0=−2, theta=2, sigma0=1, sigma=2
True positive fraction
False positive fraction
Method (AUROC)
Random (0.500)
RSS−NET (0.978)
RSS−E (0.978)
RSS−E−baseline (0.445)
Pascal−max (0.505)
Pascal−sum (0.505)
True PVE=0.2, theta0=−3, theta=2, sigma0=1, sigma=2
Method (AUROC)
Random (0.500)
RSS−NET (0.881)
RSS−E (0.890)
RSS−E−baseline (0.588)
Pascal−max (0.506)
Pascal−sum (0.507)
True PVE=0.2, theta0=−4, theta=2, sigma0=1, sigma=2
Method (AUROC)
Random (0.500)
RSS−NET (0.734)
RSS−E (0.747)
RSS−E−baseline (0.623)
Pascal−max (0.517)
Pascal−sum (0.519)
True PVE=0.2, theta0=−2, theta=2, sigma0=1, sigma=2
Precision
Recall
Method (AUPRC)
Random (0.883)
RSS−NET (0.997)
RSS−E (0.997)
RSS−E−baseline (0.865)
Pascal−max (0.885)
Pascal−sum (0.885)
True PVE=0.2, theta0=−3, theta=2, sigma0=1, sigma=2
Method (AUPRC)
Random (0.656)
RSS−NET (0.922)
RSS−E (0.932)
RSS−E−baseline (0.684)
Pascal−max (0.663)
Pascal−sum (0.664)
True PVE=0.2, theta0=−4, theta=2, sigma0=1, sigma=2
Method (AUPRC)
Random (0.158)
RSS−NET (0.316)
RSS−E (0.337)
RSS−E−baseline (0.209)
Pascal−max (0.174)
Pascal−sum (0.175)
True PVE=0.6, theta0=−2, theta=2, sigma0=1, sigma=2
Method (AUROC)
Random (0.500)
RSS−NET (0.979)
RSS−E (0.978)
RSS−E−baseline (0.445)
Pascal−max (0.513)
Pascal−sum (0.515)
True PVE=0.6, theta0=−3, theta=2, sigma0=1, sigma=2
Method (AUROC)
Random (0.500)
RSS−NET (0.882)
RSS−E (0.890)
RSS−E−baseline (0.589)
Pascal−max (0.515)
Pascal−sum (0.516)
True PVE=0.6, theta0=−4, theta=2, sigma0=1, sigma=2
Method (AUROC)
Random (0.500)
RSS−NET (0.746)
RSS−E (0.747)
RSS−E−baseline (0.623)
Pascal−max (0.537)
Pascal−sum (0.539)
True PVE=0.6, theta0=−2, theta=2, sigma0=1, sigma=2
Method (AUPRC)
Random (0.883)
RSS−NET (0.997)
RSS−E (0.997)
RSS−E−baseline (0.865)
Pascal−max (0.889)
Pascal−sum (0.890)
True PVE=0.6, theta0=−3, theta=2, sigma0=1, sigma=2
Method (AUPRC)
Random (0.656)
RSS−NET (0.924)
RSS−E (0.932)
RSS−E−baseline (0.684)
Pascal−max (0.673)
Pascal−sum (0.674)
True PVE=0.6, theta0=−4, theta=2, sigma0=1, sigma=2
Method (AUPRC)
Random (0.158)
RSS−NET (0.345)
RSS−E (0.337)
RSS−E−baseline (0.209)
Pascal−max (0.200)
Pascal−sum (0.200)

#### Supplementary Figure 8

We repeat the simulations in **Supplementary Figure 7** on GWAS summary statistics simulated in **Supplementary Figure 2** and the vagina regulatory network.

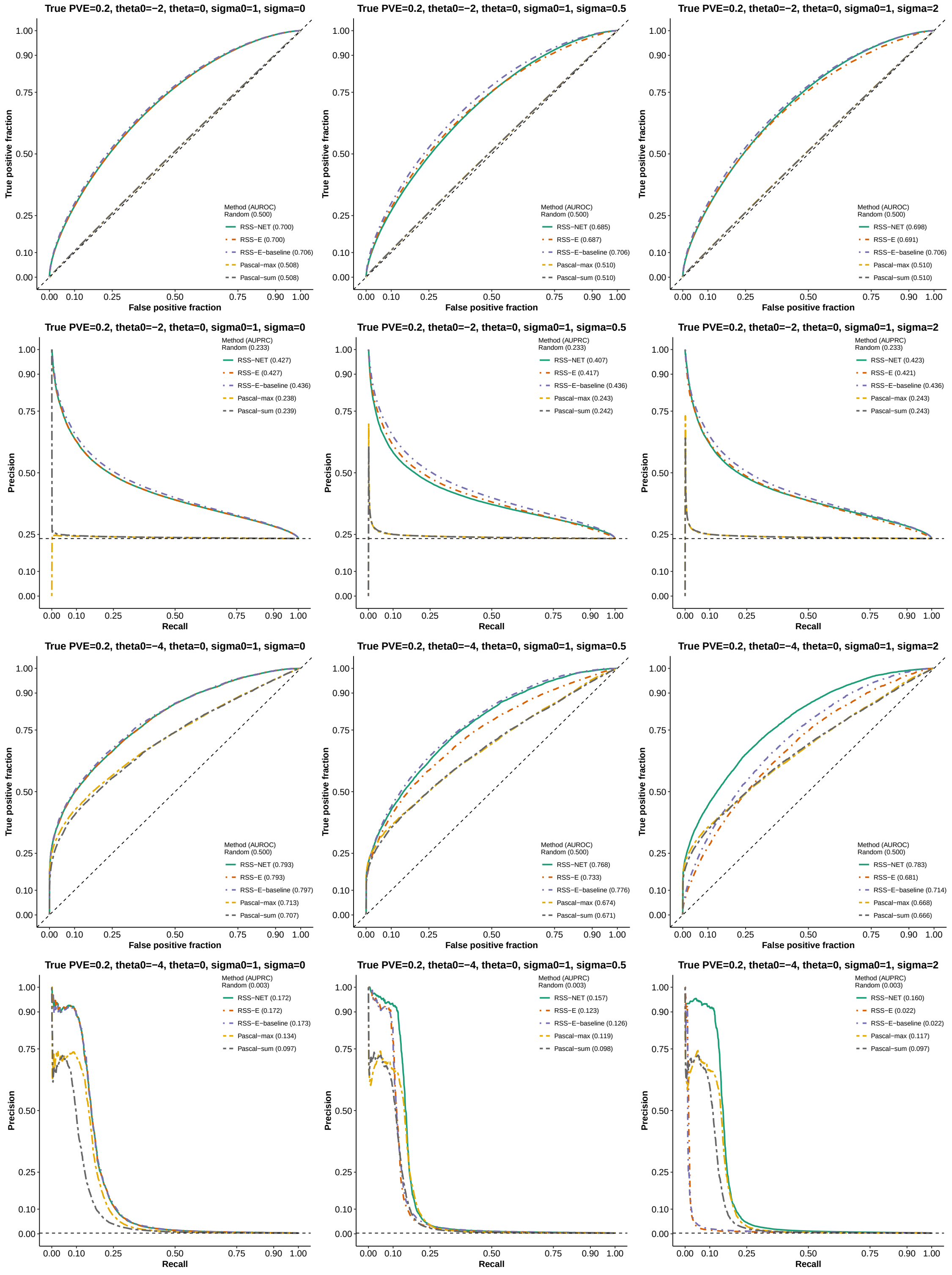

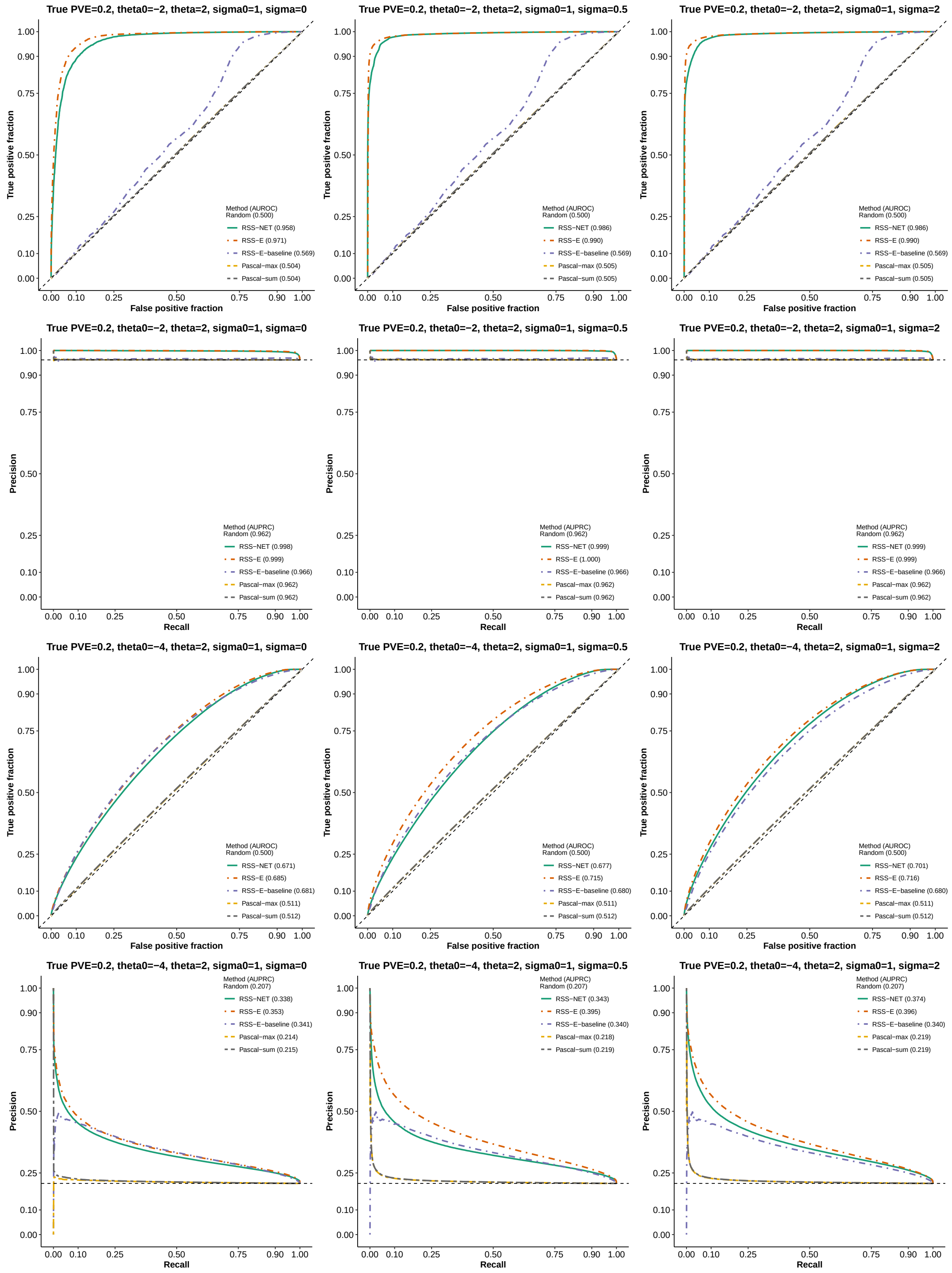
True PVE=0.2, theta0=−2, theta=2, sigma0=1, sigma=0
True positive fraction
False positive fraction
Method (AUROC)
Random (0.500)
RSS−NET (0.958)
RSS−E (0.971)
RSS−E−baseline (0.569)
Pascal−max (0.504)
Pascal−sum (0.504)
True PVE=0.2, theta0=−2, theta=2, sigma0=1, sigma=0.5
Method (AUROC)
Random (0.500)
RSS−NET (0.986)
RSS−E (0.990)
RSS−E−baseline (0.569)
Pascal−max (0.505)
Pascal−sum (0.505)
True PVE=0.2, theta0=−2, theta=2, sigma0=1, sigma=2
Method (AUROC)
Random (0.500)
RSS−NET (0.986)
RSS−E (0.990)
RSS−E−baseline (0.569)
Pascal−max (0.505)
Pascal−sum (0.505)
True PVE=0.2, theta0=−2, theta=2, sigma0=1, sigma=0
Precision
Recall
Method (AUPRC)
Random (0.962)
RSS−NET (0.998)
RSS−E (0.999)
RSS−E−baseline (0.966)
Pascal−max (0.962)
Pascal−sum (0.962)
True PVE=0.2, theta0=−2, theta=2, sigma0=1, sigma=0.5
Method (AUPRC)
Random (0.962)
RSS−NET (0.999)
RSS−E (1.000)
RSS−E−baseline (0.966)
Pascal−max (0.962)
Pascal−sum (0.962)
True PVE=0.2, theta0=−2, theta=2, sigma0=1, sigma=2
Method (AUPRC)
Random (0.962)
RSS−NET (0.999)
RSS−E (0.999)
RSS−E−baseline (0.966)
Pascal−max (0.962)
Pascal−sum (0.962)
True PVE=0.2, theta0=−4, theta=2, sigma0=1, sigma=0
Method (AUROC)
Random (0.500)
RSS−NET (0.671)
RSS−E (0.685)
RSS−E−baseline (0.681)
Pascal−max (0.511)
Pascal−sum (0.512)
True PVE=0.2, theta0=−4, theta=2, sigma0=1, sigma=0.5
Method (AUROC)
Random (0.500)
RSS−NET (0.677)
RSS−E (0.715)
RSS−E−baseline (0.680)
Pascal−max (0.511)
Pascal−sum (0.512)
True PVE=0.2, theta0=−4, theta=2, sigma0=1, sigma=2
Method (AUROC)
Random (0.500)
RSS−NET (0.701)
RSS−E (0.716)
RSS−E−baseline (0.680)
Pascal−max (0.511)
Pascal−sum (0.512)
True PVE=0.2, theta0=−4, theta=2, sigma0=1, sigma=0
Method (AUPRC)
Random (0.207)
RSS−NET (0.338)
RSS−E (0.353)
RSS−E−baseline (0.341)
Pascal−max (0.214)
Pascal−sum (0.215)
True PVE=0.2, theta0=−4, theta=2, sigma0=1, sigma=0.5
Method (AUPRC)
Random (0.207)
RSS−NET (0.343)
RSS−E (0.395)
RSS−E−baseline (0.340)
Pascal−max (0.218)
Pascal−sum (0.219)
True PVE=0.2, theta0=−4, theta=2, sigma0=1, sigma=2
Method (AUPRC)
Random (0.207)
RSS−NET (0.374)
RSS−E (0.396)
RSS−E−baseline (0.340)
Pascal−max (0.219)
Pascal−sum (0.219)

#### Supplementary Figure 9

**RSS-NET simulations based on 1 million common SNPs.** This part is the same as **Supplementary Figure 1**, except that here we use the haplotype data of 503 European-ancestry individuals on 1,030,397 common SNPs (MAF $\geq 1\%$) from 1000 Genomes Project Consortium (2015) to specify the genotype matrix $X$. Results below (**a**: timing; **b**: network enrichment; **c**: gene-level association) correspond to simulations with true hyper-parameters of positive datasets being $\theta_0 = -4$, $\theta = 2$, $\sigma_0 = 1$, $\sigma = 2$ and PVE = 0.6.

As shown in Panel **a**, the computation time of RSS-NET increases as the number of genome-wide SNPs analyzed increases. On average, when the number of SNPs increases from 348,965 to 1,030,397, the total computation time per dataset is four times longer (one-sided Wilcoxon $P = 8.02 \times 10^{-132}$). Due to the large-scale simulations conducted in this study, we mostly use 348,965 genome-wide SNPs in order to reduce computation.

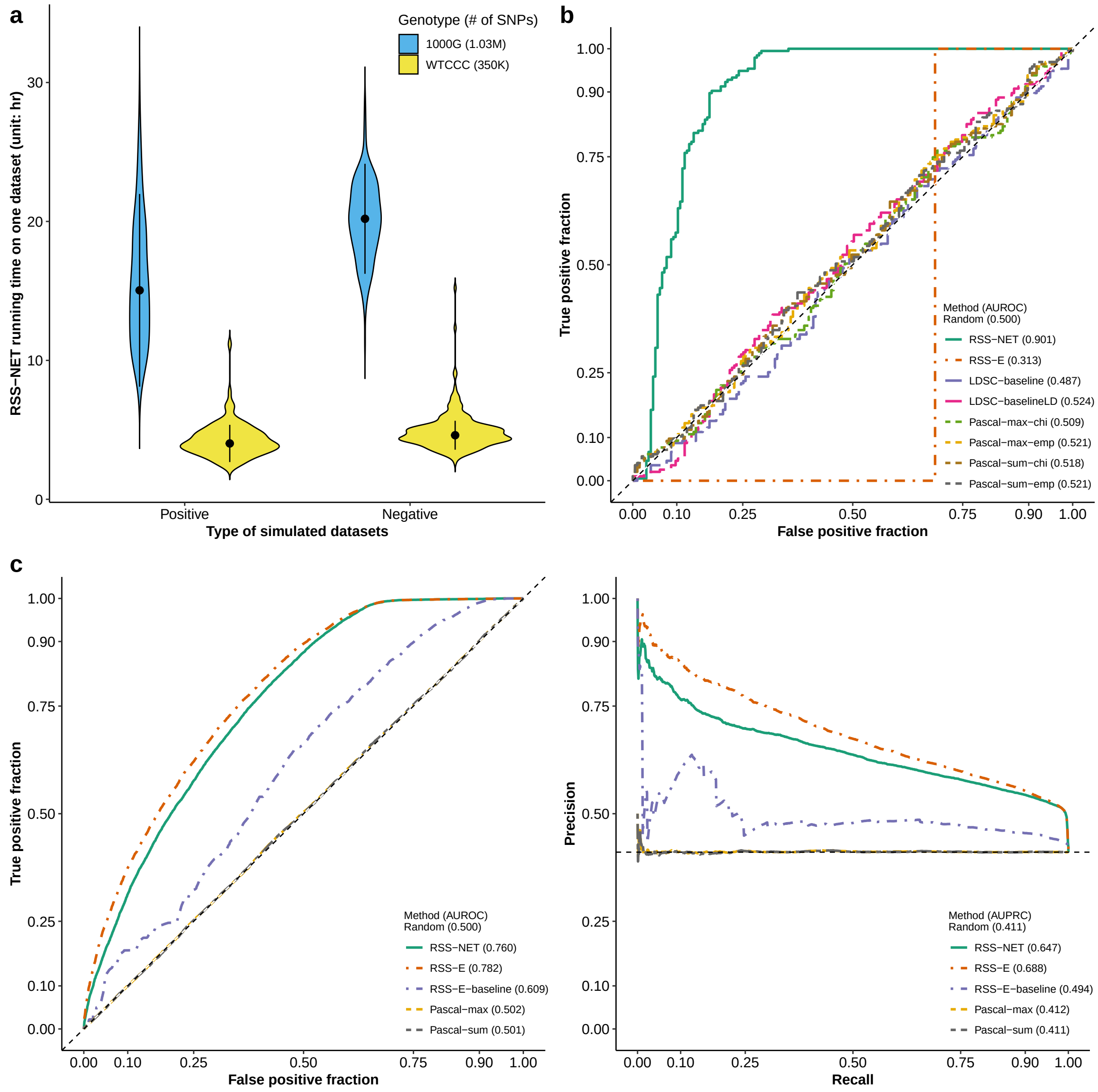

#### Supplementary Figure 10

**Robustness of RSS-NET under noisy networks.** This part aims to assess the robustness of RSS-NET to model mis-specification where the GWAS summary data are simulated from a real target network and the enrichment testing is performed on a noisy version of this real target. Details of this part are almost identical to those in **Supplementary Figure 1**. Here we only highlight the differences.

The real target network used to simulate GWAS summary data is the B cell regulatory network in **Supplementary Figure 1**. We create a noisy network for this target by randomly subsetting a given fraction of its TF-TG edges and the associated REs. Here the fraction of TF-TG edges to subset are $\{1\%, 5\%, 10\%, 50\%\}$. For each pair of negative and positive GWAS datasets, we simulate a new noisy network and use it in RSS-NET.

The GWAS summary data (both negative and positive) used here are the same as those used in **Supplementary Figure 1**, with true $\theta_0 = -4$, $\theta = 2$, $\sigma_0 = 1$, $\sigma = 2$ and PVE $= 0.2$. Unlike **Supplementary Figure 1**, here we do not test the enrichment of B cell network (i.e. the real target network), but test its random subsets instead.

Panels **a-c** Comparison of network enrichment results between using the real target network and using its noisy versions in RSS-NET.

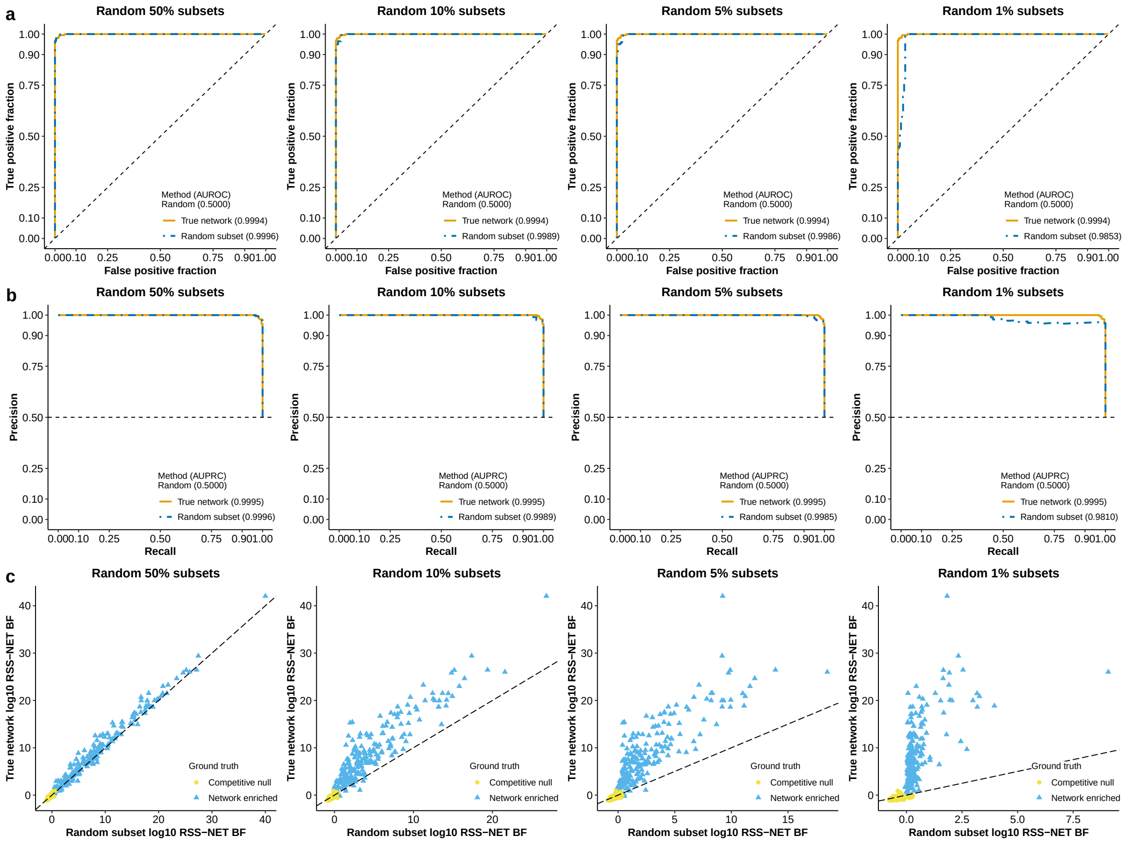

Panels **d-e** Comparison of gene-level association results between using the real target network and using its noisy versions in RSS-NET. Note that the AUROC based on true network shown below is slightly different from the AUROC shown in **Supplementary Figure 7**, although they are generated from the same simulated datasets. This is because AUROC shown in **Supplementary Figure 7** is evaluated based on 16,954 autosomal protein-coding genes that are available in Pascal-required database, and AUROC shown below is evaluated based on all available autosomal protein-coding genes.

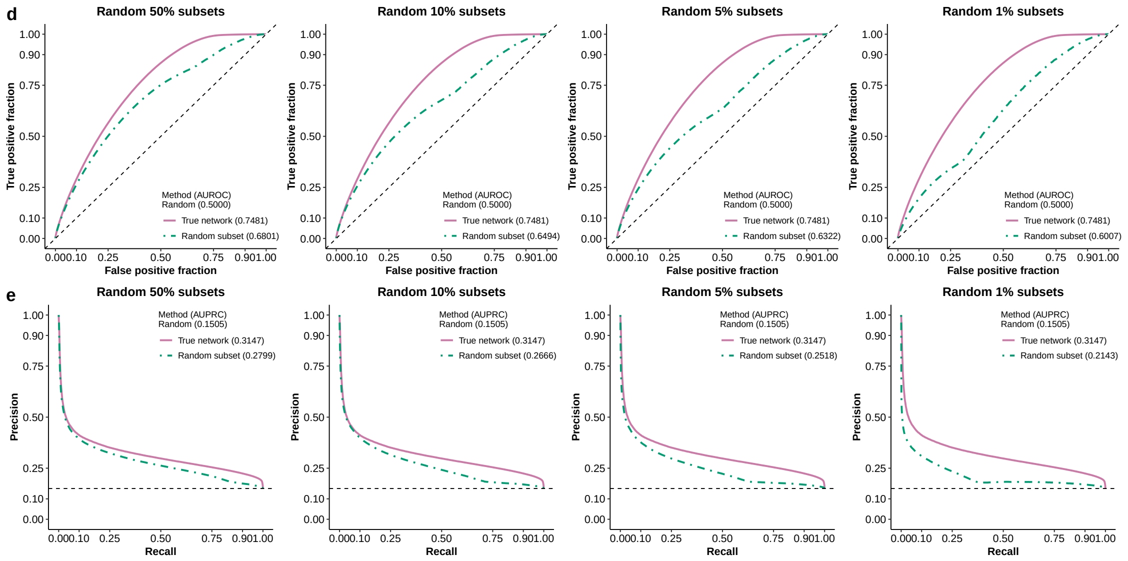


Panels **f-i** Descriptive statistics of noisy networks under different subsetting fractions. Each box plot below is based on 200 noisy networks.

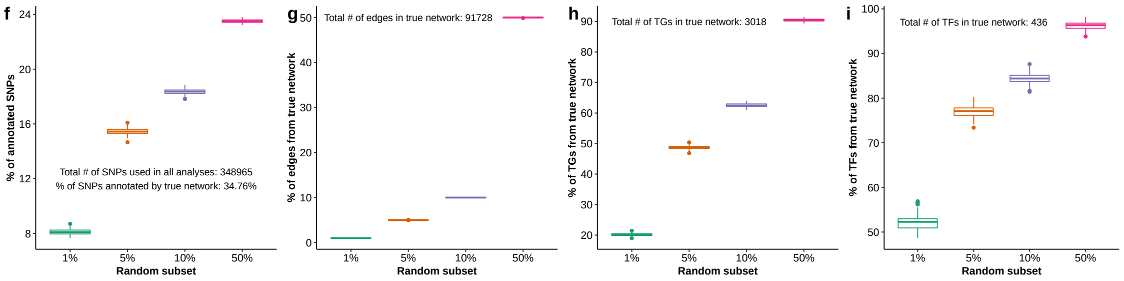

#### Supplementary Figure 11

**Analysis details and additional results for Figure 5(a).** Here we perform clustering analysis of 38 regulatory networks based on principal component analysis (PCA), $t$-distributed stochastic neighbor embedding (t-SNE), and hierarchical clustering. To perform PCA, we use the R built-in function `prcomp`. To perform t-SNE, we use the R package `Rtsne`. To perform hierarchical clustering, we use the R built-in function `hclust`, with distance = 1 − Pearson correlation, and average method. **Figure 5(a)** corresponds to the t-SNE plot in Panel **e**.

Panel **a** Here the input data matrix has 1,289,786 rows (SNPs) and 38 columns (networks), where the $(i, j)$-th entry is 1 if SNP $i$ is within 100 kb of any associated RE or transcribed region of any member gene of network $j$, and it is 0 otherwise.

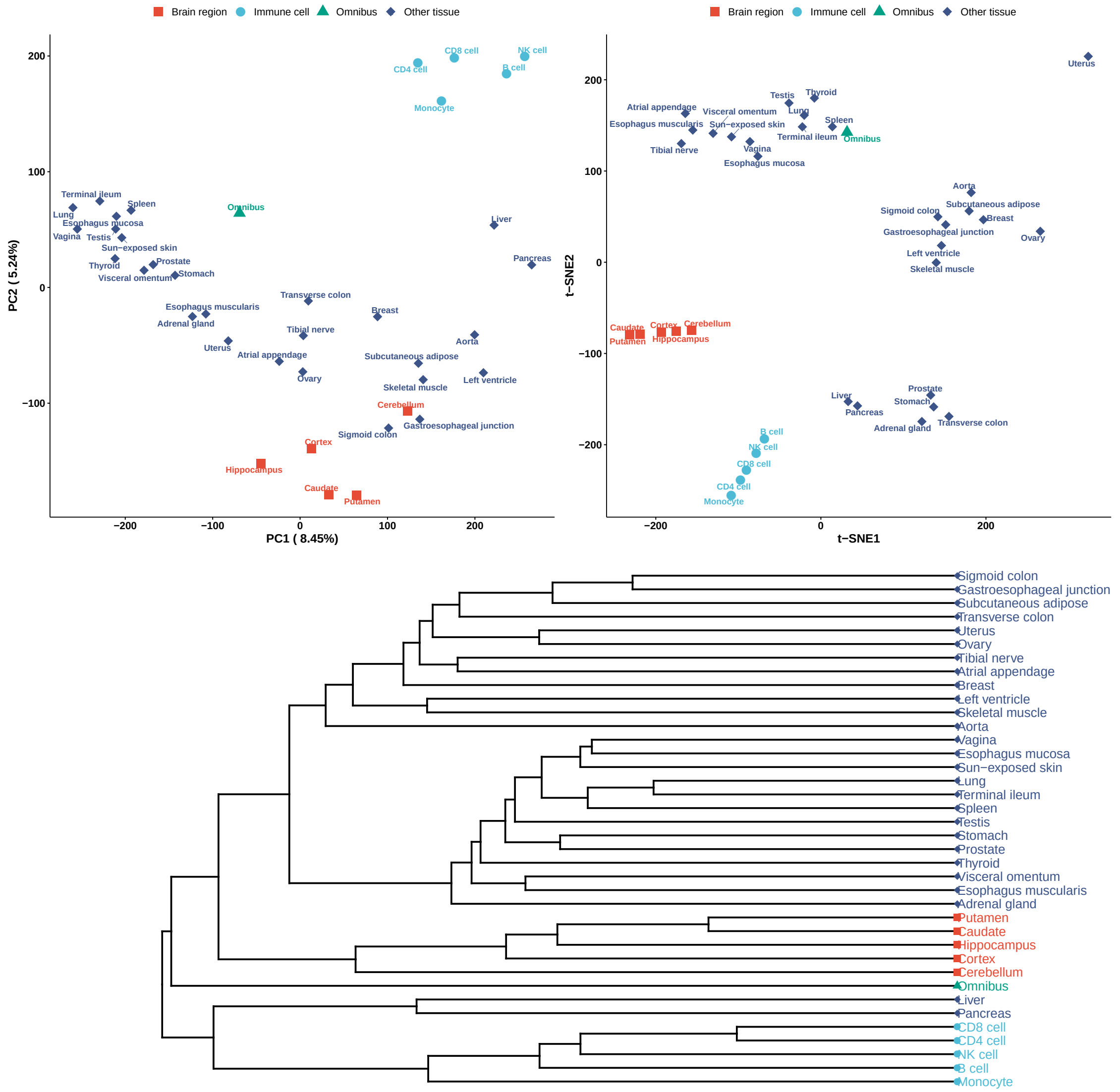

Panel **b** Here the input data matrix has 605 rows (TFs) and 38 columns (networks), where the $(i, j)$-th entry is 1 if TF $i$ belongs to network $j$, and it is 0 otherwise.

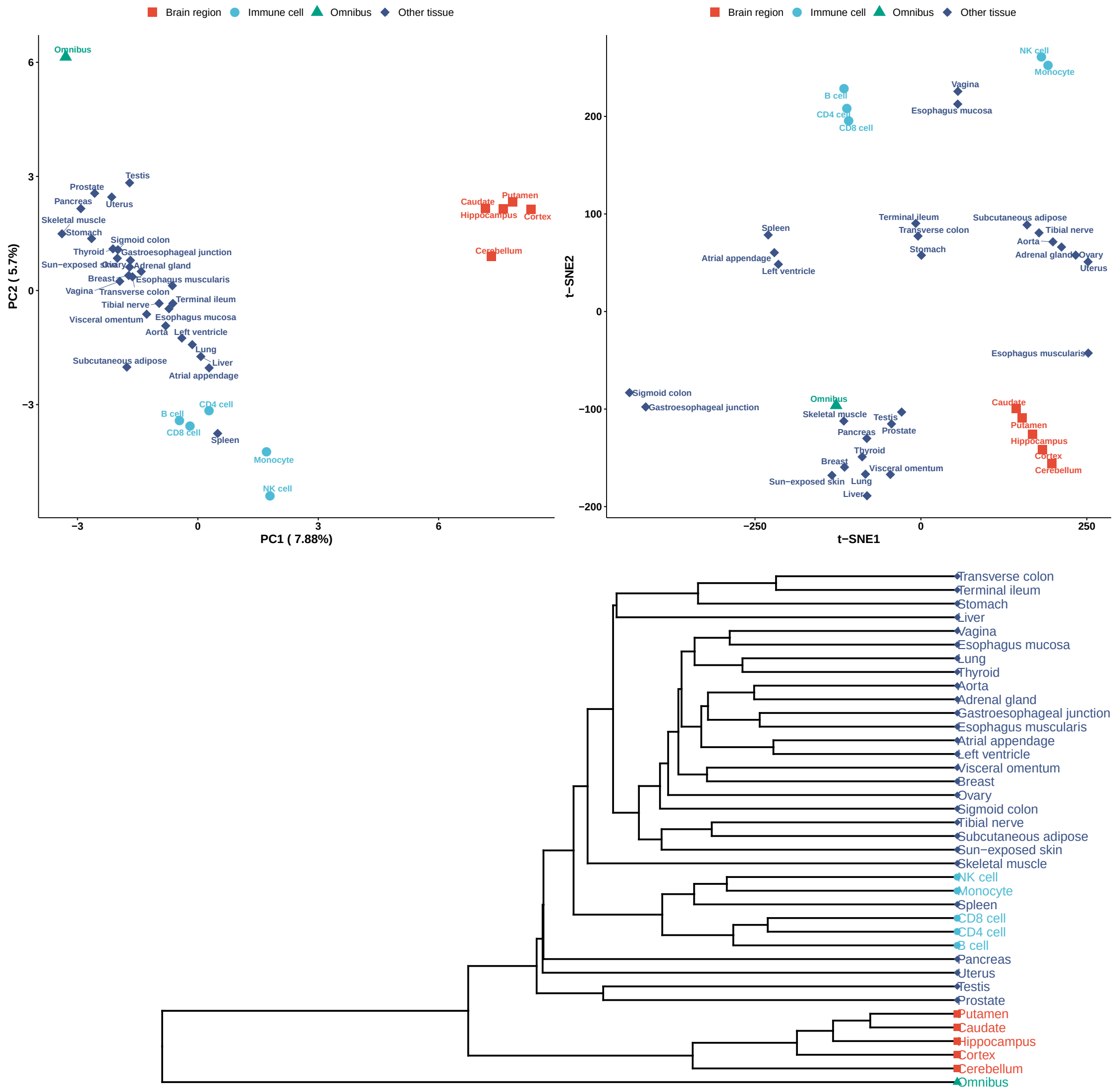

Panel **c** Here the input data matrix has 12,963 rows (TGs) and 38 columns (networks), where the $(i, j)$-th entry is 1 if TG $i$ belongs to network $j$, and it is 0 otherwise.

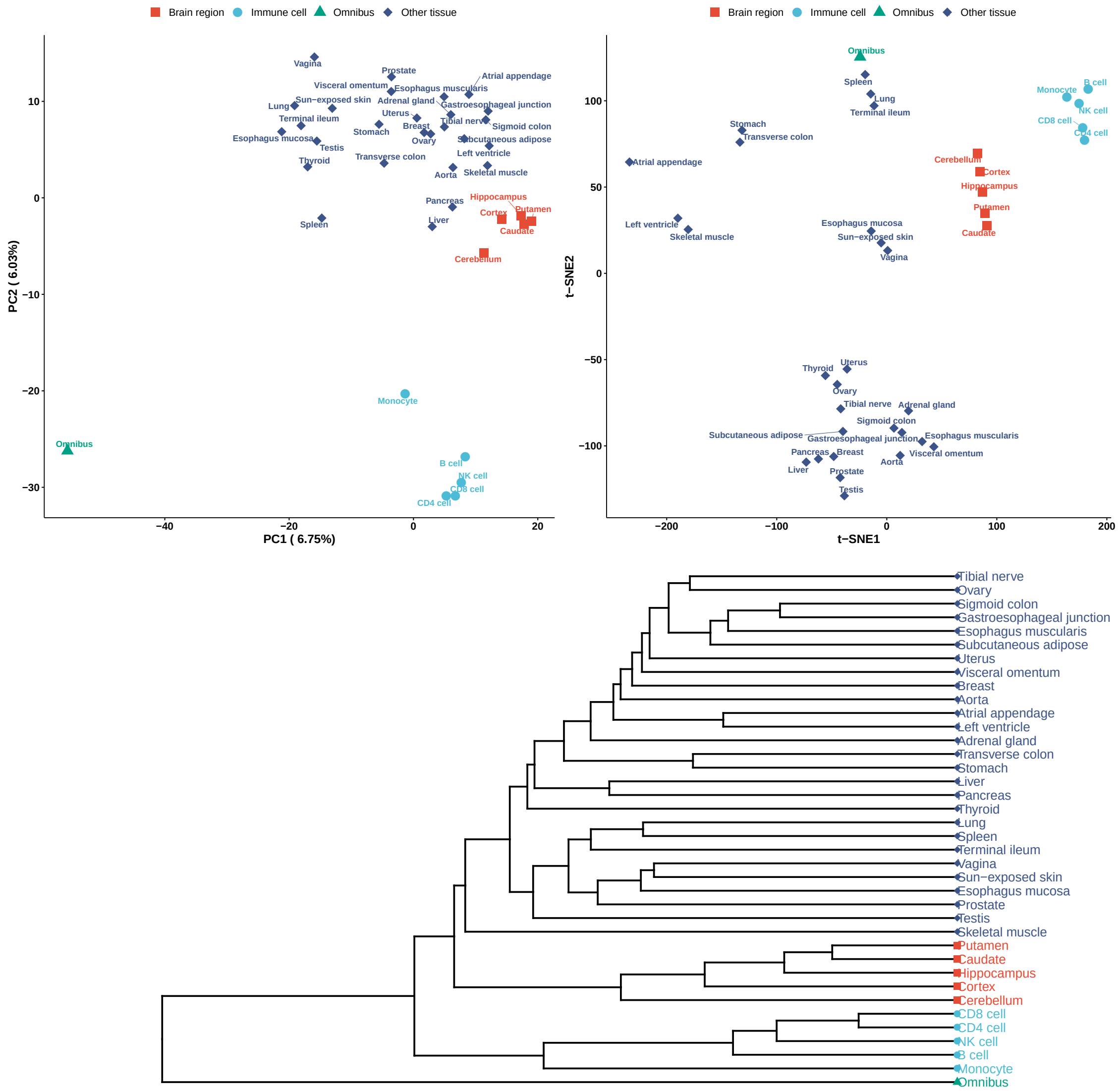

Panel **d** Here the input data matrix has 880,777 rows (TF-TG edges) and 38 columns (networks), where the $(i, j)$-th entry is the weight of TF-TG edge $i$ in network $j$, and it is 0 if TF-TG edge $i$ is not available in network $j$.

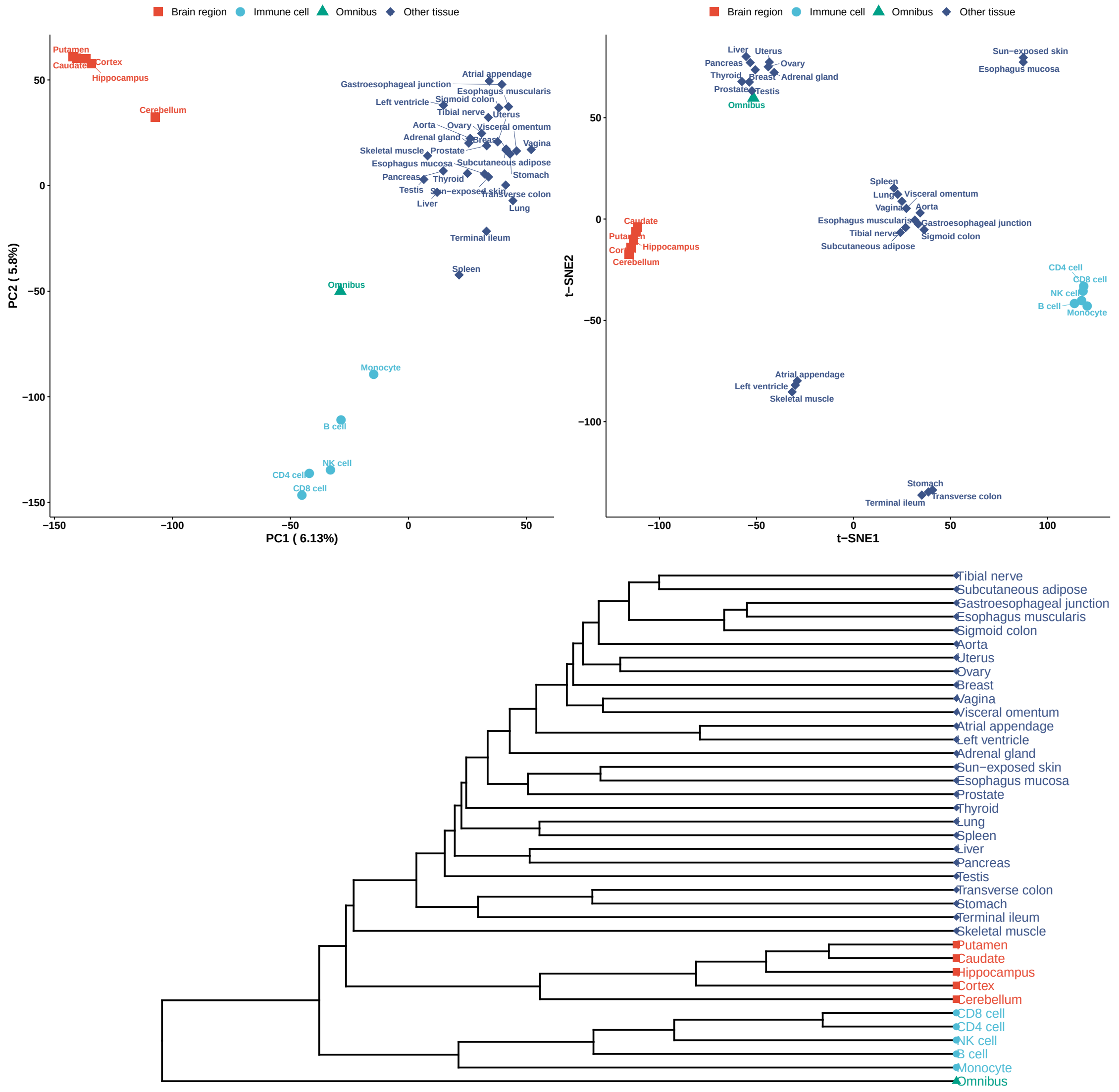

Panel **e** Here the input data matrix is obtained by concatenating the previous 4 data matrices vertically, with a total of 2,184,131 rows and 38 columns (networks).

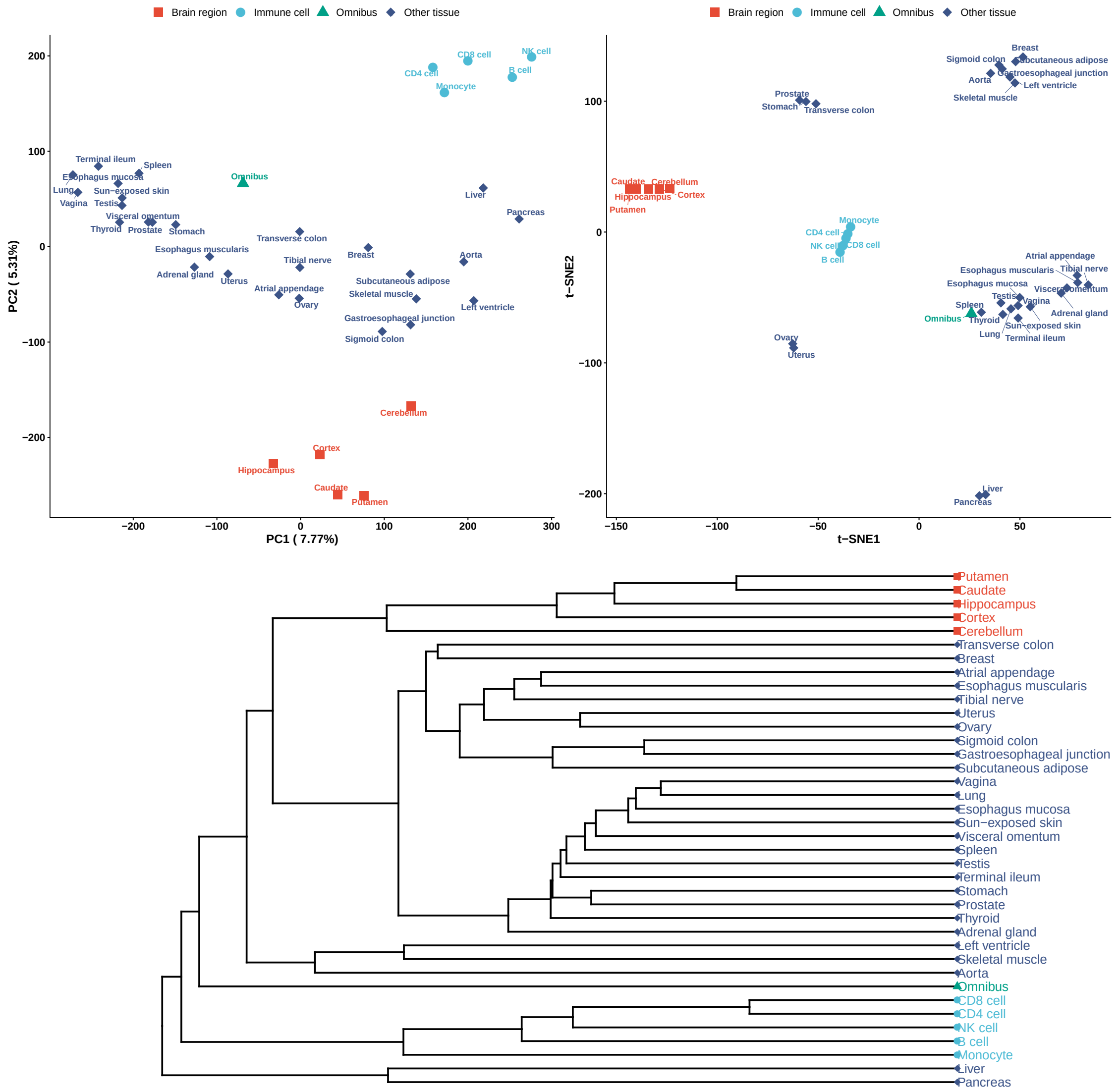

#### Supplementary Figure 12

**Additional results for Figure 5(b).** Each panel below shows the node and edge similarities between a given cell type- or tissue-specific PECA-based network (panel caption: cell type or tissue) and CAGE-based networks for 394 cell types and tissues (Marbach et al. 2016) inferred from independent CAGE data (Andersson et al. 2014; The FANTOM Consortium and the RIKEN PMI and CLST (DGT) 2014). In each panel, each point represents a context-specific CAGE-based network, where $x$- and $y$-axis values indicate its node and edge set similarity with the given PECA network respectively. For both node and edge sets, the similarity is measured by Jaccard index (the size of the intersection divided by the size of the union).

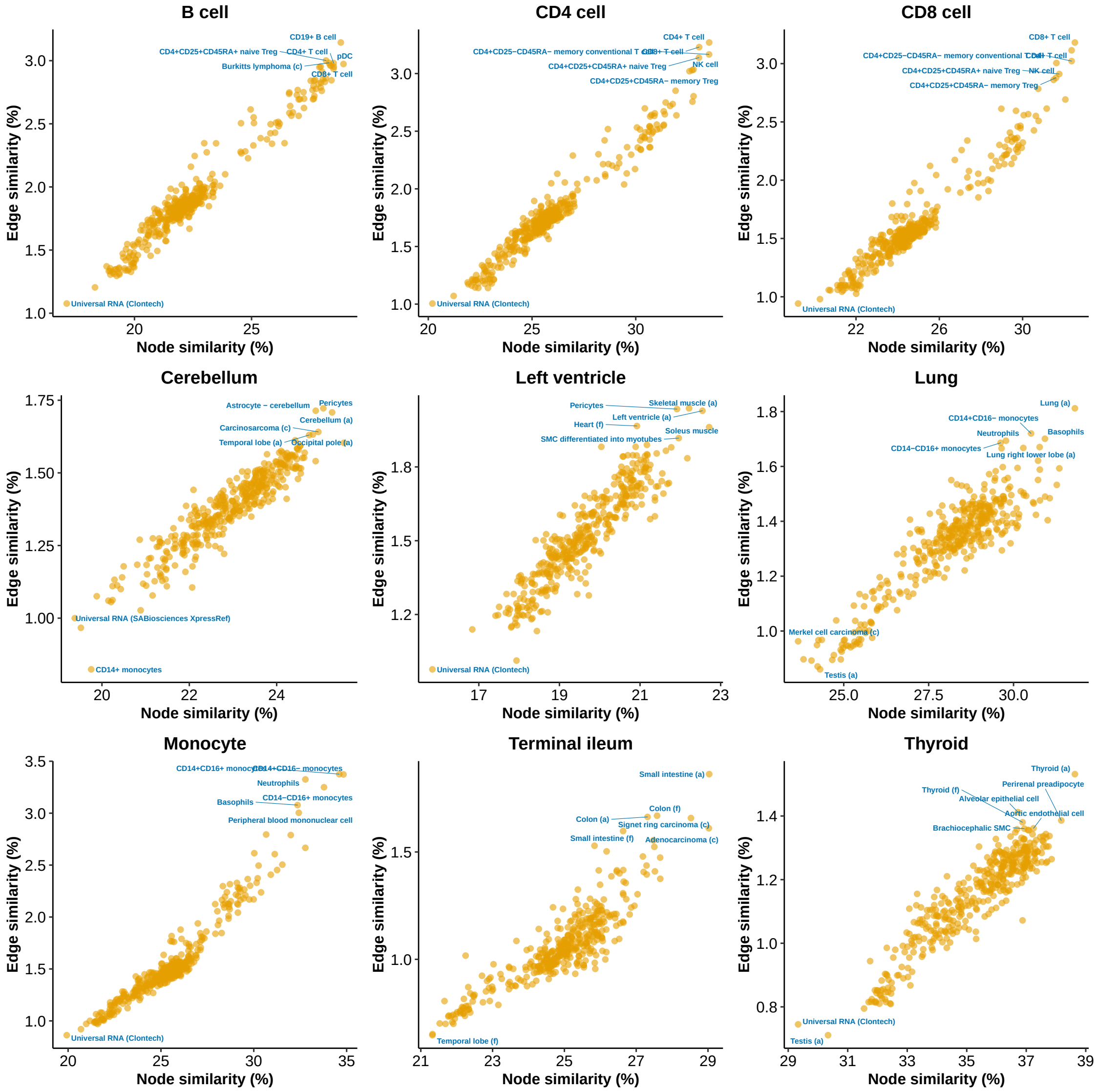

#### Supplementary Figure 13

**RSS-NET results with zero TF-TG edge weights.** Here we repeat RSS-NET analyses of 11 GWAS traits on 38 TF-TG regulatory networks by setting all TF-TG edge weights zero (i.e., $v_{gt}=0$ in main text Equation 5). Each panel below corresponds to one of 11 GWAS traits that pass the near-gene enrichment control for all 38 networks in our original analyses using actual TF-TG edge weights (**Supplementary Table 3**). We remove myocardial infarction since it is a subtype of coronary artery disease. In each panel, each point represents one of the 38 regulatory networks. The $x$- and $y$-axis values of each point are log 10 BFs ($M_1$ against $M_0$) based on zero and actual edge weights of a network respectively. All diagonal lines have slope 1 and intercept 0. Numerical values are available at https://suwonglab.github.io/rss-net/results.html.

For 10 out 11 traits, BFs based on actual edge weights are unanimously larger than BFs based on zero edge weights across all 38 networks (i.e., all dots above the diagonal line). The only exception is low-density lipoprotein cholesterol, for which BFs based on actual edge weights and BFs based on zero edge weights are not significantly different from zero across 38 networks (Wilcoxon $P=0.91$).

Of note, setting TF-TG edge weights zero is different from setting the edge enrichment parameter $\sigma^2$ zero (see main text Equations 2 and 5-6). Setting edge weights zero induces the following effect size distribution: $\beta_j \sim \pi_j \cdot \mathcal{N}(\sum_{g\in\mathbf{G}_j} c_{jg}\cdot\gamma_{jg},\ \sigma_0^2)+(1-\pi_j)\cdot\delta_0$, $\gamma_{jg}\sim\mathcal{N}(0,\ \sigma^2)$. where $\{\mathbf{G}_j, c_{jg}, \gamma_{jg}\}$ are defined in main text Equations 5-6. Setting $\sigma^2=0$ induces a different effect size distribution: $\beta_j\sim\pi_j\cdot\mathcal{N}(0,\ \sigma_0^2)+(1-\pi_j)\cdot\delta_0$.

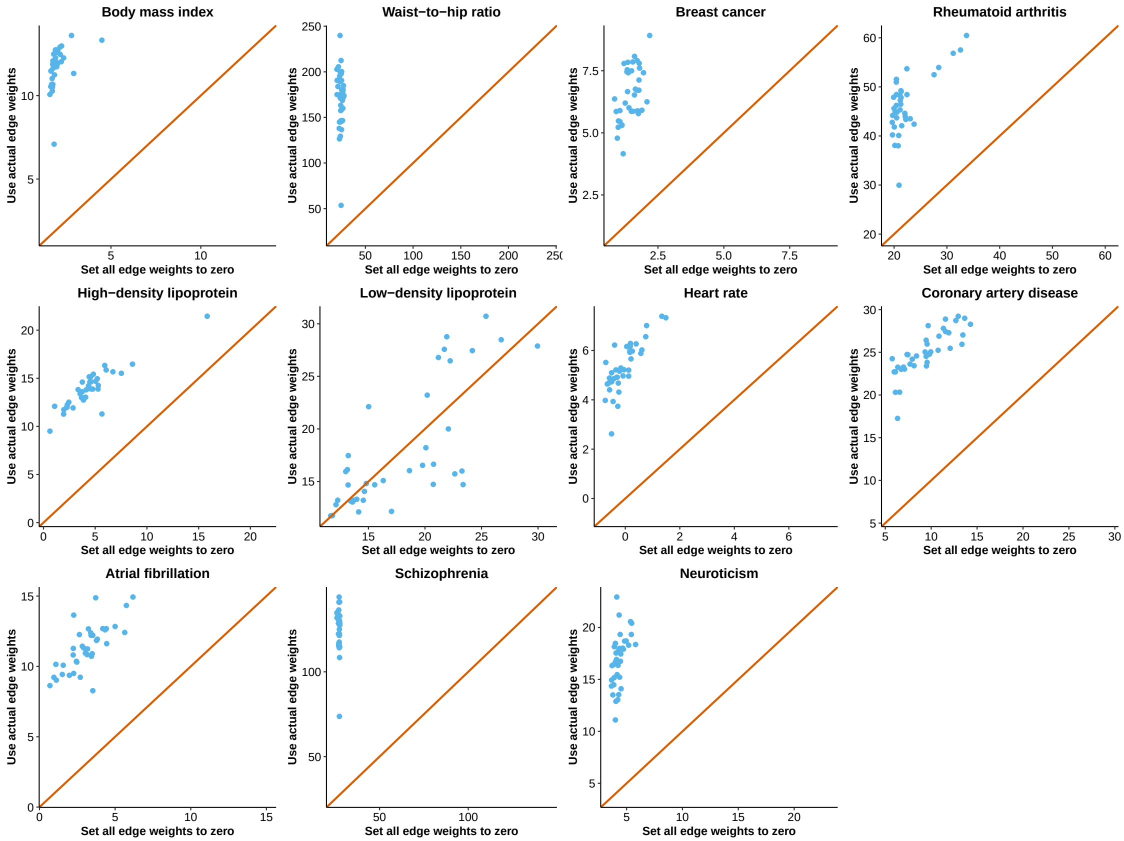

#### Supplementary Figure 14

**Additional results for Figure 5(d).** Each panel below shows the comparison of PECA- and CAGE-based network enrichments on the same cell types or tissues (labels) and the same GWAS data (caption). In each panel, each point represents a context (cell type or tissue), where *x*- and *y*-axis show the RSS-NET log 10 enrichment BFs for the CAGE- and PECA-based networks in this context respectively. Given a context, the analyzed CAGE- and PECA-based networks have the same TF-TG edge counts, and their edge weights are on the same scale (see main text Methods section). The RSS-NET analyses of CAGE- and PECA-based networks use the same hyper-parameter grids (**Supplementary Table 19**). All dashed lines have slope 1 and intercept 0. Numerical values are available at https://suwonglab.github.io/rss-net/results.html.

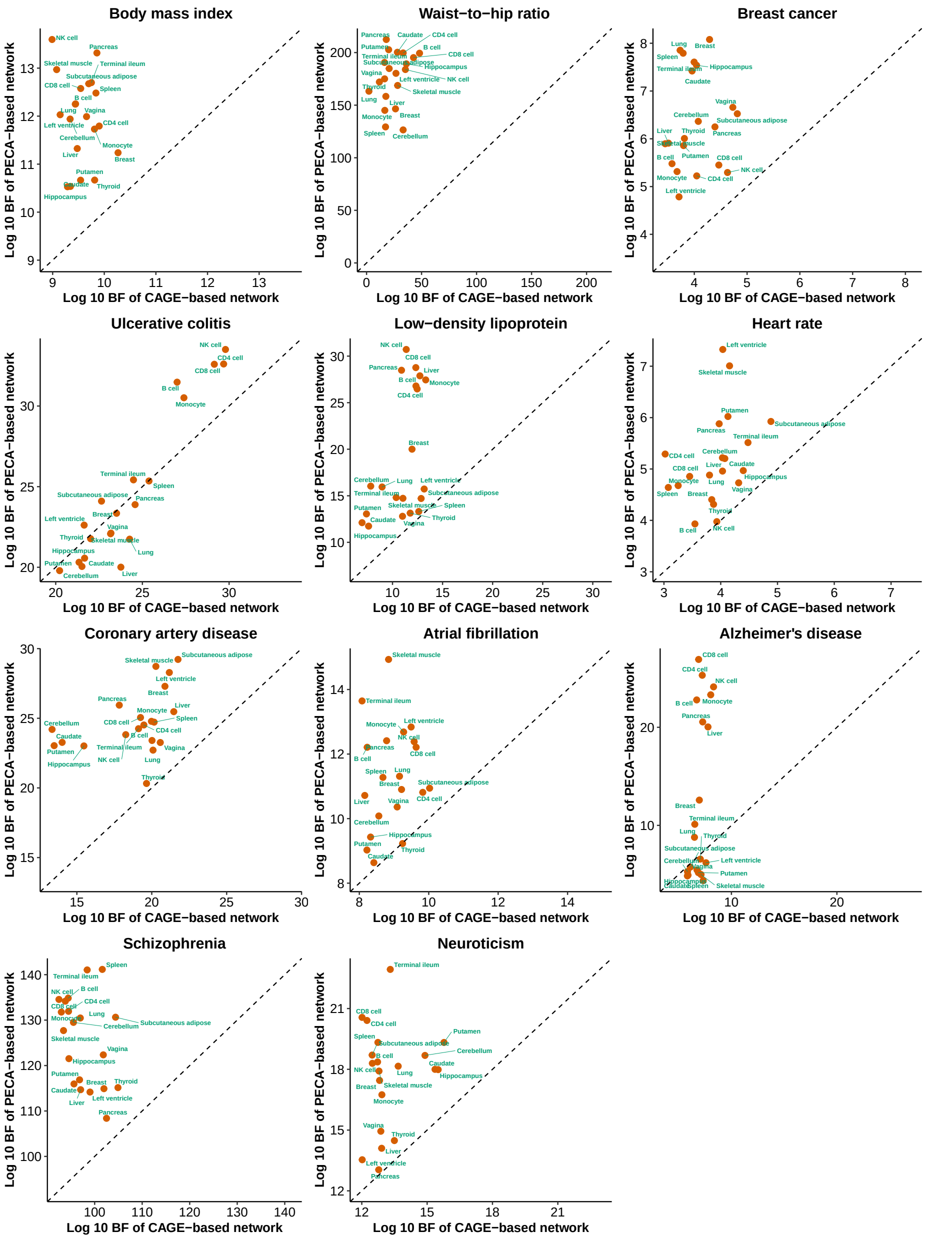

#### Supplementary Figure 15

**Overlap between RSS-NET prioritized genes and genes from external databases.** In each panel, each point represents a pair of GWAS trait and knockout mouse phenotype (left) or human Mendelian disorder (right). For each point, the log odds ratio (OR) is produced by running R built-in function `fisher.test` on the following 2 by 2 table:

| | Query genes | Other genes |
|---|---|---|
| Genes with $P_1^{\mathrm{bma}} \geq 0.9$ | | |
| Genes with $P_1^{\mathrm{bma}} < 0.9$ | | |

For the left panel, "query genes" correspond to genes that are implicated in one of 27 categories of knockout mouse phenotypes (Bult et al. 2019), and "other genes" correspond to genes that are not implicated in any mouse phenotype. For the right panel, "query genes" correspond to genes that are implicated in one of 19 categories of Mendelian disorders (Freund et al. 2018; Amberger et al. 2019), and "other genes" correspond to genes that are not implicated in any Mendelian disorder. For all points, the log ORs shown on the $x$-axis are estimated from all genes, and the log ORs shown on the $y$-axis are estimated after excluding genes that are implicated in GWAS of the same trait at the time of analysis (**Supplementary Table 10**). The OR values and associated $P$-values are available in **Supplementary Tables 12-13**. The diagonal lines have slope 1 and intercept 0.

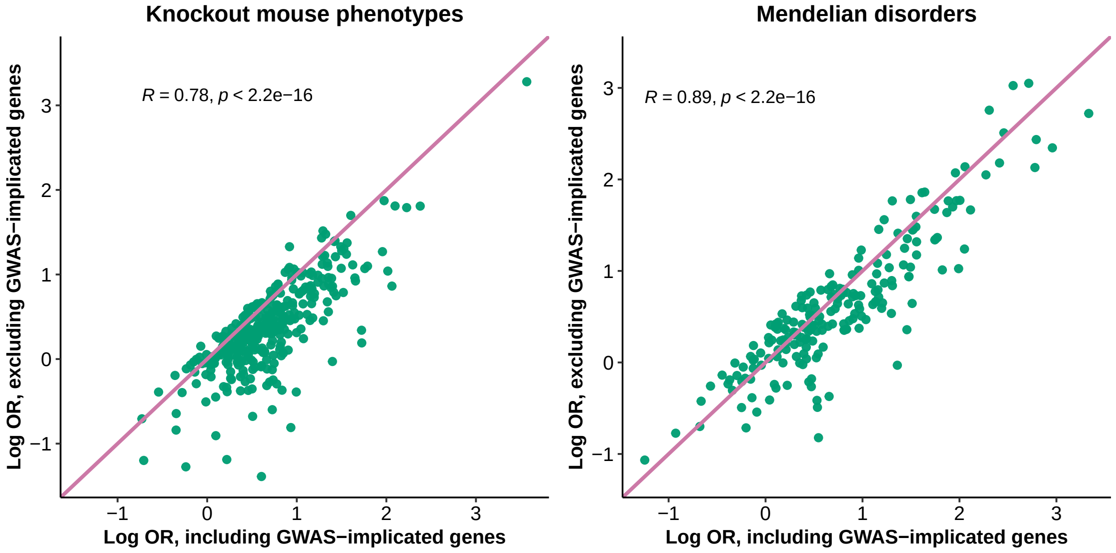

#### Supplementary Figure 16

**Posterior estimation of $\{\sigma,\theta\}$ in simulations.** Similar to our calculation of $P_1$ (20), we estimate the posterior means of $\{\sigma,\theta\}$ as:

$$E(\sigma \mid \mathbf{D}) \approx \sum_{s=1}^{n_1} \sigma^{(s)} \cdot \omega^{\star}(\theta_0^{(s)},\theta^{(s)},\sigma_0^{(s)},\sigma^{(s)}),$$

$$E(\theta \mid \mathbf{D}) \approx \sum_{s=1}^{n_1} \theta^{(s)} \cdot \omega^{\star}(\theta_0^{(s)},\theta^{(s)},\sigma_0^{(s)},\sigma^{(s)}),$$

where $\omega^{\star}(\theta_0,\theta,\sigma_0,\sigma)$ is a discrete approximation to the joint posterior of hyper-parameters $\{\theta_0,\theta,\sigma_0,\sigma\}$; see (16). The $x$-axis indicates the simulation scenarios (true hyper-parameter values of the positive datasets and the target network); see **Supplementary Figures 1-2**. Each box contains results of 200 independent positive datasets for a given scenario. Horizontal dashed lines indicate true values of $\{\sigma,\theta\}$.

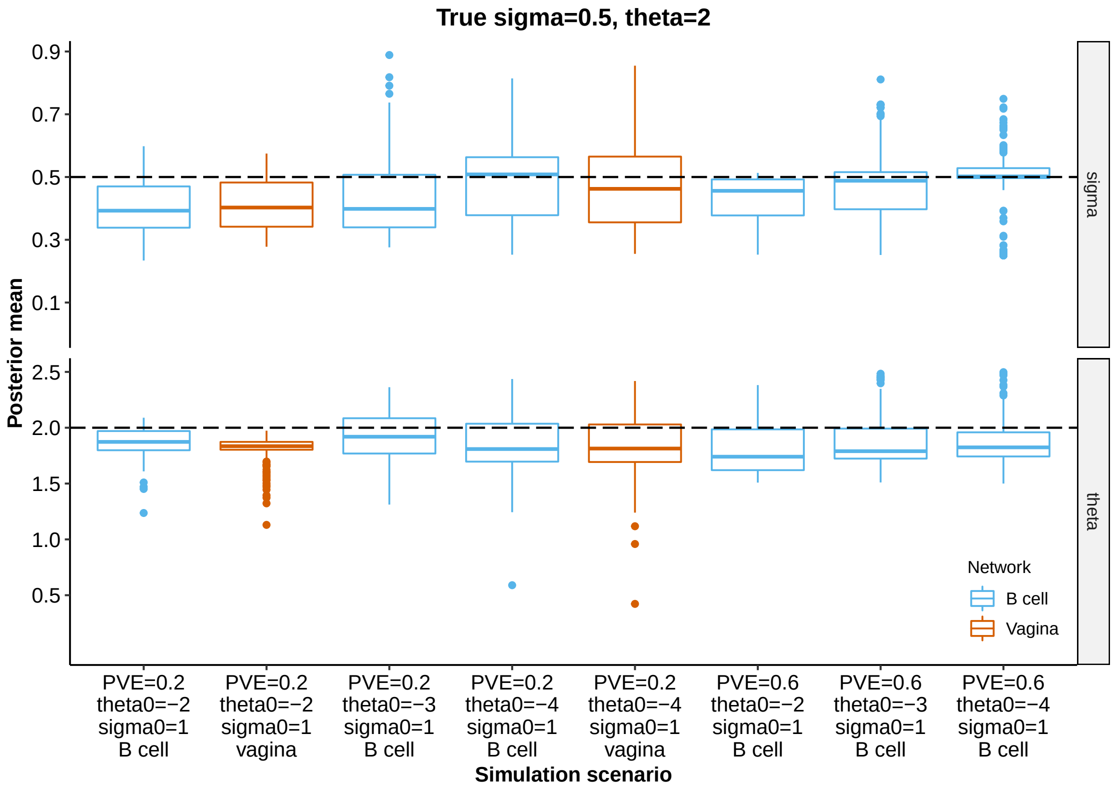

#### Supplementary Figure 17

**Robustness of RSS-NET to $\sigma$ specification in simulations.** Here we repeat RSS-NET analyses of simulated datasets with a fixed value of $\sigma \in \{0.25, 0.5, 0.75, 1\}$. In our original analyses we use a model averaging approach (18) to account for the uncertainty of hyper-parameters.

Panel **a** The simulated negative and positive GWAS summary data are the same as those from **Supplementary Figures 1-2**. The caption of each plot shows the true hyper-parameter values of positive datasets and the target network for enrichment testing.

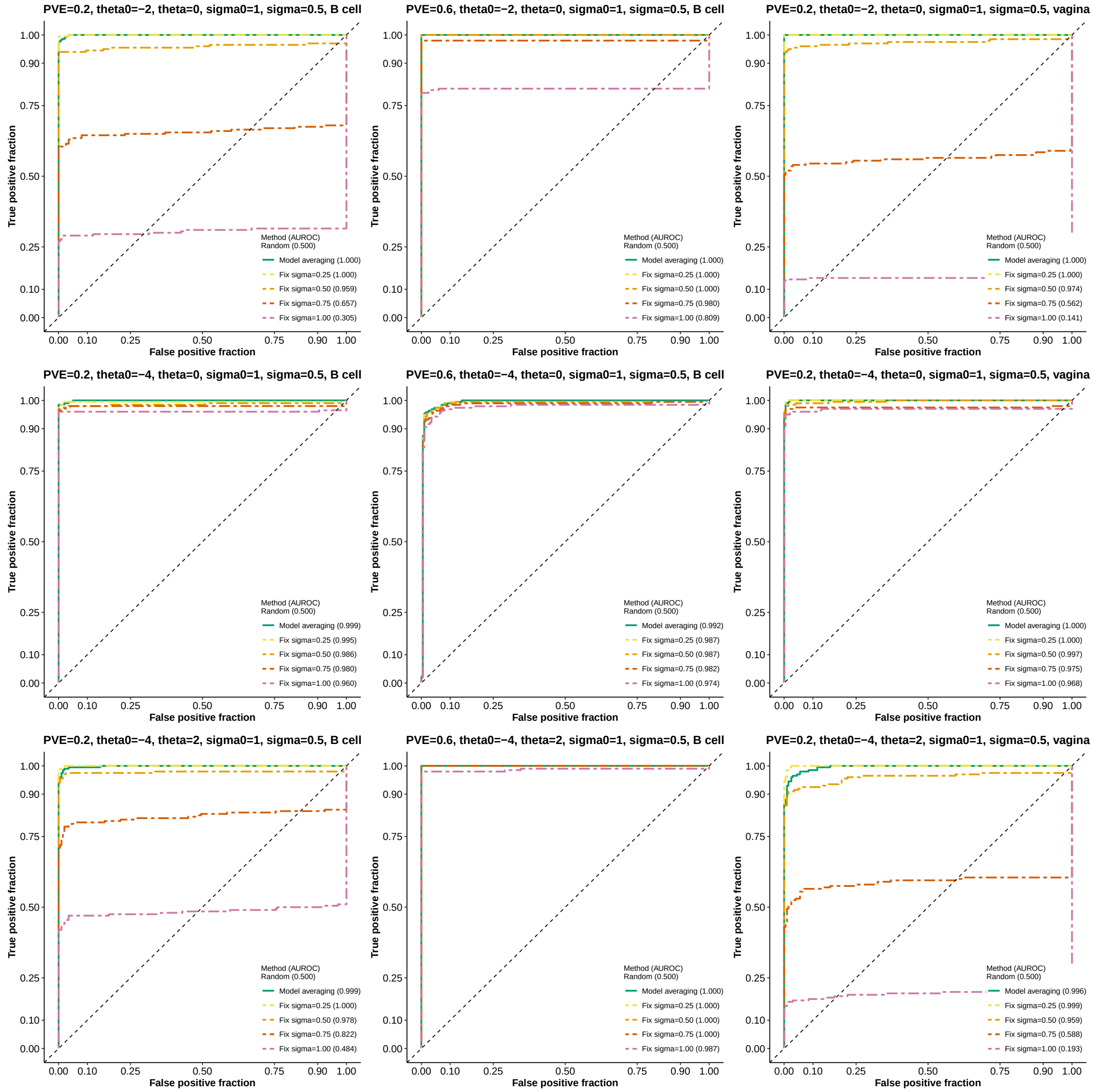

Panel **b** Each plot corresponds to a positive simulation scenario with 200 independent datasets, and its caption indicates the true hyper-parameter values and the target network. In each plot, each point represent a simulated positive dataset. The $y$-axis value of each point is the log 10 enrichment BF when the value of $\sigma$ is fixed as 0.5 (ground truth). The $x$-axis value of each point is the log 10 BF under other $\sigma$ specifications (differentiated by point shape and color). The dashed lines have slope 1 and intercept 0. The number "($a$ / $200 - a$)" in each legend indicates that BFs based on $\sigma = 0.5$ are higher than BFs based on other $\sigma$ specifications in $a$ simulated datasets for this scenario.

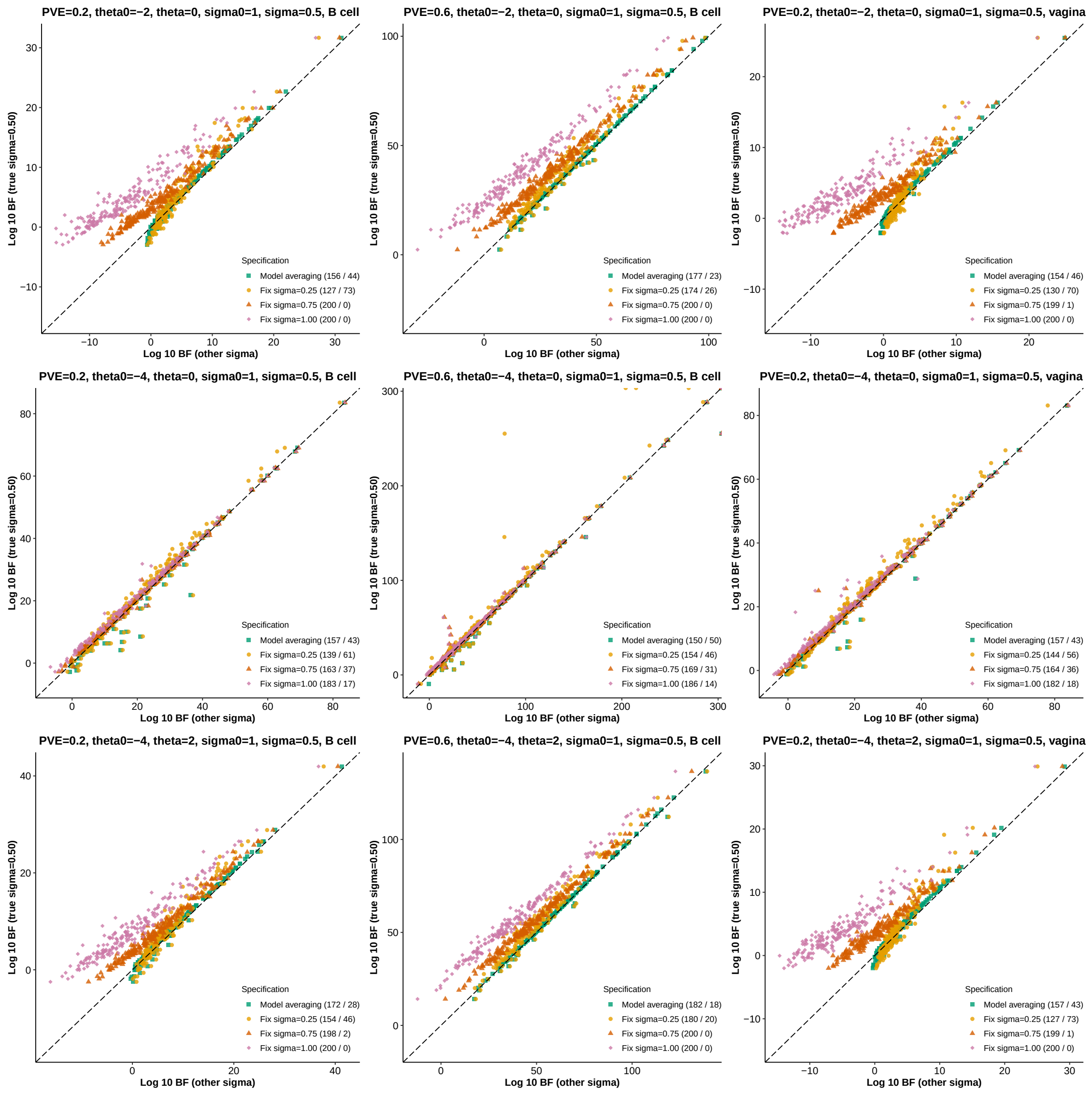


**Concluding Remark:** Apart from hyper-parameters, RSS-NET uses approximations in the likelihood function (Zhu and Stephens 2017) and the variational inference (Zhu and Stephens 2018). Hence, setting $\sigma$ as ground truth does not guarantee the optimal outcome in some simulated datasets. Instead we recommend the model averaging approach (18) as used in our main analyses.

#### Supplementary Figure 18

**Robustness of RSS-NET to hyper-parameter grid choice in real data.** For three GWAS datasets, we run RSS-NET on 38 networks with two different hyper-parameter grids (grid 1: coarse; grid 2: focused), and then compare results of network enrichments (BF, top panels) and gene associations ($P_1^{\text{net}}$, bottom panels). In each enrichment comparison plot, each point represents a network. In each association comparison plot, each point represent a gene-network pair. All diagonal lines have slope 1 and intercept 0.

Panel **a** Alzheimer's disease (Lambert et al. 2013). In grid 1, $\theta \in (0 : 0.5 : 4)$. In grid 2, $\theta \in (0 : 0.25 : 1)$. For both grids, $\eta = 0.6$, $\rho \in (0 : 0.2 : 0.8)$ and $\theta_0 \in (-5.25 : 0.1 : -4.75)$. Panel **b** low-density lipoprotein (Teslovich et al. 2010). In grid 1, $\theta \in (0 : 0.5 : 3)$. In grid 2, $\theta \in (0 : 0.25 : 1)$. For both grids, $\eta = 0.3$, $\rho \in (0 : 0.2 : 0.8)$ and $\theta_0 \in (-3.75 : 0.05 : -3.5)$. Panel **c** inflammatory bowel disease (Liu et al. 2015). In grid 1, $\theta \in (0 : 0.5 : 3)$. In grid 2, $\theta \in (0 : 0.25 : 1)$. For both grids, $\eta = 0.3$, $\rho \in (0 : 0.2 : 0.8)$ and $\theta_0 \in (-3 : 0.05 : -2.8)$.

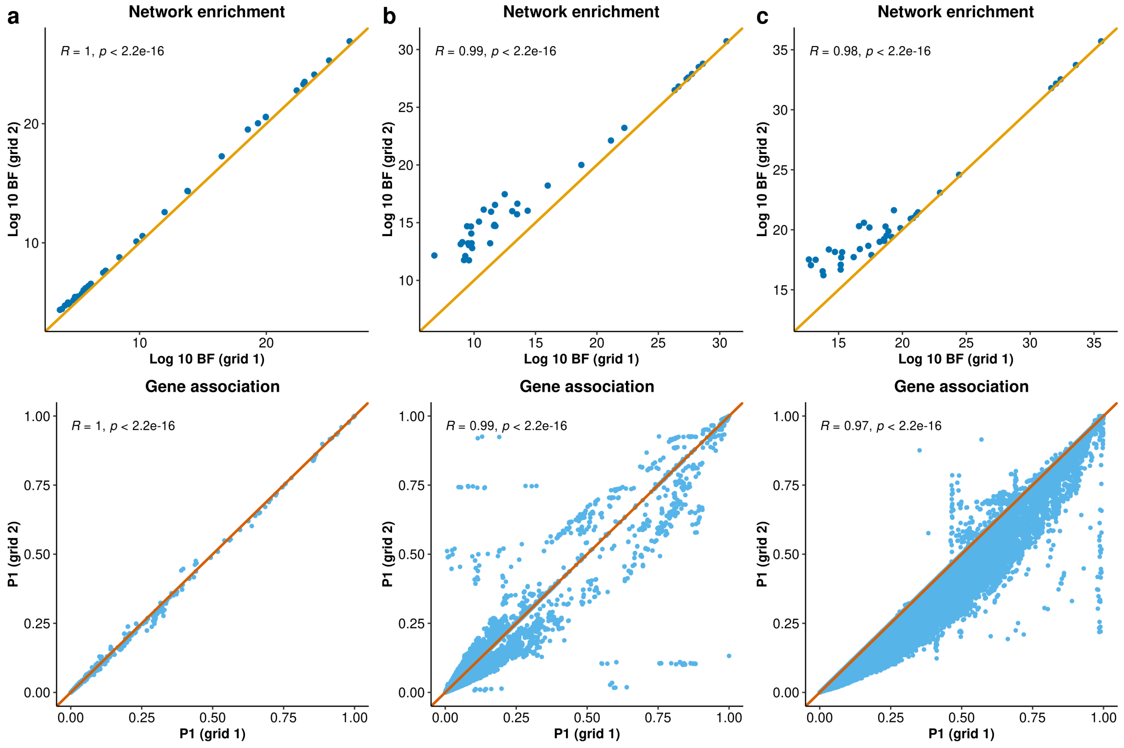

#### Supplementary Figure 19

**Comparison of two LDSC output statistics in simulations.** On each simulated dataset two LDSC methods, LDSC-baseline (Finucane et al. 2015) and LDSC-baselineLD (Gazal et al. 2017), are used to test network enrichments. Each LDSC method produces two statistics, enrichment $P$-value (denoted as `Enrichment_p` in the software) and enrichment coefficient $Z$-score (`Coefficient_z-score`), both of which can be used to rank the significance of enrichments. In each panel, each point represent a simulation scenario that contains 200 positive and 200 negative independent datasets (**Supplementary Figures 1-6 and 9**). The $x$- and $y$-axis of each point show AUROC values evaluated on the basis of enrichment $P$-value and coefficient $Z$-score respectively. In total there are 89 simulation scenarios of assessing network enrichments in this study, and thus each panel below consists of 89 points. The diagonal lines in both panels have slope 1 and intercept 0.

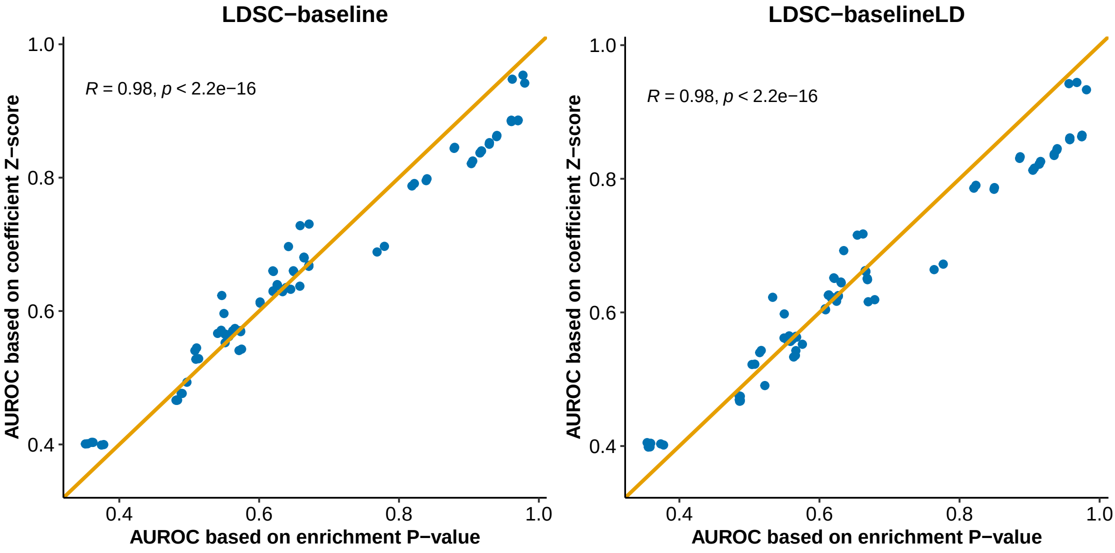

### Supplementary Tables

#### Supplementary Table 1

Paired high-throughput sequencing data of gene expression and chromatin accessibility from 207 biosamples that are used to construct 38 TF-TG regulatory networks.

| ID | Cell type or tissue | Expression ID | Accessibility ID | Network |
|---|---|---|---|---|
| 1 | Hematopoietic MPP cell | ENCSR000CUA | ENCSR115YPI | Omnibus |
| 2 | G-CSF-mobilized HSC | GSM909310 | GSM530657 | Omnibus |
| 3 | Fetal adrenal gland | GSM1220584 | GSM530653 | Omnibus |
| 4 | CD4 primary cell | GSM1059487 | GSM665839 | CD4 cell; Omnibus |
| 5 | CD8 primary cell | GSM1060237 | GSM665838 | CD8 cell; Omnibus |
| 6 | Primary monocyte | GSM1220575 | GSM701541 | Monocyte; Omnibus |
| 7 | Primary B cell | GSM1220576 | GSM701507 | B cell; Omnibus |
| 8 | Primary T cell | GSM1220574 | GSM701516 | Omnibus |
| 9 | CD4 primary cell | GSM1060238 | GSM701539 | CD4 cell; Omnibus |
| 10 | Primary NK cell | GSM1220577 | GSM701508 | NK cell; Omnibus |
| 11 | CD8 primary cell | GSM1060239 | GSM701540 | CD8 cell; Omnibus |
| 12 | Fetal intestine large | GSM1059485 | GSM701495 | Omnibus |
| 13 | Fetal intestine large | GSM1059509 | GSM701531 | Omnibus |
| 14 | Fetal intestine small | GSM1059486 | GSM701496 | Omnibus |
| 15 | Fetal intestine small | GSM1059508 | GSM701530 | Omnibus |
| 16 | Fetal kidney | GSM1059512 | GSM701515 | Omnibus |
| 17 | Fetal renal cortex | GSM1059496 | GSM701529 | Omnibus |
| 18 | Fetal renal cortex | GSM1059515 | GSM701532 | Omnibus |
| 19 | Fetal renal cortex, right | GSM1059498 | GSM701517 | Omnibus |
| 20 | Fetal renal pelvis, left | GSM1059514 | GSM701520 | Omnibus |
| 21 | Fetal renal pelvis, right | GSM1059500 | GSM701518 | Omnibus |
| 22 | Fetal lung, left | GSM1101684 | GSM701524 | Omnibus |
| 23 | Fetal lung, left | GSM1101685 | GSM701527 | Omnibus |
| 24 | Fetal lung, left | GSM1101687 | GSM701534 | Omnibus |
| 25 | Fetal lung, right | GSM1101683 | GSM701523 | Omnibus |
| 26 | Fetal muscle, arm | GSM1101689 | GSM701535 | Omnibus |
| 27 | Fetal muscle, back | GSM1101690 | GSM701536 | Omnibus |
| 28 | Fetal muscle trunk | GSM1101688 | GSM701533 | Omnibus |
| 29 | Fetal muscle, upper limb | GSM1101686 | GSM701522 | Omnibus |
| 30 | Fetal spleen | GSM1220588 | GSM701509 | Omnibus |
| 31 | Fetal stomach | GSM1220587 | GSM701521 | Omnibus |
| 32 | Fetal stomach | GSM1220583 | GSM701538 | Omnibus |
| 33 | Fetal stomach | GSM1220594 | GSM701538 | Omnibus |
| 34 | Fetal thymus | GSM1220591 | GSM701497 | Omnibus |
| 35 | Fetal thymus | GSM1220593 | GSM701537 | Omnibus |
| 36 | Fetal thymus | GSM1220593 | GSM774230 | Omnibus |
| 37 | Fetal intestine large | GSM1059516 | GSM774213 | Omnibus |
| 38 | Fetal intestine large | GSM1059518 | GSM774220 | Omnibus |
| 39 | Fetal intestine large | GSM1059520 | GSM774228 | Omnibus |
| 40 | Treg primary cell | GSM1059522 | GSM774233 | Omnibus |
| 41 | Treg primary cell | GSM1059507 | GSM774216 | Omnibus |
| 42 | Fetal renal pelvis | GSM1059524 | GSM774222 | Omnibus |
| 43 | Fetal lung, right | GSM1101691 | GSM774227 | Omnibus |
| 44 | Fetal lung, right | GSM1101692 | GSM774231 | Omnibus |
| 45 | Fetal muscle, arm | GSM1101694 | GSM774226 | Omnibus |

*(continued)*

| ID | Cell type or tissue | Expression ID | Accessibility ID | Network |
|---|---|---|---|---|
| 46 | Fetal muscle, arm | GSM1101695 | GSM774234 | Omnibus |
| 47 | Fetal muscle, arm | GSM1101682 | GSM774239 | Omnibus |
| 48 | Fetal muscle, back | GSM1220596 | GSM774224 | Omnibus |
| 49 | Fetal muscle, back | GSM1101696 | GSM774235 | Omnibus |
| 50 | Fetal muscle leg | GSM1101698 | GSM774242 | Omnibus |
| 51 | Fetal muscle, lower limb | GSM1101697 | GSM774238 | Omnibus |
| 52 | Treg primary cell | GSM1220590 | GSM774212 | Omnibus |
| 53 | Treg primary cell | GSM1220597 | GSM774232 | Omnibus |
| 54 | FK primary cell | GSM941745 | GSM817196 | Omnibus |
| 55 | FK primary cell | GSM941745 | GSM817197 | Omnibus |
| 56 | FF primary cell | GSM941744 | GSM817169 | Omnibus |
| 57 | FF primary cell | GSM941744 | GSM817170 | Omnibus |
| 58 | Melanocyte | GSM941743 | GSM1027307 | Omnibus |
| 59 | Melanocyte | GSM941743 | GSM1027312 | Omnibus |
| 60 | Fetal adrenal gland | GSM1059505 | GSM817165 | Omnibus |
| 61 | Fetal adrenal gland | GSM1059525 | GSM817167 | Omnibus |
| 62 | Fetal heart | GSM1059495 | GSM817220 | Omnibus |
| 63 | Fetal kidney | GSM1059510 | GSM817159 | Omnibus |
| 64 | Fetal kidney, left | GSM1059526 | GSM817181 | Omnibus |
| 65 | Fetal renal cortex, left | GSM1059528 | GSM817190 | Omnibus |
| 66 | Fetal renal cortex, left | GSM1059489 | GSM817203 | Omnibus |
| 67 | Fetal renal cortex, left | GSM1059492 | GSM817211 | Omnibus |
| 68 | Fetal renal cortex, right | GSM1059488 | GSM817202 | Omnibus |
| 69 | Fetal renal cortex, right | GSM1059499 | GSM817210 | Omnibus |
| 70 | Treg primary cell | GSM1059493 | GSM817218 | Omnibus |
| 71 | Fetal renal pelvis, left | GSM1059491 | GSM817193 | Omnibus |
| 72 | Fetal renal pelvis, left | GSM1059504 | GSM817205 | Omnibus |
| 73 | Fetal renal pelvis, left | GSM1059503 | GSM817209 | Omnibus |
| 74 | Fetal renal pelvis, right | GSM1059490 | GSM817192 | Omnibus |
| 75 | Fetal renal pelvis, right | GSM1059502 | GSM817204 | Omnibus |
| 76 | Fetal renal pelvis, right | GSM1059501 | GSM817208 | Omnibus |
| 77 | Fetal lung, left | GSM1101693 | GSM817168 | Omnibus |
| 78 | Fetal lung, left | GSM1101699 | GSM817175 | Omnibus |
| 79 | Fetal lung, left | GSM1101708 | GSM817185 | Omnibus |
| 80 | Fetal lung, right | GSM1101707 | GSM817174 | Omnibus |
| 81 | Fetal lung, right | GSM1101709 | GSM817186 | Omnibus |
| 82 | Fetal muscle, arm | GSM1101701 | GSM817178 | Omnibus |
| 83 | Fetal muscle, arm | GSM1101710 | GSM817191 | Omnibus |
| 84 | Fetal muscle, arm | GSM1101674 | GSM817214 | Omnibus |
| 85 | Fetal muscle, arm | GSM1101675 | GSM817216 | Omnibus |
| 86 | Fetal muscle, back | GSM1101703 | GSM817171 | Omnibus |
| 87 | Fetal muscle, back | GSM1101704 | GSM817182 | Omnibus |
| 88 | Fetal muscle, back | GSM1101661 | GSM817200 | Omnibus |
| 89 | Fetal muscle, back | GSM1101663 | GSM817207 | Omnibus |
| 90 | Fetal muscle, back | GSM1101677 | GSM817217 | Omnibus |
| 91 | Fetal muscle leg | GSM1101705 | GSM817179 | Omnibus |
| 92 | Fetal muscle leg | GSM1101706 | GSM817183 | Omnibus |
| 93 | Fetal muscle leg | GSM1101664 | GSM817206 | Omnibus |
| 94 | Fetal muscle leg | GSM1101671 | GSM817213 | Omnibus |
| 95 | Fetal muscle, back | GSM1101676 | GSM817212 | Omnibus |

*(continued)*

| ID | Cell type or tissue | Expression ID | Accessibility ID | Network |
|---|---|---|---|---|
| 96 | Fetal spinal cord | GSM1101711 | GSM817189 | Omnibus |
| 97 | Treg primary cell | GSM1220598 | GSM817173 | Omnibus |
| 98 | Treg primary cell | GSM1220600 | GSM817199 | Omnibus |
| 99 | Fetal thymus | GSM1220599 | GSM817172 | Omnibus |
| 100 | Fibroblasts, fetal skin | GSM1101681 | GSM878635 | Omnibus |
| 101 | Fibroblasts, fetal skin | GSM1101666 | GSM878623 | Omnibus |
| 102 | Fibroblasts, fetal skin | GSM1101667 | GSM878634 | Omnibus |
| 103 | Fibroblasts, fetal skin | GSM1101668 | GSM878633 | Omnibus |
| 104 | Fetal heart | GSM1059494 | GSM878630 | Omnibus |
| 105 | Fetal renal cortex | GSM1059497 | GSM878629 | Omnibus |
| 106 | Fetal muscle, arm | GSM1101700 | GSM878610 | Omnibus |
| 107 | Fetal muscle, arm | GSM1101673 | GSM878625 | Omnibus |
| 108 | Fetal muscle, back | GSM1101678 | GSM878632 | Omnibus |
| 109 | Fetal muscle leg | GSM1101670 | GSM878626 | Omnibus |
| 110 | Fetal muscle leg | GSM1101672 | GSM878631 | Omnibus |
| 111 | Fetal muscle leg | GSM1220579 | GSM878653 | Omnibus |
| 112 | Fetal muscle trunk | GSM1220582 | GSM878654 | Omnibus |
| 113 | Treg primary cell | GSM1220581 | GSM878659 | Omnibus |
| 114 | Fetal spinal cord | GSM1220580 | GSM878661 | Omnibus |
| 115 | Fetal thymus | GSM1220578 | GSM878657 | Omnibus |
| 116 | Pancreas | GSM1010966 | GSM1027326 | Omnibus |
| 117 | Gastric | GSM1010960 | GSM1027325 | Omnibus |
| 118 | Ovary | GSM1010948 | GSM1027334 | Omnibus |
| 119 | Gastric | GSM1120306 | GSM1027320 | Omnibus |
| 120 | Psoas Muscle | GSM1120310 | GSM1027321 | Omnibus |
| 121 | Pancreas | GSM1120309 | GSM1027335 | Omnibus |
| 122 | Fetal adrenal gland | GSM1060240 | GSM1027310 | Omnibus |
| 123 | Fetal adrenal gland | GSM1060241 | GSM1027311 | Omnibus |
| 124 | Fetal ovary | GSM1101662 | GSM1027306 | Omnibus |
| 125 | Fetal spinal cord | GSM1101679 | GSM1027308 | Omnibus |
| 126 | Fetal kidney | GSM1059523 | GSM774221 | Omnibus |
| 127 | BE2C | ENCSR000BYK | ENCFF001CNX | Omnibus |
| 128 | BJ | ENCSR000COP | ENCFF001COH | Omnibus |
| 129 | GM12878 | ENCSR000AEE | ENCFF000SKV | Omnibus |
| 130 | H1 BMP4-derived TB cell | ENCSR762CJN | ENCFF983HYM | Omnibus |
| 131 | H1-hESC | ENCSR000COU | ENCFF876IHU | Omnibus |
| 132 | H7-hESC | ENCSR490SQH | ENCFF000SOL | Omnibus |
| 133 | HeLa-S3 | ENCSR552EGO | ENCFF000SPR | Omnibus |
| 134 | HepG2 | ENCSR931WGT | ENCFF000SQJ | Omnibus |
| 135 | Jurkat | ENCSR000BXX | ENCFF001DOT | Omnibus |
| 136 | K562 | ENCSR000CPH | ENCFF000SWS | Omnibus |
| 137 | MCF-7 | ENCSR000CPT | ENCFF000SYX | Omnibus |
| 138 | Panc1 | ENCSR000BYM | ENCFF001EDO | Omnibus |
| 139 | T47D | ENCSR000BYF | ENCFF000TET | Omnibus |
| 140 | Testis | ENCFF673KOG | ENCFF048IOT | Testis; Omnibus |
| 141 | Testis | ENCFF438ZIY | ENCFF066TTB | Testis; Omnibus |
| 142 | Stomach | ENCFF155AGD | ENCFF025QEQ | Stomach; Omnibus |
| 143 | Stomach | ENCFF930JZC | ENCFF184XKI | Stomach; Omnibus |
| 144 | Stomach | ENCFF535HFG | ENCFF758MDE | Stomach; Omnibus |
| 145 | Ovary | ENCFF232UER | ENCFF420ECZ | Ovary; Omnibus |

*(continued)*

| ID | Cell type or tissue | Expression ID | Accessibility ID | Network |
|---|---|---|---|---|
| 146 | Ovary | ENCFF957VSL | ENCFF247HQX | Ovary; Omnibus |
| 147 | Uterus | ENCFF624VKT | ENCFF678QTC | Uterus; Omnibus |
| 148 | Vagina | ENCFF855LJY | ENCFF243NYB | Vagina; Omnibus |
| 149 | Liver | ENCFF347TXW | ENCFF244IRQ | Liver; Omnibus |
| 150 | Pancreas | ENCFF204BZU | ENCFF011ZXN | Pancreas; Omnibus |
| 151 | Pancreas | ENCFF368OME | ENCFF646IYT | Pancreas; Omnibus |
| 152 | Transverse colon | ENCFF874CRV | ENCFF629OKO | Transverse colon; Omnibus |
| 153 | Transverse colon | ENCFF505DGK | ENCFF159DOO | Transverse colon; Omnibus |
| 154 | Transverse colon | ENCFF332OML | ENCFF746GDH | Transverse colon; Omnibus |
| 155 | Transverse colon | ENCFF014IOZ | ENCFF715LGJ | Transverse colon; Omnibus |
| 156 | Sigmoid colon | ENCFF934VJV | ENCFF482HAC | Sigmoid colon; Omnibus |
| 157 | Sigmoid colon | ENCFF192TOO | ENCFF387OFS | Sigmoid colon; Omnibus |
| 158 | Sigmoid colon | ENCFF179AVF | ENCFF692DLP | Sigmoid colon; Omnibus |
| 159 | Sigmoid colon | ENCFF146ISW | ENCFF754DCA | Sigmoid colon; Omnibus |
| 160 | Terminal ileum | ENCFF336EQS | ENCFF748WZP | Terminal ileum; Omnibus |
| 161 | Terminal ileum | ENCFF187NLN | ENCFF474HAC | Terminal ileum; Omnibus |
| 162 | Terminal ileum | ENCFF715EVL | ENCFF603CAV | Terminal ileum; Omnibus |
| 163 | Tibial nerve | ENCFF904FEZ | ENCFF226ZCG | Tibial nerve; Omnibus |
| 164 | Tibial nerve | ENCFF241VXT | ENCFF372VAD | Tibial nerve; Omnibus |
| 165 | Aorta | ENCFF127YOW | ENCFF984OPE | Aorta; Omnibus |
| 166 | Thyroid | ENCFF625QLJ | ENCFF942YPH | Thyroid; Omnibus |
| 167 | Thyroid | ENCFF376HXE | ENCFF743DWF | Thyroid; Omnibus |
| 168 | Thyroid | ENCFF194VCC | ENCFF611LJR | Thyroid; Omnibus |
| 169 | Thyroid | ENCFF509HKT | ENCFF530YPZ | Thyroid; Omnibus |
| 170 | Left ventricle | ENCFF470AAE | ENCFF702IJE | Left ventricle; Omnibus |
| 171 | Left ventricle | ENCFF870BQB | ENCFF906UQF | Left ventricle; Omnibus |
| 172 | Spleen | ENCFF087VIO | ENCFF294ZCT | Spleen; Omnibus |
| 173 | Spleen | ENCFF916TWO | ENCFF183HFJ | Spleen; Omnibus |
| 174 | Subcutaneous adipose | ENCFF967YNU | ENCFF159RKV | Subcutaneous adipose; Omnibus |
| 175 | Prostate | ENCFF454GAP | ENCFF007NTA | Prostate; Omnibus |
| 176 | Prostate | ENCFF110THT | ENCFF670GFY | Prostate; Omnibus |
| 177 | Adrenal gland | ENCFF254VJQ | ENCFF970LXV | Adrenal gland; Omnibus |
| 178 | Adrenal gland | ENCFF250FNI | ENCFF374OSU | Adrenal gland; Omnibus |
| 179 | Adrenal gland | ENCFF587WDK | ENCFF279GVQ | Adrenal gland; Omnibus |
| 180 | Adrenal gland | ENCFF036ERC | ENCFF430AHU | Adrenal gland; Omnibus |
| 181 | Sun-exposed skin | ENCFF153NMV | ENCFF238BRB | Sun-exposed skin; Omnibus |
| 182 | Gastroesophageal junction | ENCFF338HIM | ENCFF293BRN | Gastroesophageal junction; Omnibus |
| 183 | Gastroesophageal junction | ENCFF749AJA | ENCFF765DRX | Gastroesophageal junction; Omnibus |
| 184 | Esophagus muscularis | ENCFF660MOE | ENCFF208ILP | Esophagus muscularis; Omnibus |
| 185 | Esophagus muscularis | ENCFF650KLZ | ENCFF762MNF | Esophagus muscularis; Omnibus |
| 186 | Atrial appendage | ENCFF406YAV | ENCFF513LVE | Atrial appendage; Omnibus |
| 187 | Atrial appendage | ENCFF701KTC | ENCFF939RTU | Atrial appendage; Omnibus |
| 188 | Esophagus mucosa | ENCFF560ZBX | ENCFF846JID | Esophagus mucosa; Omnibus |
| 189 | Esophagus mucosa | ENCFF301YAH | ENCFF542ZZY | Esophagus mucosa; Omnibus |
| 190 | Breast | ENCFF557XAK | ENCFF589ACB | Breast; Omnibus |
| 191 | Breast | ENCFF912TDO | ENCFF585YAB | Breast; Omnibus |
| 192 | Lung | ENCFF049ETK | ENCFF022PJN | Lung; Omnibus |
| 193 | Lung | ENCFF841RGX | ENCFF985OXZ | Lung; Omnibus |
| 194 | Lung | ENCFF360GZR | ENCFF803ZER | Lung; Omnibus |
| 195 | Visceral omentum | ENCFF324LPM | ENCFF519QGD | Visceral omentum; Omnibus |

*(continued)*

| ID | Cell type or tissue | Expression ID | Accessibility ID | Network |
|---|---|---|---|---|
| 196 | Visceral omentum | ENCFF766ECG | ENCFF880CAD | Visceral omentum; Omnibus |
| 197 | Skeletal muscle | ENCFF342MON | ENCFF508WVY | Skeletal muscle; Omnibus |
| 198 | Skeletal muscle | ENCFF728KWY | ENCFF086TID | Skeletal muscle; Omnibus |
| 199 | Skeletal muscle | ENCFF092VEZ | ENCFF461GFF | Skeletal muscle; Omnibus |
| 200 | Skeletal muscle | ENCFF151LCI | ENCFF858PLS | Skeletal muscle; Omnibus |
| 201 | Skeletal muscle | ENCFF149OTO | ENCFF478ARL | Skeletal muscle; Omnibus |
| 202 | Cortex | GTEx V7 | ENCSR000EIY | Cortex |
| 203 | Cortex | GTEx V7 | ENCSR000EIK | Cortex |
| 204 | Cerebellum | GTEx V7 | ENCSR000EIJ | Cerebellum |
| 205 | Caudate | GTEx V7 | ENCSR015BGH | Caudate |
| 206 | Putamen | GTEx V7 | ENCSR493VDS | Putamen |
| 207 | Hippocampus | GTEx V7 | ENCSR584LUZ | Hippocampus |

#### Supplementary Table 2

GWAS publications, total sample sizes and numbers of genetic variants analyzed by RSS-NET for 18 human traits. Click blue links to view publications online.

| Trait (abbreviation) | PMID | # of SNPs (analyzed) | Sample size (cases+controls) |
|---|---|---|---|
| Alzheimer's disease (LOAD) | 24162737 | 1,136,997 | 17,008+37,154 |
| Neuroticism (NEU) | 27089181 | 1,119,108 | 170,911 |
| Schizophrenia (SCZ) | 25056061 | 1,113,442 | 152,805 |
| Body mass index (BMI) | 25673413 | 1,012,465 | 234,069 |
| Height (HEIGHT) | 25282103 | 1,064,575 | 253,288 |
| Waist-to-hip ratio (WAIST) | 25673412 | 1,008,898 | 142,762 |
| Crohn's disease (CD) | 26192919 | 1,064,533 | 5,956+14,927 |
| Inflammatory bowel disease (IBD) | 26192919 | 1,081,481 | 12,882+21,770 |
| Rheumatoid arthritis (RA) | 24390342 | 1,158,064 | 14,361+43,923 |
| Ulcerative colitis (UC) | 26192919 | 1,092,170 | 6,968+20,464 |
| Breast cancer (BC) | 23535729 | 1,188,902 | 15,863+40,022 |
| Atrial fibrillation (AF) | 28416818 | 1,175,059 | 17,931+115,142 |
| Coronary artery disease (CAD) | 26343387 | 1,121,322 | 60,801+123,504 |
| High-density lipoprotein (HDL) | 20686565 | 1,032,214 | 99,900 |
| Heart rate (HR) | 23583979 | 1,066,168 | 92,355 |
| Low-density lipoprotein (LDL) | 20686565 | 1,030,397 | 95,454 |
| Myocardial infarction (MI) | 26343387 | 1,111,568 | 42,561+123,504 |
| Type 2 diabetes (T2D) | 22885922 | 1,047,618 | 12,171+56,862 |

#### Supplementary Table 3

To confirm that RSS-NET network enrichment results are unlikely to be driven by generic regulatory enrichments harbored in the vicinity of genes, we perform a "near-gene" control analysis. Specifically, we create a "near-gene" control network with 18,334 protein-coding autosomal genes as nodes and no edges, and then analyze this control with RSS-NET on the same GWAS data for 18 traits. The near-gene control analysis is formalized as:

$$\begin{aligned}
\widehat{\boldsymbol{\beta}} &\sim \mathcal{N}(\widehat{\mathbf{S}}\widehat{\mathbf{R}}\widehat{\mathbf{S}}^{-1}\boldsymbol{\beta},\ \widehat{\mathbf{S}}\widehat{\mathbf{R}}\widehat{\mathbf{S}}),\\
\beta_j &\sim \pi_j \cdot \mathcal{N}(\mu_j,\ \sigma_0^2) + (1-\pi_j)\cdot\delta_0,\\
\pi_j &= 1\,/\,[1+10^{-(\theta_0+a_j\theta)}],\\
\mu_j &= \textstyle\sum_{g\in G_j} c_{jg}\cdot\gamma_{jg},\\
\gamma_{jg} &\sim \mathcal{N}(0,\ \sigma^2),
\end{aligned}$$

where $a_j = 1$ if SNP $j$ is within 100 kb of transcribed region of any protein-coding autosomal gene and $a_j = 0$ otherwise, $G_j$ denotes the set of all protein-coding genes within 1 Mb window of SNP $j$, and $c_{jg}$ measures the relative impact of SNP $j$ on gene $g$ (derived from eQTLGen *cis*-eQTL results; see **Supplementary Notes**).

Panel **a** Using the same GWAS data of each trait and the same hyper-parameter grid, we compare BFs between each network and the near-gene control under four enrichment models: (1) $M_{11}$: $\theta > 0$ and $\sigma^2 = 0$; (2) $M_{12}$: $\theta = 0$ and $\sigma^2 > 0$; (3) $M_{13}$: $\theta > 0$ and $\sigma^2 > 0$; (4) $M_1$: $\theta > 0$ or $\sigma^2 > 0$. For all 38 networks and the near-gene control, BF computations are based on the same baseline model ($M_0$: $\theta = 0$ and $\sigma^2 = 0$). Each row below reports the number of networks with BFs greater than near-gene control BF for each trait and each enrichment model. Trait abbreviations are defined in **Supplementary Table 2**.

| | # of networks | | | |
|---|---|---|---|---|
| Trait | $M_{11}$ | $M_{12}$ | $M_{13}$ | $M_1$ |
| AF | 32 | 38 | 38 | 38 |
| BC | 33 | 38 | 38 | 38 |
| BMI | 9 | 38 | 38 | 38 |
| CAD | 38 | 38 | 38 | 38 |
| HDL | 38 | 38 | 38 | 38 |
| HR | 32 | 38 | 38 | 38 |
| LDL | 38 | 38 | 38 | 38 |
| MI | 38 | 38 | 38 | 38 |
| NEU | 33 | 38 | 38 | 38 |
| RA | 38 | 38 | 38 | 38 |
| SCZ | 9 | 38 | 38 | 38 |
| WAIST | 38 | 38 | 38 | 38 |
| UC | 38 | 22 | 34 | 34 |
| LOAD | 38 | 13 | 11 | 11 |
| IBD | 38 | 0 | 6 | 6 |
| T2D | 2 | 0 | 21 | 5 |
| CD | 38 | 0 | 5 | 0 |
| HEIGHT | 38 | 0 | 0 | 0 |

Panel **b** Using the same GWAS data of each trait and the external methods (LDSC, Pascal), we compare enrichment $P$-values between each network and the near-gene control. Each row below reports the number of networks with enrichment $P$-value smaller than near-gene control enrichment $P$-value for each trait and each external method. Numerical values of LDSC and Pascal analyses are available at https://suwonglab.github.io/rss-net/results.html.

| | # of networks | | | | | |
|---|---|---|---|---|---|---|
| | LDSC methods | | Pascal methods | | | |
| Trait | baseline | baselineLD | max-chi | max-emp | sum-chi | sum-emp |
| RA | 38 | 38 | 0 | 0 | 2 | 0 |
| IBD | 36 | 37 | 0 | 0 | 0 | 0 |
| UC | 37 | 37 | 1 | 0 | 1 | 0 |
| HR | 31 | 36 | 0 | 0 | 0 | 0 |
| WAIST | 37 | 36 | 1 | 0 | 27 | 24 |
| CD | 32 | 34 | 0 | 0 | 0 | 0 |
| BC | 28 | 32 | 0 | 2 | 1 | 0 |
| MI | 31 | 30 | 0 | 0 | 15 | 15 |
| CAD | 23 | 23 | 0 | 0 | 8 | 0 |
| T2D | 25 | 22 | 0 | 0 | 5 | 6 |
| LDL | 24 | 18 | 0 | 0 | 1 | 0 |
| NEU | 18 | 16 | 0 | 0 | 0 | 0 |
| HEIGHT | 6 | 11 | 0 | 0 | 0 | 0 |
| HDL | 6 | 10 | 0 | 0 | 1 | 0 |
| AF | 5 | 5 | 0 | 0 | 7 | 0 |
| LOAD | 1 | 2 | 0 | 0 | 1 | 1 |
| SCZ | 2 | 2 | 0 | 0 | 0 | 0 |
| BMI | 0 | 0 | 0 | 0 | 0 | 0 |

#### Supplementary Table 4

We compute Pearson correlations between five network features and log 10 enrichment BFs, either across 512 trait-network pairs that pass the near-gene control (**Supplementary Table 3**), or across all 684 trait-network pairs (18 traits and 38 networks). We use R built-in function `cor.test` for all correlation analyses. All tests are two-sided.

| | Pearson $R$ ($P$-value) | |
|---|---|---|
| Network feature | 512 pairs | 684 pairs |
| % of SNPs in a network | -0.0304 (0.4925) | -0.0403 (0.2924) |
| # of TF-TG edges | -0.0092 (0.8353) | 0.0025 (0.9479) |
| # of nodes (TF or TG) | -0.0535 (0.2272) | -0.0422 (0.2702) |
| # of TGs | -0.0531 (0.2308) | -0.0424 (0.2686) |
| # of TFs | -0.0403 (0.3625) | -0.0184 (0.6312) |

#### Supplementary Table 5

We compute Pearson correlations between 73 binnary functional annotations in LDSC baselineLD v2.1 (Gazal et al. 2017) and log 10 enrichment BFs, either across 512 trait-network pairs that pass the near-gene control (**Supplementary Table 3**), or across all 684 trait-network pairs (18 traits and 38 networks). Specifically, for a given functional annotation, we estimate the correlation between log 10 BFs and proportion of SNPs falling into both a network and this functional category, across all trait-network pairs. Rows are ranked by Pearson $P$-values based on 512 trait-network pairs. The Bonferroni cutoff is $0.05/73 = 6.8 \times 10^{-4}$. The rest is the same as **Supplementary Table 4**.

| | Pearson $R$ (- log 10 $P$-value) | |
|---|---|---|
| Functional annotation | 512 pairs | 684 pairs |
| BivFlnk | -0.1295 (2.4786) | -0.1252 (2.9849) |
| TSS_Hoffman | -0.1256 (2.3543) | -0.1547 (4.3146) |
| BivFlnk_500 | -0.1195 (2.1668) | -0.1166 (2.6468) |
| TSS_Hoffman_500 | -0.1178 (2.1193) | -0.1465 (3.9183) |
| H3K9ac_peaks_Trynka | -0.0795 (1.1410) | -0.1041 (2.1907) |
| Promoter_UCSC_500 | -0.0777 (1.1022) | -0.1137 (2.5363) |
| Promoter_UCSC | -0.0772 (1.0926) | -0.1134 (2.5254) |
| PromoterFlanking_Hoffman_500 | -0.0742 (1.0296) | -0.1125 (2.4941) |
| Enhancer_Hoffman | -0.0707 (0.9579) | -0.0907 (1.7518) |
| H3K9ac_Trynka | -0.0704 (0.9521) | -0.0939 (1.8543) |
| PromoterFlanking_Hoffman | -0.0696 (0.9357) | -0.1144 (2.5616) |
| H3K4me3_peaks_Trynka | -0.0681 (0.9063) | -0.0941 (1.8597) |
| Enhancer_Hoffman_500 | -0.0671 (0.8879) | -0.0891 (1.7044) |
| H3K4me3_Trynka | -0.0647 (0.8428) | -0.0903 (1.7393) |
| WeakEnhancer_Hoffman | -0.0641 (0.8313) | -0.0798 (1.4331) |
| H3K9ac_Trynka_500 | -0.0606 (0.7666) | -0.0852 (1.5871) |
| SuperEnhancer_Hnisz_500 | -0.0596 (0.7490) | -0.0806 (1.4540) |
| SuperEnhancer_Hnisz | -0.0595 (0.7481) | -0.0804 (1.4496) |
| Enhancer_Andersson | -0.0590 (0.7386) | -0.0747 (1.2932) |
| WeakEnhancer_Hoffman_500 | -0.0564 (0.6929) | -0.0750 (1.3024) |
| UTR_5_UCSC_500 | -0.0562 (0.6897) | -0.0933 (1.8339) |
| Enhancer_Andersson_500 | -0.0557 (0.6807) | -0.0701 (1.1735) |
| H3K4me3_Trynka_500 | -0.0536 (0.6465) | -0.0796 (1.4284) |
| TFBS_ENCODE | -0.0522 (0.6230) | -0.0767 (1.3473) |
| UTR_5_UCSC | -0.0497 (0.5823) | -0.0787 (1.4009) |
| UTR_3_UCSC_500 | -0.0480 (0.5559) | -0.0813 (1.4738) |
| MAFbin1 | -0.0474 (0.5461) | -0.0508 (0.7331) |
| H3K4me1_peaks_Trynka | -0.0460 (0.5249) | -0.0704 (1.1819) |
| H3K27ac_PGC2 | -0.0453 (0.5144) | -0.0712 (1.2030) |
| CTCF_Hoffman_500 | -0.0447 (0.5043) | -0.0655 (1.0606) |
| DGF_ENCODE | -0.0432 (0.4828) | -0.0676 (1.1124) |
| H3K27ac_PGC2_500 | -0.0429 (0.4775) | -0.0692 (1.1512) |
| Coding_UCSC_500 | -0.0424 (0.4710) | -0.0753 (1.3093) |

*(continued)*

| Functional annotation | Pearson $R$ (- log 10 $P$-value) 512 pairs | 684 pairs |
|---|---|---|
| CTCF_Hoffman | -0.0421 (0.4667) | -0.0618 (0.9732) |
| MAFbin2 | -0.0420 (0.4642) | -0.0530 (0.7797) |
| H3K27ac_Hnisz | -0.0412 (0.4531) | -0.0665 (1.0860) |
| TFBS_ENCODE_500 | -0.0408 (0.4474) | -0.0659 (1.0700) |
| H3K27ac_Hnisz_500 | -0.0402 (0.4386) | -0.0658 (1.0682) |
| H3K4me1_Trynka | -0.0377 (0.4040) | -0.0643 (1.0314) |
| FetalDHS_Trynka | -0.0375 (0.4013) | -0.0615 (0.9672) |
| DHS_peaks_Trynka | -0.0365 (0.3869) | -0.0610 (0.9557) |
| Transcr_Hoffman | -0.0359 (0.3794) | -0.0728 (1.2426) |
| MAFbin3 | -0.0353 (0.3710) | -0.0557 (0.8365) |
| DGF_ENCODE_500 | -0.0350 (0.3675) | -0.0609 (0.9532) |
| FetalDHS_Trynka_500 | -0.0347 (0.3635) | -0.0589 (0.9067) |
| DHS_Trynka | -0.0347 (0.3629) | -0.0594 (0.9191) |
| H3K4me1_Trynka_500 | -0.0336 (0.3491) | -0.0606 (0.9464) |
| Intron_UCSC_500 | -0.0330 (0.3412) | -0.0661 (1.0752) |
| Intron_UCSC | -0.0330 (0.3408) | -0.0662 (1.0768) |
| non_synonymous | -0.0323 (0.3313) | -0.0639 (1.0221) |
| DHS_Trynka_500 | -0.0318 (0.3259) | -0.0575 (0.8757) |
| MAFbin4 | -0.0310 (0.3156) | -0.0557 (0.8374) |
| Vertebrate_phastCons46way_500 | -0.0309 (0.3136) | -0.0583 (0.8935) |
| Mammal_phastCons46way_500 | -0.0305 (0.3084) | -0.0577 (0.8810) |
| Transcr_Hoffman_500 | -0.0297 (0.2992) | -0.0602 (0.9356) |
| LindbladToh_500 | -0.0296 (0.2972) | -0.0554 (0.8312) |
| Primate_phastCons46way_500 | -0.0293 (0.2933) | -0.0565 (0.8541) |
| Vertebrate_phastCons46way | -0.0285 (0.2838) | -0.0529 (0.7776) |
| MAFbin5 | -0.0268 (0.2635) | -0.0561 (0.8447) |
| Mammal_phastCons46way | -0.0267 (0.2618) | -0.0511 (0.7392) |
| MAFbin6 | -0.0263 (0.2581) | -0.0576 (0.8780) |
| LindbladToh | -0.0248 (0.2402) | -0.0482 (0.6828) |
| MAFbin9 | -0.0244 (0.2348) | -0.0580 (0.8876) |
| Primate_phastCons46way | -0.0244 (0.2348) | -0.0498 (0.7131) |
| MAFbin8 | -0.0241 (0.2314) | -0.0580 (0.8862) |
| MAFbin7 | -0.0241 (0.2313) | -0.0558 (0.8396) |
| MAFbin10 | -0.0237 (0.2267) | -0.0567 (0.8579) |
| Coding_UCSC | -0.0236 (0.2266) | -0.0542 (0.8045) |
| GERP.RSsup4 | -0.0227 (0.2163) | -0.0461 (0.6402) |
| UTR_3_UCSC | -0.0220 (0.2079) | -0.0531 (0.7807) |
| Repressed_Hoffman_500 | -0.0176 (0.1599) | -0.0429 (0.5803) |
| Repressed_Hoffman | -0.0158 (0.1418) | -0.0411 (0.5472) |
| synonymous | -0.0135 (0.1191) | -0.0425 (0.5734) |

#### Supplementary Table 6

For 512 network-trait pairs passing the near-gene enrichment control (**Supplementary Table 3**), we compute BFs comparing the baseline model ($M_0$: $\theta = 0$ and $\sigma^2 = 0$) against three disjoint enrichment models: (1) $M_{11}$: $\theta > 0$ and $\sigma^2 = 0$; (2) $M_{12}$: $\theta = 0$ and $\sigma^2 > 0$; (3) $M_{13}$: $\theta > 0$ and $\sigma^2 > 0$. For each network-trait pair, we compare BFs based on $M_{11}$, $M_{12}$ and $M_{13}$, and define the "best" enrichment model as the one with the largest BF. Each row below shows, for each trait, the number of networks with $M_{11}$, $M_{12}$ and $M_{13}$ being the "best" model, respectively. Trait abbreviations are defined in **Supplementary Table 2**. **Figure 5(c)** is based on the table below.

| | # of networks | | | |
|---|---|---|---|---|
| Trait | $M_{11}$ | $M_{12}$ | $M_{13}$ | Total |
| AF | 0 | 0 | 38 | 38 |
| BC | 0 | 10 | 28 | 38 |
| BMI | 0 | 14 | 24 | 38 |
| CAD | 0 | 0 | 38 | 38 |
| HDL | 0 | 0 | 38 | 38 |
| HR | 0 | 0 | 38 | 38 |
| LDL | 0 | 0 | 38 | 38 |
| MI | 0 | 0 | 38 | 38 |
| NEU | 0 | 0 | 38 | 38 |
| RA | 0 | 0 | 38 | 38 |
| SCZ | 0 | 38 | 0 | 38 |
| WAIST | 0 | 37 | 1 | 38 |
| UC | 0 | 0 | 34 | 34 |
| LOAD | 0 | 0 | 11 | 11 |
| IBD | 0 | 0 | 6 | 6 |
| T2D | 2 | 0 | 3 | 5 |
| Total | 2 | 99 | 411 | 512 |

#### Supplementary Table 7

Correlations of RSS-NET enrichment log 10 BFs between 18 traits are computed over all 38 networks. Rows are ranked by Pearson $P$-values. The Bonferroni cutoff is $0.05/153 = 3.3 \times 10^{-4}$. Trait abbreviations are defined in **Supplementary Table 2**. Genetic correlations computed on the same GWAS summary data are retrieved from Watanabe et al. (2019), gwasATLAS_v20191115_GC.txt.gz (https://atlas.ctglab.nl/, accessed June 8, 2020).

| | | Estimate (- log 10 $P$-value) | |
|---|---|---|---|
| Trait 1 | Trait 2 | Log 10 BF correlation | Genetic correlation |
| CD | IBD | 0.9551 (19.8723) | 0.9058 (Inf) |
| IBD | UC | 0.9204 (15.5240) | 0.9017 (Inf) |
| LDL | LOAD | 0.8969 (13.5933) | 0.0682 (0.7100) |
| CAD | MI | 0.8954 (13.4826) | NA (NA) |
| CD | UC | 0.8342 (10.1112) | 0.6166 (31.1960) |
| CD | LDL | 0.7969 (8.6674) | -0.0706 (1.3799) |
| RA | UC | 0.7853 (8.2783) | 0.0550 (0.4411) |
| CD | LOAD | 0.7791 (8.0795) | -0.1196 (0.7964) |
| RA | SCZ | 0.7676 (7.7266) | -0.0430 (0.7792) |
| IBD | LOAD | 0.7548 (7.3575) | -0.1240 (0.8598) |
| CAD | HR | 0.7526 (7.2965) | -0.0116 (0.0980) |
| NEU | SCZ | 0.7335 (6.7908) | 0.1936 (6.5036) |
| BMI | SCZ | 0.7289 (6.6741) | -0.0789 (3.9818) |
| IBD | LDL | 0.7237 (6.5469) | -0.0552 (1.0600) |
| BMI | MI | 0.7131 (6.2964) | NA (NA) |
| IBD | RA | 0.7014 (6.0293) | 0.0522 (0.4496) |
| BMI | CAD | 0.6858 (5.6952) | 0.2046 (12.4028) |
| AF | BMI | 0.6827 (5.6303) | 0.2008 (18.0579) |
| HR | MI | 0.6743 (5.4617) | NA (NA) |
| BMI | RA | 0.6671 (5.3205) | 0.0500 (1.2590) |
| CD | RA | 0.6562 (5.1122) | 0.0393 (0.2860) |
| AF | CAD | 0.6520 (5.0354) | 0.1773 (7.4074) |
| AF | MI | 0.6491 (4.9826) | NA (NA) |
| CAD | HEIGHT | 0.6390 (4.8016) | -0.0963 (4.5005) |
| HEIGHT | MI | 0.6269 (4.5958) | NA (NA) |
| HDL | LDL | 0.6244 (4.5539) | -0.0375 (0.4878) |
| NEU | RA | 0.6050 (4.2420) | -0.0303 (0.2968) |
| LOAD | UC | 0.5880 (3.9847) | -0.1587 (1.0109) |
| LDL | UC | 0.5587 (3.5744) | -0.0548 (0.8752) |
| AF | HEIGHT | 0.5499 (3.4575) | 0.2466 (33.5271) |
| AF | RA | 0.5291 (3.1956) | 0.0864 (1.9809) |
| LDL | RA | 0.5131 (3.0062) | 0.0002 (0.0021) |
| BMI | HR | 0.4912 (2.7597) | -0.0548 (1.1392) |
| CAD | SCZ | 0.4684 (2.5202) | -0.0250 (0.4286) |
| BMI | NEU | 0.4586 (2.4222) | -0.0059 (0.0742) |

*(continued)*

| Trait 1 | Trait 2 | Estimate (- log 10 $P$-value) | |
|---|---|---|---|
| | | Log 10 BF correlation | Genetic correlation |
| HDL | MI | 0.4563 (2.3999) | NA (NA) |
| MI | SCZ | 0.4541 (2.3784) | NA (NA) |
| AF | SCZ | 0.4516 (2.3544) | -0.0131 (0.2132) |
| SCZ | UC | 0.4371 (2.2164) | 0.1122 (2.5792) |
| BC | IBD | -0.4332 (2.1810) | 0.2826 (2.6405) |
| AF | HR | 0.4170 (2.0364) | -0.0823 (1.0538) |
| BMI | LDL | 0.4158 (2.0256) | 0.0215 (0.3918) |
| LDL | MI | 0.4149 (2.0181) | NA (NA) |
| NEU | UC | 0.4034 (1.9198) | 0.0998 (1.0578) |
| BMI | HDL | 0.4029 (1.9161) | -0.2589 (18.2255) |
| HEIGHT | HR | 0.4012 (1.9018) | -0.0962 (2.6964) |
| HDL | LOAD | 0.4008 (1.8985) | 0.1559 (1.7379) |
| CAD | HDL | 0.3992 (1.8848) | -0.2526 (14.6796) |
| LOAD | RA | 0.3956 (1.8554) | -0.1284 (0.7667) |
| CD | BC | -0.3949 (1.8495) | 0.1349 (0.8437) |
| BMI | WAIST | 0.3773 (1.7091) | -0.0776 (2.5595) |
| BMI | UC | 0.3739 (1.6832) | -0.1015 (2.2981) |
| CD | HDL | 0.3616 (1.5904) | -0.0542 (0.8791) |
| BC | SCZ | 0.3603 (1.5807) | -0.0501 (0.4057) |
| BMI | HEIGHT | 0.3542 (1.5359) | -0.0615 (2.9427) |
| HR | WAIST | 0.3500 (1.5051) | 0.0601 (0.8069) |
| MI | RA | 0.3476 (1.4883) | NA (NA) |
| HR | SCZ | 0.3396 (1.4321) | 0.0623 (1.0736) |
| SCZ | WAIST | 0.3370 (1.4140) | -0.0044 (0.0574) |
| BMI | IBD | 0.3311 (1.3735) | -0.0424 (0.8890) |
| CD | MI | 0.3303 (1.3683) | NA (NA) |
| AF | UC | 0.3275 (1.3496) | -0.0244 (0.2665) |
| MI | WAIST | 0.3259 (1.3389) | NA (NA) |
| AF | IBD | 0.3241 (1.3270) | 0.0035 (0.0341) |
| BMI | CD | 0.3204 (1.3025) | 0.0089 (0.1156) |
| IBD | MI | 0.3115 (1.2443) | NA (NA) |
| BC | LOAD | -0.3000 (1.1721) | 0.1410 (0.4095) |
| NEU | WAIST | 0.2963 (1.1493) | 0.1167 (3.4213) |
| BC | HR | 0.2956 (1.1455) | -0.0508 (0.2254) |
| HDL | IBD | 0.2956 (1.1450) | -0.0274 (0.3364) |
| HDL | HEIGHT | 0.2921 (1.1240) | -0.0095 (0.1868) |
| BC | UC | -0.2857 (1.0859) | 0.3221 (2.5085) |
| CAD | WAIST | 0.2847 (1.0801) | 0.2011 (8.1403) |
| IBD | SCZ | 0.2804 (1.0545) | 0.1115 (3.3010) |
| CAD | T2D | -0.2705 (0.9982) | 0.3372 (9.7169) |
| HEIGHT | LDL | 0.2692 (0.9905) | -0.0770 (3.2569) |

*(continued)*

| | | Estimate (- log 10 *P*-value) | |
|---|---|---|---|
| Trait 1 | Trait 2 | Log 10 BF correlation | Genetic correlation |
| HDL | RA | 0.2680 (0.9841) | 0.0085 (0.1005) |
| IBD | NEU | 0.2674 (0.9804) | 0.0633 (0.8834) |
| MI | UC | 0.2648 (0.9661) | NA (NA) |
| HR | LOAD | -0.2619 (0.9500) | 0.0794 (0.4073) |
| AF | NEU | 0.2586 (0.9320) | 0.0152 (0.1957) |
| HR | UC | -0.2579 (0.9284) | 0.1382 (1.5419) |
| HR | IBD | -0.2560 (0.9178) | 0.0722 (0.7780) |
| AF | CD | 0.2526 (0.8997) | 0.0165 (0.1688) |
| AF | LDL | 0.2501 (0.8865) | -0.0171 (0.2734) |
| CD | NEU | 0.2468 (0.8687) | -0.0002 (0.0013) |
| AF | HDL | 0.2466 (0.8680) | -0.0949 (3.7918) |
| CD | SCZ | 0.2461 (0.8653) | 0.0924 (2.2076) |
| RA | WAIST | 0.2434 (0.8511) | -0.0367 (0.4518) |
| CAD | BC | 0.2379 (0.8228) | 0.0699 (0.4598) |
| CAD | NEU | 0.2361 (0.8139) | 0.0782 (1.5651) |
| CAD | RA | 0.2355 (0.8107) | 0.0779 (1.4365) |
| MI | T2D | -0.2351 (0.8089) | NA (NA) |
| BC | NEU | 0.2308 (0.7868) | 0.0186 (0.1021) |
| BMI | BC | 0.2249 (0.7578) | -0.0182 (0.1336) |
| HEIGHT | NEU | -0.2242 (0.7544) | -0.0524 (1.5010) |
| BC | LDL | -0.2177 (0.7230) | 0.0712 (0.6930) |
| MI | NEU | 0.2138 (0.7046) | NA (NA) |
| LOAD | MI | 0.2129 (0.7003) | NA (NA) |
| CD | HR | -0.2046 (0.6620) | 0.0069 (0.0481) |
| AF | WAIST | 0.2024 (0.6517) | -0.0459 (0.9825) |
| LDL | SCZ | 0.2008 (0.6445) | -0.0170 (0.3287) |
| HEIGHT | LOAD | 0.1980 (0.6317) | -0.1321 (1.9932) |
| HDL | SCZ | 0.1973 (0.6289) | 0.0336 (0.8680) |
| CD | WAIST | 0.1951 (0.6190) | -0.0349 (0.3788) |
| BC | HDL | -0.1847 (0.5735) | 0.0398 (0.3261) |
| HDL | HR | 0.1833 (0.5674) | -0.0277 (0.3379) |
| HR | T2D | -0.1819 (0.5615) | 0.0508 (0.4013) |
| HDL | T2D | -0.1722 (0.5211) | -0.3058 (12.5307) |
| CAD | LDL | 0.1717 (0.5191) | 0.2115 (7.1408) |
| LDL | WAIST | 0.1622 (0.4805) | 0.1095 (3.5518) |
| LDL | T2D | -0.1611 (0.4764) | 0.0505 (0.6992) |
| BC | WAIST | 0.1553 (0.4536) | -0.0291 (0.1636) |
| LOAD | T2D | -0.1488 (0.4289) | 0.0509 (0.1909) |
| UC | WAIST | 0.1412 (0.4003) | -0.0487 (0.5153) |
| HEIGHT | WAIST | -0.1389 (0.3919) | -0.0209 (0.3626) |
| HDL | UC | 0.1318 (0.3662) | 0.0264 (0.2749) |
| SCZ | T2D | -0.1309 (0.3631) | -0.0092 (0.0871) |

*(continued)*

| | | Estimate (- log 10 $P$-value) | |
|---|---|---|---|
| Trait 1 | Trait 2 | Log 10 BF correlation | Genetic correlation |
| AF | LOAD | 0.1305 (0.3618) | -0.0636 (0.5952) |
| HEIGHT | T2D | -0.1298 (0.3591) | -0.0100 (0.1005) |
| HR | NEU | 0.1289 (0.3560) | 0.0577 (0.8406) |
| IBD | WAIST | 0.1254 (0.3438) | -0.0418 (0.5297) |
| HEIGHT | IBD | 0.1239 (0.3386) | 0.0690 (1.3587) |
| BMI | LOAD | 0.1200 (0.3251) | -0.0294 (0.2396) |
| AF | BC | 0.1062 (0.2793) | -0.0153 (0.0882) |
| BC | MI | 0.0984 (0.2543) | NA (NA) |
| HEIGHT | RA | 0.0973 (0.2509) | 0.0477 (1.0714) |
| CD | HEIGHT | 0.0949 (0.2434) | 0.0494 (0.8913) |
| HEIGHT | SCZ | 0.0869 (0.2190) | 0.0055 (0.1102) |
| BMI | T2D | -0.0833 (0.2082) | 0.3267 (15.5429) |
| T2D | WAIST | 0.0793 (0.1965) | 0.2340 (5.6054) |
| CAD | LOAD | -0.0752 (0.1846) | -0.0374 (0.2298) |
| AF | T2D | 0.0708 (0.1722) | 0.1023 (1.7296) |
| BC | T2D | -0.0689 (0.1669) | -0.2116 (1.2632) |
| NEU | T2D | -0.0648 (0.1554) | -0.0468 (0.4010) |
| HEIGHT | UC | 0.0626 (0.1495) | 0.0579 (0.8229) |
| BC | RA | -0.0485 (0.1121) | 0.0442 (0.2305) |
| HR | LDL | -0.0477 (0.1101) | 0.0849 (1.6197) |
| HR | RA | -0.0457 (0.1051) | 0.0373 (0.3053) |
| HDL | NEU | 0.0456 (0.1048) | -0.0061 (0.0888) |
| T2D | UC | 0.0447 (0.1024) | 0.0520 (0.3240) |
| RA | T2D | 0.0391 (0.0885) | -0.0477 (0.4284) |
| LOAD | NEU | -0.0390 (0.0881) | 0.0789 (0.4814) |
| IBD | T2D | 0.0251 (0.0549) | -0.0021 (0.0144) |
| HDL | WAIST | -0.0236 (0.0515) | -0.2476 (13.2806) |
| LOAD | SCZ | 0.0212 (0.0460) | 0.0336 (0.2694) |
| LDL | NEU | 0.0210 (0.0455) | 0.0088 (0.1071) |
| CD | T2D | -0.0159 (0.0341) | -0.0162 (0.1260) |
| CAD | CD | 0.0112 (0.0238) | 0.0497 (0.5809) |
| BC | HEIGHT | 0.0099 (0.0209) | 0.0051 (0.0354) |
| CAD | IBD | 0.0055 (0.0116) | 0.0435 (0.4331) |
| CAD | UC | 0.0050 (0.0103) | 0.0090 (0.0617) |
| LOAD | WAIST | 0.0001 (0.0002) | 0.0254 (0.1522) |

#### Supplementary Table 8

For each trait or trait-network pair, we compute the proportion of genes with higher enrichment $P_1$ estimates ($P_1^{\mathsf{bma}}$ or $P_1^{\mathsf{net}}$) than reference estimates ($P_1^{\mathsf{base}}$ or $P_1^{\mathsf{near}}$), among genes with reference $P_1$ estimates higher than a given cutoff.

Panel **a** Median proportion of genes with $P_1^{\mathsf{bma}}$ higher than reference estimates ($P_1^{\mathsf{base}}$ or $P_1^{\mathsf{near}}$), among genes with reference estimates higher than a given cutoff. Medians shown below are across 16 traits that have multiple networks more enriched than the near-gene enrichment control (**Supplementary Table 3**). **Figure 5(e)** is based on this panel.

| | All genes | | Only TFs | | Only TGs | |
|---|---|---|---|---|---|---|
| Cutoff | $P_1^{\mathsf{base}}$ | $P_1^{\mathsf{near}}$ | $P_1^{\mathsf{base}}$ | $P_1^{\mathsf{near}}$ | $P_1^{\mathsf{base}}$ | $P_1^{\mathsf{near}}$ |
| 0 | 0.9828 | 0.9401 | 0.9556 | 0.8829 | 0.9842 | 0.9423 |
| 0.1 | 0.9531 | 0.8811 | 0.9837 | 0.8593 | 0.9499 | 0.8920 |
| 0.2 | 0.9180 | 0.8590 | 0.9514 | 0.8333 | 0.9166 | 0.8707 |
| 0.3 | 0.8856 | 0.8342 | 0.9564 | 0.8000 | 0.8898 | 0.8360 |
| 0.4 | 0.8754 | 0.8061 | 0.9667 | 0.7500 | 0.8784 | 0.8084 |
| 0.5 | 0.8695 | 0.7871 | 0.9444 | 0.7321 | 0.8771 | 0.7979 |
| 0.6 | 0.8499 | 0.7815 | 0.9348 | 0.7321 | 0.8607 | 0.7872 |
| 0.7 | 0.8647 | 0.7795 | 0.9583 | 0.7321 | 0.8624 | 0.7831 |
| 0.8 | 0.8399 | 0.7578 | 1.0000 | 0.6667 | 0.8521 | 0.7657 |
| 0.9 | 0.8167 | 0.7443 | 1.0000 | 0.6870 | 0.8259 | 0.7245 |

Panel **b** Median proportion of genes with $P_1^{\mathsf{net}}$ higher than reference estimates ($P_1^{\mathsf{base}}$ or $P_1^{\mathsf{near}}$), among genes with reference estimates higher than a given cutoff. Medians shown below are across 512 network-trait pairs passing the near-gene enrichment control (**Supplementary Table 3**).

| | All genes | | Only TFs | | Only TGs | |
|---|---|---|---|---|---|---|
| Cutoff | $P_1^{\mathsf{base}}$ | $P_1^{\mathsf{near}}$ | $P_1^{\mathsf{base}}$ | $P_1^{\mathsf{near}}$ | $P_1^{\mathsf{base}}$ | $P_1^{\mathsf{near}}$ |
| 0 | 0.9637 | 0.9046 | 0.8926 | 0.7058 | 0.9730 | 0.9325 |
| 0.1 | 0.9203 | 0.8613 | 0.8549 | 0.6667 | 0.9340 | 0.8898 |
| 0.2 | 0.9081 | 0.8494 | 0.8571 | 0.6667 | 0.9114 | 0.8696 |
| 0.3 | 0.8623 | 0.8276 | 0.8313 | 0.6667 | 0.8745 | 0.8537 |
| 0.4 | 0.8622 | 0.7971 | 0.8000 | 0.6000 | 0.8750 | 0.8225 |
| 0.5 | 0.8756 | 0.7891 | 0.7656 | 0.6079 | 0.8827 | 0.8053 |
| 0.6 | 0.8707 | 0.7895 | 0.7895 | 0.6154 | 0.8757 | 0.8000 |
| 0.7 | 0.8750 | 0.7906 | 0.7895 | 0.6206 | 0.8788 | 0.8077 |
| 0.8 | 0.8462 | 0.7679 | 0.7471 | 0.5862 | 0.8583 | 0.7863 |
| 0.9 | 0.8333 | 0.7500 | 0.7230 | 0.5000 | 0.8468 | 0.7619 |

#### Supplementary Table 9

For each trait we compute the correlation between RSS-NET enrichment statistic (log10 BF) and the percentage of network genes with $P_1^{\text{net}}$ higher than the reference estimates ($P_1^{\text{base}}$ or $P_1^{\text{near}}$), across all networks. Here we only consider 12 traits where all 38 networks show stronger enrichment than the near-gene control (**Supplementary Table 3**). Rows are ranked by Pearson $P$-values. The Bonferroni cutoff is $0.05/12 = 4.2 \times 10^{-3}$. Trait abbreviations are defined in **Supplementary Table 2**.

Panel **a** The reference $P_1$ is $P_1^{\text{base}}$.

| | Pearson $R$ (- log 10 $P$-value) | | |
|---|---|---|---|
| Trait | All genes | Only TFs | Only TGs |
| LDL | 0.8992 (13.7584) | 0.9419 (17.9148) | 0.8947 (13.4362) |
| HR | 0.8166 (9.3903) | 0.7301 (6.7046) | 0.8090 (9.0992) |
| BMI | 0.6810 (5.5959) | 0.6628 (5.2378) | 0.7271 (6.6306) |
| AF | 0.6687 (5.3518) | 0.7041 (6.0910) | 0.6492 (4.9850) |
| MI | 0.5851 (3.9420) | 0.5452 (3.3972) | 0.6125 (4.3594) |
| SCZ | 0.5722 (3.7584) | 0.6440 (4.8916) | 0.6173 (4.4368) |
| BC | 0.5579 (3.5637) | 0.6122 (4.3543) | 0.5412 (3.3468) |
| RA | 0.3848 (1.7679) | 0.3427 (1.4540) | 0.4013 (1.9022) |
| WAIST | 0.3672 (1.6316) | 0.4298 (2.1496) | 0.3946 (1.8468) |
| HDL | 0.2629 (0.9555) | 0.4662 (2.4981) | 0.2658 (0.9714) |
| CAD | 0.1948 (0.6175) | 0.0261 (0.0574) | 0.3409 (1.4413) |
| NEU | 0.1461 (0.4184) | 0.0523 (0.1219) | 0.2634 (0.9581) |

Panel **b** The reference $P_1$ is $P_1^{\text{near}}$. This panel is reported in the main text.

| | Pearson $R$ (- log 10 $P$-value) | | |
|---|---|---|---|
| Trait | All genes | Only TFs | Only TGs |
| LDL | 0.9100 (14.6053) | 0.9432 (18.0902) | 0.9086 (14.4901) |
| HR | 0.8392 (10.3326) | 0.8105 (9.1548) | 0.8323 (10.0295) |
| BMI | 0.7247 (6.5719) | 0.7711 (7.8321) | 0.7873 (8.3440) |
| MI | 0.7079 (6.1755) | 0.7797 (8.0986) | 0.7372 (6.8854) |
| AF | 0.7069 (6.1523) | 0.7894 (8.4116) | 0.6956 (5.9022) |
| NEU | 0.5618 (3.6150) | 0.6441 (4.8921) | 0.6092 (4.3072) |
| CAD | 0.5268 (3.1675) | 0.5975 (4.1260) | 0.6206 (4.4905) |
| SCZ | 0.5218 (3.1084) | 0.6433 (4.8791) | 0.5812 (3.8855) |
| BC | 0.5124 (2.9978) | 0.5914 (4.0355) | 0.4724 (2.5613) |
| RA | 0.5110 (2.9809) | 0.5191 (3.0761) | 0.5098 (2.9673) |
| WAIST | 0.2877 (1.0977) | 0.4315 (2.1654) | 0.3785 (1.7184) |
| HDL | 0.2831 (1.0705) | 0.4966 (2.8189) | 0.3151 (1.2679) |

#### Supplementary Table 10

Here we quantify overlap between RSS-NET prioritized genes ($P_1^{\text{bma}} \geq 0.9$) and genes implicated in the GWAS Catalog (Buniello et al. 2019), for each of 16 traits that pass the near-gene enrichment control (**Supplementary Table 3**). The "Same GWAS" column reports the overlap between RSS-NET prioritized genes and genes implicated in the same GWAS. The "Later GWAS" column reports the overlap between RSS-NET prioritized genes and genes implicated in a later GWAS of the same trait with increased sample size. The "No GWAS" column reports the number of RSS-NET prioritized genes that were not reported in either the same or the later GWAS at the time of analysis. For example, there are 38 genome-wide significant genes in GWAS Catalog for the same GWAS of HDL (Teslovich et al. 2010) that are analyzed by RSS-NET, and 32 of them have $P_1^{\text{bma}} \geq 0.9$; there are 20 new genome-wide significant genes in GWAS Catalog for a later GWAS of HDL (Global Lipids Genetics Consortium 2013), and 3 of them have $P_1^{\text{bma}} \geq 0.9$ based on RSS-NET analysis of previous GWAS (Teslovich et al. 2010); there are 142 genes with $P_1^{\text{bma}} \geq 0.9$ that are not reported in either the same or the later GWAS of HDL. All $P$-values are generated by R built-in function `fisher.test`. Rows are sorted by $P$-values in the "Same GWAS" column, then $P$-values in the "Later GWAS" column. Trait abbreviations are defined in **Supplementary Table 2**.

It is important to note that some published GWAS hits are not necessarily genome-wide significant in the corresponding summary data files. For example, rs34856868 shows $P = 9.80 \times 10^{-9}$ for association with IBD in Table 2 of Liu et al. (2015); however, the $P$-value of the same SNP is 0.27 in the corresponding summary data file (`EUR.IBD.gwas.assoc.gz`). For this SNP, the result in Table 2 of Liu et al. (2015) was indeed obtained from a combined analysis of data on both GWAS and Immunochip arrays, whereas the result in the summary data file was only based on GWAS arrays. Because of this type of potential discrepancy between summary data files and corresponding publications, the overlap fractions shown in the "Same GWAS" column are not very high for certain traits.

| | Fraction (- log 10 $P$-value) | | |
|---|---|---|---|
| Trait | Same GWAS | Later GWAS | No GWAS |
| RA | 44 / 61 (63.75) | 4 / 7 (5.79) | 190 |
| HDL | 32 / 38 (54.66) | 3 / 20 (2.87) | 142 |
| LDL | 26 / 27 (48.62) | 1 / 18 (0.75) | 141 |
| SCZ | 47 / 58 (45.93) | 138 / 330 (82.51) | 638 |
| AF | 16 / 18 (36.05) | 10 / 83 (12.79) | 35 |
| CAD | 18 / 20 (35.64) | 21 / 140 (21.75) | 72 |
| IBD | 33 / 80 (35.50) | 4 / 12 (4.60) | 101 |
| WAIST | 16 / 33 (29.73) | 5 / 233 (3.11) | 36 |
| HR | 10 / 15 (22.08) | 2 / 43 (2.33) | 29 |
| BC | 8 / 9 (21.03) | 4 / 107 (4.82) | 15 |
| BMI | 13 / 59 (19.61) | 4 / 461 (1.39) | 33 |
| MI | 8 / 9 (17.45) | NA | 61 |
| T2D | 6 / 16 (12.26) | 5 / 110 (6.29) | 8 |
| LOAD | 6 / 18 (11.12) | 11 / 29 (22.14) | 18 |
| UC | 6 / 17 (7.92) | 2 / 5 (3.02) | 126 |
| NEU | 3 / 5 (6.58) | 9 / 122 (10.40) | 28 |

#### Supplementary Table 11

Examples of RSS-NET highlighted genes that were reported in GWAS of the same data. The "mouse trait" column is based on the Mouse Genome Informatics (Bult et al. 2019). The "therapeutic/clinical evidence" column is based on the Online Mendelian Inheritance in Man (Amberger et al. 2019) and Therapeutic Target Database (Wang et al. 2020). Click blue links to view details online. Drugs are highlighted in yellow. CS: cardiovascular system; Metab.: metabolism; NS: nervous system.

| Trait | Gene (Role) | $P_1^{\text{base}}$ | $P_1^{\text{near}}$ | $P_1^{\text{bma}}$ | $P_1^{\text{net}}$ (Network, BF) | Mouse trait | Therapeutic/clinical evidence |
|---|---|---|---|---|---|---|---|
| BMI | *ADCY3* (TG) | 1.00 | 1.00 | 1.00 | 1.00 (Ileum, $4.99 \times 10^{12}$) | Growth | Severe obesity |
| | *NPC1* (TG) | 0.72 | 0.93 | 0.97 | 0.97 (Colon, $5.49 \times 10^{12}$) | Growth | Niemann-Pick disease |
| | *RPL27A* (TG) | 0.66 | 0.70 | 0.86 | 0.90 (Pancreas, $2.07 \times 10^{13}$) | Growth | |
| WAIST | *PIGC* (TG) | 1.00 | 1.00 | 1.00 | 1.00 (Esophagus, $6.78 \times 10^{239}$) | | GPIBD16 |
| | *TBX15* (TF) | 1.00 | 1.00 | 1.00 | 1.00 (Pancreas, $2.72 \times 10^{212}$) | Growth | Cousin syndrome |
| | *PPARG* (TF) | 0.94 | 0.94 | 0.96 | 0.96 (Esophagus, $6.78 \times 10^{239}$) | Metab. | Insulin resistance, Obesity |
| BC | *FGFR2* (TG) | 1.00 | 1.00 | 1.00 | 1.00 (Lung, $7.14 \times 10^{7}$) | Growth | Pfeiffer syndrome |
| | *PTHLH* (TG) | 1.00 | 1.00 | 1.00 | 1.00 (Aorta, $8.27 \times 10^{8}$) | Growth | Brachydactyly E2 |
| | *MKL1* (TG) | 0.80 | 0.80 | 0.91 | 0.92 (Spleen, $6.19 \times 10^{7}$) | | AMKL |
| RA | *CD40* (TG) | 1.00 | 1.00 | 1.00 | 1.00 (B cell, $3.31 \times 10^{57}$) | Immune | HIGM3 |
| | *CTLA4* (TG) | 1.00 | 1.00 | 1.00 | 1.00 (CD8, $6.96 \times 10^{56}$) | Immune | ALPS5 |
| | *ICOS* (TG) | 1.00 | 1.00 | 1.00 | 1.00 (CD8, $6.96 \times 10^{56}$) | Immune | Immunodeficiency CV1 |
| | *IL2RA* (TG) | 1.00 | 1.00 | 1.00 | 1.00 (NK cell, $2.95 \times 10^{60}$) | Immune | Immunodeficiency 41 |
| | *IL6R* (TG) | 1.00 | 1.00 | 1.00 | 1.00 (Monocyte, $8.91 \times 10^{53}$) | Immune | |
| | *RASGRP1* (TG) | 1.00 | 1.00 | 1.00 | 1.00 (NK cell, $2.95 \times 10^{60}$) | Immune | Immunodeficiency 64 |
| | *STAT4* (TF) | 1.00 | 1.00 | 1.00 | 1.00 (NK cell, $2.95 \times 10^{60}$) | Immune | |
| | *TYK2* (TG) | 1.00 | 1.00 | 1.00 | 1.00 (NK cell, $2.95 \times 10^{60}$) | Immune | Immunodeficiency 35 |
| IBD | *ATG16L1* (TG) | 1.00 | 1.00 | 1.00 | 1.00 (NK cell, $5.07 \times 10^{35}$) | Immune | |
| | *BACH2* (TF) | 1.00 | 1.00 | 1.00 | 1.00 (NK cell, $5.07 \times 10^{35}$) | Immune | Immunodeficiency 60 |
| | *FCGR2A* (TG) | 1.00 | 1.00 | 1.00 | 1.00 (Monocyte, $6.28 \times 10^{31}$) | Immune | Lupus nephritis |
| | *JAK2* (TG) | 1.00 | 1.00 | 1.00 | 1.00 (Monocyte, $6.28 \times 10^{31}$) | Immune | Baricitinib |
| | *STAT3* (TF) | 1.00 | 1.00 | 1.00 | 1.00 (NK cell, $5.07 \times 10^{35}$) | Immune | ADMIO1 |
| | *TYK2* (TG) | 0.96 | 0.98 | 0.98 | 0.98 (NK cell, $5.07 \times 10^{35}$) | Immune | Immunodeficiency 35 |
| | *STAT4* (TF) | 0.77 | 0.82 | 0.91 | 0.91 (NK cell, $5.07 \times 10^{35}$) | Immune | Susceptibility to SLE |
| HDL | *ABCA1* (TG) | 1.00 | 1.00 | 1.00 | 1.00 (Liver, $2.81 \times 10^{21}$) | Metab. | Tangier disease, Probucol |
| | *APOB* (TG) | 1.00 | 1.00 | 1.00 | 1.00 (Liver, $2.81 \times 10^{21}$) | Metab. | FCHL2, FHBL1, Mipomersen |
| | *GALNT2* (TG) | 1.00 | 1.00 | 1.00 | 1.00 (Liver, $2.81 \times 10^{21}$) | Metab. | |
| | *LIPG* (TG) | 1.00 | 1.00 | 1.00 | 1.00 (Liver, $2.81 \times 10^{21}$) | Metab. | GSK-264220A |
| | *LPL* (TG) | 1.00 | 1.00 | 1.00 | 1.00 (Omentum, $8.70 \times 10^{13}$) | Metab. | Clofibrate Gemfibrozil |
| | *SCARB1* (TG) | 1.00 | 1.00 | 1.00 | 1.00 (Liver, $2.81 \times 10^{21}$) | Metab. | |
| LDL | *APOB* (TG) | 1.00 | 1.00 | 1.00 | 1.00 (Liver, $7.66 \times 10^{27}$) | Metab. | FCHL2, FHBL1, Mipomersen |
| | *HMGCR* (TG) | 1.00 | 1.00 | 1.00 | 1.00 (CD8, $5.86 \times 10^{28}$) | Metab. | |
| | *HNF1A* (TF) | 1.00 | 1.00 | 1.00 | 1.00 (CD8, $5.86 \times 10^{28}$) | Metab. | MODY3, Hepatic adenomas |
| | *LDLR* (TG) | 1.00 | 1.00 | 1.00 | 1.00 (Pancreas, $3.04 \times 10^{28}$) | Metab. | FHCL1 |
| | *NPC1L1* (TG) | 1.00 | 1.00 | 1.00 | 1.00 (Liver, $7.66 \times 10^{27}$) | Metab. | Response to ezetimibe |
| T2D | *TCF7L2* (TF) | 1.00 | 1.00 | 1.00 | 1.00 (NK cell, $1.49 \times 10^{77}$) | Metab. | Susceptibility to T2D |
| | *IGF2BP2* (TG) | 0.99 | 1.00 | 1.00 | 1.00 (Ileum, $4.52 \times 10^{62}$) | Metab. | Susceptibility to T2D |
| | *PPARG* (TF) | 0.98 | 1.00 | 1.00 | 1.00 (Prostate, $5.64 \times 10^{66}$) | Metab. | Insulin resistance, Obesity |
| | *PROX1* (TG) | 0.32 | 0.92 | 0.92 | 0.92 (Thyroid, $3.61 \times 10^{61}$) | CS, Metab. | |
| HR | *MYH6* (TG) | 1.00 | 1.00 | 1.00 | 1.00 (Heart, $2.12 \times 10^{7}$) | CS, Muscle | Cardiomyopathy D1EE, H14 |
| | *GJA1* (TG) | 1.00 | 1.00 | 1.00 | 1.00 (Uterus, $1.51 \times 10^{6}$) | CS, Muscle | AVSD3, HLHS1 |
| | *EPHB4* (TG) | 1.00 | 1.00 | 1.00 | 1.00 (Aorta, $2.43 \times 10^{7}$) | CS, Muscle | CMAVM2, LMPHM7 |
| | *TTN* (TG) | 1.00 | 1.00 | 1.00 | 1.00 (Aorta, $2.43 \times 10^{7}$) | CS, Muscle | Cardiomyopathy D1G, H9 |
| | *GNB4* (TG) | 0.55 | 0.64 | 0.91 | 0.91 (Aorta, $2.43 \times 10^{7}$) | CS | CMTDIF |
| CAD | *SMAD3* (TF) | 0.99 | 1.00 | 1.00 | 1.00 (Adipose, $1.67 \times 10^{29}$) | CS, Blood | Loeys-Dietz syndrome 3 |
| | *APOB* (TG) | 0.94 | 0.98 | 0.99 | 0.99 (Liver, $2.99 \times 10^{25}$) | CS, Metab. | FCHL2, FHBL1, Mipomersen |
| | *ATP2B1* (TG) | 0.91 | 0.96 | 0.98 | 0.99 (Heart, $1.93 \times 10^{28}$) | CS, Muscle | |
| | *MRAS* (TG) | 0.89 | 0.96 | 0.98 | 0.98 (Adipose, $1.67 \times 10^{29}$) | Immune | Noonan syndrome 11 |
| | *APOE* (TG) | 0.80 | 0.92 | 0.96 | 0.96 (Adipose, $1.67 \times 10^{29}$) | CS, Metab. | Hyperlipoproteinemia 3 |
| | *ABHD2* (TG) | 0.81 | 0.92 | 0.95 | 0.95 (Adipose, $1.67 \times 10^{29}$) | CS | |
| | *IL6R* (TG) | 0.77 | 0.91 | 0.93 | 0.94 (Heart, $1.93 \times 10^{28}$) | Immune | Serum level of IL6 |
| | *FURIN* (TG) | 0.69 | 0.86 | 0.93 | 0.95 (Heart, $1.93 \times 10^{28}$) | CS | |
| AF | *TBX5* (TF) | 1.00 | 1.00 | 1.00 | 1.00 (Heart, $2.15 \times 10^{14}$) | CS, Muscle | Holt-Oram syndrome |
| | *PITX2* (TF) | 1.00 | 1.00 | 1.00 | 1.00 (Muscle, $8.55 \times 10^{14}$) | CS, Muscle | |
| | *CAV1* (TG) | 1.00 | 1.00 | 1.00 | 1.00 (Muscle, $8.55 \times 10^{14}$) | CS, Muscle | PP hypertension 3 |
| | *SH3PXD2A* (TG) | 1.00 | 1.00 | 1.00 | 1.00 (Muscle, $8.55 \times 10^{14}$) | CS, Muscle | |

*(continued)*

| Trait | Gene (Role) | $P_1^{base}$ | $P_1^{near}$ | $P_1^{bma}$ | $P_1^{net}$ (Network, BF) | Mouse trait | Therapeutic/clinical evidence |
|---|---|---|---|---|---|---|---|
| | *ASAH1* (TG) | 1.00 | 1.00 | 1.00 | 1.00 (Muscle, $8.55 \times 10^{14}$) | Metab. | Farber lipogranulomatosis |
| | *TTN* (TG) | 0.95 | 0.97 | 0.99 | 0.99 (Muscle, $8.55 \times 10^{14}$) | CS, Muscle | Cardiomyopathy D1G, H9 |
| LOAD | *APOE* (TG) | 1.00 | 1.00 | 1.00 | 1.00 (Liver, $1.09 \times 10^{20}$) | Liver, NS | Hyperlipoproteinemia 3 |
| | *BIN1* (TG) | 1.00 | 1.00 | 1.00 | 1.00 (CD8, $8.31 \times 10^{26}$) | Muscle | Centronuclear myopathy 2 |
| | *CLU* (TG) | 1.00 | 1.00 | 1.00 | 1.00 (Aorta, $3.24 \times 10^{23}$) | CS, Muscle | |
| | *EPHA1* (TG) | 0.99 | 1.00 | 1.00 | 1.00 (CD8, $8.31 \times 10^{26}$) | Immune | |
| | *CD2AP* (TG) | 0.70 | 0.95 | 0.98 | 0.98 (Pancreas, $3.53 \times 10^{20}$) | Metab. | FSGS3 |
| SCZ | *CACNA1C* (TG) | 1.00 | 1.00 | 1.00 | 1.00 (Spleen, $1.44 \times 10^{141}$) | Immune, NS | Timothy syndrome |
| | *TCF4* (TF) | 1.00 | 1.00 | 1.00 | 1.00 (Colon, $1.20 \times 10^{144}$) | Immune, NS | Pitt-Hopkins syndrome |
| | *DPYD* (TG) | 1.00 | 1.00 | 1.00 | 1.00 (Spleen, $1.44 \times 10^{141}$) | Neuron | DPD deficiency |
| NEU | *LINGO1* (TG) | 0.99 | 1.00 | 1.00 | 1.00 (Putamen, $2.12 \times 10^{19}$) | NS | Mental retardation |
| | *SBF2* (TG) | 0.94 | 0.97 | 0.99 | 0.99 (Lung, $1.42 \times 10^{18}$) | Neuron, NS | CMT4B2 |
| | *PAFAH1B1* (TG) | 0.77 | 0.93 | 1.00 | 1.00 (Cerebellum, $4.85 \times 10^{18}$) | Neuron, NS | Lissencephaly |

#### Supplementary Table 12

For each of 14 GWAS traits, we quantify overlap between RSS-NET prioritized genes ($P_1^{bma} \geq 0.9$) and genes implicated in 27 categories of knockout mouse phenotypes (Bult et al. 2019). These 14 traits are obtained by removing 2 disease subtypes (MI for CAD, UC for IBD) from the 16 traits that pass the near-gene control (**Supplementary Table 3**). Analysis details are provided in **Supplementary Figure 15**. Within a trait, rows are sorted by $P$-values and then odds ratios from the analysis of all genes (the 4th column). Trait abbreviations are defined in **Supplementary Table 2**. We use the 1st row "41 / 184 (19.17)" to explain the meaning of "GWAS overlap" (the 3rd column): 184 genes are implicated by GWAS of AF (Buniello et al. 2019) at the time of analysis, 41 of them belong to the "Muscle" category, and the $-\log 10$ Fisher exact test $P$-value is 19.17.

| | | | Odds ratio (- log 10 $P$-value) | |
|---|---|---|---|---|
| Trait | Category | GWAS overlap | All genes | Non-GWAS genes |
| AF | Muscle | 41 / 184 (19.17) | 3.88 (3.36) | 2.42 (1.12) |
| AF | Skeleton | 29 / 184 (7.65) | 3.17 (2.74) | 2.10 (0.88) |
| AF | Cardiovascular system | 44 / 184 (12.68) | 2.92 (2.65) | 1.92 (0.79) |
| AF | Neoplasm | 10 / 184 (4.00) | 4.11 (2.30) | 2.20 (0.72) |
| AF | Respiratory system | 26 / 184 (10.80) | 3.25 (2.25) | 1.63 (0.48) |
| AF | Renal/urinary system | 10 / 184 (2.41) | 3.05 (1.96) | 1.69 (0.50) |
| AF | Growth/size/body region | 53 / 184 (10.57) | 2.35 (1.93) | 1.66 (0.65) |
| AF | Craniofacial | 20 / 184 (8.13) | 3.00 (1.79) | 2.34 (0.79) |
| AF | Nervous system | 36 / 184 (7.15) | 2.33 (1.79) | 1.62 (0.58) |
| AF | Behavior/neurological | 45 / 184 (9.40) | 2.25 (1.71) | 1.33 (0.30) |
| AF | Reproductive system | 29 / 184 (8.19) | 2.29 (1.40) | 1.04 (-0.00) |
| AF | Endocrine/exocrine gland | 27 / 184 (6.82) | 2.26 (1.34) | 1.65 (0.52) |
| AF | Liver/biliary system | 19 / 184 (6.73) | 2.57 (1.34) | 1.61 (0.30) |
| AF | Digestive/alimentary | 17 / 184 (5.52) | 2.52 (1.32) | 1.58 (0.30) |
| AF | Mortality/aging | 61 / 184 (10.74) | 1.92 (1.30) | 1.36 (0.26) |
| AF | Homeostasis/metabolism | 63 / 184 (11.65) | 1.90 (1.18) | 1.08 (-0.00) |
| AF | Embryo | 20 / 184 (5.16) | 2.08 (1.06) | 1.44 (0.25) |
| AF | Hematopoietic system | 27 / 184 (4.09) | 1.86 (1.06) | 1.55 (0.43) |
| AF | Integument | 23 / 184 (6.23) | 1.97 (0.84) | 1.37 (0.25) |
| AF | Immune system | 28 / 184 (4.32) | 1.73 (0.77) | 1.39 (0.31) |

*(continued)*

| Trait | Category | GWAS overlap | Odds ratio (- log 10 $P$-value) | |
|---|---|---|---|---|
| | | | All genes | Non-GWAS genes |
| AF | Limbs/digits/tail | 11 / 184 (3.51) | 2.09 (0.74) | 1.05 (-0.00) |
| AF | Vision/eye | 13 / 184 (2.26) | 1.86 (0.69) | 1.45 (0.25) |
| AF | Adipose tissue | 15 / 184 (5.13) | 1.83 (0.66) | 0.91 (-0.00) |
| AF | Taste/olfaction | 1 / 184 (0.56) | 3.64 (0.60) | 4.55 (0.67) |
| AF | Cellular | 48 / 184 (9.38) | 1.58 (0.59) | 1.06 (-0.00) |
| AF | Hearing/vestibular/ear | 8 / 184 (2.91) | 1.79 (0.38) | 0.00 (0.21) |
| AF | Pigmentation | 1 / 184 (-0.00) | 0.93 (-0.00) | 1.16 (0.23) |
| BC | Cardiovascular system | 39 / 364 (3.37) | 4.60 (2.80) | 3.68 (1.54) |
| BC | Skeleton | 41 / 364 (5.33) | 4.19 (2.20) | 3.35 (1.36) |
| BC | Renal/urinary system | 27 / 364 (5.66) | 5.07 (2.12) | 3.05 (0.88) |
| BC | Limbs/digits/tail | 21 / 364 (4.41) | 5.24 (1.96) | 2.52 (0.61) |
| BC | Neoplasm | 21 / 364 (6.37) | 5.85 (1.85) | 0.00 (-0.00) |
| BC | Integument | 37 / 364 (5.56) | 3.82 (1.77) | 1.97 (0.40) |
| BC | Adipose tissue | 28 / 364 (6.58) | 4.57 (1.75) | 2.20 (0.53) |
| BC | Mortality/aging | 83 / 364 (6.67) | 2.92 (1.61) | 2.25 (0.75) |
| BC | Respiratory system | 22 / 364 (3.19) | 4.05 (1.57) | 0.97 (-0.00) |
| BC | Muscle | 32 / 364 (6.26) | 3.44 (1.34) | 2.48 (0.72) |
| BC | Hearing/vestibular/ear | 8 / 364 (0.81) | 4.49 (1.27) | 3.60 (0.82) |
| BC | Vision/eye | 29 / 364 (3.50) | 2.90 (1.12) | 2.79 (0.92) |
| BC | Liver/biliary system | 22 / 364 (3.23) | 3.21 (1.11) | 1.92 (0.46) |
| BC | Immune system | 44 / 364 (2.75) | 2.47 (1.02) | 1.11 (-0.00) |
| BC | Growth/size/body region | 72 / 364 (6.83) | 2.29 (0.93) | 1.22 (0.13) |
| BC | Homeostasis/metabolism | 67 / 364 (3.96) | 2.16 (0.86) | 1.56 (0.26) |
| BC | Craniofacial | 21 / 364 (3.60) | 2.80 (0.84) | 0.00 (0.00) |
| BC | Nervous system | 45 / 364 (3.09) | 2.26 (0.83) | 1.16 (-0.00) |
| BC | Hematopoietic system | 45 / 364 (3.07) | 2.17 (0.80) | 0.75 (-0.00) |
| BC | Behavior/neurological | 60 / 364 (5.64) | 2.37 (0.80) | 1.42 (0.14) |
| BC | Cellular | 57 / 364 (3.97) | 2.11 (0.75) | 0.95 (0.00) |
| BC | Digestive/alimentary | 18 / 364 (1.81) | 2.36 (0.70) | 0.00 (0.23) |
| BC | Embryo | 28 / 364 (3.07) | 2.31 (0.61) | 0.69 (-0.00) |
| BC | Reproductive system | 38 / 364 (5.07) | 2.07 (0.57) | 1.24 (0.17) |
| BC | Endocrine/exocrine gland | 39 / 364 (4.84) | 1.88 (0.52) | 0.00 (0.47) |
| BC | Pigmentation | 7 / 364 (1.40) | 2.33 (0.42) | 0.00 (-0.00) |
| BC | Taste/olfaction | 4 / 364 (1.95) | 0.00 (-0.00) | 0.00 (-0.00) |
| BMI | Renal/urinary system | 65 / 889 (6.26) | 4.77 (4.16) | 3.95 (2.70) |
| BMI | Liver/biliary system | 79 / 889 (9.66) | 3.87 (3.05) | 2.62 (1.19) |
| BMI | Embryo | 114 / 889 (12.90) | 3.24 (2.60) | 2.68 (1.65) |
| BMI | Vision/eye | 105 / 889 (10.35) | 2.79 (2.01) | 2.16 (0.96) |
| BMI | Growth/size/body region | 270 / 889 (24.06) | 2.24 (1.76) | 1.42 (0.38) |
| BMI | Adipose tissue | 74 / 889 (10.18) | 2.93 (1.75) | 1.27 (0.14) |
| BMI | Reproductive system | 107 / 889 (7.57) | 2.50 (1.65) | 1.69 (0.53) |

*(continued)*

| Trait | Category | GWAS overlap | Odds ratio (- log 10 $P$-value) | |
|---|---|---|---|---|
| | | | All genes | Non-GWAS genes |
| BMI | Mortality/aging | 301 / 889 (21.97) | 2.00 (1.45) | 1.35 (0.35) |
| BMI | Taste/olfaction | 8 / 889 (1.84) | 7.36 (1.43) | 0.00 (-0.00) |
| BMI | Respiratory system | 85 / 889 (11.32) | 2.59 (1.35) | 1.88 (0.67) |
| BMI | Homeostasis/metabolism | 255 / 889 (13.97) | 1.90 (1.18) | 1.50 (0.47) |
| BMI | Endocrine/exocrine gland | 134 / 889 (13.01) | 2.07 (1.14) | 1.75 (0.66) |
| BMI | Nervous system | 220 / 889 (22.42) | 1.94 (1.09) | 1.50 (0.42) |
| BMI | Behavior/neurological | 238 / 889 (22.82) | 1.90 (1.04) | 1.10 (0.08) |
| BMI | Hematopoietic system | 178 / 889 (12.03) | 1.74 (0.77) | 1.44 (0.41) |
| BMI | Cardiovascular system | 154 / 889 (13.34) | 1.68 (0.70) | 1.24 (0.20) |
| BMI | Integument | 107 / 889 (9.73) | 1.75 (0.66) | 1.26 (0.23) |
| BMI | Cellular | 224 / 889 (15.69) | 1.58 (0.59) | 1.10 (0.08) |
| BMI | Pigmentation | 27 / 889 (3.92) | 1.87 (0.50) | 1.08 (0.21) |
| BMI | Immune system | 170 / 889 (10.05) | 1.48 (0.50) | 1.28 (0.18) |
| BMI | Limbs/digits/tail | 64 / 889 (8.95) | 1.67 (0.49) | 0.97 (-0.00) |
| BMI | Digestive/alimentary | 72 / 889 (7.07) | 1.57 (0.42) | 1.82 (0.65) |
| BMI | Hearing/vestibular/ear | 40 / 889 (4.98) | 1.79 (0.38) | 1.39 (0.18) |
| BMI | Craniofacial | 76 / 889 (11.06) | 1.49 (0.29) | 1.30 (0.14) |
| BMI | Skeleton | 140 / 889 (14.46) | 1.30 (0.20) | 0.86 (-0.00) |
| BMI | Muscle | 87 / 889 (9.77) | 1.10 (0.11) | 0.64 (0.12) |
| BMI | Neoplasm | 45 / 889 (6.29) | 0.58 (-0.00) | 0.68 (-0.00) |
| CAD | Cardiovascular system | 70 / 308 (18.88) | 2.45 (3.65) | 1.59 (0.99) |
| CAD | Liver/biliary system | 36 / 308 (13.08) | 2.71 (2.86) | 1.37 (0.39) |
| CAD | Mortality/aging | 101 / 308 (16.94) | 1.85 (2.24) | 1.14 (0.22) |
| CAD | Embryo | 43 / 308 (12.48) | 2.15 (2.07) | 1.41 (0.55) |
| CAD | Renal/urinary system | 26 / 308 (8.03) | 2.25 (1.93) | 1.12 (0.08) |
| CAD | Hematopoietic system | 61 / 308 (11.49) | 1.81 (1.70) | 1.11 (0.11) |
| CAD | Cellular | 68 / 308 (10.97) | 1.73 (1.57) | 1.14 (0.17) |
| CAD | Respiratory system | 33 / 308 (11.21) | 2.01 (1.47) | 0.76 (0.08) |
| CAD | Limbs/digits/tail | 22 / 308 (7.33) | 2.04 (1.32) | 1.18 (0.19) |
| CAD | Integument | 42 / 308 (11.43) | 1.84 (1.31) | 0.93 (-0.00) |
| CAD | Digestive/alimentary | 33 / 308 (11.13) | 1.95 (1.27) | 0.89 (-0.00) |
| CAD | Immune system | 54 / 308 (8.90) | 1.64 (1.19) | 0.99 (-0.00) |
| CAD | Neoplasm | 18 / 308 (6.99) | 2.07 (1.15) | 0.55 (0.24) |
| CAD | Growth/size/body region | 79 / 308 (13.82) | 1.53 (1.08) | 0.77 (0.34) |
| CAD | Endocrine/exocrine gland | 40 / 308 (8.64) | 1.66 (1.07) | 0.88 (0.07) |
| CAD | Muscle | 40 / 308 (13.81) | 1.70 (1.00) | 0.91 (-0.00) |
| CAD | Reproductive system | 40 / 308 (9.30) | 1.65 (0.99) | 1.27 (0.31) |
| CAD | Nervous system | 51 / 308 (8.19) | 1.54 (0.99) | 1.03 (0.05) |
| CAD | Homeostasis/metabolism | 87 / 308 (13.23) | 1.45 (0.91) | 0.81 (0.31) |
| CAD | Vision/eye | 33 / 308 (7.91) | 1.54 (0.77) | 0.98 (-0.00) |
| CAD | Pigmentation | 7 / 308 (2.26) | 2.07 (0.74) | 0.88 (-0.00) |
| CAD | Craniofacial | 29 / 308 (10.32) | 1.65 (0.70) | 0.70 (0.19) |

*(continued)*

| Trait | Category | GWAS overlap | Odds ratio (- log 10 $P$-value) | |
|---|---|---|---|---|
| | | | All genes | Non-GWAS genes |
| CAD | Adipose tissue | 28 / 308 (9.67) | 1.45 (0.53) | 0.69 (0.19) |
| CAD | Skeleton | 47 / 308 (11.49) | 1.31 (0.38) | 0.78 (0.22) |
| CAD | Taste/olfaction | 1 / 308 (0.38) | 1.60 (0.32) | 0.00 (-0.00) |
| CAD | Behavior/neurological | 44 / 308 (5.13) | 1.20 (0.31) | 0.72 (0.45) |
| CAD | Hearing/vestibular/ear | 7 / 308 (1.35) | 0.79 (0.00) | 0.28 (0.60) |
| HDL | Liver/biliary system | 12 / 104 (3.13) | 2.23 (2.89) | 1.74 (1.38) |
| HDL | Digestive/alimentary | 13 / 104 (3.51) | 2.18 (2.83) | 1.89 (1.71) |
| HDL | Homeostasis/metabolism | 35 / 104 (3.83) | 1.72 (2.76) | 1.43 (1.17) |
| HDL | Embryo | 17 / 104 (3.88) | 1.77 (1.84) | 1.32 (0.49) |
| HDL | Cardiovascular system | 24 / 104 (4.49) | 1.64 (1.79) | 1.22 (0.40) |
| HDL | Immune system | 17 / 104 (1.33) | 1.51 (1.39) | 1.37 (0.82) |
| HDL | Hematopoietic system | 17 / 104 (1.33) | 1.49 (1.29) | 1.30 (0.65) |
| HDL | Respiratory system | 9 / 104 (1.82) | 1.65 (1.16) | 1.38 (0.59) |
| HDL | Mortality/aging | 31 / 104 (2.83) | 1.38 (1.08) | 1.11 (0.21) |
| HDL | Vision/eye | 13 / 104 (2.26) | 1.54 (1.06) | 1.25 (0.40) |
| HDL | Endocrine/exocrine gland | 10 / 104 (0.73) | 1.49 (1.04) | 1.34 (0.61) |
| HDL | Reproductive system | 10 / 104 (0.93) | 1.43 (0.89) | 1.18 (0.23) |
| HDL | Cellular | 20 / 104 (1.36) | 1.35 (0.86) | 1.14 (0.23) |
| HDL | Adipose tissue | 5 / 104 (0.63) | 1.49 (0.82) | 1.45 (0.63) |
| HDL | Growth/size/body region | 20 / 104 (1.33) | 1.30 (0.70) | 1.18 (0.34) |
| HDL | Renal/urinary system | 11 / 104 (2.86) | 1.38 (0.57) | 1.25 (0.30) |
| HDL | Limbs/digits/tail | 7 / 104 (1.60) | 1.38 (0.52) | 1.30 (0.33) |
| HDL | Muscle | 9 / 104 (1.46) | 1.26 (0.41) | 0.94 (-0.00) |
| HDL | Behavior/neurological | 13 / 104 (0.45) | 1.17 (0.34) | 1.11 (0.19) |
| HDL | Integument | 10 / 104 (1.19) | 1.17 (0.23) | 1.05 (0.05) |
| HDL | Hearing/vestibular/ear | 1 / 104 (-0.00) | 0.76 (0.17) | 0.67 (0.18) |
| HDL | Neoplasm | 3 / 104 (0.36) | 1.19 (0.17) | 1.15 (0.17) |
| HDL | Skeleton | 12 / 104 (1.27) | 1.09 (0.15) | 1.00 (-0.00) |
| HDL | Nervous system | 15 / 104 (1.01) | 1.08 (0.13) | 0.95 (0.04) |
| HDL | Pigmentation | 3 / 104 (0.92) | 0.71 (0.10) | 0.52 (0.23) |
| HDL | Craniofacial | 6 / 104 (0.89) | 0.95 (-0.00) | 0.95 (-0.00) |
| HDL | Taste/olfaction | 0 / 104 (-0.00) | 0.92 (-0.00) | 1.02 (0.00) |
| HR | Endocrine/exocrine gland | 13 / 123 (1.81) | 4.45 (2.73) | 3.78 (1.87) |
| HR | Muscle | 18 / 123 (5.89) | 4.73 (2.45) | 3.45 (1.34) |
| HR | Immune system | 17 / 123 (1.59) | 3.71 (2.34) | 3.40 (1.73) |
| HR | Reproductive system | 10 / 123 (1.17) | 4.01 (2.21) | 2.60 (0.82) |
| HR | Nervous system | 24 / 123 (3.59) | 3.61 (2.14) | 2.59 (1.05) |
| HR | Mortality/aging | 30 / 123 (3.00) | 3.22 (2.12) | 2.30 (0.85) |
| HR | Cardiovascular system | 28 / 123 (6.51) | 3.61 (2.09) | 3.07 (1.40) |
| HR | Vision/eye | 10 / 123 (1.55) | 3.99 (2.07) | 2.32 (0.61) |
| HR | Behavior/neurological | 24 / 123 (3.15) | 3.30 (2.02) | 2.07 (0.61) |
| HR | Cellular | 27 / 123 (3.42) | 3.17 (1.92) | 2.38 (0.95) |

*(continued)*

| Trait | Category | GWAS overlap | Odds ratio (- log 10 *P*-value) | |
|---|---|---|---|---|
| | | | All genes | Non-GWAS genes |
| HR | Respiratory system | 10 / 123 (2.45) | 4.17 (1.84) | 4.05 (1.57) |
| HR | Hematopoietic system | 12 / 123 (0.63) | 3.20 (1.84) | 2.80 (1.25) |
| HR | Liver/biliary system | 8 / 123 (1.67) | 4.13 (1.83) | 4.01 (1.55) |
| HR | Digestive/alimentary | 7 / 123 (1.12) | 4.05 (1.80) | 2.36 (0.70) |
| HR | Renal/urinary system | 5 / 123 (0.66) | 3.62 (1.46) | 3.38 (1.17) |
| HR | Homeostasis/metabolism | 29 / 123 (2.84) | 2.59 (1.34) | 1.95 (0.53) |
| HR | Limbs/digits/tail | 5 / 123 (0.76) | 3.59 (1.27) | 4.19 (1.42) |
| HR | Growth/size/body region | 22 / 123 (1.93) | 2.62 (1.24) | 2.29 (0.93) |
| HR | Hearing/vestibular/ear | 4 / 123 (1.01) | 3.85 (1.15) | 2.99 (0.73) |
| HR | Neoplasm | 2 / 123 (0.16) | 3.75 (1.13) | 4.38 (1.25) |
| HR | Adipose tissue | 3 / 123 (0.13) | 3.13 (1.12) | 2.74 (0.82) |
| HR | Integument | 16 / 123 (3.62) | 2.81 (1.04) | 2.19 (0.59) |
| HR | Skeleton | 7 / 123 (0.32) | 2.39 (0.92) | 2.79 (1.03) |
| HR | Embryo | 13 / 123 (2.39) | 2.47 (0.81) | 2.89 (1.11) |
| HR | Craniofacial | 5 / 123 (0.70) | 2.40 (0.73) | 1.87 (0.46) |
| HR | Pigmentation | 2 / 123 (0.54) | 2.00 (0.37) | 0.00 (-0.00) |
| HR | Taste/olfaction | 0 / 123 (-0.00) | 0.00 (0.00) | 0.00 (0.00) |
| IBD | Hematopoietic system | 100 / 374 (17.69) | 2.35 (4.48) | 1.86 (2.07) |
| IBD | Neoplasm | 23 / 374 (6.95) | 3.44 (4.41) | 2.70 (2.23) |
| IBD | Immune system | 104 / 374 (18.96) | 2.27 (4.14) | 1.76 (1.68) |
| IBD | Digestive/alimentary | 44 / 374 (12.09) | 2.85 (4.13) | 2.67 (3.08) |
| IBD | Homeostasis/metabolism | 104 / 374 (10.69) | 2.08 (3.47) | 1.74 (1.77) |
| IBD | Adipose tissue | 20 / 374 (2.80) | 2.74 (3.04) | 2.79 (2.87) |
| IBD | Endocrine/exocrine gland | 57 / 374 (10.02) | 2.20 (2.95) | 1.88 (1.70) |
| IBD | Cardiovascular system | 56 / 374 (7.29) | 2.03 (2.64) | 1.61 (1.15) |
| IBD | Nervous system | 48 / 374 (3.21) | 2.02 (2.61) | 1.92 (1.95) |
| IBD | Renal/urinary system | 26 / 374 (4.60) | 2.36 (2.37) | 1.63 (0.80) |
| IBD | Cellular | 91 / 374 (11.24) | 1.74 (1.91) | 1.41 (0.77) |
| IBD | Liver/biliary system | 31 / 374 (6.31) | 2.02 (1.74) | 2.04 (1.57) |
| IBD | Respiratory system | 26 / 374 (4.24) | 1.93 (1.56) | 1.67 (0.88) |
| IBD | Behavior/neurological | 53 / 374 (3.34) | 1.69 (1.53) | 1.30 (0.48) |
| IBD | Limbs/digits/tail | 19 / 374 (3.13) | 2.08 (1.53) | 1.60 (0.57) |
| IBD | Skeleton | 41 / 374 (4.75) | 1.74 (1.53) | 1.41 (0.55) |
| IBD | Reproductive system | 36 / 374 (3.72) | 1.77 (1.42) | 1.47 (0.61) |
| IBD | Mortality/aging | 97 / 374 (8.64) | 1.56 (1.41) | 1.37 (0.70) |
| IBD | Growth/size/body region | 77 / 374 (7.04) | 1.54 (1.26) | 1.32 (0.51) |
| IBD | Embryo | 37 / 374 (5.41) | 1.61 (1.11) | 1.40 (0.59) |
| IBD | Integument | 45 / 374 (7.69) | 1.57 (0.99) | 1.26 (0.30) |
| IBD | Vision/eye | 24 / 374 (1.66) | 1.55 (0.91) | 1.55 (0.76) |
| IBD | Muscle | 28 / 374 (4.15) | 1.61 (0.86) | 1.35 (0.44) |
| IBD | Craniofacial | 19 / 374 (2.55) | 1.53 (0.72) | 1.52 (0.52) |

*(continued)*

| Trait | Category | GWAS overlap | Odds ratio (- log 10 $P$-value) | |
|---|---|---|---|---|
| | | | All genes | Non-GWAS genes |
| IBD | Pigmentation | 8 / 374 (1.63) | 1.29 (0.26) | 1.16 (0.13) |
| IBD | Hearing/vestibular/ear | 6 / 374 (0.19) | 1.11 (0.10) | 1.31 (0.23) |
| IBD | Taste/olfaction | 3 / 374 (1.18) | 0.00 (-0.00) | 0.00 (-0.00) |
| LDL | Liver/biliary system | 15 / 92 (3.81) | 2.16 (2.75) | 1.48 (0.80) |
| LDL | Digestive/alimentary | 9 / 92 (1.33) | 1.97 (2.16) | 1.79 (1.48) |
| LDL | Skeleton | 10 / 92 (0.38) | 0.48 (1.63) | 0.49 (1.45) |
| LDL | Homeostasis/metabolism | 30 / 92 (1.75) | 1.26 (0.63) | 1.04 (0.08) |
| LDL | Growth/size/body region | 21 / 92 (0.91) | 1.21 (0.48) | 1.15 (0.27) |
| LDL | Renal/urinary system | 7 / 92 (0.74) | 1.29 (0.44) | 1.18 (0.21) |
| LDL | Embryo | 6 / 92 (0.09) | 1.21 (0.36) | 1.30 (0.47) |
| LDL | Respiratory system | 5 / 92 (0.22) | 1.23 (0.34) | 1.39 (0.57) |
| LDL | Nervous system | 15 / 92 (0.52) | 1.14 (0.28) | 1.07 (0.13) |
| LDL | Cellular | 18 / 92 (0.58) | 1.13 (0.26) | 1.05 (0.08) |
| LDL | Cardiovascular system | 12 / 92 (0.46) | 0.87 (0.19) | 0.86 (0.21) |
| LDL | Hematopoietic system | 16 / 92 (0.64) | 1.09 (0.17) | 1.03 (0.04) |
| LDL | Behavior/neurological | 17 / 92 (0.61) | 1.10 (0.17) | 1.01 (-0.00) |
| LDL | Hearing/vestibular/ear | 4 / 92 (0.54) | 1.14 (0.16) | 1.28 (0.28) |
| LDL | Limbs/digits/tail | 5 / 92 (0.43) | 0.79 (0.14) | 0.89 (-0.00) |
| LDL | Endocrine/exocrine gland | 9 / 92 (0.27) | 1.07 (0.09) | 0.96 (-0.00) |
| LDL | Immune system | 14 / 92 (0.41) | 1.06 (0.08) | 1.02 (0.04) |
| LDL | Neoplasm | 1 / 92 (0.14) | 0.83 (0.07) | 0.93 (-0.00) |
| LDL | Mortality/aging | 20 / 92 (0.35) | 1.05 (0.07) | 1.03 (0.04) |
| LDL | Craniofacial | 5 / 92 (0.39) | 0.89 (0.06) | 1.00 (-0.00) |
| LDL | Adipose tissue | 9 / 92 (1.50) | 1.04 (0.06) | 0.88 (0.06) |
| LDL | Integument | 10 / 92 (0.71) | 1.04 (0.05) | 0.93 (0.05) |
| LDL | Muscle | 6 / 92 (0.33) | 0.98 (0.00) | 0.96 (-0.00) |
| LDL | Vision/eye | 8 / 92 (0.43) | 0.99 (-0.00) | 1.05 (0.05) |
| LDL | Reproductive system | 8 / 92 (0.17) | 0.99 (-0.00) | 0.83 (0.24) |
| LDL | Pigmentation | 3 / 92 (0.74) | 0.89 (-0.00) | 0.75 (0.10) |
| LDL | Taste/olfaction | 0 / 92 (-0.00) | 0.86 (-0.00) | 0.97 (-0.00) |
| LOAD | Pigmentation | 2 / 98 (0.51) | 3.82 (1.20) | 3.11 (0.78) |
| LOAD | Muscle | 9 / 98 (1.51) | 2.54 (1.16) | 0.44 (0.16) |
| LOAD | Homeostasis/metabolism | 24 / 98 (1.56) | 1.96 (1.06) | 1.41 (0.31) |
| LOAD | Integument | 7 / 98 (0.49) | 2.34 (1.00) | 1.07 (-0.00) |
| LOAD | Immune system | 19 / 98 (1.92) | 1.94 (0.89) | 0.99 (-0.00) |
| LOAD | Vision/eye | 4 / 98 (0.00) | 2.18 (0.82) | 1.90 (0.48) |
| LOAD | Hearing/vestibular/ear | 4 / 98 (0.96) | 2.45 (0.80) | 2.00 (0.52) |
| LOAD | Renal/urinary system | 7 / 98 (1.14) | 2.26 (0.77) | 0.00 (0.43) |
| LOAD | Taste/olfaction | 4 / 98 (3.44) | 5.24 (0.73) | 0.00 (-0.00) |
| LOAD | Cardiovascular system | 17 / 98 (2.22) | 1.83 (0.65) | 0.25 (0.52) |
| LOAD | Behavior/neurological | 13 / 98 (0.46) | 1.62 (0.58) | 0.79 (0.11) |
| LOAD | Embryo | 7 / 98 (0.50) | 0.30 (0.50) | 0.00 (0.89) |

*(continued)*

| Trait | Category | GWAS overlap | Odds ratio (- log 10 $P$-value) | |
|---|---|---|---|---|
| | | | All genes | Non-GWAS genes |
| LOAD | Digestive/alimentary | 6 / 98 (0.81) | 1.67 (0.49) | 0.00 (0.42) |
| LOAD | Liver/biliary system | 5 / 98 (0.43) | 1.66 (0.48) | 0.51 (-0.00) |
| LOAD | Growth/size/body region | 20 / 98 (1.37) | 1.50 (0.41) | 0.83 (-0.00) |
| LOAD | Limbs/digits/tail | 5 / 98 (0.74) | 1.65 (0.36) | 1.35 (0.18) |
| LOAD | Craniofacial | 8 / 98 (1.84) | 1.49 (0.33) | 1.22 (0.17) |
| LOAD | Skeleton | 9 / 98 (0.59) | 0.49 (0.27) | 0.30 (0.51) |
| LOAD | Reproductive system | 7 / 98 (0.32) | 1.40 (0.25) | 1.02 (-0.00) |
| LOAD | Hematopoietic system | 17 / 98 (1.55) | 1.30 (0.20) | 0.80 (0.11) |
| LOAD | Cellular | 20 / 98 (1.57) | 1.25 (0.19) | 0.68 (0.23) |
| LOAD | Endocrine/exocrine gland | 14 / 98 (1.99) | 1.24 (0.11) | 0.30 (0.51) |
| LOAD | Nervous system | 10 / 98 (0.26) | 0.71 (0.10) | 0.43 (0.45) |
| LOAD | Mortality/aging | 18 / 98 (0.62) | 1.10 (0.08) | 0.40 (0.61) |
| LOAD | Respiratory system | 5 / 98 (0.43) | 0.86 (0.00) | 0.00 (0.43) |
| LOAD | Adipose tissue | 5 / 98 (0.66) | 0.99 (-0.00) | 0.60 (-0.00) |
| LOAD | Neoplasm | 2 / 98 (0.16) | 0.74 (-0.00) | 0.00 (0.21) |
| NEU | Endocrine/exocrine gland | 33 / 376 (3.62) | 2.57 (1.37) | 2.54 (1.21) |
| NEU | Mortality/aging | 77 / 376 (6.51) | 2.05 (1.21) | 2.13 (1.10) |
| NEU | Muscle | 23 / 376 (3.38) | 2.63 (1.18) | 2.90 (1.48) |
| NEU | Cellular | 48 / 376 (2.94) | 2.16 (1.17) | 2.22 (1.23) |
| NEU | Growth/size/body region | 62 / 376 (5.32) | 2.08 (1.15) | 2.14 (1.04) |
| NEU | Reproductive system | 32 / 376 (3.78) | 2.27 (0.99) | 2.18 (0.82) |
| NEU | Skeleton | 39 / 376 (5.44) | 2.03 (0.90) | 2.23 (0.98) |
| NEU | Embryo | 22 / 376 (1.87) | 2.20 (0.82) | 2.43 (1.10) |
| NEU | Respiratory system | 22 / 376 (3.70) | 2.21 (0.76) | 2.43 (0.80) |
| NEU | Digestive/alimentary | 12 / 376 (0.70) | 2.14 (0.75) | 2.36 (0.79) |
| NEU | Taste/olfaction | 3 / 376 (1.36) | 4.97 (0.71) | 5.47 (0.74) |
| NEU | Nervous system | 65 / 376 (9.14) | 1.76 (0.61) | 1.36 (0.21) |
| NEU | Hematopoietic system | 44 / 376 (3.44) | 1.69 (0.60) | 1.86 (0.78) |
| NEU | Immune system | 36 / 376 (1.77) | 1.68 (0.59) | 1.85 (0.78) |
| NEU | Behavior/neurological | 65 / 376 (7.69) | 1.61 (0.58) | 1.42 (0.33) |
| NEU | Liver/biliary system | 14 / 376 (1.18) | 1.75 (0.51) | 1.92 (0.55) |
| NEU | Cardiovascular system | 32 / 376 (2.35) | 1.67 (0.50) | 1.84 (0.69) |
| NEU | Vision/eye | 32 / 376 (5.05) | 1.58 (0.43) | 1.74 (0.46) |
| NEU | Limbs/digits/tail | 14 / 376 (1.87) | 1.71 (0.37) | 1.88 (0.39) |
| NEU | Craniofacial | 14 / 376 (1.50) | 1.53 (0.34) | 1.68 (0.36) |
| NEU | Adipose tissue | 23 / 376 (4.94) | 1.49 (0.33) | 1.64 (0.36) |
| NEU | Hearing/vestibular/ear | 8 / 376 (0.87) | 0.00 (0.21) | 0.00 (0.21) |
| NEU | Neoplasm | 12 / 376 (2.40) | 1.59 (0.20) | 1.75 (0.45) |
| NEU | Renal/urinary system | 15 / 376 (1.62) | 1.38 (0.15) | 1.52 (0.34) |
| NEU | Integument | 28 / 376 (3.38) | 1.19 (0.12) | 1.31 (0.12) |
| NEU | Homeostasis/metabolism | 59 / 376 (3.30) | 1.18 (0.08) | 1.29 (0.18) |
| NEU | Pigmentation | 10 / 376 (3.03) | 0.00 (0.00) | 0.00 (-0.00) |

*(continued)*

| Trait | Category | GWAS overlap | Odds ratio (- log 10 $P$-value) | |
|---|---|---|---|---|
| | | | All genes | Non-GWAS genes |
| RA | Hematopoietic system | 44 / 123 (11.12) | 2.14 (5.66) | 1.40 (1.09) |
| RA | Immune system | 42 / 123 (10.20) | 2.05 (5.02) | 1.36 (1.00) |
| RA | Endocrine/exocrine gland | 25 / 123 (6.73) | 2.14 (4.48) | 1.41 (0.95) |
| RA | Neoplasm | 10 / 123 (4.43) | 2.87 (4.20) | 2.22 (2.22) |
| RA | Cellular | 33 / 123 (5.49) | 1.75 (3.19) | 1.32 (0.85) |
| RA | Digestive/alimentary | 12 / 123 (3.49) | 2.10 (3.11) | 1.46 (0.80) |
| RA | Liver/biliary system | 12 / 123 (3.57) | 1.95 (2.42) | 1.62 (1.30) |
| RA | Homeostasis/metabolism | 38 / 123 (5.32) | 1.49 (1.89) | 1.15 (0.37) |
| RA | Respiratory system | 10 / 123 (2.58) | 1.76 (1.79) | 1.49 (0.92) |
| RA | Cardiovascular system | 16 / 123 (2.37) | 1.48 (1.42) | 1.24 (0.47) |
| RA | Nervous system | 17 / 123 (1.86) | 1.45 (1.39) | 1.24 (0.57) |
| RA | Behavior/neurological | 18 / 123 (1.76) | 1.40 (1.25) | 1.19 (0.42) |
| RA | Skeleton | 20 / 123 (4.60) | 1.42 (1.07) | 0.93 (0.04) |
| RA | Reproductive system | 15 / 123 (2.80) | 1.42 (0.94) | 1.00 (0.00) |
| RA | Vision/eye | 16 / 123 (4.03) | 1.40 (0.90) | 0.96 (0.00) |
| RA | Renal/urinary system | 10 / 123 (2.78) | 1.42 (0.78) | 0.96 (0.00) |
| RA | Pigmentation | 4 / 123 (1.63) | 1.69 (0.75) | 1.23 (0.20) |
| RA | Growth/size/body region | 26 / 123 (3.21) | 1.26 (0.73) | 1.00 (-0.00) |
| RA | Integument | 13 / 123 (2.40) | 1.31 (0.69) | 1.05 (0.09) |
| RA | Mortality/aging | 31 / 123 (3.49) | 1.24 (0.67) | 0.96 (0.07) |
| RA | Limbs/digits/tail | 4 / 123 (0.54) | 1.43 (0.66) | 1.09 (0.13) |
| RA | Muscle | 7 / 123 (1.07) | 1.27 (0.51) | 1.08 (0.11) |
| RA | Embryo | 8 / 123 (0.92) | 1.25 (0.47) | 1.05 (0.10) |
| RA | Adipose tissue | 9 / 123 (2.48) | 1.24 (0.41) | 0.72 (0.37) |
| RA | Hearing/vestibular/ear | 3 / 123 (0.66) | 1.20 (0.23) | 1.17 (0.24) |
| RA | Taste/olfaction | 0 / 123 (0.00) | 0.00 (0.19) | 0.00 (0.20) |
| RA | Craniofacial | 5 / 123 (0.73) | 1.04 (0.05) | 0.81 (0.20) |
| SCZ | Nervous system | 121 / 780 (2.38) | 2.04 (13.97) | 2.08 (12.69) |
| SCZ | Behavior/neurological | 130 / 780 (2.19) | 1.90 (11.46) | 1.91 (10.08) |
| SCZ | Growth/size/body region | 130 / 780 (0.87) | 1.77 (9.89) | 1.86 (10.09) |
| SCZ | Muscle | 52 / 780 (1.34) | 2.09 (9.12) | 2.13 (8.33) |
| SCZ | Mortality/aging | 167 / 780 (1.67) | 1.51 (5.61) | 1.53 (5.11) |
| SCZ | Cardiovascular system | 95 / 780 (1.79) | 1.62 (5.60) | 1.61 (4.74) |
| SCZ | Craniofacial | 36 / 780 (0.55) | 1.84 (4.98) | 1.95 (5.18) |
| SCZ | Homeostasis/metabolism | 151 / 780 (0.85) | 1.48 (4.94) | 1.49 (4.41) |
| SCZ | Hematopoietic system | 100 / 780 (0.54) | 1.48 (4.13) | 1.49 (3.64) |
| SCZ | Skeleton | 82 / 780 (1.82) | 1.55 (4.10) | 1.60 (4.10) |
| SCZ | Cellular | 132 / 780 (1.37) | 1.45 (4.09) | 1.48 (3.91) |
| SCZ | Reproductive system | 71 / 780 (1.12) | 1.55 (3.77) | 1.59 (3.72) |
| SCZ | Hearing/vestibular/ear | 22 / 780 (0.41) | 1.82 (3.39) | 1.93 (3.53) |
| SCZ | Adipose tissue | 37 / 780 (0.62) | 1.65 (3.39) | 1.85 (4.44) |

*(continued)*

| Trait | Category | GWAS overlap | Odds ratio (- log 10 $P$-value) | |
|---|---|---|---|---|
| | | | All genes | Non-GWAS genes |
| SCZ | Neoplasm | 24 / 780 (0.55) | 1.81 (3.35) | 1.81 (2.97) |
| SCZ | Endocrine/exocrine gland | 68 / 780 (0.47) | 1.46 (3.08) | 1.49 (2.90) |
| SCZ | Vision/eye | 53 / 780 (0.33) | 1.50 (3.03) | 1.48 (2.51) |
| SCZ | Digestive/alimentary | 39 / 780 (0.34) | 1.55 (2.93) | 1.61 (2.96) |
| SCZ | Respiratory system | 42 / 780 (0.70) | 1.53 (2.67) | 1.61 (2.91) |
| SCZ | Renal/urinary system | 41 / 780 (0.89) | 1.54 (2.64) | 1.58 (2.64) |
| SCZ | Embryo | 60 / 780 (0.88) | 1.42 (2.30) | 1.37 (1.72) |
| SCZ | Immune system | 93 / 780 (0.12) | 1.32 (2.21) | 1.31 (1.86) |
| SCZ | Taste/olfaction | 1 / 780 (0.42) | 2.51 (2.08) | 2.96 (2.57) |
| SCZ | Liver/biliary system | 42 / 780 (0.71) | 1.38 (1.62) | 1.33 (1.22) |
| SCZ | Integument | 53 / 780 (0.16) | 1.31 (1.54) | 1.33 (1.51) |
| SCZ | Limbs/digits/tail | 28 / 780 (0.17) | 1.32 (1.05) | 1.44 (1.55) |
| SCZ | Pigmentation | 7 / 780 (0.66) | 0.70 (0.59) | 0.82 (0.23) |
| T2D | Liver/biliary system | 43 / 300 (12.22) | 10.78 (3.64) | 6.10 (1.25) |
| T2D | Digestive/alimentary | 35 / 300 (8.11) | 9.27 (3.04) | 5.99 (1.23) |
| T2D | Integument | 49 / 300 (9.70) | 7.51 (2.77) | 2.83 (0.55) |
| T2D | Respiratory system | 33 / 300 (7.30) | 8.14 (2.54) | 6.11 (1.25) |
| T2D | Taste/olfaction | 7 / 300 (4.35) | 35.42 (2.47) | 26.56 (1.26) |
| T2D | Cardiovascular system | 64 / 300 (10.34) | 5.81 (2.34) | 2.92 (0.46) |
| T2D | Muscle | 50 / 300 (13.97) | 7.08 (2.28) | 3.56 (0.68) |
| T2D | Adipose tissue | 46 / 300 (15.52) | 7.86 (2.24) | 2.37 (0.36) |
| T2D | Growth/size/body region | 106 / 300 (15.13) | 5.20 (2.16) | 2.61 (0.38) |
| T2D | Renal/urinary system | 37 / 300 (9.88) | 7.21 (2.10) | 6.51 (1.31) |
| T2D | Endocrine/exocrine gland | 68 / 300 (14.73) | 5.61 (2.03) | 1.21 (-0.00) |
| T2D | Vision/eye | 39 / 300 (6.42) | 5.99 (1.99) | 3.00 (0.58) |
| T2D | Embryo | 44 / 300 (8.22) | 5.60 (1.88) | 1.41 (-0.00) |
| T2D | Mortality/aging | 118 / 300 (14.20) | 4.21 (1.73) | 2.11 (0.36) |
| T2D | Neoplasm | 21 / 300 (6.07) | 7.15 (1.55) | 0.00 (0.00) |
| T2D | Homeostasis/metabolism | 113 / 300 (13.25) | 3.87 (1.33) | 1.75 (0.18) |
| T2D | Behavior/neurological | 80 / 300 (10.29) | 3.68 (1.27) | 2.37 (0.42) |
| T2D | Immune system | 68 / 300 (8.03) | 3.66 (1.26) | 1.57 (0.19) |
| T2D | Craniofacial | 29 / 300 (6.74) | 4.59 (1.12) | 0.00 (-0.00) |
| T2D | Hematopoietic system | 65 / 300 (7.43) | 3.15 (1.00) | 1.58 (0.19) |
| T2D | Nervous system | 61 / 300 (6.41) | 2.79 (0.80) | 1.68 (0.20) |
| T2D | Cellular | 87 / 300 (10.57) | 2.70 (0.74) | 0.68 (-0.00) |
| T2D | Reproductive system | 40 / 300 (5.16) | 2.85 (0.73) | 1.43 (-0.00) |
| T2D | Skeleton | 54 / 300 (9.28) | 3.12 (0.68) | 2.35 (0.23) |
| T2D | Hearing/vestibular/ear | 15 / 300 (3.35) | 2.51 (0.41) | 3.78 (0.51) |
| T2D | Limbs/digits/tail | 25 / 300 (5.89) | 1.76 (0.30) | 0.00 (-0.00) |
| T2D | Pigmentation | 14 / 300 (4.82) | 0.00 (0.00) | 0.00 (-0.00) |
| WAIST | Limbs/digits/tail | 46 / 391 (16.62) | 4.46 (3.63) | 2.93 (1.44) |
| WAIST | Muscle | 72 / 391 (24.63) | 3.17 (2.41) | 2.24 (1.05) |

*(continued)*

| Trait | Category | GWAS overlap | Odds ratio (- log 10 $P$-value) | |
|---|---|---|---|---|
| | | | All genes | Non-GWAS genes |
| WAIST | Embryo | 71 / 391 (19.09) | 2.66 (1.88) | 1.61 (0.39) |
| WAIST | Vision/eye | 52 / 391 (10.74) | 2.46 (1.64) | 1.35 (0.24) |
| WAIST | Skeleton | 96 / 391 (25.48) | 2.30 (1.62) | 1.51 (0.35) |
| WAIST | Integument | 64 / 391 (14.99) | 2.32 (1.55) | 1.52 (0.38) |
| WAIST | Respiratory system | 49 / 391 (14.22) | 2.58 (1.50) | 2.62 (1.19) |
| WAIST | Immune system | 102 / 391 (17.34) | 2.07 (1.46) | 1.58 (0.54) |
| WAIST | Craniofacial | 59 / 391 (22.39) | 2.64 (1.37) | 1.73 (0.51) |
| WAIST | Pigmentation | 19 / 391 (7.64) | 3.31 (1.32) | 2.17 (0.58) |
| WAIST | Behavior/neurological | 109 / 391 (17.77) | 1.99 (1.27) | 1.51 (0.40) |
| WAIST | Mortality/aging | 146 / 391 (20.15) | 1.77 (1.08) | 1.35 (0.35) |
| WAIST | Growth/size/body region | 135 / 391 (22.39) | 1.80 (1.04) | 1.65 (0.63) |
| WAIST | Renal/urinary system | 42 / 391 (11.60) | 2.09 (1.00) | 0.78 (-0.00) |
| WAIST | Nervous system | 91 / 391 (14.22) | 1.71 (0.86) | 1.64 (0.55) |
| WAIST | Cardiovascular system | 94 / 391 (20.34) | 1.76 (0.79) | 1.42 (0.32) |
| WAIST | Digestive/alimentary | 47 / 391 (13.07) | 1.94 (0.77) | 0.73 (-0.00) |
| WAIST | Hematopoietic system | 98 / 391 (16.35) | 1.64 (0.72) | 1.15 (0.09) |
| WAIST | Neoplasm | 26 / 391 (8.32) | 2.06 (0.58) | 1.35 (0.18) |
| WAIST | Liver/biliary system | 57 / 391 (18.79) | 1.70 (0.57) | 1.11 (0.13) |
| WAIST | Cellular | 117 / 391 (17.93) | 1.49 (0.55) | 1.10 (0.08) |
| WAIST | Reproductive system | 62 / 391 (12.15) | 1.46 (0.45) | 0.96 (0.00) |
| WAIST | Adipose tissue | 50 / 391 (16.85) | 1.61 (0.43) | 1.27 (0.14) |
| WAIST | Endocrine/exocrine gland | 83 / 391 (19.81) | 1.49 (0.43) | 0.87 (-0.00) |
| WAIST | Hearing/vestibular/ear | 37 / 391 (15.85) | 1.58 (0.35) | 0.69 (-0.00) |
| WAIST | Homeostasis/metabolism | 136 / 391 (17.99) | 1.22 (0.22) | 1.10 (0.08) |
| WAIST | Taste/olfaction | 6 / 391 (3.29) | 0.00 (-0.00) | 0.00 (-0.00) |

#### Supplementary Table 13

For each GWAS trait we quantify overlap between RSS-NET prioritized genes ($P_1^{\text{bma}} \geq 0.9$) and genes causing 19 categories of Mendelian disorders (Freund et al. 2018; Amberger et al. 2019). The rest is the same as **Supplementary Table 12**.

| | | | Odds ratio (- log 10 $P$-value) | |
|---|---|---|---|---|
| Trait | Category | GWAS overlap | All genes | Non-GWAS genes |
| AF | Arrythmia | 16 / 184 (10.78) | 8.28 (4.77) | 5.29 (2.02) |
| AF | Cardiovascular | 15 / 184 (5.47) | 3.31 (2.06) | 1.81 (0.60) |
| AF | Diabetes | 3 / 184 (1.10) | 3.21 (0.86) | 2.04 (0.40) |
| AF | Positive mood | 3 / 184 (2.18) | 5.02 (0.73) | 6.39 (0.82) |
| AF | Renal | 10 / 184 (1.55) | 1.73 (0.64) | 0.44 (0.14) |
| AF | Immune | 4 / 184 (0.26) | 1.53 (0.34) | 1.30 (0.17) |
| AF | MODY | 7 / 184 (1.41) | 1.51 (0.34) | 1.28 (0.17) |
| AF | Psychiatric | 5 / 184 (1.78) | 1.13 (0.23) | 1.44 (0.29) |
| AF | Reproductive | 1 / 184 (-0.00) | 1.03 (0.20) | 1.31 (0.27) |
| AF | Hematologic | 5 / 184 (0.63) | 1.04 (0.14) | 0.66 (-0.00) |
| AF | Growth | 7 / 184 (0.91) | 0.40 (0.14) | 0.00 (0.59) |
| AF | Insulin | 8 / 184 (1.60) | 0.91 (0.00) | 0.58 (-0.00) |
| AF | Platelet | 4 / 184 (0.51) | 0.63 (0.00) | 0.00 (0.20) |
| AF | Uric acid | 1 / 184 (-0.00) | 0.00 (0.00) | 0.00 (-0.00) |
| AF | Neurologic | 3 / 184 (0.91) | 0.00 (0.00) | 0.00 (-0.00) |
| AF | Development | 4 / 184 (0.25) | 0.51 (-0.00) | 0.65 (-0.00) |
| AF | Autism | 0 / 184 (-0.00) | 0.00 (-0.00) | 0.00 (-0.00) |
| AF | Weight | 3 / 184 (1.47) | 0.00 (-0.00) | 0.00 (-0.00) |
| AF | Microalbumin | 0 / 184 (-0.00) | 0.00 (-0.00) | 0.00 (-0.00) |
| BC | Insulin | 16 / 364 (3.90) | 9.71 (4.35) | 7.77 (2.41) |
| BC | Cardiovascular | 8 / 364 (0.78) | 7.09 (2.77) | 7.94 (2.44) |
| BC | Platelet | 10 / 364 (2.32) | 5.71 (1.65) | 5.33 (1.17) |
| BC | Renal | 22 / 364 (5.16) | 4.15 (1.59) | 2.90 (0.75) |
| BC | Hematologic | 10 / 364 (1.56) | 4.71 (1.45) | 4.40 (1.03) |
| BC | Uric acid | 6 / 364 (1.97) | 7.48 (1.45) | 0.00 (-0.00) |
| BC | Immune | 10 / 364 (1.57) | 4.59 (1.42) | 4.29 (1.01) |
| BC | MODY | 8 / 364 (0.83) | 4.55 (1.41) | 4.25 (1.00) |
| BC | Reproductive | 6 / 364 (1.47) | 6.23 (1.31) | 0.00 (-0.00) |
| BC | Growth | 16 / 364 (3.21) | 3.66 (1.19) | 1.71 (0.33) |
| BC | Positive mood | 4 / 364 (2.58) | 15.10 (1.16) | 21.12 (1.29) |
| BC | Weight | 2 / 364 (0.55) | 6.40 (0.81) | 0.00 (-0.00) |
| BC | Development | 13 / 364 (2.75) | 3.10 (0.81) | 2.17 (0.41) |
| BC | Microalbumin | 3 / 364 (0.84) | 5.64 (0.76) | 0.00 (-0.00) |
| BC | Diabetes | 3 / 364 (0.72) | 4.80 (0.70) | 0.00 (-0.00) |
| BC | Psychiatric | 7 / 364 (2.15) | 3.40 (0.57) | 4.76 (0.69) |
| BC | Arrythmia | 8 / 364 (2.61) | 0.00 (-0.00) | 0.00 (-0.00) |
| BC | Autism | 0 / 364 (-0.00) | 0.00 (-0.00) | 0.00 (-0.00) |
| BC | Neurologic | 6 / 364 (1.95) | 0.00 (-0.00) | 0.00 (0.00) |

*(continued)*

| Trait | Category | GWAS overlap | Odds ratio (- log 10 $P$-value) All genes | Non-GWAS genes |
|---|---|---|---|---|
| BMI | Insulin | 38 / 889 (2.71) | 2.54 (1.22) | 2.08 (0.71) |
| BMI | Weight | 10 / 889 (1.59) | 4.75 (1.14) | 3.24 (0.56) |
| BMI | Platelet | 29 / 889 (2.75) | 2.10 (0.73) | 1.90 (0.53) |
| BMI | Psychiatric | 19 / 889 (2.42) | 2.51 (0.70) | 1.71 (0.34) |
| BMI | Autism | 11 / 889 (2.69) | 3.00 (0.54) | 0.00 (-0.00) |
| BMI | Renal | 48 / 889 (2.77) | 1.52 (0.46) | 1.04 (0.14) |
| BMI | Microalbumin | 9 / 889 (0.73) | 2.07 (0.41) | 0.00 (0.00) |
| BMI | Hematologic | 28 / 889 (1.22) | 1.73 (0.37) | 1.58 (0.42) |
| BMI | Cardiovascular | 26 / 889 (0.58) | 1.56 (0.35) | 1.41 (0.18) |
| BMI | Neurologic | 14 / 889 (1.39) | 1.53 (0.31) | 2.09 (0.41) |
| BMI | Growth | 42 / 889 (2.54) | 1.35 (0.31) | 1.22 (0.17) |
| BMI | Uric acid | 11 / 889 (0.58) | 1.37 (0.28) | 1.85 (0.37) |
| BMI | Arrythmia | 11 / 889 (0.21) | 1.11 (0.22) | 1.52 (0.31) |
| BMI | Development | 46 / 889 (5.79) | 1.14 (0.16) | 1.55 (0.42) |
| BMI | Immune | 30 / 889 (1.59) | 1.12 (0.15) | 1.53 (0.41) |
| BMI | MODY | 26 / 889 (0.79) | 1.11 (0.15) | 0.76 (-0.00) |
| BMI | Reproductive | 20 / 889 (2.26) | 0.00 (-0.00) | 0.00 (-0.00) |
| BMI | Diabetes | 14 / 889 (2.15) | 0.00 (-0.00) | 0.00 (-0.00) |
| BMI | Positive mood | 7 / 889 (1.96) | 0.00 (-0.00) | 0.00 (-0.00) |
| CAD | Cardiovascular | 26 / 308 (11.19) | 3.70 (4.01) | 2.31 (1.37) |
| CAD | Platelet | 17 / 308 (7.00) | 3.19 (2.44) | 2.21 (1.05) |
| CAD | Insulin | 25 / 308 (10.10) | 2.82 (2.44) | 1.60 (0.59) |
| CAD | Uric acid | 10 / 308 (5.10) | 4.19 (2.35) | 3.48 (1.48) |
| CAD | Hematologic | 15 / 308 (4.54) | 2.62 (1.93) | 1.45 (0.44) |
| CAD | Neurologic | 6 / 308 (2.20) | 3.89 (1.92) | 0.97 (-0.00) |
| CAD | Renal | 23 / 308 (6.48) | 2.31 (1.87) | 1.44 (0.35) |
| CAD | Development | 12 / 308 (2.73) | 2.30 (1.50) | 1.44 (0.27) |
| CAD | Microalbumin | 7 / 308 (3.69) | 3.15 (1.11) | 2.63 (0.74) |
| CAD | MODY | 17 / 308 (5.62) | 1.96 (0.99) | 1.75 (0.65) |
| CAD | Reproductive | 8 / 308 (2.81) | 2.30 (0.97) | 2.15 (0.76) |
| CAD | Autism | 2 / 308 (0.72) | 3.02 (0.83) | 1.88 (0.38) |
| CAD | Immune | 14 / 308 (4.00) | 1.69 (0.74) | 1.41 (0.27) |
| CAD | Growth | 10 / 308 (1.15) | 1.58 (0.64) | 1.68 (0.56) |
| CAD | Positive mood | 2 / 308 (1.05) | 2.77 (0.51) | 0.00 (-0.00) |
| CAD | Diabetes | 9 / 308 (4.96) | 1.77 (0.50) | 2.20 (0.62) |
| CAD | Arrythmia | 7 / 308 (2.33) | 1.67 (0.36) | 0.00 (0.39) |
| CAD | Weight | 4 / 308 (1.83) | 1.17 (0.24) | 0.00 (-0.00) |
| CAD | Psychiatric | 5 / 308 (1.34) | 1.25 (0.17) | 0.78 (-0.00) |
| HDL | Cardiovascular | 10 / 104 (4.09) | 2.69 (3.24) | 1.66 (0.73) |
| HDL | Diabetes | 5 / 104 (3.31) | 4.39 (3.07) | 2.56 (1.10) |
| HDL | Insulin | 10 / 104 (3.95) | 2.46 (2.60) | 1.61 (0.72) |
| HDL | Weight | 5 / 104 (4.01) | 4.38 (2.44) | 2.55 (0.91) |

*(continued)*

| Trait | Category | GWAS overlap | Odds ratio (- log 10 $P$-value) All genes | Non-GWAS genes |
|---|---|---|---|---|
| HDL | MODY | 7 / 104 (2.39) | 2.37 (2.22) | 1.58 (0.60) |
| HDL | Microalbumin | 2 / 104 (0.99) | 3.17 (1.59) | 2.96 (1.27) |
| HDL | Renal | 4 / 104 (0.27) | 1.49 (0.78) | 1.47 (0.59) |
| HDL | Growth | 9 / 104 (2.87) | 1.48 (0.72) | 1.10 (0.16) |
| HDL | Hematologic | 6 / 104 (1.87) | 1.56 (0.70) | 0.81 (-0.00) |
| HDL | Immune | 4 / 104 (0.92) | 1.52 (0.68) | 1.18 (0.19) |
| HDL | Platelet | 5 / 104 (1.70) | 1.47 (0.46) | 0.98 (0.00) |
| HDL | Positive mood | 0 / 104 (-0.00) | 1.65 (0.34) | 1.92 (0.38) |
| HDL | Neurologic | 1 / 104 (0.28) | 1.37 (0.31) | 1.07 (0.15) |
| HDL | Arrythmia | 0 / 104 (-0.00) | 1.34 (0.26) | 1.55 (0.47) |
| HDL | Development | 3 / 104 (0.35) | 1.20 (0.17) | 1.00 (-0.00) |
| HDL | Uric acid | 0 / 104 (-0.00) | 1.23 (0.13) | 1.43 (0.33) |
| HDL | Autism | 0 / 104 (-0.00) | 0.89 (-0.00) | 1.04 (0.21) |
| HDL | Psychiatric | 1 / 104 (0.23) | 0.74 (-0.00) | 0.87 (-0.00) |
| HDL | Reproductive | 2 / 104 (0.60) | 0.68 (-0.00) | 0.79 (-0.00) |
| HR | Arrythmia | 9 / 123 (5.41) | 7.16 (2.93) | 5.86 (1.72) |
| HR | Cardiovascular | 10 / 123 (3.50) | 2.64 (1.10) | 1.80 (0.49) |
| HR | Neurologic | 1 / 123 (0.23) | 1.94 (0.38) | 2.64 (0.49) |
| HR | Renal | 6 / 123 (0.79) | 1.45 (0.33) | 1.32 (0.18) |
| HR | Psychiatric | 2 / 123 (0.54) | 1.58 (0.32) | 2.16 (0.42) |
| HR | Immune | 3 / 123 (0.31) | 0.00 (0.19) | 0.00 (0.21) |
| HR | MODY | 4 / 123 (0.53) | 0.00 (0.19) | 0.00 (0.21) |
| HR | Development | 1 / 123 (0.14) | 1.45 (0.19) | 1.97 (0.54) |
| HR | Growth | 3 / 123 (0.12) | 0.57 (0.00) | 0.77 (-0.00) |
| HR | Diabetes | 1 / 123 (0.28) | 0.00 (0.00) | 0.00 (-0.00) |
| HR | Platelet | 3 / 123 (0.37) | 0.88 (-0.00) | 1.20 (0.24) |
| HR | Hematologic | 1 / 123 (0.14) | 0.73 (-0.00) | 0.99 (-0.00) |
| HR | Insulin | 2 / 123 (-0.00) | 0.64 (-0.00) | 0.87 (-0.00) |
| HR | Autism | 0 / 123 (-0.00) | 0.00 (-0.00) | 0.00 (-0.00) |
| HR | Weight | 1 / 123 (0.39) | 0.00 (-0.00) | 0.00 (-0.00) |
| HR | Uric acid | 0 / 123 (-0.00) | 0.00 (-0.00) | 0.00 (0.00) |
| HR | Reproductive | 0 / 123 (0.20) | 0.00 (-0.00) | 0.00 (-0.00) |
| HR | Microalbumin | 1 / 123 (0.32) | 0.00 (-0.00) | 0.00 (0.00) |
| HR | Positive mood | 3 / 123 (2.52) | 0.00 (-0.00) | 0.00 (-0.00) |
| IBD | Immune | 27 / 374 (9.51) | 4.32 (6.22) | 3.87 (4.14) |
| IBD | MODY | 21 / 374 (5.77) | 3.92 (4.95) | 4.10 (4.40) |
| IBD | Psychiatric | 5 / 374 (0.68) | 5.16 (3.80) | 6.44 (4.46) |
| IBD | Uric acid | 13 / 374 (5.97) | 4.46 (3.06) | 2.83 (1.21) |
| IBD | Renal | 15 / 374 (1.25) | 2.45 (2.42) | 2.05 (1.39) |
| IBD | Growth | 14 / 374 (1.41) | 2.45 (2.12) | 2.61 (2.18) |
| IBD | Neurologic | 3 / 374 (0.30) | 3.48 (1.46) | 3.25 (1.13) |

*(continued)*

| | | | Odds ratio (- log 10 $P$-value) | |
|---|---|---|---|---|
| Trait | Category | GWAS overlap | All genes | Non-GWAS genes |
| IBD | Insulin | 19 / 374 (4.13) | 1.99 (1.18) | 1.52 (0.52) |
| IBD | Development | 10 / 374 (0.84) | 1.91 (1.07) | 1.48 (0.40) |
| IBD | Arrythmia | 6 / 374 (1.14) | 2.11 (0.87) | 2.01 (0.70) |
| IBD | Autism | 1 / 374 (-0.00) | 3.10 (0.83) | 1.93 (0.38) |
| IBD | Cardiovascular | 8 / 374 (0.38) | 1.75 (0.83) | 1.65 (0.56) |
| IBD | Hematologic | 17 / 374 (3.84) | 1.72 (0.81) | 1.10 (0.10) |
| IBD | Platelet | 12 / 374 (2.47) | 1.67 (0.56) | 1.81 (0.67) |
| IBD | Diabetes | 5 / 374 (1.35) | 1.69 (0.47) | 1.05 (0.21) |
| IBD | Reproductive | 2 / 374 (0.11) | 1.46 (0.33) | 1.82 (0.62) |
| IBD | Weight | 2 / 374 (0.41) | 1.25 (0.25) | 1.59 (0.32) |
| IBD | Microalbumin | 1 / 374 (-0.00) | 0.86 (-0.00) | 1.07 (0.21) |
| IBD | Positive mood | 0 / 374 (-0.00) | 0.00 (-0.00) | 0.00 (-0.00) |
| LDL | Cardiovascular | 10 / 92 (4.51) | 3.40 (4.96) | 2.38 (1.93) |
| LDL | MODY | 5 / 92 (1.53) | 2.35 (2.06) | 1.90 (1.01) |
| LDL | Hematologic | 9 / 92 (4.07) | 2.24 (1.83) | 1.52 (0.62) |
| LDL | Uric acid | 4 / 92 (2.32) | 2.67 (1.51) | 2.07 (0.86) |
| LDL | Platelet | 3 / 92 (0.83) | 2.03 (1.28) | 2.10 (1.20) |
| LDL | Insulin | 6 / 92 (1.87) | 1.79 (1.17) | 1.52 (0.58) |
| LDL | Renal | 10 / 92 (3.36) | 1.59 (0.96) | 1.43 (0.60) |
| LDL | Reproductive | 2 / 92 (0.68) | 0.00 (0.93) | 0.00 (0.75) |
| LDL | Neurologic | 0 / 92 (-0.00) | 1.97 (0.81) | 2.30 (0.97) |
| LDL | Development | 2 / 92 (0.16) | 1.65 (0.73) | 1.50 (0.47) |
| LDL | Immune | 6 / 92 (2.11) | 1.63 (0.72) | 1.26 (0.31) |
| LDL | Microalbumin | 3 / 92 (1.90) | 2.00 (0.70) | 2.33 (0.83) |
| LDL | Arrythmia | 2 / 92 (0.71) | 1.43 (0.43) | 1.25 (0.13) |
| LDL | Weight | 2 / 92 (1.25) | 1.50 (0.41) | 0.00 (0.20) |
| LDL | Diabetes | 2 / 92 (0.99) | 1.70 (0.37) | 0.66 (-0.00) |
| LDL | Psychiatric | 0 / 92 (0.00) | 0.40 (0.28) | 0.46 (0.14) |
| LDL | Positive mood | 0 / 92 (-0.00) | 0.00 (0.00) | 0.00 (0.00) |
| LDL | Autism | 0 / 92 (-0.00) | 0.95 (-0.00) | 1.11 (0.22) |
| LDL | Growth | 4 / 92 (0.55) | 0.86 (-0.00) | 0.83 (0.08) |
| LOAD | Immune | 6 / 98 (2.30) | 7.32 (3.77) | 2.79 (0.74) |
| LOAD | Renal | 5 / 98 (1.08) | 5.73 (3.49) | 3.82 (1.49) |
| LOAD | Platelet | 5 / 98 (2.07) | 7.78 (3.48) | 3.45 (0.88) |
| LOAD | MODY | 4 / 98 (1.16) | 6.19 (2.98) | 2.75 (0.73) |
| LOAD | Cardiovascular | 5 / 98 (1.58) | 5.79 (2.84) | 3.86 (1.25) |
| LOAD | Arrythmia | 1 / 98 (0.28) | 6.58 (1.84) | 5.85 (1.26) |
| LOAD | Hematologic | 3 / 98 (0.73) | 4.30 (1.68) | 1.43 (0.29) |
| LOAD | Microalbumin | 2 / 98 (1.16) | 7.69 (1.48) | 0.00 (-0.00) |
| LOAD | Insulin | 4 / 98 (1.03) | 3.67 (1.48) | 2.45 (0.66) |
| LOAD | Neurologic | 1 / 98 (0.35) | 5.88 (1.28) | 3.92 (0.62) |
| LOAD | Uric acid | 2 / 98 (0.92) | 5.06 (1.17) | 0.00 (-0.00) |

*(continued)*

| Trait | Category | GWAS overlap | Odds ratio (- log 10 $P$-value) | |
|---|---|---|---|---|
| | | | All genes | Non-GWAS genes |
| LOAD | Weight | 0 / 98 (0.00) | 4.46 (0.68) | 5.93 (0.78) |
| LOAD | Diabetes | 0 / 98 (-0.00) | 3.21 (0.56) | 4.28 (0.66) |
| LOAD | Psychiatric | 2 / 98 (0.80) | 2.43 (0.46) | 0.00 (-0.00) |
| LOAD | Growth | 3 / 98 (0.35) | 1.67 (0.45) | 0.00 (-0.00) |
| LOAD | Development | 1 / 98 (-0.00) | 1.10 (0.22) | 1.47 (0.29) |
| LOAD | Reproductive | 3 / 98 (1.35) | 0.00 (0.00) | 0.00 (-0.00) |
| LOAD | Autism | 1 / 98 (0.59) | 0.00 (-0.00) | 0.00 (0.00) |
| LOAD | Positive mood | 0 / 98 (0.00) | 0.00 (-0.00) | 0.00 (-0.00) |
| NEU | Arrythmia | 4 / 376 (0.49) | 4.74 (1.50) | 4.94 (1.55) |
| NEU | Positive mood | 4 / 376 (2.40) | 7.82 (0.90) | 8.48 (0.93) |
| NEU | Platelet | 9 / 376 (1.41) | 1.97 (0.55) | 2.04 (0.57) |
| NEU | Development | 16 / 376 (3.64) | 1.61 (0.43) | 0.84 (-0.00) |
| NEU | Immune | 12 / 376 (2.16) | 1.58 (0.43) | 1.64 (0.44) |
| NEU | Neurologic | 7 / 376 (2.28) | 2.15 (0.42) | 2.25 (0.43) |
| NEU | Renal | 11 / 376 (0.58) | 1.61 (0.36) | 1.67 (0.36) |
| NEU | Psychiatric | 3 / 376 (0.13) | 1.76 (0.35) | 0.00 (-0.00) |
| NEU | Cardiovascular | 4 / 376 (0.08) | 1.46 (0.19) | 1.52 (0.40) |
| NEU | Growth | 13 / 376 (1.55) | 1.26 (0.17) | 1.31 (0.18) |
| NEU | Hematologic | 6 / 376 (0.19) | 0.81 (0.00) | 0.84 (-0.00) |
| NEU | MODY | 10 / 376 (1.34) | 0.78 (0.00) | 0.81 (-0.00) |
| NEU | Diabetes | 3 / 376 (0.62) | 0.00 (0.00) | 0.00 (-0.00) |
| NEU | Insulin | 8 / 376 (0.41) | 0.71 (-0.00) | 0.74 (-0.00) |
| NEU | Autism | 3 / 376 (1.07) | 0.00 (-0.00) | 0.00 (-0.00) |
| NEU | Weight | 2 / 376 (0.48) | 0.00 (-0.00) | 0.00 (-0.00) |
| NEU | Uric acid | 2 / 376 (-0.00) | 0.00 (-0.00) | 0.00 (0.00) |
| NEU | Reproductive | 6 / 376 (1.26) | 0.00 (-0.00) | 0.00 (-0.00) |
| NEU | Microalbumin | 1 / 376 (-0.00) | 0.00 (-0.00) | 0.00 (-0.00) |
| RA | Immune | 12 / 123 (5.69) | 3.11 (5.04) | 1.94 (1.40) |
| RA | MODY | 8 / 123 (2.89) | 2.25 (2.47) | 1.43 (0.51) |
| RA | Hematologic | 4 / 123 (0.87) | 2.05 (1.85) | 1.80 (1.18) |
| RA | Uric acid | 1 / 123 (0.27) | 2.61 (1.80) | 2.74 (1.73) |
| RA | Platelet | 5 / 123 (1.64) | 1.81 (1.19) | 1.18 (0.19) |
| RA | Renal | 4 / 123 (0.27) | 1.61 (1.18) | 1.50 (0.75) |
| RA | Positive mood | 0 / 123 (-0.00) | 2.61 (0.73) | 3.13 (0.85) |
| RA | Insulin | 5 / 123 (1.13) | 1.42 (0.66) | 0.99 (-0.00) |
| RA | Psychiatric | 0 / 123 (-0.00) | 0.29 (0.57) | 0.34 (0.27) |
| RA | Growth | 3 / 123 (0.13) | 1.37 (0.50) | 1.38 (0.56) |
| RA | Cardiovascular | 0 / 123 (0.57) | 1.21 (0.32) | 1.45 (0.61) |
| RA | Development | 3 / 123 (0.34) | 1.20 (0.24) | 1.28 (0.27) |
| RA | Neurologic | 0 / 123 (0.00) | 1.07 (0.12) | 1.28 (0.29) |
| RA | Arrythmia | 0 / 123 (-0.00) | 1.04 (0.10) | 1.24 (0.24) |
| RA | Reproductive | 0 / 123 (-0.00) | 0.79 (0.00) | 0.95 (-0.00) |

*(continued)*

| Trait | Category | GWAS overlap | Odds ratio (- log 10 $P$-value) | |
|---|---|---|---|---|
| | | | All genes | Non-GWAS genes |
| RA | Microalbumin | 2 / 123 (0.96) | 0.96 (-0.00) | 0.00 (0.38) |
| RA | Diabetes | 1 / 123 (0.33) | 0.82 (-0.00) | 0.49 (0.14) |
| RA | Autism | 0 / 123 (-0.00) | 0.69 (-0.00) | 0.83 (-0.00) |
| RA | Weight | 1 / 123 (0.44) | 0.54 (-0.00) | 0.00 (0.39) |
| SCZ | Development | 28 / 780 (1.92) | 2.23 (6.26) | 2.22 (5.35) |
| SCZ | Psychiatric | 17 / 780 (2.63) | 2.22 (3.21) | 2.14 (2.47) |
| SCZ | Positive mood | 7 / 780 (2.43) | 2.25 (1.18) | 1.42 (0.32) |
| SCZ | Autism | 5 / 780 (0.60) | 1.79 (0.96) | 1.42 (0.35) |
| SCZ | Uric acid | 3 / 780 (0.65) | 0.51 (0.92) | 0.50 (0.87) |
| SCZ | Arrythmia | 4 / 780 (0.70) | 1.38 (0.68) | 1.36 (0.64) |
| SCZ | Weight | 4 / 780 (0.10) | 1.54 (0.65) | 1.81 (1.08) |
| SCZ | Immune | 11 / 780 (0.53) | 1.24 (0.63) | 1.15 (0.35) |
| SCZ | Reproductive | 9 / 780 (0.14) | 1.34 (0.59) | 1.39 (0.65) |
| SCZ | Hematologic | 19 / 780 (0.36) | 1.23 (0.57) | 1.23 (0.49) |
| SCZ | Diabetes | 3 / 780 (0.30) | 1.38 (0.53) | 1.35 (0.46) |
| SCZ | Platelet | 11 / 780 (0.11) | 1.17 (0.36) | 1.27 (0.55) |
| SCZ | Insulin | 14 / 780 (0.34) | 1.14 (0.35) | 1.15 (0.32) |
| SCZ | Renal | 19 / 780 (0.41) | 1.13 (0.34) | 1.07 (0.14) |
| SCZ | Growth | 18 / 780 (0.19) | 0.88 (0.23) | 0.94 (0.07) |
| SCZ | Microalbumin | 4 / 780 (-0.00) | 0.78 (0.15) | 0.61 (0.38) |
| SCZ | MODY | 9 / 780 (1.05) | 1.03 (0.07) | 1.05 (0.08) |
| SCZ | Neurologic | 11 / 780 (1.16) | 0.87 (0.06) | 0.68 (0.32) |
| SCZ | Cardiovascular | 14 / 780 (0.28) | 0.96 (0.03) | 0.97 (-0.00) |
| T2D | Immune | 19 / 300 (4.39) | 11.17 (4.07) | 8.85 (2.03) |
| T2D | Weight | 18 / 300 (13.61) | 28.03 (3.49) | 15.19 (1.14) |
| T2D | Diabetes | 23 / 300 (16.01) | 19.27 (3.03) | 10.44 (0.98) |
| T2D | Neurologic | 12 / 300 (4.84) | 16.09 (2.81) | 8.41 (0.90) |
| T2D | Psychiatric | 8 / 300 (1.91) | 11.68 (2.43) | 12.29 (1.77) |
| T2D | MODY | 29 / 300 (9.87) | 7.42 (2.36) | 5.88 (1.20) |
| T2D | Cardiovascular | 26 / 300 (7.49) | 6.89 (2.25) | 5.47 (1.15) |
| T2D | Insulin | 40 / 300 (16.01) | 6.49 (2.16) | 5.15 (1.11) |
| T2D | Microalbumin | 9 / 300 (3.95) | 12.85 (1.85) | 20.60 (2.19) |
| T2D | Reproductive | 11 / 300 (3.20) | 8.07 (1.48) | 0.00 (-0.00) |
| T2D | Positive mood | 3 / 300 (1.35) | 20.43 (1.27) | 0.00 (0.00) |
| T2D | Renal | 22 / 300 (3.07) | 3.70 (1.17) | 5.84 (1.57) |
| T2D | Platelet | 16 / 300 (4.04) | 4.75 (1.09) | 3.74 (0.59) |
| T2D | Development | 12 / 300 (1.30) | 3.58 (0.89) | 2.82 (0.48) |
| T2D | Growth | 23 / 300 (4.57) | 2.99 (0.77) | 2.36 (0.43) |
| T2D | Uric acid | 10 / 300 (3.54) | 4.85 (0.70) | 0.00 (-0.00) |
| T2D | Hematologic | 18 / 300 (3.90) | 1.86 (0.36) | 0.00 (-0.00) |
| T2D | Arrythmia | 3 / 300 (0.00) | 0.00 (-0.00) | 0.00 (-0.00) |

*(continued)*

| | | | Odds ratio (- log 10 $P$-value) | |
|---|---|---|---|---|
| Trait | Category | GWAS overlap | All genes | Non-GWAS genes |
| T2D | Autism | 4 / 300 (1.41) | 0.00 (-0.00) | 0.00 (0.00) |
| WAIST | Platelet | 14 / 391 (2.11) | 3.34 (1.64) | 1.92 (0.53) |
| WAIST | Diabetes | 8 / 391 (2.26) | 5.09 (1.60) | 0.00 (-0.00) |
| WAIST | Cardiovascular | 17 / 391 (2.03) | 2.47 (1.19) | 2.12 (0.73) |
| WAIST | Renal | 30 / 391 (4.75) | 2.18 (0.92) | 2.08 (0.83) |
| WAIST | Uric acid | 10 / 391 (2.79) | 2.61 (0.73) | 1.87 (0.37) |
| WAIST | Insulin | 21 / 391 (3.24) | 1.93 (0.56) | 0.69 (-0.00) |
| WAIST | Growth | 24 / 391 (3.37) | 1.71 (0.51) | 0.61 (-0.00) |
| WAIST | Weight | 5 / 391 (1.41) | 2.23 (0.43) | 0.00 (-0.00) |
| WAIST | Microalbumin | 6 / 391 (1.54) | 1.97 (0.39) | 0.00 (-0.00) |
| WAIST | Development | 15 / 391 (1.45) | 0.00 (0.37) | 0.00 (0.20) |
| WAIST | Immune | 17 / 391 (2.23) | 1.60 (0.36) | 0.77 (-0.00) |
| WAIST | MODY | 14 / 391 (1.32) | 1.59 (0.35) | 0.00 (0.20) |
| WAIST | Neurologic | 6 / 391 (0.84) | 1.45 (0.30) | 2.07 (0.41) |
| WAIST | Psychiatric | 4 / 391 (0.10) | 1.19 (0.24) | 1.70 (0.34) |
| WAIST | Reproductive | 12 / 391 (2.89) | 1.08 (0.22) | 0.00 (0.00) |
| WAIST | Arrythmia | 7 / 391 (0.91) | 1.06 (0.21) | 1.51 (0.31) |
| WAIST | Hematologic | 13 / 391 (1.00) | 1.10 (0.15) | 0.79 (-0.00) |
| WAIST | Autism | 0 / 391 (0.38) | 0.00 (-0.00) | 0.00 (0.00) |
| WAIST | Positive mood | 2 / 391 (0.58) | 0.00 (-0.00) | 0.00 (-0.00) |

**Supplementary Table 14**

Descriptive statistics of proportion of GWAS SNPs inside a network ($a_j = 1$) over 18 traits for each of 38 regulatory networks. Rows are sorted by the "Median" column.

| | Summary of $\sum_{j=1}^{p} a_j/p$ | | | | |
|---|---|---|---|---|---|
| Network | Min | Q1 | Median | Q3 | Max |
| Vagina | 0.6400 | 0.6423 | 0.6438 | 0.6481 | 0.6505 |
| Lung | 0.6377 | 0.6399 | 0.6414 | 0.6460 | 0.6485 |
| Thyroid | 0.6287 | 0.6305 | 0.6324 | 0.6368 | 0.6394 |
| Terminal ileum | 0.6145 | 0.6170 | 0.6188 | 0.6230 | 0.6256 |
| Testis | 0.6109 | 0.6129 | 0.6149 | 0.6193 | 0.6220 |
| Esophagus mucosa | 0.6068 | 0.6090 | 0.6107 | 0.6153 | 0.6180 |
| Sun-exposed skin | 0.6014 | 0.6037 | 0.6056 | 0.6101 | 0.6129 |
| Spleen | 0.5965 | 0.5988 | 0.6003 | 0.6056 | 0.6085 |
| Visceral omentum | 0.5875 | 0.5901 | 0.5912 | 0.5963 | 0.5989 |
| Prostate | 0.5738 | 0.5762 | 0.5788 | 0.5828 | 0.5856 |
| Stomach | 0.5632 | 0.5655 | 0.5677 | 0.5725 | 0.5753 |
| Adrenal gland | 0.5576 | 0.5605 | 0.5621 | 0.5673 | 0.5699 |
| Uterus | 0.5508 | 0.5534 | 0.5551 | 0.5602 | 0.5631 |
| Omnibus | 0.5475 | 0.5494 | 0.5516 | 0.5567 | 0.5597 |
| Esophagus muscularis | 0.5442 | 0.5469 | 0.5489 | 0.5535 | 0.5561 |
| Hippocampus | 0.5429 | 0.5450 | 0.5463 | 0.5516 | 0.5545 |
| Cortex | 0.5068 | 0.5083 | 0.5101 | 0.5164 | 0.5197 |
| Ovary | 0.4999 | 0.5028 | 0.5049 | 0.5103 | 0.5135 |
| Atrial appendage | 0.4976 | 0.5001 | 0.5017 | 0.5070 | 0.5099 |
| Transverse colon | 0.4930 | 0.4956 | 0.4981 | 0.5034 | 0.5067 |
| Caudate | 0.4914 | 0.4930 | 0.4946 | 0.5001 | 0.5033 |
| Tibial nerve | 0.4876 | 0.4900 | 0.4918 | 0.4971 | 0.5002 |
| Putamen | 0.4701 | 0.4721 | 0.4743 | 0.4793 | 0.4824 |
| CD4 cell | 0.4468 | 0.4487 | 0.4521 | 0.4578 | 0.4614 |
| Breast | 0.4426 | 0.4451 | 0.4477 | 0.4532 | 0.4564 |
| Sigmoid colon | 0.4397 | 0.4425 | 0.4446 | 0.4501 | 0.4533 |
| Cerebellum | 0.4366 | 0.4385 | 0.4405 | 0.4467 | 0.4503 |
| CD8 cell | 0.4198 | 0.4213 | 0.4249 | 0.4307 | 0.4345 |
| Subcutaneous adipose | 0.4170 | 0.4193 | 0.4219 | 0.4275 | 0.4309 |
| Gastroesophageal junction | 0.4150 | 0.4180 | 0.4202 | 0.4260 | 0.4294 |
| Skeletal muscle | 0.4116 | 0.4145 | 0.4169 | 0.4225 | 0.4258 |
| Monocyte | 0.4093 | 0.4111 | 0.4148 | 0.4202 | 0.4236 |
| Aorta | 0.3766 | 0.3786 | 0.3817 | 0.3869 | 0.3901 |
| Left ventricle | 0.3713 | 0.3742 | 0.3768 | 0.3826 | 0.3859 |
| B cell | 0.3669 | 0.3691 | 0.3730 | 0.3786 | 0.3822 |
| Liver | 0.3611 | 0.3640 | 0.3676 | 0.3730 | 0.3763 |
| NK cell | 0.3494 | 0.3508 | 0.3549 | 0.3605 | 0.3643 |
| Pancreas | 0.3241 | 0.3262 | 0.3300 | 0.3355 | 0.3390 |

#### Supplementary Table 15

Descriptive statistics of 38 regulatory networks. Rows are sorted by "# of TF-TG edges".

| Network | # of TF-TG edges | # of TGs | # of TFs |
|---|---|---|---|
| Stomach | 96135 | 3727 | 450 |
| Omnibus | 96008 | 9317 | 571 |
| Esophagus mucosa | 95797 | 4891 | 419 |
| Terminal ileum | 95596 | 3700 | 408 |
| Pancreas | 95446 | 2941 | 481 |
| Transverse colon | 95446 | 3696 | 435 |
| Lung | 95361 | 4123 | 415 |
| Breast | 95355 | 3260 | 442 |
| Left ventricle | 95030 | 2546 | 424 |
| Sun-exposed skin | 94988 | 4144 | 448 |
| Gastroesophageal junction | 94838 | 2380 | 443 |
| Atrial appendage | 94749 | 2564 | 399 |
| Vagina | 94592 | 4325 | 431 |
| Sigmoid colon | 94549 | 2545 | 460 |
| Esophagus muscularis | 94408 | 2993 | 429 |
| Visceral omentum | 94334 | 3701 | 428 |
| Tibial nerve | 94133 | 3214 | 431 |
| Subcutaneous adipose | 94005 | 2718 | 434 |
| Ovary | 93822 | 3659 | 430 |
| Liver | 93706 | 3398 | 412 |
| Spleen | 93494 | 4358 | 397 |
| Aorta | 93408 | 3295 | 414 |
| Uterus | 93057 | 3569 | 453 |
| Skeletal muscle | 92832 | 2308 | 490 |
| NK cell | 92399 | 3105 | 376 |
| Thyroid | 92263 | 5369 | 450 |
| CD8 cell | 91925 | 3439 | 424 |
| B cell | 91728 | 3018 | 436 |
| Prostate | 91722 | 3165 | 474 |
| CD4 cell | 91569 | 3643 | 433 |
| Monocyte | 91401 | 3702 | 384 |
| Testis | 90804 | 4646 | 474 |
| Cortex | 90579 | 2972 | 333 |
| Adrenal gland | 90401 | 2758 | 434 |
| Caudate | 89754 | 2547 | 374 |
| Cerebellum | 89371 | 3301 | 336 |
| Hippocampus | 88948 | 2592 | 366 |
| Putamen | 88634 | 2380 | 365 |

**Supplementary Table 16**

Descriptive statistics of TF-TG edge weights $\{v_{gt}\}$ over all TF-TG edges available in each of 38 regulatory networks. Rows are sorted by the "Median" column.

| | Summary of $v_{gt}$ | | | | |
|---|---|---|---|---|---|
| Network | Min | Q1 | Median | Q3 | Max |
| Cortex | 0.6946 | 0.7105 | 0.7315 | 0.7636 | 1.0000 |
| Putamen | 0.6797 | 0.6972 | 0.7201 | 0.7560 | 0.9944 |
| Cerebellum | 0.6817 | 0.6970 | 0.7170 | 0.7484 | 1.0000 |
| Caudate | 0.6704 | 0.6880 | 0.7113 | 0.7469 | 1.0000 |
| Hippocampus | 0.6678 | 0.6843 | 0.7063 | 0.7403 | 0.9977 |
| Atrial appendage | 0.6489 | 0.6674 | 0.6920 | 0.7314 | 1.0000 |
| CD8 cell | 0.6385 | 0.6553 | 0.6775 | 0.7125 | 1.0000 |
| Subcutaneous adipose | 0.6339 | 0.6520 | 0.6769 | 0.7160 | 1.0000 |
| Omnibus | 0.6278 | 0.6484 | 0.6748 | 0.7143 | 1.0000 |
| Transverse colon | 0.6353 | 0.6518 | 0.6740 | 0.7099 | 1.0000 |
| Terminal ileum | 0.6362 | 0.6522 | 0.6736 | 0.7082 | 1.0000 |
| Adrenal gland | 0.6296 | 0.6445 | 0.6650 | 0.6987 | 1.0000 |
| CD4 cell | 0.6266 | 0.6430 | 0.6647 | 0.6989 | 1.0000 |
| Gastroesophageal junction | 0.6233 | 0.6413 | 0.6645 | 0.7020 | 1.0000 |
| Stomach | 0.6276 | 0.6431 | 0.6640 | 0.6984 | 1.0000 |
| Skeletal muscle | 0.6187 | 0.6366 | 0.6609 | 0.6998 | 1.0000 |
| Lung | 0.6243 | 0.6393 | 0.6597 | 0.6927 | 1.0000 |
| NK cell | 0.6138 | 0.6324 | 0.6568 | 0.6949 | 1.0000 |
| B cell | 0.6101 | 0.6278 | 0.6515 | 0.6885 | 1.0000 |
| Breast | 0.6087 | 0.6263 | 0.6497 | 0.6870 | 1.0000 |
| Spleen | 0.6122 | 0.6279 | 0.6492 | 0.6831 | 1.0000 |
| Monocyte | 0.6088 | 0.6259 | 0.6488 | 0.6844 | 1.0000 |
| Sigmoid colon | 0.6079 | 0.6250 | 0.6482 | 0.6849 | 1.0000 |
| Prostate | 0.6047 | 0.6210 | 0.6427 | 0.6780 | 1.0000 |
| Left ventricle | 0.6006 | 0.6185 | 0.6418 | 0.6788 | 1.0000 |
| Esophagus muscularis | 0.5972 | 0.6157 | 0.6405 | 0.6797 | 1.0000 |
| Uterus | 0.5977 | 0.6142 | 0.6369 | 0.6728 | 1.0000 |
| Sun-exposed skin | 0.5984 | 0.6149 | 0.6368 | 0.6724 | 1.0000 |
| Ovary | 0.5938 | 0.6122 | 0.6368 | 0.6763 | 1.0000 |
| Tibial nerve | 0.5920 | 0.6104 | 0.6350 | 0.6740 | 1.0000 |
| Liver | 0.5867 | 0.6060 | 0.6312 | 0.6707 | 1.0000 |
| Pancreas | 0.5847 | 0.6046 | 0.6305 | 0.6722 | 1.0000 |
| Vagina | 0.5852 | 0.6010 | 0.6228 | 0.6581 | 1.0000 |
| Visceral omentum | 0.5814 | 0.5977 | 0.6198 | 0.6552 | 1.0000 |
| Aorta | 0.5646 | 0.5826 | 0.6067 | 0.6438 | 1.0000 |
| Esophagus mucosa | 0.5643 | 0.5809 | 0.6034 | 0.6400 | 1.0000 |
| Thyroid | 0.5621 | 0.5766 | 0.5966 | 0.6288 | 0.9587 |
| Testis | 0.5512 | 0.5675 | 0.5896 | 0.6252 | 1.0000 |

**Supplementary Table 17**

Descriptive statistics of *cis*-eQTL databases used in this study. Rows are sorted by the "# of SNP-gene pairs" column.

| Source of eQTL | # of SNP-gene pairs | # of genes | # of SNPs |
|---|---|---|---|
| Testis, GTEx (V7) | 15,759,533 | 17,366 | 1,230,746 |
| Prostate, GTEx (V7) | 14,585,826 | 16,085 | 1,219,002 |
| Terminal ileum, GTEx (V7) | 14,546,128 | 16,041 | 1,215,324 |
| Lung, GTEx (V7) | 14,483,585 | 15,973 | 1,214,950 |
| Sun-exposed skin, GTEx (V7) | 14,404,353 | 15,869 | 1,213,959 |
| Transverse colon, GTEx (V7) | 14,402,835 | 15,875 | 1,215,596 |
| Breast, GTEx (V7) | 14,392,706 | 15,872 | 1,212,389 |
| Vagina, GTEx (V7) | 14,380,972 | 15,869 | 1,217,316 |
| Thyroid, GTEx (V7) | 14,336,449 | 15,810 | 1,214,645 |
| Tibial nerve, GTEx (V7) | 14,288,824 | 15,761 | 1,217,709 |
| Cortex, GTEx (V7) | 14,261,947 | 15,745 | 1,226,087 |
| Hippocampus, GTEx (V7) | 14,233,460 | 15,719 | 1,224,980 |
| Stomach, GTEx (V7) | 14,220,507 | 15,684 | 1,213,739 |
| Caudate, GTEx (V7) | 14,209,653 | 15,689 | 1,226,945 |
| Sigmoid colon, GTEx (V7) | 14,202,126 | 15,668 | 1,219,231 |
| Cerebellum, GTEx (V7) | 14,192,621 | 15,679 | 1,224,520 |
| Visceral omentum, GTEx (V7) | 14,151,283 | 15,611 | 1,210,453 |
| Uterus, GTEx (V7) | 14,144,127 | 15,599 | 1,208,996 |
| Spleen, GTEx (V7) | 14,088,656 | 15,568 | 1,203,063 |
| Ovary, GTEx (V7) | 14,083,563 | 15,543 | 1,211,776 |
| Subcutaneous adipose, GTEx (V7) | 14,074,828 | 15,528 | 1,213,026 |
| Esophagus mucosa, GTEx (V7) | 14,050,266 | 15,504 | 1,201,919 |
| Putamen, GTEx (V7) | 14,036,948 | 15,503 | 1,226,334 |
| Gastroesophageal junction, GTEx (V7) | 13,998,654 | 15,446 | 1,217,148 |
| Esophagus muscularis, GTEx (V7) | 13,940,389 | 15,390 | 1,211,409 |
| Adrenal gland, GTEx (V7) | 13,918,735 | 15,368 | 1,215,183 |
| Aorta, GTEx (V7) | 13,913,807 | 15,356 | 1,207,409 |
| Pancreas, GTEx (V7) | 13,707,801 | 15,139 | 1,200,808 |
| Atrial appendage, GTEx (V7) | 13,683,429 | 15,122 | 1,207,966 |
| Liver, GTEx (V7) | 13,481,989 | 14,908 | 1,190,839 |
| Left ventricle, GTEx (V7) | 13,120,834 | 14,507 | 1,195,904 |
| Skeletal muscle, GTEx (V7) | 13,077,954 | 14,458 | 1,193,409 |
| CD8 cell, DICE (DB1) | 12,467,472 | 18,258 | 944,172 |
| CD4 cell, DICE (DB1) | 12,464,676 | 18,258 | 943,869 |
| NK cell, DICE (DB1) | 12,379,934 | 18,258 | 937,629 |
| Blood, eQTLGen | 12,372,754 | 13,588 | 1,173,301 |
| B cell, DICE (DB1) | 12,344,429 | 18,258 | 934,954 |
| Monocyte, DICE (DB1) | 12,344,429 | 18,258 | 934,954 |

**Supplementary Table 18**

Descriptive statistics of relative *cis* impact of SNPs on genes $\{c_{jg}\}$ over all SNP-gene pairs available in *cis*-eQTL databases. Rows are sorted by the "Median" column.

| | Summary of $(c_{jg} - 1)$ | | | | |
|---|---|---|---|---|---|
| Source of eQTL | Min | Q1 | Median | Q3 | Max |
| Uterus, GTEx (V7) | 0 | 0.033 | 0.069 | 0.118 | 0.935 |
| Vagina, GTEx (V7) | 0 | 0.032 | 0.067 | 0.115 | 0.932 |
| Putamen, GTEx (V7) | 0 | 0.032 | 0.067 | 0.114 | 0.923 |
| Hippocampus, GTEx (V7) | 0 | 0.031 | 0.066 | 0.114 | 0.912 |
| CD8 cell, DICE (DB1) | 0 | 0.024 | 0.065 | 0.118 | 0.774 |
| CD4 cell, DICE (DB1) | 0 | 0.023 | 0.064 | 0.118 | 0.771 |
| Monocyte, DICE (DB1) | 0 | 0.023 | 0.064 | 0.118 | 0.792 |
| Terminal ileum, GTEx (V7) | 0 | 0.030 | 0.064 | 0.109 | 0.935 |
| NK cell, DICE (DB1) | 0 | 0.023 | 0.064 | 0.117 | 0.762 |
| Ovary, GTEx (V7) | 0 | 0.030 | 0.063 | 0.108 | 0.931 |
| B cell, DICE (DB1) | 0 | 0.021 | 0.063 | 0.117 | 0.777 |
| Cortex, GTEx (V7) | 0 | 0.029 | 0.061 | 0.105 | 0.915 |
| Prostate, GTEx (V7) | 0 | 0.029 | 0.061 | 0.104 | 0.943 |
| Spleen, GTEx (V7) | 0 | 0.028 | 0.059 | 0.101 | 0.942 |
| Caudate, GTEx (V7) | 0 | 0.028 | 0.059 | 0.101 | 0.908 |
| Cerebellum, GTEx (V7) | 0 | 0.027 | 0.058 | 0.101 | 0.952 |
| Liver, GTEx (V7) | 0 | 0.027 | 0.057 | 0.098 | 0.921 |
| Adrenal gland, GTEx (V7) | 0 | 0.026 | 0.054 | 0.093 | 0.945 |
| Sigmoid colon, GTEx (V7) | 0 | 0.024 | 0.050 | 0.087 | 0.942 |
| Gastroesophageal junction, GTEx (V7) | 0 | 0.023 | 0.049 | 0.085 | 0.948 |
| Pancreas, GTEx (V7) | 0 | 0.023 | 0.049 | 0.084 | 0.939 |
| Testis, GTEx (V7) | 0 | 0.023 | 0.049 | 0.084 | 0.927 |
| Stomach, GTEx (V7) | 0 | 0.022 | 0.046 | 0.080 | 0.942 |
| Transverse colon, GTEx (V7) | 0 | 0.022 | 0.046 | 0.079 | 0.943 |
| Breast, GTEx (V7) | 0 | 0.021 | 0.045 | 0.078 | 0.949 |
| Atrial appendage, GTEx (V7) | 0 | 0.021 | 0.045 | 0.077 | 0.943 |
| Aorta, GTEx (V7) | 0 | 0.021 | 0.045 | 0.077 | 0.948 |
| Left ventricle, GTEx (V7) | 0 | 0.021 | 0.044 | 0.076 | 0.943 |
| Visceral omentum, GTEx (V7) | 0 | 0.019 | 0.041 | 0.071 | 0.939 |
| Esophagus muscularis, GTEx (V7) | 0 | 0.019 | 0.040 | 0.070 | 0.955 |
| Tibial nerve, GTEx (V7) | 0 | 0.019 | 0.040 | 0.069 | 0.964 |
| Esophagus mucosa, GTEx (V7) | 0 | 0.018 | 0.039 | 0.068 | 0.938 |
| Subcutaneous adipose, GTEx (V7) | 0 | 0.018 | 0.038 | 0.066 | 0.952 |
| Thyroid, GTEx (V7) | 0 | 0.018 | 0.038 | 0.065 | 0.955 |
| Lung, GTEx (V7) | 0 | 0.018 | 0.038 | 0.065 | 0.928 |
| Sun-exposed skin, GTEx (V7) | 0 | 0.017 | 0.037 | 0.064 | 0.948 |
| Skeletal muscle, GTEx (V7) | 0 | 0.016 | 0.034 | 0.058 | 0.918 |
| Blood, eQTLGen | 0 | 0.003 | 0.007 | 0.014 | 0.855 |

#### Supplementary Table 19

RSS-NET hyper-parameter grids used in the present study. For all traits, the grid for $\theta$ is $(0 : 0.25 : 1)$ and the grid for $\rho$ is $(0 : 0.2 : 0.8)$. Here $(j : i : k)$ denotes a regularly-spaced vector that starts at $j$, uses $i$ as the increment, and (roughly) stops at $k$. RSS-NET analyses of the same trait across different networks use the same hyper-parameter grid specified here. Trait abbreviations are defined in **Supplementary Table 2**.

| Trait | $\theta_0$ | $\eta$ |
|---|---|---|
| LOAD | $(-5.25:0.1:-4.75)$ | 0.6 |
| NEU | $(-4.4:0.05:-4)$ | 0.3 |
| SCZ | $(-2.175:0.025:-2.05)$ | 0.3 |
| BMI | $(-4.1:0.025:-3.95)$ | 0.3 |
| HEIGHT | $(-2.05:0.05:-1.95)$ | 0.3 |
| WAIST | $(-3:0.025:-2.95)$ | 0.3 |
| CD | $(-3:0.05:-2.9)$ | 0.3 |
| IBD | $(-3:0.05:-2.8)$ | 0.3 |
| RA | $(-3.25:0.025:-3.125)$ | 0.3 |
| UC | $(-3.05:0.025:-2.95)$ | 0.3 |
| BC | $(-4.25:0.05:-4)$ | 0.2 |
| AF | $(-4.5:0.25:-4)$ | 0.2, 0.3 |
| CAD | $(-4:0.025:-3.85)$ | 0.3 |
| HDL | $(-3.575:0.025:-3.45)$ | 0.3 |
| HR | $(-4.45:0.05:-4.25)$ | 0.3, 0.4 |
| LDL | $(-3.75:0.05:-3.5)$ | 0.3 |
| MI | $(-4.3:0.025:-4.1)$ | 0.3 |
| T2D | $(-4.65:0.05:-4.55)$ | 0.4, 0.5, 0.6 |

#### Supplementary Table 20

False positive rates of RSS-NET under different log 10 enrichment BF cutoffs across 47 simulation scenarios of negative datasets.

Panel **a** Descriptive statistics of RSS-NET false positive rates across 17 negative scenarios in **Supplementary Figures 1-2 & 9**.

| Cutoff | Min | Q1 | Median | Q3 | Max |
|---|---|---|---|---|---|
| 0 | 0.0000 | 0.0000 | 0.0200 | 0.1250 | 0.3800 |
| 1 | 0.0000 | 0.0000 | 0.0000 | 0.0250 | 0.1800 |
| 2 | 0.0000 | 0.0000 | 0.0000 | 0.0000 | 0.1150 |
| 3 | 0.0000 | 0.0000 | 0.0000 | 0.0000 | 0.0900 |
| 4 | 0.0000 | 0.0000 | 0.0000 | 0.0000 | 0.0600 |
| 5 | 0.0000 | 0.0000 | 0.0000 | 0.0000 | 0.0550 |
| 6 | 0.0000 | 0.0000 | 0.0000 | 0.0000 | 0.0450 |
| 7 | 0.0000 | 0.0000 | 0.0000 | 0.0000 | 0.0400 |
| 8 | 0.0000 | 0.0000 | 0.0000 | 0.0000 | 0.0400 |
| 9 | 0.0000 | 0.0000 | 0.0000 | 0.0000 | 0.0350 |

Panel **b** Descriptive statistics of RSS-NET false positive rates across 6 negative scenarios in **Supplementary Figure 3**.

| Cutoff | Min | Q1 | Median | Q3 | Max |
|---|---|---|---|---|---|
| 0 | 0.0150 | 0.0438 | 0.3300 | 0.6050 | 0.6300 |
| 1 | 0.0000 | 0.0000 | 0.0475 | 0.1362 | 0.2000 |
| 2 | 0.0000 | 0.0000 | 0.0075 | 0.0300 | 0.0750 |
| 3 | 0.0000 | 0.0000 | 0.0000 | 0.0037 | 0.0150 |
| 4 | 0.0000 | 0.0000 | 0.0000 | 0.0000 | 0.0050 |
| 5 | 0.0000 | 0.0000 | 0.0000 | 0.0000 | 0.0050 |
| 6 | 0.0000 | 0.0000 | 0.0000 | 0.0000 | 0.0000 |
| 7 | 0.0000 | 0.0000 | 0.0000 | 0.0000 | 0.0000 |
| 8 | 0.0000 | 0.0000 | 0.0000 | 0.0000 | 0.0000 |
| 9 | 0.0000 | 0.0000 | 0.0000 | 0.0000 | 0.0000 |

Panel **c** Descriptive statistics of RSS-NET false positive rates across 6 negative scenarios in **Supplementary Figure 4**.

| Cutoff | Min | Q1 | Median | Q3 | Max |
|---|---|---|---|---|---|
| 0 | 0.0200 | 0.0563 | 0.4225 | 0.8337 | 0.8700 |
| 1 | 0.0000 | 0.0000 | 0.1450 | 0.3013 | 0.4200 |
| 2 | 0.0000 | 0.0000 | 0.0100 | 0.0650 | 0.1400 |
| 3 | 0.0000 | 0.0000 | 0.0025 | 0.0125 | 0.0200 |
| 4 | 0.0000 | 0.0000 | 0.0000 | 0.0000 | 0.0050 |
| 5 | 0.0000 | 0.0000 | 0.0000 | 0.0000 | 0.0000 |
| 6 | 0.0000 | 0.0000 | 0.0000 | 0.0000 | 0.0000 |
| 7 | 0.0000 | 0.0000 | 0.0000 | 0.0000 | 0.0000 |
| 8 | 0.0000 | 0.0000 | 0.0000 | 0.0000 | 0.0000 |
| 9 | 0.0000 | 0.0000 | 0.0000 | 0.0000 | 0.0000 |

Panel **d** Descriptive statistics of RSS-NET false positive rates across 6 negative scenarios in **Supplementary Figure 5**.

| Cutoff | Min | Q1 | Median | Q3 | Max |
|---|---|---|---|---|---|
| 0 | 0.0150 | 0.0287 | 0.0950 | 0.1462 | 0.2350 |
| 1 | 0.0000 | 0.0000 | 0.0125 | 0.0287 | 0.0350 |
| 2 | 0.0000 | 0.0000 | 0.0000 | 0.0000 | 0.0050 |
| 3 | 0.0000 | 0.0000 | 0.0000 | 0.0000 | 0.0000 |
| 4 | 0.0000 | 0.0000 | 0.0000 | 0.0000 | 0.0000 |
| 5 | 0.0000 | 0.0000 | 0.0000 | 0.0000 | 0.0000 |
| 6 | 0.0000 | 0.0000 | 0.0000 | 0.0000 | 0.0000 |
| 7 | 0.0000 | 0.0000 | 0.0000 | 0.0000 | 0.0000 |
| 8 | 0.0000 | 0.0000 | 0.0000 | 0.0000 | 0.0000 |
| 9 | 0.0000 | 0.0000 | 0.0000 | 0.0000 | 0.0000 |

Panel **e** Descriptive statistics of RSS-NET false positive rates across 12 negative scenarios in **Supplementary Figure 6**.

| Cutoff | Min | Q1 | Median | Q3 | Max |
|---|---|---|---|---|---|
| 0 | 0.2100 | 0.4625 | 0.8075 | 0.9950 | 1.0000 |
| 1 | 0.0400 | 0.0900 | 0.5675 | 0.9413 | 1.0000 |
| 2 | 0.0050 | 0.0138 | 0.3825 | 0.8712 | 1.0000 |
| 3 | 0.0000 | 0.0050 | 0.2675 | 0.7650 | 0.9900 |
| 4 | 0.0000 | 0.0050 | 0.2050 | 0.6262 | 0.9800 |
| 5 | 0.0000 | 0.0050 | 0.1425 | 0.4888 | 0.9700 |
| 6 | 0.0000 | 0.0050 | 0.0775 | 0.3700 | 0.9550 |
| 7 | 0.0000 | 0.0000 | 0.0400 | 0.2500 | 0.9250 |
| 8 | 0.0000 | 0.0000 | 0.0275 | 0.1675 | 0.8900 |
| 9 | 0.0000 | 0.0000 | 0.0225 | 0.1038 | 0.8550 |

Panel **f** Descriptive statistics of RSS-NET false positive rates across 47 negative scenarios from Panels **a** to **e** above.

| Cutoff | Min | Q1 | Median | Q3 | Max |
|---|---|---|---|---|---|
| 0 | 0.0000 | 0.0200 | 0.1350 | 0.6150 | 1.0000 |
| 1 | 0.0000 | 0.0000 | 0.0300 | 0.1900 | 1.0000 |
| 2 | 0.0000 | 0.0000 | 0.0000 | 0.0750 | 1.0000 |
| 3 | 0.0000 | 0.0000 | 0.0000 | 0.0150 | 0.9900 |
| 4 | 0.0000 | 0.0000 | 0.0000 | 0.0050 | 0.9800 |
| 5 | 0.0000 | 0.0000 | 0.0000 | 0.0050 | 0.9700 |
| 6 | 0.0000 | 0.0000 | 0.0000 | 0.0025 | 0.9550 |
| 7 | 0.0000 | 0.0000 | 0.0000 | 0.0000 | 0.9250 |
| 8 | 0.0000 | 0.0000 | 0.0000 | 0.0000 | 0.8900 |
| 9 | 0.0000 | 0.0000 | 0.0000 | 0.0000 | 0.8550 |

#### Supplementary Table 21

False positive rates of RSS-NET under different $P_1$ cutoffs across 49 simulation scenarios in **Supplementary Figures 7-9**.

Panel **a** Descriptive statistics of RSS-NET false positive rates across 8 scenarios with $\theta = 0$ and $\sigma^2 = 0$.

| Cutoff | Min | Q1 | Median | Q3 | Max |
|---|---|---|---|---|---|
| 0.50 | 1.9158e-05 | 4.3152e-05 | 1.3280e-04 | 3.1445e-02 | 6.6337e-02 |
| 0.55 | 1.9158e-06 | 3.9181e-05 | 1.1041e-04 | 1.7910e-02 | 3.7976e-02 |
| 0.60 | 9.5791e-07 | 3.7196e-05 | 9.1061e-05 | 1.0309e-02 | 2.1019e-02 |
| 0.65 | 7.1843e-07 | 3.2524e-05 | 8.2365e-05 | 5.7166e-03 | 1.1068e-02 |
| 0.70 | 7.1843e-07 | 2.8846e-05 | 7.1198e-05 | 3.0852e-03 | 5.6593e-03 |
| 0.75 | 0.0000e+00 | 2.5692e-05 | 6.2483e-05 | 1.5269e-03 | 2.9909e-03 |
| 0.80 | 0.0000e+00 | 2.3357e-05 | 5.5691e-05 | 6.9265e-04 | 1.4404e-03 |
| 0.85 | 0.0000e+00 | 1.9970e-05 | 5.2003e-05 | 2.7399e-04 | 6.2126e-04 |
| 0.90 | 0.0000e+00 | 1.7868e-05 | 4.4425e-05 | 8.9064e-05 | 1.8040e-04 |
| 0.95 | 0.0000e+00 | 1.4094e-05 | 1.5193e-05 | 2.1983e-05 | 6.0961e-05 |

Panel **b** Descriptive statistics of RSS-NET false positive rates across 16 scenarios with $\theta = 0$ and $\sigma^2 > 0$.

| Cutoff | Min | Q1 | Median | Q3 | Max |
|---|---|---|---|---|---|
| 0.50 | 3.4568e-05 | 7.9590e-05 | 1.8260e-04 | 3.9777e-02 | 1.3713e-01 |
| 0.55 | 3.1765e-05 | 7.3369e-05 | 1.3957e-04 | 2.4210e-02 | 9.0496e-02 |
| 0.60 | 2.8028e-05 | 6.3212e-05 | 1.3148e-04 | 1.4952e-02 | 5.6791e-02 |
| 0.65 | 2.5926e-05 | 5.2579e-05 | 1.2136e-04 | 8.2891e-03 | 3.3777e-02 |
| 0.70 | 2.5225e-05 | 4.7513e-05 | 1.1207e-04 | 4.4175e-03 | 1.8742e-02 |
| 0.75 | 2.4525e-05 | 4.1115e-05 | 1.0169e-04 | 2.1514e-03 | 9.8169e-03 |
| 0.80 | 2.1722e-05 | 3.7478e-05 | 9.0797e-05 | 9.1384e-04 | 4.7398e-03 |
| 0.85 | 2.1255e-05 | 3.2583e-05 | 7.9758e-05 | 3.8602e-04 | 2.0535e-03 |
| 0.90 | 1.9620e-05 | 2.8757e-05 | 7.1112e-05 | 1.4587e-04 | 6.6205e-04 |
| 0.95 | 1.7518e-05 | 2.1299e-05 | 3.8781e-05 | 8.3458e-05 | 1.1029e-04 |

Panel **c** Descriptive statistics of RSS-NET false positive rates across 8 scenarios with $\theta > 0$ and $\sigma^2 = 0$.

| Cutoff | Min | Q1 | Median | Q3 | Max |
|---|---|---|---|---|---|
| 0.50 | 3.1391e-04 | 2.7308e-03 | 8.4924e-02 | 1.0921e-01 | 1.6779e-01 |
| 0.55 | 1.4283e-04 | 1.3149e-03 | 6.0600e-02 | 8.4700e-02 | 1.3078e-01 |
| 0.60 | 5.7572e-05 | 6.2062e-04 | 4.6203e-02 | 6.5058e-02 | 9.8028e-02 |
| 0.65 | 2.6867e-05 | 2.9597e-04 | 3.4367e-02 | 4.9285e-02 | 6.9542e-02 |
| 0.70 | 1.5078e-05 | 1.3498e-04 | 2.4858e-02 | 3.5227e-02 | 4.6563e-02 |
| 0.75 | 8.2246e-06 | 5.4906e-05 | 1.7402e-02 | 2.3246e-02 | 2.8511e-02 |
| 0.80 | 3.0157e-06 | 2.1513e-05 | 1.1155e-02 | 1.3445e-02 | 1.6516e-02 |
| 0.85 | 5.4831e-07 | 5.1493e-06 | 6.1871e-03 | 6.8245e-03 | 1.0139e-02 |
| 0.90 | 0.0000e+00 | 1.0088e-06 | 2.1403e-03 | 3.0779e-03 | 4.6428e-03 |
| 0.95 | 0.0000e+00 | 2.8646e-07 | 2.5865e-04 | 8.9441e-04 | 1.1598e-03 |

Panel **d** Descriptive statistics of RSS-NET false positive rates across 17 scenarios with $\theta > 0$ and $\sigma^2 > 0$.

| Cutoff | Min | Q1 | Median | Q3 | Max |
|---|---|---|---|---|---|
| 0.50 | 3.6369e-03 | 1.4967e-02 | 9.4407e-02 | 1.1433e-01 | 2.8226e-01 |
| 0.55 | 1.8042e-03 | 9.9008e-03 | 7.1041e-02 | 9.6921e-02 | 2.4531e-01 |
| 0.60 | 8.8415e-04 | 6.4193e-03 | 5.1986e-02 | 8.7748e-02 | 2.0722e-01 |
| 0.65 | 4.2384e-04 | 3.9577e-03 | 4.0081e-02 | 6.6512e-02 | 1.6879e-01 |
| 0.70 | 2.0918e-04 | 2.2908e-03 | 3.0449e-02 | 5.0644e-02 | 1.3095e-01 |
| 0.75 | 1.1240e-04 | 1.1916e-03 | 2.1810e-02 | 3.5411e-02 | 9.5022e-02 |
| 0.80 | 6.4700e-05 | 5.0790e-04 | 1.4876e-02 | 2.5534e-02 | 6.2574e-02 |
| 0.85 | 3.1671e-05 | 1.8625e-04 | 8.6225e-03 | 1.2890e-02 | 3.6571e-02 |
| 0.90 | 1.7724e-05 | 9.7325e-05 | 4.3930e-03 | 6.0918e-03 | 2.0671e-02 |
| 0.95 | 1.1623e-05 | 4.5801e-05 | 7.0200e-04 | 1.6252e-03 | 8.4746e-03 |

Panel **e** Descriptive statistics of RSS-NET false positive rates across 49 scenarios from Panels **a** to **d** above.

| Cutoff | Min | Q1 | Median | Q3 | Max |
|---|---|---|---|---|---|
| 0.50 | 1.9158e-05 | 1.5303e-04 | 2.3029e-02 | 9.9199e-02 | 2.8226e-01 |
| 0.55 | 1.9158e-06 | 1.1734e-04 | 1.3126e-02 | 7.1041e-02 | 2.4531e-01 |
| 0.60 | 9.5791e-07 | 8.9924e-05 | 7.9050e-03 | 5.1986e-02 | 2.0722e-01 |
| 0.65 | 7.1843e-07 | 7.9649e-05 | 4.4754e-03 | 3.7725e-02 | 1.6879e-01 |
| 0.70 | 7.1843e-07 | 7.3107e-05 | 2.5532e-03 | 2.8071e-02 | 1.3095e-01 |
| 0.75 | 0.0000e+00 | 6.8903e-05 | 1.3102e-03 | 2.0162e-02 | 9.5022e-02 |
| 0.80 | 0.0000e+00 | 4.7177e-05 | 6.6729e-04 | 1.2758e-02 | 6.2574e-02 |
| 0.85 | 0.0000e+00 | 3.4724e-05 | 3.1254e-04 | 6.7839e-03 | 3.6571e-02 |
| 0.90 | 0.0000e+00 | 2.6867e-05 | 1.2389e-04 | 2.6863e-03 | 2.0671e-02 |
| 0.95 | 0.0000e+00 | 1.8642e-05 | 4.5801e-05 | 3.8126e-04 | 8.4746e-03 |
